# Transmural Single-Nucleus Transcriptomics Nominates Candidate Disease Programs in Human Necrotizing Enterocolitis

**DOI:** 10.64898/2026.10.29.762618

**Authors:** Mickael F. Meyer, Caden W. Sweet, Keith Breau, Joseph Burclaff, R. Jarrett Bliton, Hailey M. Dodson, Gabrielle Cannon, Pauline Bougaran, Vikram Puri, Lauren C. Frazer, Lauren R. Bolzan, Catherine R. Rizzuto, Danielle E. Sklar, Corey M. Jania, Gergely D. Mozes, Amy B. Hair, Camilia R. Martin, Troy A. Markel, Brian Scottoline, Natalia S. Akopyants, Jennifer Canvasser, Neil McCarthy, Deepika Sharma, Victoria Bautch, Laurianne Van Landeghem, Misty Good, Scott T. Magness

## Abstract

**Background & Aims:** Necrotizing enterocolitis (NEC) is a life-threatening disease of premature infants marked by inflammation, ischemia, mucosal injury, and, in advanced cases, necrosis extending into deeper layers of the intestinal wall. Because NEC affects specialized, interacting cells in a developing intestine, we used single-nucleus transcriptomics to build a transmural, cell-type-resolved map and nominate hypothesis-generating disease programs distinct from maturation.

**Methods:** Full-thickness intestinal specimens from fetal/prenatal controls, acute NEC cases, and infant controls were analyzed by single-nucleus RNA sequencing (snRNA-seq). Fetal and infant samples served as developmental references. Analyses included cell annotation, NEC-versus-infant differential expression, gene ontology (GO), cross-lineage recurrence, cell death signatures, and developmental maturity.

**Results:** We profiled 53,735 nuclei by snRNA-seq and resolved 34 epithelial, stromal, vascular, immune, neuronal, glial, and smooth muscle populations. We used a morphology-based nuclei scoring system for sample selection that predicted high-quality transcript recovery. NEC nuclei were represented across this cellular landscape, with recurrent gene-expression changes suggesting shared inflammatory recruitment, antimicrobial/redox defense, stress adaptation, complement/vascular activation, reduced homeostatic signaling, and cell death-associated regulation. Developmental maturity analysis identified fetal-like expression levels in subsets of NEC-regulated genes. Proliferative crypt populations exhibit DNA repair/cell-cycle programs with inflammatory, hypoxic, and ER-stress features, while absorptive enterocytes showed reduced digestive, transport, detoxification, and terminal villus-function programs. Secretory-lineage changes implicated goblet mucus-barrier responses with reduced glycosylation, trafficking, and hydration; enteroendocrine changes in epithelial defense, metabolic stress, and neural signaling; and Paneth cell retention of antimicrobial effector transcripts despite altered ER quality control, autophagy, and granule delivery in NEC. Non-epithelial programs implicated stromal Wnt/BMP niche loss, impaired lymphatic drainage/scavenging, macrophage cytokine activation with reduced clearance machinery, neuroglial circuit defects, and altered smooth muscle contraction-relaxation balance.

**Conclusions:** This single nuclear intestinal cell atlas nominates multicellular programs that may govern NEC initiation, progression toward tissue destruction, and resolution toward functional recovery.

## Introduction

### Clinical Burden and Biological Complexity of NEC

Necrotizing enterocolitis (NEC) is a life-threatening inflammatory disease of the neonatal intestine characterized by mucosal injury, ischemia, and, in severe cases, transmural necrosis.^1–4^ It occurs predominantly in premature and low birth weight infants, with incidence increasing sharply as birth weight and gestational age decline.^5–7^ NEC affects approximately 1 in 1,000 live births overall.^1^ In one large extremely preterm cohort (gestational age 22-28 weeks), NEC occurred in 11% of infants.^7^ Recent systematic analyses estimate an overall mortality of approximately 23.5%, with fatality exceeding 50% among extremely low birth weight (birth weight less than 1000 grams) infants requiring surgical intervention.^8^ Among survivors, particularly those with surgical NEC, long-term morbidity remains substantial, including neurodevelopmental impairment and intestinal failure.^9–10^ These outcomes underscore the need to understand not only the mechanisms that initiate intestinal injury, but also the multicellular processes that amplify tissue destruction and determine whether the intestine progresses toward recovery or irreversible failure.

NEC arises in a developmentally immature intestine that is vulnerable to microbial dysbiosis, exaggerated innate immune activation, epithelial-barrier injury, vascular/metabolic stress, and failed mucosal repair. Conversely, breastmilk feeding, partly through maternal milk-derived IgA coating of gut bacteria, restriction of IgA-unbound *Enterobacteriaceae*, and the inhibition of TLR4 signaling is associated with reduced NEC risk.^2–4,11–13^ Prospective human microbiome studies indicate that gut bacterial dysbiosis can precede NEC, including enrichment of Gammaproteobacteria and reduced strict anaerobes in very low birth weight infants before NEC onset.^14^ Broader NEC microbiome literature supports links among dysbiosis, barrier disruption, inflammation, and epithelial injury.^15^ In experimental mouse NEC, epithelial Toll-like receptor 4 (TLR4) signaling has been linked to microbial sensing, impaired intestinal stem cell proliferation, crypt apoptosis, defective mucosal regeneration, and endoplasmic reticulum (ER)-stress-associated stem cell apoptosis.^16–17^ Human single-cell transcriptomic data similarly show broad epithelial inflammatory activation and depletion of differentiated villus epithelial populations in NEC tissue.^18^ Together, these findings support a model in which immature intestinal cells interact with microbial, nutritional, and immune-modulatory stressors, but the cellular programs that determine disease onset, propagation, and recovery remain incompletely defined.

### NEC as a Multicellular Disease Beyond Epithelium and Immunity

Many mechanistic frameworks for NEC have emphasized epithelial immaturity, microbial sensing, and immune activation.^2–3^ However, the intestine is a structurally integrated organ in which epithelial, stromal, endothelial, immune, neuronal, glial, and smooth muscle populations exchange trophic, inflammatory, metabolic, mechanical, and reparative signals. For example, enteric glia regulate epithelial barrier function, modulate epithelial proliferation and differentiation through paracrine mediators and contribute to gastrointestinal motility regulation.^19–24^ Stromal and extracellular matrix-producing populations provide niche-associated, positional, and trophic signals required for epithelial homeostasis and regeneration.^25–29^ Lymphatic endothelial cells can also provide repair-associated lymphangiocrine signals during intestinal injury responses.^30^ By contrast, lymphocyte populations may have context-dependent effects in NEC, with regulatory T-cell programs potentially limiting inflammation and tissue injury, while TLR4-associated lymphocyte influx and IL-17-producing T-cell responses can amplify epithelial injury.^31–32^ Because severe NEC can extend from the mucosa through the submucosa, muscularis, and serosa, the disease cannot be fully understood through analysis of mucosal epithelial or immune populations alone.^1–2^ It remains unclear which transcriptional responses are broadly shared across intestinal lineages, which are specific to individual cell populations, and which may contribute to NEC pathological mechanisms of initiation, progression toward tissue destruction, or resolution.

Single-cell RNA sequencing has begun to reveal important features of the human NEC intestine. Human NEC tissue contains increased inflammatory CD16+CD163+ monocytes and extensive immune activation detected by single-cell RNA sequencing and imaging mass cytometry.^33^ Other studies reported altered intestinal immune cell states in NEC, including T-cell TNF-α and mTOR pathway enrichment.^34^ Human NEC single-cell data also show loss of villus-tip epithelial populations and inflammatory activation of epithelial, endothelial, fibroblast, and immune compartments.^18^ Computational analyses have implicated macrophage pyroptosis-associated programs in NEC, although transcriptomic associations alone do not establish execution of pyroptotic cell death.^35^ Collectively, these studies establish immune-epithelial dysfunction as a central feature of NEC, but provide more limited resolution of epithelial progenitors, differentiated epithelial subtypes, stromal and vascular populations, enteric neural and glial populations, and smooth muscle.

### Study Design

A major obstacle to comprehensive cellular analysis is the nature of NEC tissue itself. Surgical specimens are scarce, clinically heterogeneous, and often obtained during nights or weekends when research personnel may not be available to immediately process fresh tissue, necessitating cryopreservation for subsequent analysis.^18,36–37^ These challenges are amplified for conventional single-cell RNA sequencing, where tissue dissociation and recovery of intact viable cells can induce stress-associated transcriptional changes, reduce cell viability, and distort cell-type representation through the loss of fragile or difficult-to-dissociate populations.^38–39^ These limitations are especially important when trying to generate a representative transcriptomic profile across all layers of the intestinal wall.

Single-nucleus RNA sequencing (snRNA-seq) provides a practical alternative for frozen tissue by using nuclei as the transcriptomic unit of the cell, allowing cellular states to be inferred without recovery of intact viable cells.^40^ Single-nucleus and whole-cell RNA sequencing capture partially different transcript pools and can differ in cell-type representation. Nuclear profiling is enriched for nuclear and nascent RNA features, but it enables broader profiling of rare, archived NEC surgical specimens while reducing dependence on dissociation-sensitive viable-cell workflows.^41^ This approach is particularly useful for profiling full-thickness intestinal specimens, as it allows characterization of diverse cell populations throughout the intestinal wall, including epithelial, endothelial, neuronal, glial, stromal, and smooth muscle cells.^39–40^

Given these advantages, our study used single-nucleus RNA sequencing of flash-frozen, full-thickness intestinal specimens obtained through the NEC Biorepository, a multicenter resource that includes samples contributed from multiple medical centers across the United States. Nuclei-based profiling avoided the requirement for viable whole-cell recovery, allowed nuclear quality to be assessed before library preparation, and enabled donor samples to be pooled for split-pool combinatorial barcoding and high-throughput sequencing. We analyzed prenatal intestine (fetal controls), surgical NEC specimens, and infant control tissue spanning a range of ages during prenatal and postnatal development. The fetal and infant control groups served as developmental references, allowing us to evaluate NEC-associated transcriptional changes in the context of gene-expression changes that normally occur as the neonatal intestine matures.

### Analytical Strategy

This study design enabled transcriptomic profiling of epithelial, stromal, endothelial, immune, neuronal, glial, and smooth muscle cell populations from transmural intestinal sections. We first identify and annotate cell populations to establish the anatomical and cellular composition of the dataset, then use this organization to examine NEC-associated biology within individual populations and across the intestine. To orient the reader, we provide an analytical primer that defines how we interpret differential expression, recurrent gene regulation, GO term analysis, curated gene programs, and maturity analyses throughout the manuscript. This primer also explains how fetal controls, NEC samples, and infant controls are used together to distinguish NEC-associated regulation from normal developmental maturation.

The first analytical layer asks whether NEC produces shared transcriptional responses across the intestine. We identify genes recurrently upregulated or downregulated across otherwise distinct cell types, allowing broad tissue-level injury programs to be distinguished from population-restricted responses. These shared programs provide a global view of NEC biology before the manuscript turns to individual lineages, and help identify recurring themes such as innate immune activation, epithelial defense, inflammatory recruitment, vascular and stromal activation, and stress adaptation.

We next apply a similar cross-cell-type strategy to regulated cell death pathways. Apoptosis, necroptosis, pyroptosis, and ferroptosis can intersect during inflammatory injury, but their execution depends on mechanisms such as caspase activation, RIPK3/MLKL signaling, gasdermin-mediated membrane disruption, and iron-dependent lipid peroxidation.^42–45^ Because pathway execution is largely post-transcriptional, RNA changes are interpreted as evidence of death-related priming, vulnerability, restraint, or protective buffering and do not establish that a death pathway has occurred.^46^

The manuscript then moves from global analyses into a systematic examination of each major intestinal compartment and its constituent cell types. Like a world atlas, the manuscript is organized into self-contained cell-type entries that readers can consult independently while using the same interpretive framework throughout. Each cell-type section follows a repeated structure that begins with normal anatomy and NEC relevance, then integrates the strongest NEC-associated differentially expressed genes (DEGs), GO-terms when supported, curated gene panels, and developmental maturity patterns. Gene-level maturity analysis places each population along the fetal-to-infant developmental axis, while gene-level and transcription factor analyses identify genes and regulatory programs that mature normally, remain fetal-like, or show errant disease-associated expression. This structure supports comparison across cell types while keeping each interpretation grounded in the population’s location, function, and potential contribution to NEC initiation, progression, resolution, or impaired recovery.

### A Hypothesis-Generating Framework for NEC Initiation, Progression, and Resolution

Surgical NEC tissue creates an important interpretive constraint. A resected intestinal segment captures severe disease at the time of surgery, but it cannot determine when each transcriptional program began, whether that program helped drive injury or arose in response to it, or whether the sampled tissue came from fully involved regions, less involved margins, or both. We therefore discuss how transcriptomic programs may contribute to NEC initiation, progression, and resolution as a hypothesis-generating framework, not as a precise timeline of disease events or an assumption that all sampled regions are equivalent. For example, microbial sensing, inflammatory recruitment, epithelial vulnerability, and stress susceptibility may relate to initiation or early amplification; sustained inflammatory, metabolic, vascular, and death-associated programs may relate to progression toward tissue destruction; and proliferation, DNA repair, chromatin regulation, matrix remodeling, trophic signaling, and lineage differentiation may reflect attempts at resolution, including repair responses that are incomplete, overwhelmed, or insufficient for recovery.

Using this framework, we provide a deeply resolved analysis of human NEC across the full intestinal wall and identify shared and cell-type-specific programs that may shape disease behavior. Rather than reducing NEC to a single inflammatory pathway or dominant cell population, this study presents NEC as a multicellular breakdown of intestinal homeostasis, in which injury sensing, inflammatory amplification, cellular stress, death susceptibility, developmental state, and regenerative capacity are distributed unevenly across the tissue. This framework is intended to make the resource navigable and to generate testable hypotheses about why some premature intestines are susceptible to injury, while others progress toward transmural destruction or fail to exhibit epithelial restitution, cellular proliferation, and mucosal regeneration.

## Results

### Developmentally Stratified Intestinal Samples Establish a Framework for NEC-Specific Analysis

A central goal of this study was to distinguish transcriptional programs associated with NEC. To accomplish this, we assembled fetal control, surgical NEC, and infant control specimens spanning a developmental continuum from fetal preterm intestine through the gestational period represented by the earliest NEC cases and into preterm, early-term, and full-term infant development (Fig. 1A). The cohort comprised fetal controls (n = 14; 17-22 weeks gestation), infants with surgical NEC (n = 8; gestational age at birth between 23 weeks + 1 day and 27 weeks + 2 days), and infant controls (n = 8; gestational age at birth between 22 and 39 weeks). Among specimens with available collection ages, NEC tissues were obtained between 26 weeks + 5 days and 30 weeks postmenstrual age (PMA), whereas infant control tissues were obtained between 27 weeks + 6 days and 47 weeks postmenstrual age (See Methods and Fig. 39). This design provided age-relevant NEC-versus-infant comparisons while establishing a broader developmental reference for determining whether gene expression changes in NEC were due to disease or immaturity.

**Fig. 1.**
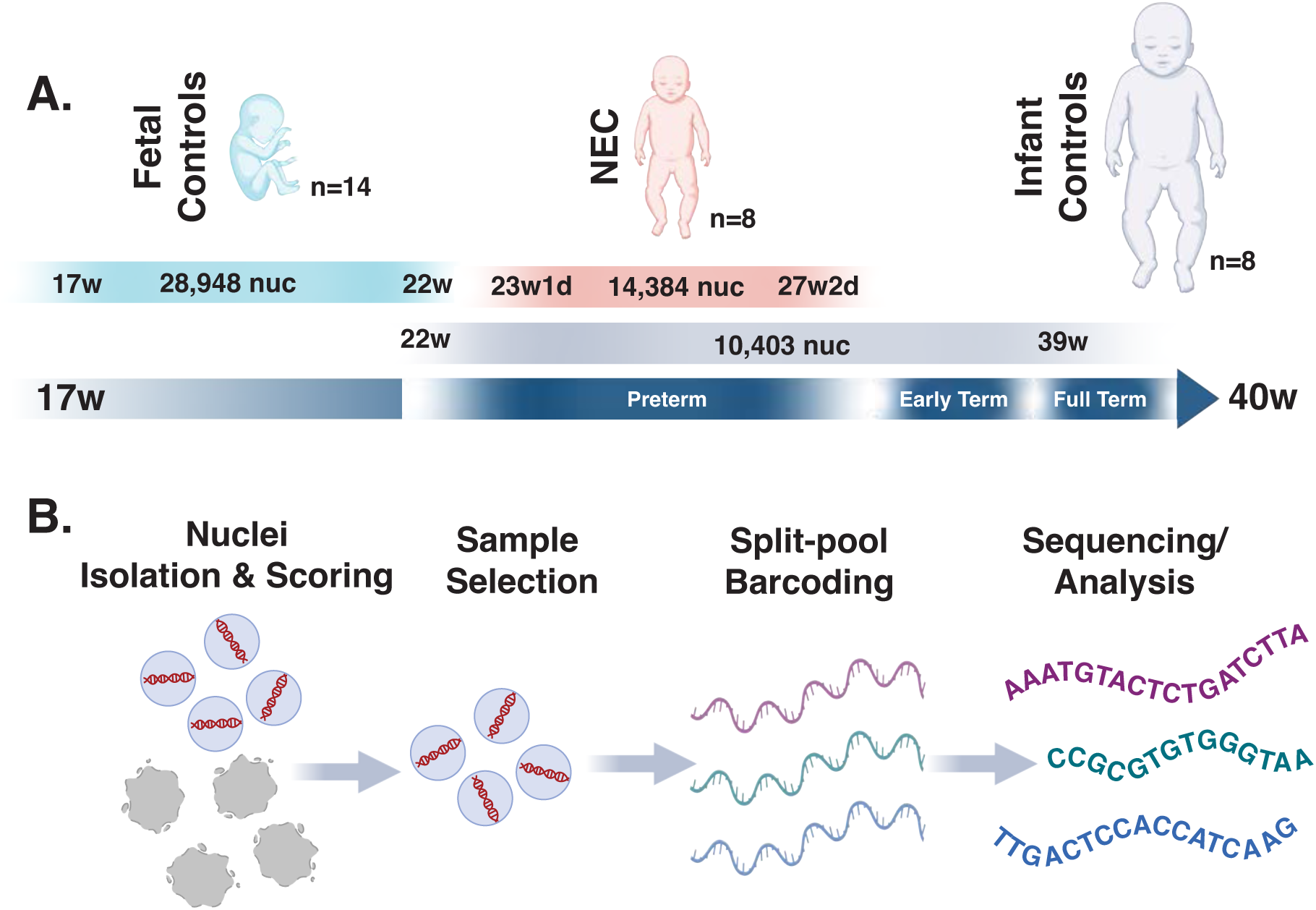
Study cohort and single-nucleus RNA-sequencing workflow. **A.** Human intestinal specimens included fetal controls (light blue), surgical NEC samples (red), and infant controls (blue-gray) spanning fetal, preterm, early-term, and full-term developmental ages. The dark blue timeline indicates the fetal-to-term developmental window. Fetal controls ranged from 17–22 weeks gestation. NEC samples came from infants born at 23 weeks, 1 day to 27 weeks, 2 days gestational age, with tissue collected at 26 weeks, 5 days to 30 weeks postmenstrual age when available. Infant controls came from infants born at 22–39 weeks gestational age, with tissue collected at 27 weeks, 6 days to 47 weeks postmenstrual age when available. Ages shown on the timeline for NEC and infant control groups indicate gestational age at birth, not age at tissue collection; fetal control ages indicate gestational age at collection. The cohort comprised 14 fetal controls with 28,948 nuclei, 8 NEC samples with 14,384 nuclei, and 8 infant controls with 10,403 nuclei. **B.** Flash-frozen full-thickness intestinal tissue was first processed for nuclei isolation and quality scoring. Samples with nuclei meeting the predefined quality threshold (see methods; blue) were selected for split-pool combinatorial barcoding and high-throughput sequencing/analysis, whereas those with nuclei below the threshold (gray) were not advanced for sequencing.

To capture the multicellular consequences of transmural NEC injury, we analyzed full-thickness intestinal specimens, enabling recovery of transcriptomic signatures from epithelial, stromal, vascular, immune, neuronal, glial, and muscular populations. To reduce the risk of sequencing poorly preserved preparations, we developed a morphology-based nuclei scoring system (see Methods; Fig. 40). Preparations from all three conditions were evaluated using this framework before selected samples underwent split-pool combinatorial barcoding, sequencing, and computational analysis (Fig. 1B). Following quality filtering, 28,948 fetal control nuclei, 14,384 NEC nuclei, and 10,403 infant control nuclei were retained, yielding 53,735 nuclei for downstream analysis (see Methods; Figs. 39,40). Together, the developmental breadth and age overlap of the cohort, transmural sampling, and quality-guided specimen selection established the foundation for identifying NEC-specific transcriptional programs while accounting for normal developmental variation.

### Single-Nucleus Analysis Resolves 34 Cell Populations Across Fetal, NEC, and Infant Intestinal Samples

Having established a developmentally stratified and quality controlled cohort, we combined data from fetal control, NEC, and infant control specimens to cluster nuclei and generate heatmaps of differentially expressed genes (DEGs) for cell-population identification. Analysis of all 53,735 nuclei produced a global transcriptional map containing six distinct tissue compartments: epithelial, stromal, endothelial, immune, neural, and smooth muscle cell populations (Fig. 2A). Tissue-level differential gene-expression analysis showed that each major intestinal tissue class retained a distinct molecular signature (Fig. 2B), supporting the anatomical and cellular organization used for annotation. Subsequent heatmaps were likewise generated from the combined fetal control, NEC, and infant control dataset.

**Fig. 2.**
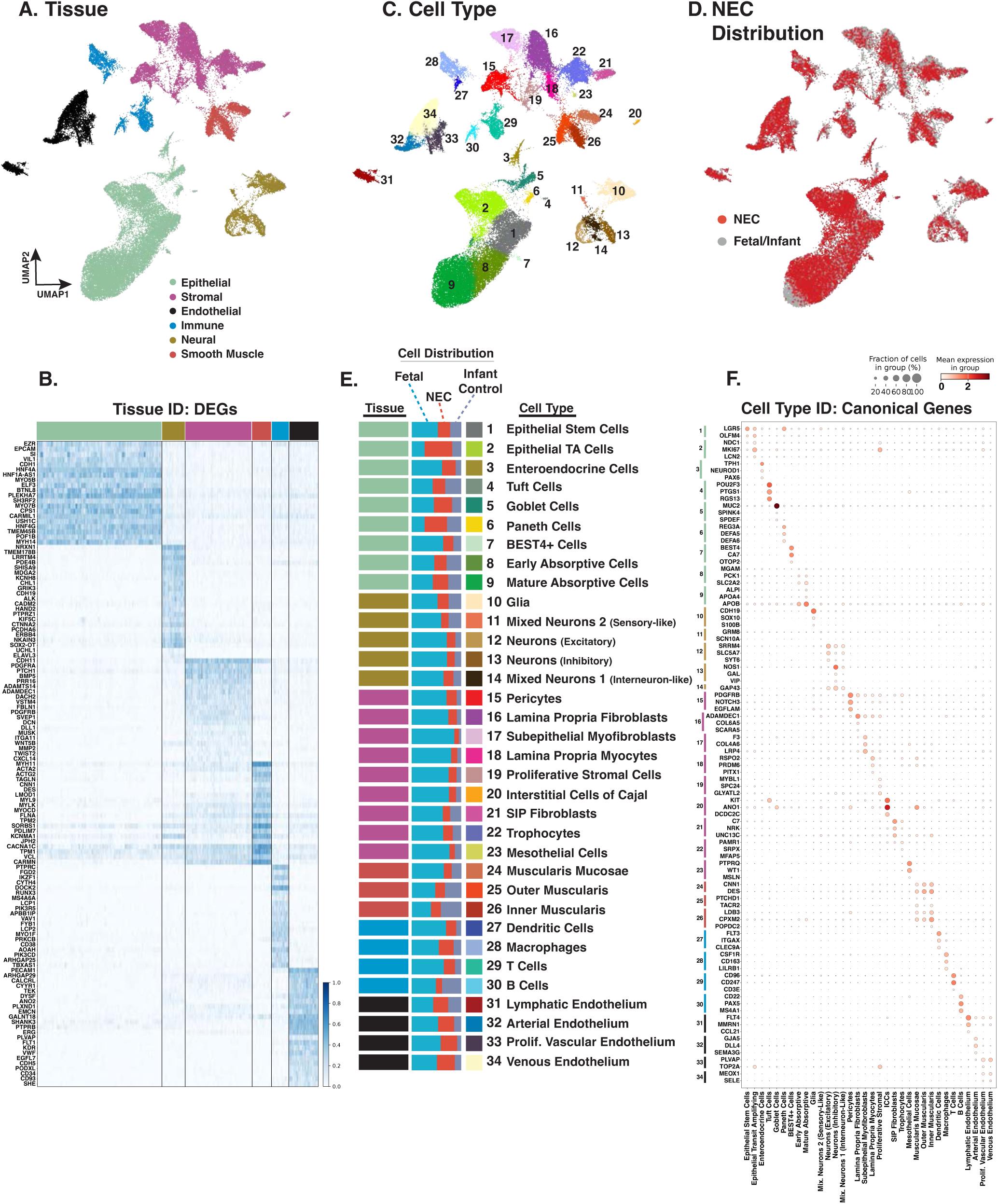
Single-nucleus analysis resolves major intestinal tissue compartments and 34 cell populations across fetal, NEC, and infant samples. **A.** Integrated UMAP of 53,735 nuclei grouped into six major tissue compartments: epithelial, stromal, endothelial, immune, neural, and smooth muscle. **B.** Tissue-level DEG heatmap showing distinct molecular signatures that support broad tissue-compartment assignment. Color key for tissue type is from Figure 2A. Gene expression was independently scaled from 0 to 1 for each gene, with 0 representing the minimum and 1 representing the maximum expression across all nuclei. **C.** Higher-resolution clustering identified 34 transcriptionally distinct cell populations across the six compartments. **D.** Overlay of NEC nuclei with fetal and infant control nuclei showing NEC representation across each cell type. **E.** Distribution of fetal, NEC, and infant control nuclei across the 34 annotated cell populations. Color key for tissue type is from Figure 2A. **F.** Dot plot of canonical marker genes supporting cell-type annotation, with dot size indicating the fraction of nuclei expressing each gene and color indicating mean expression.

Higher-resolution clustering resolved 34 transcriptionally distinct cell populations across the six tissue compartments (Fig. 2C). The epithelial compartment included intestinal stem cells (ISCs), transit-amplifying (TA) cells, early absorptive (EA) and mature absorptive (MA) cells, goblet cells, enteroendocrine cells (EECs), Paneth cells, BEST4+ cells, and tuft cells. The stromal compartment included pericytes, lamina propria fibroblasts, subepithelial myofibroblasts, lamina propria myocytes, proliferative stromal cells, interstitial cells of Cajal (ICCs), SIP fibroblasts, trophocytes, and mesothelial cells. The endothelial compartment included lymphatic, arterial, proliferative vascular, and venous endothelial populations. The immune compartment included dendritic cells, macrophages, T cells, and B cells. The neural compartment contained glia and four neuronal populations. Two neuronal populations were assigned as excitatory and inhibitory neurons, while Mixed Neuron 1 and Mixed Neuron 2 did not resolve into distinct neuronal lineages based on canonical marker expression. Subclustering suggested that subsets of Mixed Neuron 1 showed low and variable interneuron-associated markers, and subsets of Mixed Neuron 2 showed low and variable markers of intrinsic primary afferent/sensory neurons, but cell numbers and marker strength were insufficient for confident lineage assignment. The smooth muscle compartment included muscularis mucosae, inner circular muscularis, and outer longitudinal muscularis.

NEC nuclei were present in all 34 annotated populations, as were fetal and infant control nuclei, indicating that the dataset captured NEC-associated transcriptomic states across all major intestinal tissue and cell types (Fig. 2D-E). Cell-type assignments were further supported by selective expression of established canonical genes within their corresponding populations, which helped distinguish related populations within the same tissue compartment and supported identification of both abundant and less common intestinal cell states (Fig. 2F). Some population clusters, including interstitial cells of Cajal, BEST4+ cells, mesothelial cells, tuft cells, Mixed Neuron 1, Mixed Neuron 2, and dendritic cells, did not yield sufficient NEC-versus-infant differential expression to support the same depth of cell-type-specific analysis. These populations were therefore retained in the global cellular framework but were not developed into full individual NEC narratives. Together, recovery of six tissue compartments, 34 transcriptionally distinct cell populations, and NEC nuclei across every identified population established the cellular framework for the cross-lineage and cell-type-specific analyses that follow.

### Analytical Primer for NEC-Regulated Programs and Developmental Maturity

Because the same analytical framework is applied across multiple intestinal cell populations, a common interpretive approach was used throughout the study; detailed analytical procedures are provided in the Methods. Illustrative expression patterns show how gene-expression changes may suggest NEC-specific regulation, fetal-like persistence or reversion, intermediate fetal-to-infant expression consistent with incomplete maturation, developmentally preserved expression relative to the fetal and infant references, or no difference across groups (Fig. 3). Fetal-like persistence or reversion encompasses two biologically distinct possibilities: failure to acquire an infant-associated regulatory state or an NEC-associated shift toward a more fetal-like expression state. These expression patterns are used to interpret NEC-associated genes in relation to both disease and normal fetal-to-infant maturation. NEC-specific regulation refers to genes significantly changed in NEC compared to infant controls and distinct from the normal fetal-to-infant pattern, whereas other patterns identify NEC-regulated genes that remain fetal-like, extend beyond fetal or infant reference levels, or fall between the two developmental states (Fig. 3A).

**Fig. 3.**
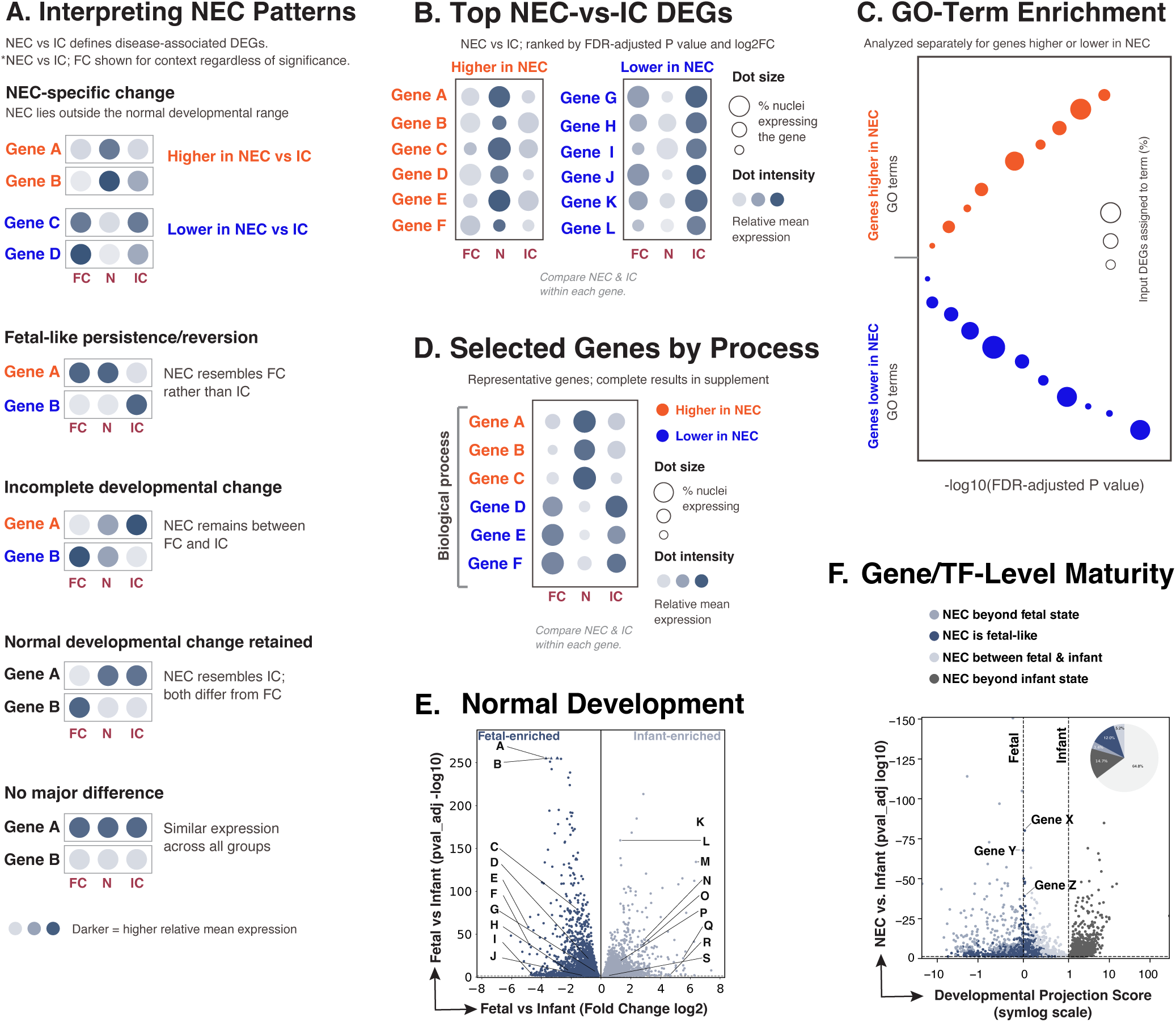
Analytical primer for NEC differential expression and developmental maturity analyses. **A.** Illustrative expression patterns used to interpret NEC-associated changes in developmental context, including NEC-specific change, fetal-like persistence/reversion, incomplete developmental change, retained normal developmental change, and no major difference. These are interpretive examples, not separate statistical categories. For NEC-associated gene analyses, significance is defined by NEC versus infant control (IC); fetal control (FC) expression is shown for developmental context regardless of significance. **B.** Top differentially expressed genes (DEGs) in the “NEC-specific change” category, where NEC expression is statistically different than both fetal and infant controls, ranked by false discovery rate (FDR)-adjusted P value and log2 fold change (log2FC). Dot size indicates the percentage of nuclei expressing each gene, and dot intensity indicates relative mean expression across groups within each gene. Genes upregulated in NEC (orange) and downregulated in NEC (blue) are shown in separate dotplots. **C.** Gene Ontology (GO)-term enrichment analysis performed using genes higher and lower in NEC compared to infant control for each cell type. Horizontal position indicates−log10(FDR-adjusted P value), and dot size indicates the percentage of input DEGs assigned to each term. GO terms enriched in NEC (orange) and depleted in NEC (blue) are both shown. **D.** Representative NEC-regulated genes grouped by biological process. Orange text indicates higher expression in NEC relative to IC, and blue text indicates lower expression. Dot size and intensity are as defined in panel B. **E.** Volcano plot of FC-versus-IC DEGs defining the normal fetal-to-infant developmental reference. **F.** Gene-level maturity projection of NEC-regulated developmental genes relative to fetal and infant reference states. Vertical dotted lines indicate fetal and infant reference scores; dot color classifies genes as beyond fetal, fetal-like, between fetal and infant, or beyond infant, based on the mean gene expression in each. The developmental projection score is displayed on a symmetric logarithmic (symlog) scale. Only genes significantly altered in NEC versus IC are included. Pie chart in corner shows percentage of each gene category. Complete DEG, GO-term, and maturity results are provided in the supplemental files. Abbreviations: N, necrotizing enterocolitis; FC, fetal control; IC, infant control; DEG, differentially expressed gene; GO, Gene Ontology; FDR, false discovery rate; symlog, symmetric logarithmic.

For each cell population, we compared NEC with both fetal and infant controls and separated significant DEGs into those upregulated and downregulated in NEC compared to both controls. The strongest examples are shown in the top-gene dot plots, ranked by statistical significance and fold-change size (Fig. 3B). In these plots, larger dots indicate a greater percentage of nuclei expressing the gene, while darker dots indicate higher relative mean expression (See Methods). Fetal control samples are shown only to provide developmental context; statistical testing was performed for NEC versus infant controls. When enough significant genes were available, GO term enrichment was performed separately for genes higher and lower in NEC compared to infant controls (Fig. 3C). Curated panels highlight representative NEC-associated genes grouped by biological process (Fig. 3D). Orange marks genes or GO terms higher in NEC compared to infant controls and blue marks those lower in NEC compared to infant controls. Complete DEG and GO term results are provided in the supplemental files.

Because NEC develops in an intestine that is still maturing, we assessed developmental maturity for each cell type at the gene level. Fetal and infant controls defined a developmental reference axis for each gene (Fig. 3E). Gene-expression patterns along this axis are consistent with known human intestinal maturation trajectories from fetal development through infancy.^47–49^ We then evaluated NEC expression along this axis for genes that changed during normal development (Fig. 3F). Dot color indicates where NEC expression falls relative to the fetal and infant control reference means (see Methods for analytical approach). *‘Fetal-like’* dots indicate NEC expression that is not statistically different from fetal controls; *‘between fetal and infant’* dots indicate expression that differs from both references and falls between them; and *‘beyond fetal’* or *‘beyond infant’* dots indicate expression that extends significantly past the corresponding reference mean (Fig. 3F). Together, these categories distinguish retained fetal-like expression or incomplete postnatal maturation from disease-associated changes outside the expected developmental range. To highlight regulatory programs linked to developmental displacement in NEC, developmentally regulated TFs were plotted on a separate graph (Fig. 3F). Genes not significantly different between NEC and infant controls are shown as not significant (N/S) and excluded from the TF plots.

### Shared NEC-Associated Transcriptional Programs Span Multiple Intestinal Lineages

We first asked whether NEC-associated transcriptional changes were confined to individual cell types or shared across the intestine. We identified NEC-associated DEGs and ranked them by the number of cell populations in which they changed in the same direction, and the 30 most broadly regulated genes with clear biological interpretability were selected for display (Fig. 4A,B). Genes were considered recurrent if they were significantly changed in the same direction in at least 16 of the 28 analyzed cell types, or about 57% of the dataset (Fig. 4A,B). This identifies NEC-associated genes that are shared across many intestinal populations and provides a tissue-wide context for the cell-type analyses that follow.

**Fig. 4.**
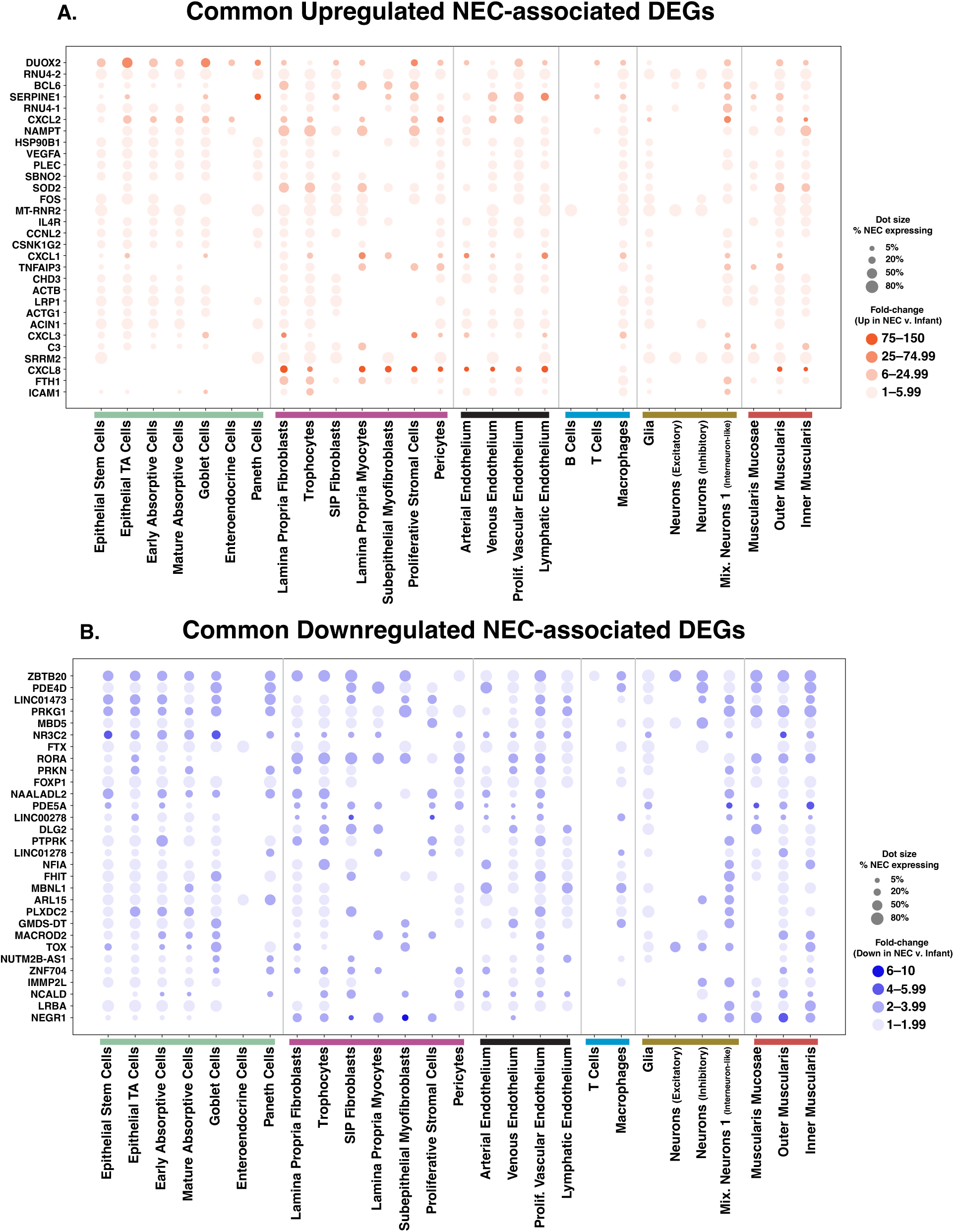
Shared NEC-associated genes across intestinal cell populations. **A.** Dot plot of broadly shared genes significantly upregulated in NEC relative to infant controls. **B.** Dot plot of broadly shared genes significantly downregulated in NEC relative to infant controls. Genes were ranked by the number of cell populations in which they were significantly altered in the same direction, and displayed genes were selected from the top 30 biologically interpretable recurrent DEGs that fit within the figure layout. Displayed genes were significant in at least 16 of 28 cell populations with NEC-associated DEGs. Cell types not shown did not have significant NEC-associated DEGs for the displayed genes, although they may contain other NEC-associated DEGs outside the selected genes. Dot size indicates the percentage of NEC nuclei expressing each gene; orange and blue indicate higher and lower expression in NEC, respectively, with darker color denoting a larger NEC-versus-infant fold change; lack of a dot indicates that the displayed gene was not significantly altered in NEC in that cell type, or that the displayed gene was significantly altered in NEC but in less than 5% of NEC cells. Cell populations are grouped by major cellular compartment. Abbreviations: NEC, necrotizing enterocolitis; DEG, differentially expressed gene; TA, transit-amplifying.

Broadly upregulated genes revealed recurrent injury-associated programs involving epithelial antimicrobial/redox defense, chemokine-mediated inflammatory recruitment, stress-adaptive metabolism and proteostasis, complement activation, and vascular inflammatory activation (Fig. 4A). *DUOX2* was most induced across epithelial populations, consistent with its role in epithelial hydrogen peroxide production and mucosal antimicrobial defense.^50^ *CXCL1*, *CXCL2*, and *CXCL3* encode neutrophil-active CXC chemokines; their broad increase across fibroblast, perivascular, and endothelial populations suggests that structural cells may contribute to inflammatory cell recruitment during NEC.^51–53^ *NAMPT*, *HSP90B1* (encoding GRP94), *SOD2*, *FTH1*, and *FOS* were increased across several compartments, marking a shared stress-adaptation program spanning inflammatory metabolism, ER proteostasis, antioxidant defense, iron-redox buffering, and immediate-early transcriptional activation.^54–58^ Vascular and perivascular populations also showed increases in *SERPINE1* (encoding PAI-1), *VEGFA*, and *ICAM1*, consistent with fibrinolytic/matrix remodeling, hypoxia-associated angiogenic signaling, and leukocyte-endothelial interaction.^59–63^ Increased *C3* further suggested local complement-associated inflammatory amplification across multiple intestinal populations.^64^ These shared increases may help cells respond to injury, but if they persist, they could maintain inflammation, oxidative stress, complement activity, and vascular activation in ways that worsen NEC progression.

Broadly downregulated genes pointed to reduced cyclic nucleotide signaling, transcriptional and chromatin-associated regulation, tissue organization, and organelle-associated homeostasis (Fig. 4B). *ZBTB20* was decreased across nearly the entire analyzed cell-type set, suggesting broad reduction of a transcriptional regulator linked to cell differentiation, immunometabolic regulation, intestinal immune homeostasis, and neuronal identity programs.^65–69^ *PDE4D*, *PDE5A*, and *PRKG1* were also broadly reduced, indicating lower cAMP- and cGMP-associated signaling capacity across multiple epithelial, stromal, endothelial, neural, and muscular populations.^70–72^ This extended to transcription factor-associated genes, including *RORA*, *FOXP1*, and *NFIA*; chromatin-associated regulation through *MBD5*; cell signaling and tissue organization through *PTPRK* and *DLG2*; and mitochondrial or organelle-associated homeostasis through *PRKN* and *IMMP2L*.^73–82^ Together, these reductions suggest that NEC broadly lowers genes that help cells maintain identity, communicate through cyclic nucleotide signaling, organize tissue structure, and preserve organelle function.

These recurrent NEC-associated genes define a tissue-wide injury signature. Increased antimicrobial/redox defense, chemokine recruitment, complement, stress-adaptation, and vascular-inflammatory genes may amplify early epithelial-microbial and immune-recruitment signals and, if sustained, reinforce inflammatory, oxidant, complement, and vascular-injury loops. Broad reductions in cyclic nucleotide signaling, transcriptional/chromatin regulation, tissue organization, and organelle homeostasis suggest weaker capacity to restore stable specialized function. Failed resolution may therefore reflect persistent injury signaling plus reduced machinery for coordinated tissue recovery.

### Regulated Cell-Death Programs Show Cell-Type-Specific Susceptibility, Restraint, and Compensation

Cell death has long been recognized as a central feature of NEC tissue injury.^83–84^ Although apoptosis, necroptosis, pyroptosis, and ferroptosis are executed largely through protein-level events, transcriptional regulation can prime pathway susceptibility.^46^ Examples include BIM induction in apoptosis, transcriptional priming of the NLRP3 inflammasome, maintenance of an MLKL expression threshold that licenses necroptosis, and BACH1-dependent control of the ferroptosis threshold.^85–88^ Prior NEC studies have implicated epithelial apoptosis in neonatal rat NEC, necroptosis in human and mouse NEC tissues, macrophage pyroptosis and pyroptosis-associated epithelial injury in NEC, and ferroptosis in NEC transcriptomic and experimental models.^89–93^ However, most studies have not resolved these programs across the full range of human intestinal cell types in NEC.

Apoptosis-associated regulation was concentrated in epithelial populations, led by mature absorptive, stem, transit-amplifying, and early absorptive cells (Fig. 5A). Early neonatal rat NEC studies showed that epithelial apoptosis precedes bowel necrosis but did not distinguish apoptosis across specific epithelial cell populations.^89^ Increases in death receptor pathway genes, including *CASP8*, *CASP10*, and *TNFRSF10B*, were concentrated in ISC, TA, and absorptive populations, consistent with greater death receptor-associated apoptotic priming.^94–96^ At the same time, selected epithelial populations showed reduced *CASP3*, *BIK*, and *BCL2L11* (encoding BIM), which may suggest transcriptional restraint of downstream apoptotic execution and mitochondrial apoptotic priming.^97–99^ Anti-apoptotic genes were also reduced in selected epithelial or vascular populations, including *BCL2*, *BCL2L1* (encoding BCL-XL and other isoforms), and *MCL1*, suggesting lower survival buffering in those cell states.^100–102^ *TNFAIP3/*A20 was increased across multiple cell types. Although A20 generally restrains TNF/NF-κB signaling, elevated epithelial A20 can paradoxically enhance TNF-induced, RIPK1-caspase-8-dependent apoptosis.^103^ Together, the coordinated increase in death-receptor/initiator-caspase signaling and loss of selected pro-survival genes suggest that NEC epithelium is transcriptionally primed for apoptosis, with reduced buffering against extrinsic death signals despite heterogeneous regulation of downstream execution pathways.

**Fig. 5.**
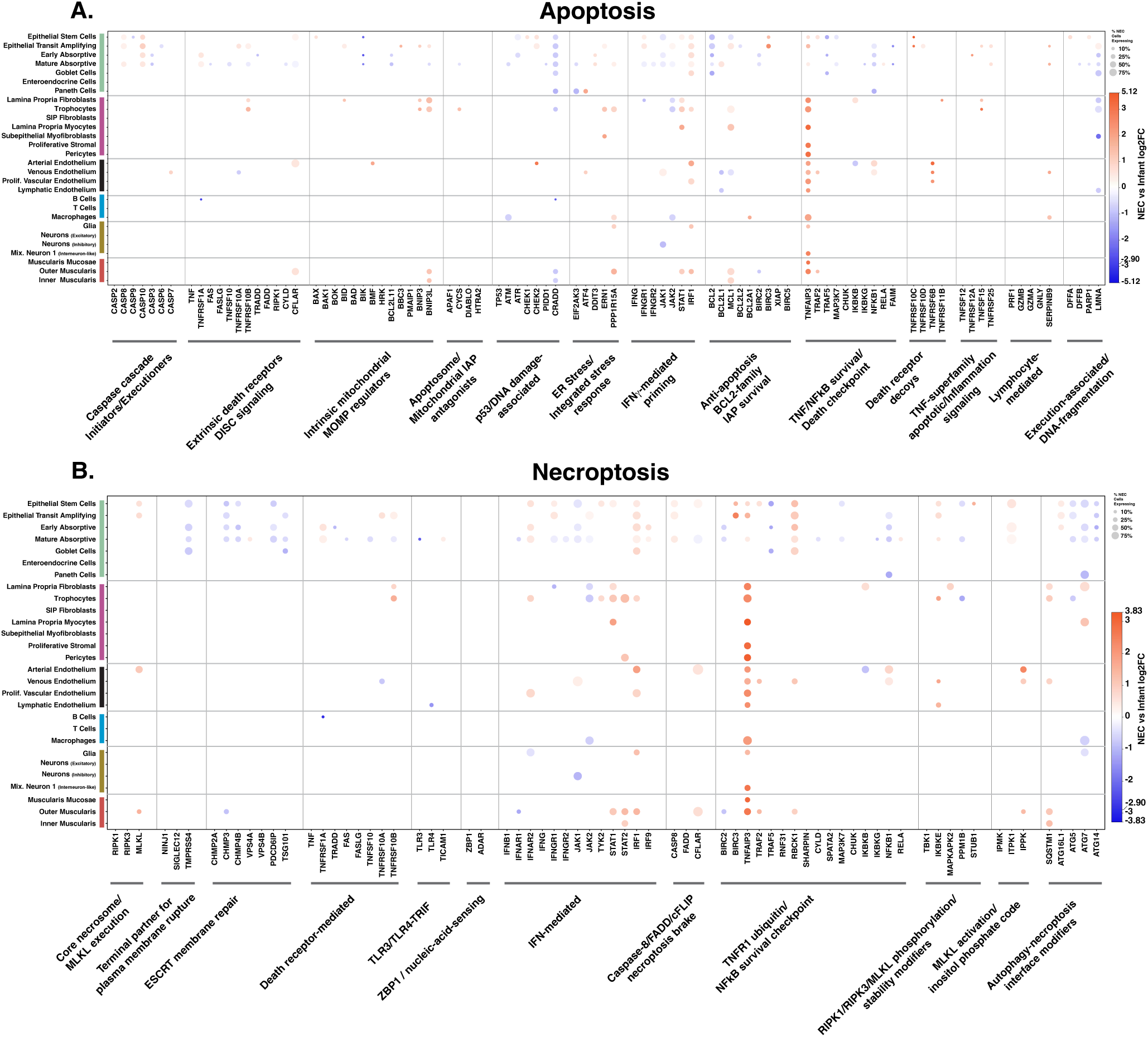
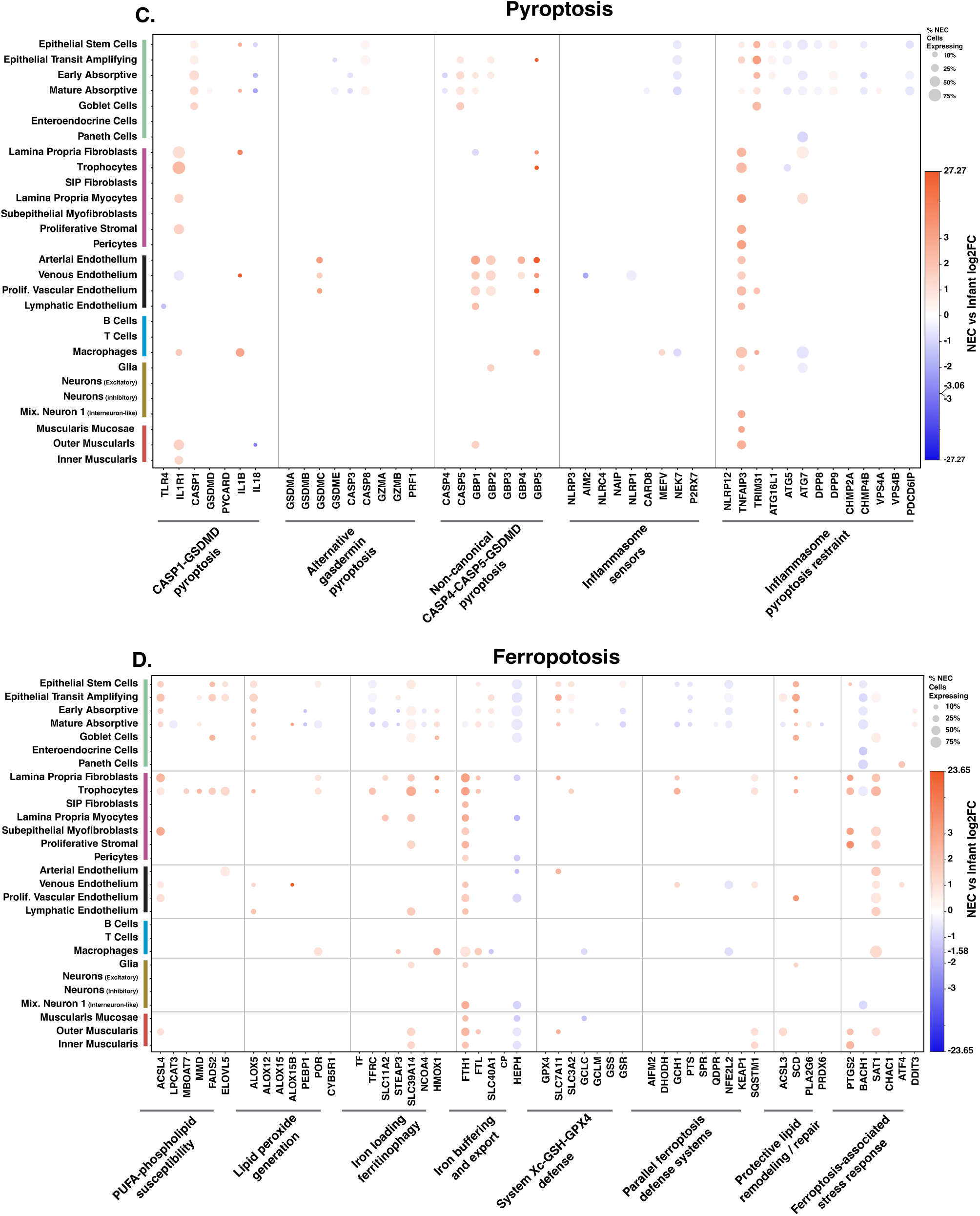
NEC-associated regulation of cell-death pathway genes across intestinal cell populations. Dot plots show NEC-versus-infant DEGs associated with **A.** apoptosis, **B.** necroptosis, **C.** pyroptosis, and **D.** ferroptosis across epithelial, stromal, vascular, immune, neural, and smooth muscle populations. Genes are grouped by pathway-related functions, with dot size indicating the fraction of NEC nuclei expressing each gene and color indicating NEC-versus-infant log2 fold change. Orange indicates genes upregulated in NEC relative to infant controls, and blue indicates genes downregulated in NEC relative to infant controls. The log2 fold change is displayed on a symmetric logarithmic (symlog) scale. These panels identify cell-type-specific patterns of death-pathway priming, restraint, and compensation.

**Fig. 6.**
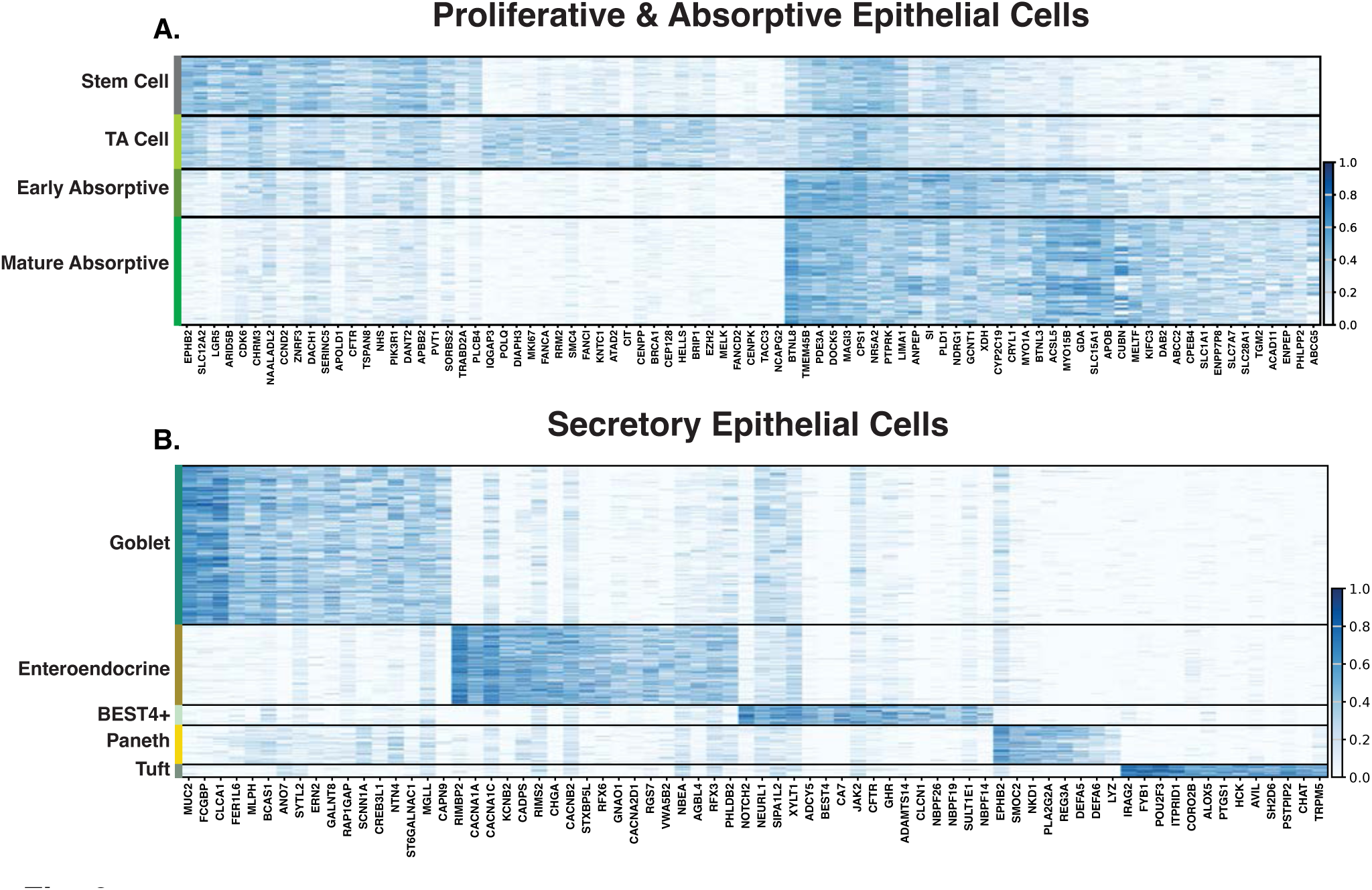
Epithelial cell-type signatures define proliferative, absorptive, and secretory-associated populations. **A.** Heatmaps show cell-type marker-gene expression across epithelial populations. Proliferative and absorptive populations include epithelial stem cells, transit-amplifying cells, early absorptive cells, and mature absorptive cells. **B.** Secretory-associated populations include goblet, enteroendocrine, BEST4+, Paneth, and tuft cells. Distinct transcriptional signatures support epithelial cell-type assignments and provide the framework for the detailed epithelial analyses that follow.

Necroptosis-associated transcriptional changes were detected across all tissue types but were most prominent in epithelial populations (Fig. 5B). Prior NEC work localized necroptosis mainly to differentiated villus epithelium but did not resolve the specific epithelial cell types involved; our data identify necroptosis-associated transcriptional changes across distinct epithelial and non-epithelial populations.^90^ In our dataset, *MLKL* increased in arterial endothelium, epithelial stem and transit-amplifying cells, and outer muscularis, suggesting greater necroptosis susceptibility across multiple tissue compartments.^104–105^ By contrast, *TLR4*, *TRADD*, and *FAS* were reduced in several populations, indicating that NEC does not produce a simple, coordinated increase in all necroptosis-pathway components.^106–108^ Reduced *CHMP3*, *CHMP4B*, *PDCD6IP*, and *TSG101* across epithelial, stromal, and muscle cell populations further suggested weaker ESCRT III-dependent membrane repair, which normally helps limit MLKL-mediated plasma membrane injury and necroptotic death.^109^

Pyroptosis-associated transcriptional changes were concentrated across epithelial and endothelial compartments (Fig. 5C). In the endothelium, arterial and venous endothelial cells showed increased *GBP1*, *GBP2*, *GBP5*, and the final effector *GSDMC*, and venous endothelium also showed increased *IL1B* with reduced *AIM2* and *NLRP1*. These changes implicate lipopolysaccharide (LPS)-responsive guanylate-binding protein (GBP) signaling, inflammasome-associated IL-1β biology, and gasdermin-linked pyroptotic priming in the vascular compartment.^110–116^ In stroma, pyroptosis-associated regulation centered more on inflammasome modifiers, inflammatory feedback, and membrane-repair/autophagy genes than canonical *CASP1-GSDMD* execution. In epithelium, mature absorptive cells increased *CASP1*, *CASP5*, *CASP8*, *IL1B*, *GBP1*, and *GSDMD*, while early absorptive and transit-amplifying cells increased *CASP1*, *CASP5*, and GBP-family genes, consistent with stronger inflammasome/gasdermin-associated priming.^110,117–119^ Macrophages showed mixed regulation, with increased *GBP5*, *IL1B*, and *MEFV* but reduced *NEK7*.^112,120–121^ Interestingly, *IL1R1* increased in multiple cell types, including macrophages and stromal populations, while changes in *TLR4* transcript abundance were limited. These findings suggest increased potential for IL-1 responsiveness across compartments, consistent with predicted IL1B-IL1R1 interactions in human NEC and the protective effects of IL-1 receptor antagonism in perimental NEC.^122^ Prior NEC work implicated macrophage NLRP3-caspase-1-IL-1β signaling; our data broaden pyroptosis-associated regulation to endothelial, epithelial, and stromal compartments.^91^

Prior NEC work combining transcriptomic analysis with experimental models implicated ferroptosis and showed reduced NEC-associated injury after ferroptosis inhibition.^93^ In our study ferroptosis-associated regulation was especially notable in stromal and epithelial compartments (Fig. 5D). Trophocytes and lamina propria fibroblasts showed prominent regulation of genes involved in iron handling, lipid peroxidation biology, and antioxidant defense. Pro-ferroptotic *ACSL4* increased in endothelial, epithelial, stromal, and muscle populations, while *ALOX5* increased in endothelial, epithelial, and stromal populations, suggesting broader lipid peroxidation susceptibility.^123–124^ Conversely, reduced *TFRC*, *NCOA4*, and *BACH1* mainly in epithelial populations may dampen distinct ferroptosis-sensitizing inputs related to iron loading, ferritinophagy, and labile-iron/glutathione regulation.^88,125–126^ Anti-ferroptotic defenses varied by cell type, with increased *SLC7A11* and *GCH1* suggesting stronger cystine/glutathione and BH4-linked protection in some populations, whereas reduced *NFE2L2*, *GCLC*, *GCLM*, *GSR*, and *GCH1* suggested weaker antioxidant and glutathione/BH4 defenses in others.^88,127–131^

The cell death-associated gene patterns suggest uneven death-pathway priming, restraint, and buffering across the NEC intestine. Conceptually, NEC initiation may involve compartments with higher transcriptional susceptibility to death receptor signaling, MLKL-associated membrane injury, inflammasome/gasdermin biology, or lipid peroxidation stress. NEC may progress when death-pathway priming occurs alongside limited capacity for cell survival, membrane repair, antioxidant defense, and anti-inflammatory control. Resolution of NEC may depend on whether vulnerable compartments maintain enough protective capacity to restrain injury and repair, or whether persistent death-pathway priming contributes to dysfunction. These RNA data prioritize cellular compartments for protein-level validation of caspase activity, MLKL phosphorylation, inflammasome activation, membrane injury, and lipid peroxidation stress. This framework may help prioritize therapies that protect vulnerable intestinal cell populations by limiting maladaptive death, thus supporting barrier repair and tissue recovery.

### Deep Cell-Type Analysis Suggests NEC-Associated Programs and Impaired Developmental Progression

We next examine each tissue compartment and its constituent cell populations in greater depth. Each section integrates anatomical context, NEC-associated genes and biological processes, selected gene-level programs, and developmental maturity. This framework enables comparison across cell types while highlighting disease-associated programs, impaired maturation, and responses linked to NEC progression, repair, or failed resolution.

### Epithelial compartment

The epithelial compartment, marked by *EPCAM* and *VIL1* (Fig. 2B), includes proliferative, absorptive, and secretory populations that support epithelial renewal, barrier function, nutrient absorption, host defense, mucus and hormone production, and chemosensory surveillance.^132–136^ Intestinal stem cells (ISCs) and transit-amplifying (TA) cells sustain crypt-based renewal and generate absorptive and secretory lineages along the crypt-villus axis.^137–141^ Absorptive enterocytes form a progressive differentiation trajectory and carry out digestive, transport, lipid-handling, and barrier-associated functions, while goblet, enteroendocrine, Paneth, BEST4+, and tuft cells selectively support mucus-barrier function, hormone signaling, antimicrobial defense, ion/pH regulation, and chemosensory immune activation.^142–151^

To define the epithelial populations in our study, heatmap-based annotation of marker-gene expression resolved nine epithelial populations, including *LGR5*+/*OLFM4*+ ISCs, *MKI67*+/*TOP2A*+ TA cells, *SI*+/*SLC5A1*+ early absorptive enterocytes, *APOA1*+/*FABP1*+ mature absorptive enterocytes, *MUC2*+/*FCGBP*+ goblet cells, *CHGA*+/*CHGB*+ enteroendocrine cells, *DEFA5*+/*REG3A*+ Paneth cells, *BEST4*+/*CA7*+ BEST4+ cells, and *POU2F3*+/*TRPM5*+ tuft cells (Fig. 2F; 6A,B).^141^ BEST4+ cells were grouped with secretory-associated populations for presentation because human trajectory analyses suggest that small-intestinal BEST4+ cells align more closely with secretory progenitors.^49,141,148^ BEST4+ and tuft cells were retained in the epithelial map, but their NEC-versus-infant differential-expression signal was not sufficient to support detailed cell-level interpretation.

The following sections examine how NEC alters epithelial populations from the crypt base to the villus surface. ISCs and TA cells capture the regenerative and proliferative response that must rebuild injured epithelium. Early absorptive (EA) and mature absorptive (MA) enterocytes extend this view along the absorptive trajectory, where stress responses, transport defects, detoxification changes, polarity defects, and terminal villus-function loss can be distinguished across differentiation. Goblet cells, enteroendocrine cells, and Paneth cells capture secretory-lineage responses involving mucus-barrier function, epithelial-neural signaling, antimicrobial defense, and intestinal stem-cell niche support. Together, these sections frame NEC epithelium as a regenerating but stressed tissue that may initiate repair and defense responses from crypt to villus, but may fail to restore a fully mature barrier, transport, and secretory epithelium.

### Intestinal Epithelial Stem Cells (ISCs)

ISCs occupy the crypt base, where they are rare but functionally essential.^139^ Clonal and functional studies estimate approximately 5-7 working stem cells per intestinal crypt.^152–153^ ISCs sustain epithelial renewal and generate transit-amplifying progenitors that replenish absorptive and secretory lineages along the crypt-villus axis (Fig. 7A).^138–139,141^ The crypt unit is highly relevant to NEC because epithelial repair after injury depends on preservation or restoration of the ISC compartment and progenitor output.^154^ Mouse NEC studies link ISC TLR4 activation to ER stress, apoptosis, reduced proliferation, and impaired repair.^16–17^ Rat NEC models also show loss of OLFM4/LGR5+ ISCs, whereas mouse and human NEC tissue show reduced ISC markers and crypt proliferation.^155–156^ Prior human NEC work used bulk RNA-seq to estimate crypt-zone enterocytes, but this approach did not establish a direct ISC-specific NEC DEG program.^18^ In experimental neonatal mouse NEC multi-omics, NEC was associated with reduced ISC/TA abundance, impaired NEC-derived organoid growth, and lower *Lgr5*, *Sox9*, and *Pcna*, further supporting impaired stem/progenitor maintenance.^157^ Thus, how NEC alters the human ISC compartment remains unresolved, including whether ISCs preserve crypt identity while adopting inflammatory-stress and repair-associated regulatory changes that could influence epithelial regeneration.

**Fig. 7.**
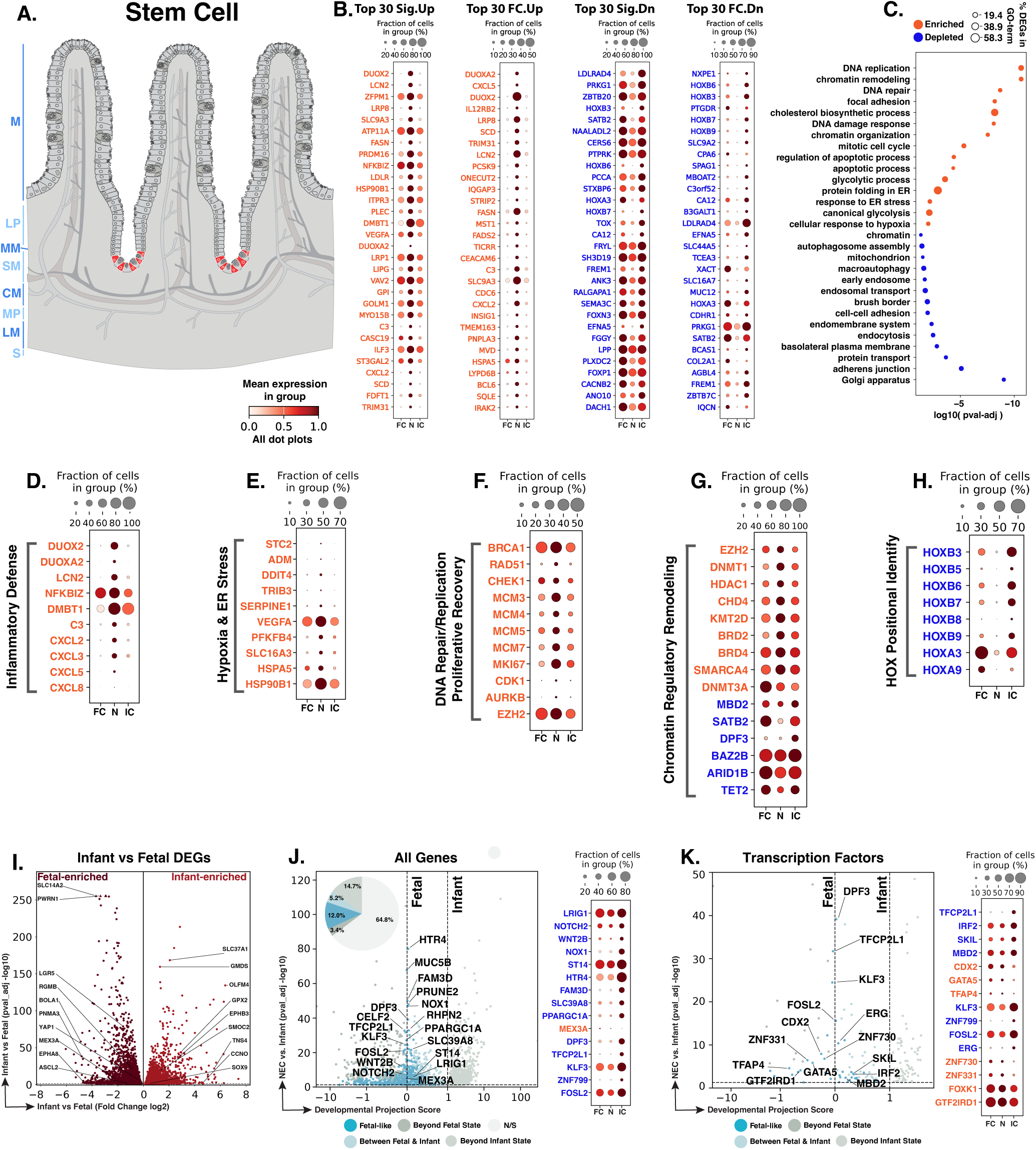
NEC intestinal stem cells show inflammatory, stress, repair, chromatin, and maturity-associated changes. **A.** Schematic showing the crypt-base location of intestinal stem cells (red). Blue text to the left of the Panel A schematic indicates general anatomical regions: M, mucosa; LP, lamina propria; MM, muscularis mucosae; SM, submucosa; CM, circular muscle; MP, myenteric plexus; LM, longitudinal muscle; S, serosa. **B.** Dot plots of top NEC-specific genes ranked by significance and fold change. **C.** GO-term analysis showing enriched and depleted biological processes among NEC-associated DEGs. **D–H.** Curated gene panels showing inflammatory-defense, hypoxia/ER-stress, DNA-repair/proliferative-recovery, chromatin-regulatory, and HOX positional-identity programs. In panels **B** and **D–H**, orange gene text indicates upregulation in NEC relative to infant controls, whereas blue gene text indicates downregulation. **I.** Volcano plot of DEGs between fetal and infant controls that define the developmental trajectory for stem cells. **J–K.** Gene-level and transcription-factor maturity analyses of fetal-versus-infant DEGs, highlighting the subset also significantly changed in NEC relative to infant controls and positioned relative to fetal and infant reference states, and their corresponding dotplots. The developmental projection score in panels **J–K** is displayed on a symmetric logarithmic (symlog) scale. For expression dot plots, dot size indicates the fraction of nuclei expressing each gene and dot color indicates mean expression. In maturity plots, vertical dotted lines indicate the fetal and infant reference scores. Colored dots classify NEC-versus-infant DEGs as Fetal-like, Between Fetal & Infant, Beyond Fetal State, or Beyond Infant State; gray N/S dots are fetal-versus-infant DEGs without a significant NEC-versus-infant change.

Differential-expression analysis of NEC ISCs compared to healthy infant controls identified 3,950 genes significantly changed, including 1,585 upregulated and 2,365 downregulated genes (Supplemental Data). Genes with the strongest adjusted significance and fold-change rankings were selected to highlight biologically interpretable DEGs increased or decreased in NEC (Fig. 7B). DEG and GO-term patterns converged on an injury-stressed ISC state. Increased programs reflected inflammatory-defense activation, hypoxia/ER-stress, DNA repair/replication, cell-cycle activity, and chromatin regulation (Fig. 7B,C). Reduced programs pointed to weaker epithelial homeostatic organization, including lower positional identity, adhesion, endomembrane/transport, and brush-border-associated maturation (Fig. 7B,C). Together, these process-level findings suggest that NEC ISCs activate inflammatory, metabolic-stress, and cell-cycle-associated repair programs while expressing lower levels of genes linked to positional identity and epithelial structural organization.

Inflammatory-defense genes were increased in NEC stem cells at the crypt base (Fig. 7D). Higher *NFKBIZ* supports NF-κB-associated inflammatory transcription through IκBζ-dependent control of inducible inflammatory genes.^158–159^ Increased *DUOX2* and *DUOXA2* point to epithelial oxidant defense linked to mucosal hydrogen peroxide generation and inflamed intestinal epithelium.^50,160^ Elevated *LCN2* and *DMBT1* extend this program to antimicrobial siderophore sequestration, mucosal defense, and epithelial immune-associated differentiation.^161–162^ Higher *CXCL2*, *CXCL3*, *CXCL5*, and *CXCL8* indicate increased chemokine-recruitment capacity, while increased *C3* suggests complement-associated inflammatory amplification.^64,163–165^ Together, these findings place inflammatory-defense activity within the ISC compartment, suggesting that ISCs contributing to NEC repair are exposed to antimicrobial, chemokine, and complement-associated inflammatory signaling.

A separate set of increased genes pointed to hypoxia-associated and ER-stress responses in NEC stem cells (Fig. 7E). Higher VEGFA, ADM, SERPINE1/PAI-1, PFKFB4, and SLC16A3/MCT4 point to hypoxia-associated angiogenic, vascular, matrix-regulatory, glycolytic, and lactate-transport responses.^60,62,166–169^ Upregulated *HSPA5*/BiP/GRP78 and *HSP90B1*/GRP94 are consistent with increased ER chaperone and protein-folding pressure, while *DDIT4* supports stress-linked mTOR regulation and *TRIB3* an ER-stress-inducible ATF4-CHOP response.^170–174^ This is particularly relevant because experimental NEC links epithelial TLR4 activation to ER stress, ISC apoptosis, and mucosal disruption.^17^ Together, these data suggest that NEC ISCs remain engaged in repair under combined hypoxic, metabolic, and ER stress that could limit effective epithelial replacement.

The NEC ISC response also included increased DNA repair and cell-cycle genes (Fig. 7F). Higher *BRCA1* and *RAD51* are consistent with homologous recombination and DNA damage repair capacity.^175–176^ Increased *CHEK1* supports replication-stress checkpoint signaling.^177^ Higher *MCM3*, *MCM4*, *MCM5*, and *MCM7*, interpreted as MCM2-7 complex components, suggest DNA-replication licensing and proliferative capacity.^178–179^ Increased *MKI67*, *CDK1*, and *AURKB* further support cycling and mitotic activity, with CDK1 and AURKB supporting mitotic entry, chromosome segregation, and anaphase dynamics.^180–182^ NEC ISCs retain cell-cycle and DNA repair capacity, but within an inflammatory, hypoxic, and ER-stressed crypt environment consistent with injury-associated repair.

Chromatin-regulatory changes in NEC stem cells followed two main patterns (Fig. 7G). Higher *EZH2*, *DNMT1*, *HDAC1*, and *CHD4* suggest increased repressive chromatin activity, including H3K27 methylation, DNA-methylation maintenance, histone deacetylation, and NuRD-associated remodeling.^183–186^ Higher *KMT2D*, *BRD2*, *BRD4*, *SMARCA4*, and *DNMT3A* extend this response to enhancer regulation, acetyl-chromatin recognition, SWI/SNF remodeling, and de novo DNA methylation.^187–191^ By contrast, lower *MBD2*, *SATB2*, *DPF3*, *BAZ2B*, *ARID1B*, and *TET2* indicate reduced expression of methyl-cytosine-phosphate-guanine (CpG)/NuRD regulators, epithelial-identity factors, chromatin readers, BAF/SWI-SNF components, and DNA-demethylation machinery.^192–199^ This pattern suggests that NEC ISCs increase some chromatin-regulatory programs that may support injury-associated adaptation.^200^ At the same time, they express lower levels of regulators linked to epithelial patterning, identity, differentiation, and chromatin stability.^192,194,199^ This is consistent with neonatal mouse NEC multi-omics showing altered epithelial chromatin accessibility and differentiation-associated programs.^157^

Reduced expression of multiple HOX regional-identity genes was another feature of NEC ISCs, including *HOXB3*, *HOXB5*, *HOXB6*, *HOXB7*, *HOXB8*, *HOXB9*, *HOXA3*, and *HOXA9* (Fig. 7H). HOX genes show position-specific expression along the gastrointestinal tract and contribute to regional patterning during development.^201–202^ Regional identity is retained within stem-cell-derived intestinal epithelium and is associated with stable region-specific transcriptional and epigenetic programs that shape differentiated epithelial function.^141,203–204^ Lower expression of these HOX genes in NEC ISCs may therefore reflect weakened regional positional identity. Disruption of this positional program could cause ISC progeny to replace lost cells without fully restoring location-specific barrier, antimicrobial, secretory, and metabolic functions.

Of the 6,575 genes distinguishing fetal from infant control ISCs, 2,317 were also dysregulated in NEC versus infant controls, including 787 with fetal-like expression (34.0% of the 2,317 genes; Fig. 7I,J). Several of the fetal-like repair-associated genes showed lower expression in NEC and were associated with weaker ISC niche control, crypt regeneration, Wnt/redox repair, barrier restoration, and serotonin-responsive epithelial signaling.^205–211^ For example, lower *FAM3D* may reduce epithelial host-defense and mucosal-homeostatic support, lower *SLC39A8* may affect manganese/divalent-metal transport and glycosylation, and lower *PPARGC1A* suggests reduced mitochondrial-metabolic support in a compartment where mitochondrial pyruvate metabolism regulates the balance between ISC proliferation and differentiation.^212–216^ By contrast, higher *MEX3A* in NEC ISCs suggests a fetal-like, slow-cycling reserve/progenitor feature that may support injury tolerance (Fig. 7J).^217–218^ Other regenerative-support genes showed a split pattern (Fig. 7J). Lower *WNT2B* and *NOTCH2* may weaken Wnt- and Notch-associated repair, whereas lower *LRIG1* may reduce negative feedback on ErbB signaling. Lower *ST14* and *HTR4* may indicate reduced support for epithelial barrier maintenance and 5-HT4-associated repair signaling.^205–207,209–211^

TF maturity analysis reinforced this interpretation, identifying NEC-regulated transcription factors linked to epithelial identity, chromatin state, developmental patterning, and stress-associated repair (Fig. 7K). Among the TF maturity genes, *CDX2* and *GATA5* connect the NEC ISC response to epithelial lineage maintenance and differentiation. Mouse *Cdx2* studies and intestinal epithelial cell systems support roles for Cdx2/CDX2 in intestinal epithelial identity and differentiation, while GATA-5 proteins regulate intestinal epithelial differentiation programs and mucin promoter activity.^219–222^ *MBD2* connects the TF maturity analysis to chromatin state control because MBD2 binds methyl-CpG DNA and functions as a methylation-dependent transcriptional repressor associated with histone-deacetylase complexes.^223–224^ *FOSL2*, which encodes Fra-2, adds a stress-repair signal because Fra-2 has been linked in nonintestinal systems to fibrotic, TGF-β-associated, and matrix-response programs.^225–226^ Consistent with the gene-level maturity analysis, these TF results suggest that many ISC identity-associated genes remain infant-like, while selected chromatin state and stress-response genes remain closer to the fetal reference.

NEC ISCs retain some infant-like gene identity but operate under clear injury stress. Their proliferation and DNA repair gene expression suggests they may still contribute to epithelial replacement, but concurrent inflammatory-defense, hypoxia-associated, ER-stress, and chromatin-regulatory changes indicate that this repair output arises from a stressed crypt base. Altered niche-support, HOX regional-identity, and maturity-associated repair programs may increase susceptibility at initiation by weakening the quality of early regenerative output. During progression, stressed ISCs may continue producing epithelial progeny that replace cells but do not fully restore organized crypt-villus function. Failed resolution may occur if ISC-derived repair remains poorly coordinated or insufficient to rebuild a mature, regionally patterned, barrier-competent epithelium.

### Transit-amplifying (TA) Cells

TA cells are rapidly cycling epithelial progenitors located above the crypt base, where stem-cell output is amplified before differentiating into absorptive and secretory lineages (Fig. 8A).^227^ In classical intestinal crypt models, the TA compartment has been estimated to include approximately 150 rapidly dividing cells per crypt, generating about 300 new epithelial cells per crypt each day.^228^ This compartment is central to NEC repair because recovery from villus injury requires not only preservation of ISCs, but also productive expansion of progenitor cells that can repopulate the epithelial surface. Prior human NEC single-cell work analyzed epithelial crypt-villus zonation using transcript markers such as *LGR5* and *MKI67* and reported increased crypt-like epithelial representation in NEC, but it did not define a dedicated TA cell cluster or TA-specific NEC transcriptional program.^18^ Experimental neonatal mouse NEC multi-omics identified reduced TA-cell abundance and proliferation, with altered progenitor-state and chromatin programs in stem/TA cells.^157^ This leaves open whether human NEC TA cells preserve a productive proliferative progenitor state or carry injury-associated stress programs into the compartment that expands epithelial repair output.

**Fig. 8.**
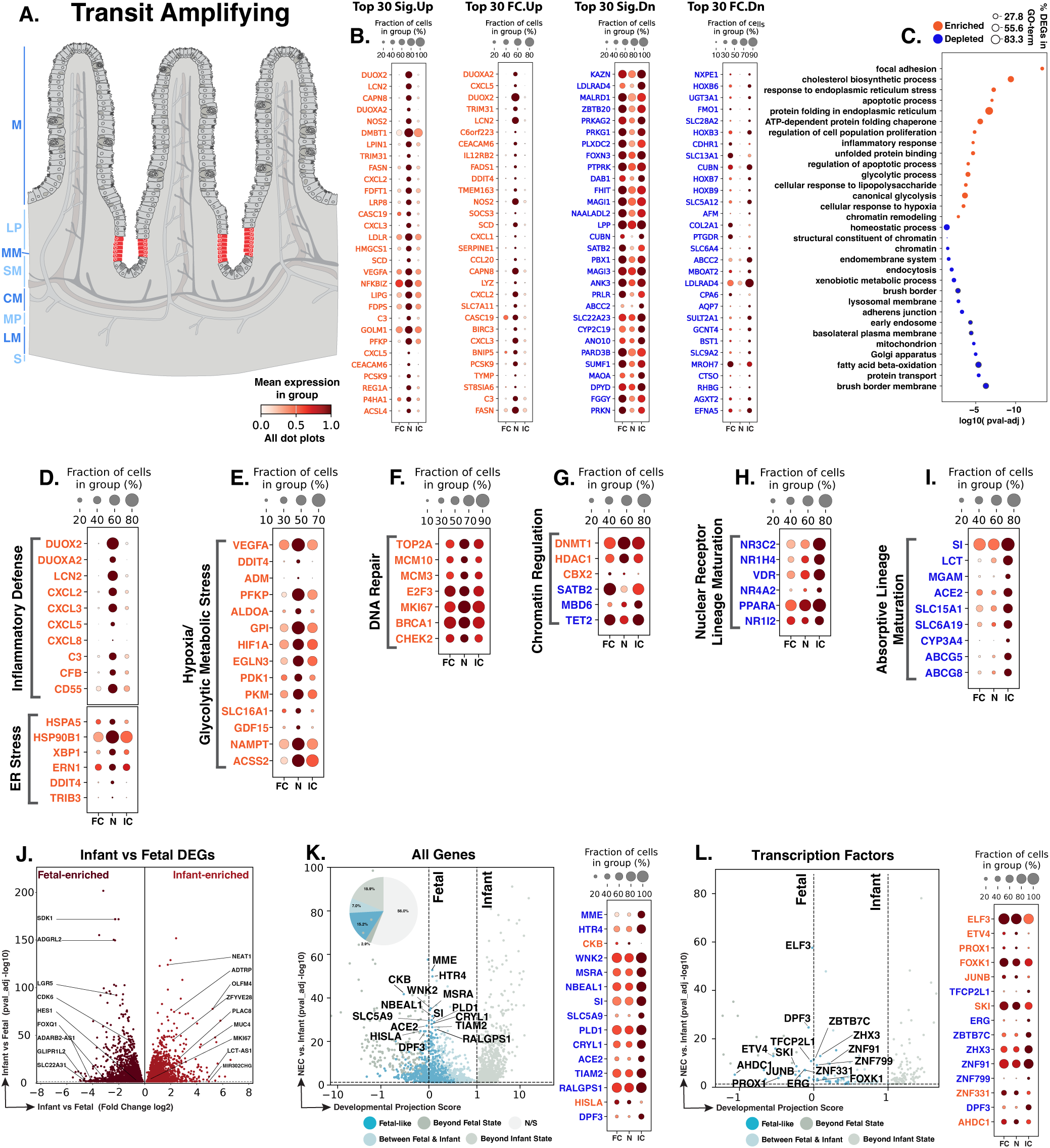
NEC transit-amplifying cells show stressed proliferative expansion and reduced absorptive-lineage maturation. **A.** Schematic showing the crypt location of transit-amplifying cells (red). Blue text to the left of the Panel A schematic indicates general anatomical regions: M, mucosa; LP, lamina propria; MM, muscularis mucosae; SM, submucosa; CM, circular muscle; MP, myenteric plexus; LM, longitudinal muscle; S, serosa. **B.** Dot plots of top NEC-specific genes ranked by significance and fold change. **C.** GO-term analysis showing enriched and depleted biological processes among NEC-associated DEGs. **D–I.** Curated gene panels showing inflammatory-defense/ER-stress, hypoxia-glycolytic/metabolic stress, DNA-repair/cell-cycle, chromatin-regulatory, nuclear-receptor, and absorptive-lineage maturation programs. In panels **B** and **D–I**, orange gene text indicates upregulation in NEC relative to infant controls, whereas blue gene text indicates downregulation. **J.** Volcano plot of DEGs between fetal and infant controls that define the developmental trajectory for TA cells. **K–L.** Gene-level and transcription-factor maturity analyses of fetal-versus-infant DEGs, highlighting the subset also significantly changed in NEC relative to infant controls and positioned relative to fetal and infant reference states, and their corresponding dotplots. The developmental projection score in panels **K–L** is displayed on a symmetric logarithmic (symlog) scale. For expression dot plots, dot size indicates the fraction of nuclei expressing each gene and dot color indicates mean expression. In maturity plots, vertical dotted lines indicate the fetal and infant reference scores. Colored dots classify NEC-versus-infant DEGs as Fetal-like, Between Fetal & Infant, Beyond Fetal State, or Beyond Infant State; gray N/S dots are fetal-versus-infant DEGs without a significant NEC-versus-infant change.

Differential-expression analysis of NEC TA cells identified 5,054 genes significantly changed in NEC compared to infant controls, including 1,788 upregulated and 3,266 downregulated genes. Top biologically interpretable DEGs, ranked by adjusted significance and fold change, are shown separately for genes increased or decreased in NEC (Fig. 8B). DEG and GO-term patterns converged on a stressed proliferative progenitor state. Increased programs reflected epithelial inflammatory-defense activation, ER protein-folding stress, hypoxia/glycolytic adaptation, DNA repair/cell-cycle activity, and chromatin regulation (Fig. 8B,C). Reduced programs pointed to weaker progression toward mature epithelial structure and absorptive function, including lower brush-border and basolateral membrane identity, vesicular/endomembrane trafficking, xenobiotic and fatty acid metabolism, mitochondrial function, and absorptive transport machinery (Fig. 8B,C). These patterns suggest that NEC TA cells remain engaged in proliferative repair under inflammatory, hypoxic, and ER stress, while showing reduced progression toward mature absorptive epithelial function.

Inflammatory-defense and ER-stress features extended from ISCs into TA cells, suggesting that NEC-associated crypt stress reaches the proliferative compartment that amplifies epithelial repair output (Fig. 8D). The TA inflammatory-defense signature included increased *DUOX2* and *DUOXA2*, supporting epithelial oxidant antimicrobial defense; *LCN2*, supporting antimicrobial iron sequestration; *CXCL2*, *CXCL3*, *CXCL5*, and *CXCL8*, indicating epithelial chemokine-recruitment capacity; and *C3* and *CFB*, suggesting complement-associated inflammatory amplification.^50,52,64,160,163,229–234^ In parallel, increased *HSPA5* and *HSP90B1* support elevated chaperone and protein-folding demand, *XBP1* and *ERN1* support IRE1-XBP1 unfolded-protein-response activation, and *DDIT4* and *TRIB3* support ER-stress/integrated stress response (ISR) signaling.^17,174,235–240^ These shared ISC-TA inflammatory and ER-stress features align with experimental NEC evidence that epithelial TLR4 activation induces ISC ER stress.^17^ NEC TA cells therefore appear to expand repair output within an environment marked by antimicrobial defense, chemokine/complement signaling, and protein-folding pressure.

A coordinated hypoxia/glycolytic-metabolic response extended this stress pattern in NEC TA cells (Fig. 8E). Increased *HIF1A*, *VEGFA*, *EGLN3*, *ADM*, and *GDF15* point to low-oxygen and stress-response signaling, while higher *PFKP*, *ALDOA*, *GPI*, *PKM*, and *PDK1* support glycolytic adaptation in the proliferative compartment.^62,241–244^ Increased *SLC16A1/MCT1* suggests greater lactate/monocarboxylate transport capacity, and higher *NAMPT*, *ACSS2*, and *DDIT4* extend this response to nicotinamide adenine dinucleotide (NAD) salvage, acetate utilization, and stress-responsive mTOR regulation.^54,172,245–247^ NEC TA cells appear metabolically adapted to keep proliferating under hypoxic and inflammatory stress. This may support epithelial replacement but could also favor stress survival over efficient maturation into absorptive cells.

Active proliferative expansion was also evident in the DNA repair and cell-cycle features of NEC TA cells. Increased *TOP2A*, *MCM3*, *MCM10*, and *MKI67* support DNA replication/licensing and cycling-cell activity, while increased *E2F3*, *BRCA1*, and *CHEK2* point to S-phase progression, homologous recombination/repair, and DNA damage checkpoint regulation (Fig. 8F).^175,178,180,248–251^ This suggests that NEC TA cells are expanding epithelial output while managing DNA damage or replication stress from inflammation, hypoxia, oxidant stress, and rapid injury-associated proliferation.

Chromatin-regulatory changes in TA cells closely paralleled those observed in ISCs (Fig. 8G). Higher *DNMT1* and *HDAC1* suggest increased DNA methylation maintenance and histone deacetylation, while higher *CBX2* adds Polycomb-associated chromatin repression.^184,252–253^ Lower *SATB2* and *TET2*, also reduced in NEC ISCs, suggest reduced expression of regulators that may help stabilize epithelial differentiation and chromatin state control, while lower *MBD6* adds a TA-associated reduction in methylated-DNA/chromatin regulation.^194,254–255^ This pattern is consistent with neonatal mouse NEC multi-omics showing epithelial chromatin-accessibility changes and altered differentiation-associated programs.^157^ This regulatory state may allow TA proliferation to continue while limiting maturation into absorptive or secretory cells, producing repair that replaces cells without fully restoring mature barrier, transport, or crypt-villus function.

Reduced nuclear receptor and lineage-marker expression point to weaker regulatory input normally needed to guide epithelial maturation in NEC TA cells (Fig. 8H). Lower *NR3C2*, *NR1H4 (encoding FXR)*, *VDR*, *NR4A2*, *PPARA*, and *NR1I2 (encoding PXR)* point to reduced electrolyte/barrier regulation, bile acid and vitamin D signaling, inflammatory control, fatty acid metabolism, and xenobiotic sensing.^256–261^ Reduced nuclear receptor programs in NEC TA cells may limit maturation toward absorptive and detoxification function, because intestinal nuclear receptors normally help coordinate epithelial differentiation with inflammation, barrier function, metabolism, microbial-metabolite responses, and xenobiotic handling.^262^

Lower expression of absorptive-lineage genes provide further evidence of reduced epithelial maturation in NEC TA cells (Fig. 8I). Lower *SI*, *LCT*, and *MGAM* suggest reduced progression toward brush-border carbohydrate-digestion programs, including sucrase-isomaltase, lactase, and maltase-glucoamylase activity.^263–264^ Lower *ACE2* and *SLC6A19/*B0AT1 suggest reduced ACE2-dependent neutral amino-acid transport, while lower *SLC15A1*/PEPT1 suggests reduced intestinal dipeptide and tripeptide uptake.^265–267^ Reduced *CYP3A4*, *ABCG5*, and *ABCG8* further point to weaker xenobiotic/drug-metabolism and sterol-efflux programs.^268–271^ TA cells generate the epithelial cells that later become absorptive and secretory lineages, so the key question is whether their progeny can mature properly. In NEC, reduced absorptive-lineage genes suggest that TA-derived cells may have limited capacity to become functional absorptive epithelium, consistent with absorptive-lineage defects discussed later.

Of the 6,108 genes distinguishing fetal from infant control TA cells, 2,687 were also dysregulated in NEC versus infant controls, including 930 with fetal-like expression (34.6% of the 2,687 genes; Fig. 8J,K). These fetal-like maturity-associated genes pointed to reduced absorptive progression, epithelial signaling, transport, oxidative resilience, and repair capacity. Reduced expression of *SI* in NEC supports diminished brush-border disaccharidase expression linked to enterocyte differentiation and carbohydrate digestion.^263,272^ Decreased *ACE2* expression may reduce ACE2/B0AT1-dependent neutral amino-acid transport, while *HTR4* downregulation suggests weaker serotonin-responsive epithelial repair signaling.^265,273^ Similarly, lower *SLC5A9* in NEC suggests reduced sodium-coupled sugar/nutrient transport capacity, and *MSRA* downregulation may reduce oxidative-protein repair capacity.^274–275^ Together, these fetal-like maturation genes characterize NEC TA cells as having lower expression of programs associated with mature metabolic, transport, and repair capacity.

TF maturity analysis identified NEC-regulated transcription factors associated with epithelial differentiation, TGF-β/SMAD signaling, and chromatin state control (Fig. 8L). Higher *ELF3* expression in NEC TA cells was fetal-like and may reflect altered regulation of epithelial morphogenesis and differentiation, consistent with mouse *Elf3* studies.^276^ Higher *SKI* may weaken TGF-β/SMAD-dependent transcriptional restraint, which could reduce signals that help progenitors mature into epithelial lineages.^277–278^ Lower *DPF3* suggests reduced histone-mark reading and BAF-associated chromatin control, which could impair transcriptional coordination as TA cells transition toward epithelial differentiation.^195^ Overall, this fetal-like TF pattern suggests that NEC TA cells retain some regulatory features closer to the fetal reference while progressing less efficiently toward mature epithelial differentiation.

NEC TA cells occupy a stressed proliferative amplification state downstream of the ISC compartment. The ISC and TA findings support a crypt-to-villus model in which injury stress is already present at the regenerative source and is carried into active proliferation. Initiation may involve inflammatory, ER-stress, DNA repair, and chromatin-regulatory pressure in ISCs, while progression is reflected in TA-cell hypoxia/glycolysis, DNA repair demand, nuclear receptor loss, and reduced absorptive-lineage maturation. This means epithelial dysfunction may arise before terminal differentiation, as proliferative repair is generated under conditions that favor stress adaptation over efficient maturation. Resolution may depend on whether TA cells can shift from stressed amplification toward production of mature, transport-competent absorptive epithelium.

### Early Absorptive (EA) Enterocytes

Early absorptive (EA) enterocytes are the first differentiated absorptive epithelial cells emerging downstream of the ISC/transit-amplifying compartment and progressing toward mature villus enterocytes along the crypt-villus axis (Fig. 9A).^144^ Human microdissection studies support a crypt-to-villus increase in absorptive function, from relatively non-absorptive upper-crypt enterocytes to absorptive mid-villus enterocytes.^279^ Human single-cell profiling also resolves distinct absorptive epithelial states across the small intestine.^141^ Prior human NEC single-cell work identified inflammatory and antimicrobial changes in lower/mid-villus enterocytes, and neonatal mouse NEC multi-omics showed impaired absorptive differentiation, but neither resolved a distinct EA population.^18,157^ EA cells therefore provide an opportunity to ask whether injury-associated signals are present as crypt-derived cells begin absorptive differentiation.

**Fig. 9.**
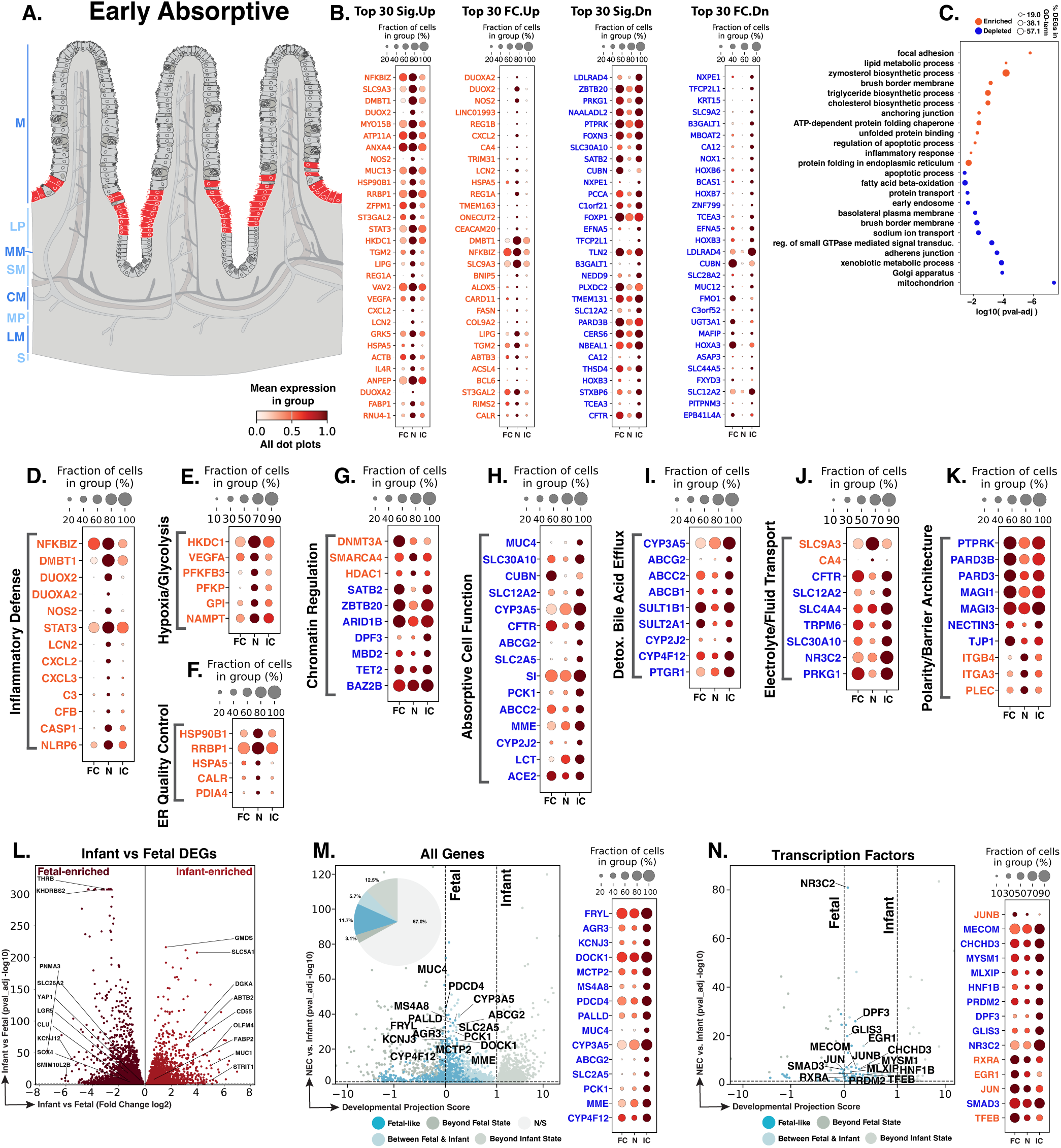
NEC early absorptive enterocytes show inflammatory and stress activation with incomplete absorptive maturation. **A.** Schematic showing the lower crypt-villus position of early absorptive enterocytes (red). Blue text to the left of the Panel A schematic indicates general anatomical regions: M, mucosa; LP, lamina propria; MM, muscularis mucosae; SM, submucosa; CM, circular muscle; MP, myenteric plexus; LM, longitudinal muscle; S, serosa. **B.** Dot plots of top NEC-specific genes ranked by significance and fold change. **C.** GO-term analysis showing enriched and depleted biological processes among NEC-associated DEGs. **D–K.** Curated gene panels showing inflammatory-defense, hypoxia/glycolysis, ER quality-control, chromatin-regulatory, absorptive-function, detoxification/bile acid efflux, electrolyte/fluid transport, and polarity/barrier-architecture programs. In panels **B** and **D–K**, orange gene text indicates upregulation in NEC relative to infant controls, whereas blue gene text indicates downregulation. **L.** Volcano plot of DEGs between fetal and infant controls that define the developmental trajectory for early absorptive cells. Triangles represent genes with extremely small adjusted P values reported as 0 because of numerical precision limits. Because −log10(0) is undefined, these genes were displayed at the upper y-axis limit with a triangle shape. **M–N.** Gene-level and transcription-factor maturity analyses of fetal-versus-infant DEGs, highlighting the subset also significantly changed in NEC relative to infant controls and positioned relative to fetal and infant reference states, and their corresponding dotplots. The developmental projection score in panels **M–N** is displayed on a symmetric logarithmic (symlog) scale. For expression dot plots, dot size indicates the fraction of nuclei expressing each gene and dot color indicates mean expression. In maturity plots, vertical dotted lines indicate the fetal and infant reference scores. Colored dots classify NEC-versus-infant DEGs as Fetal-like, Between Fetal & Infant, Beyond Fetal State, or Beyond Infant State; gray N/S dots are fetal-versus-infant DEGs without a significant NEC-versus-infant change.

In NEC EA cells, differential-expression analysis identified 3,764 genes significantly changed compared to infant controls, including 855 upregulated and 2,909 downregulated genes (Fig. 9B). The top NEC-associated genes suggested that EA cells retain epithelial inflammatory-defense and stress-response activity while expressing lower levels of genes associated with absorptive maturation, transport, xenobiotic handling, ion/fluid regulation, and epithelial surface function (Fig. 9B). GO-term analysis reinforced this split pattern, with enriched terms centered on inflammatory signaling, oxidant defense, ER protein-folding, chromatin regulation, hypoxia/metabolic stress, adhesion, and lipid/sterol biology, whereas depleted terms centered on brush-border and basolateral membrane organization, sodium transport, xenobiotic metabolism, Golgi function, and mitochondrial programs (Fig. 9C). These findings suggest that NEC EA cells carry injury-associated inflammatory and stress programs into the first differentiated absorptive stage, while showing reduced expression of genes needed for progression toward mature enterocyte function.

Building on the ISC/TA inflammatory-stress programs described above, NEC EA cells also upregulated inflammatory-defense genes previously associated with human NEC enterocytes, indicating that this response is carried into the first differentiated absorptive stage.^18^ Higher *NFKBIZ* suggests NF-κB-linked inflammatory feedback.^158,280^ Increased *DUOX2* and *DUOXA2* support epithelial oxidant defense, while higher *NOS2* adds nitric oxide-associated inflammatory stress.^50,160,281–284^ Higher *DMBT1* and *LCN2* support mucosal defense and antimicrobial iron sequestration.^161–162,229,285^ Increased *CXCL2* and *CXCL3* suggest neutrophil-chemoattractant capacity, while *C3* and *CFB* point to complement-associated inflammatory potential.^64,163,234,286^ Increased *CASP1* and *NLRP6* add inflammasome-associated epithelial stress, with *CASP1* most directly tied to cytokine processing and pyroptosis-associated priming.^114,287–289^ Together, the EA data suggest that inflammatory-defense activity persists as TA output enters absorptive differentiation, potentially amplifying epithelial inflammatory signaling before mature absorptive function is established.

Hypoxia-associated metabolic adaptation also extended into the EA compartment, but with a different gene pattern than in TA cells (Fig. 9E). In TA cells, NEC induced a broader low-oxygen program that included *HIF1A* itself, along with glycolytic, lactate-transport, NAD-salvage, acetate-utilization, and stress-responsive mTOR-regulatory genes. In EA cells, *HIF1A* was not a significant DEG, but downstream hypoxia/metabolic-associated genes were increased, including *VEGFA*, *HKDC1*, *PFKFB3*, *PFKP*, *GPI*, and *NAMPT*. Increased *HKDC1*, *PFKFB3*, *PFKP*, and *GPI* support glucose phosphorylation and glycolytic pathway activity.^290–294^ Increased *VEGFA* and *NAMPT* suggest low-oxygen/HIF-linked response signaling and NAD-salvage capacity during inflammatory or metabolic stress.^54,62,241,295^ This pattern suggests that hypoxia/metabolic stress is strongest in TA cells, where HIF1A itself is increased, but continues into EA cells through glycolytic, VEGFA, and NAD-salvage genes. This may help differentiating absorptive cells survive in an inflamed, low-oxygen environment, while also limiting the metabolic and transport programs needed for later absorptive function.

Protein-folding and ER quality control programs were increased in NEC EA cells (Fig. 9F). Upregulated *HSP90B1*, *RRBP1*, *HSPA5*, *CALR*, and *PDIA4* support increased ER protein-handling pressure through ER chaperone and protein-folding functions.^235–236,296^ Experimental mouse NEC links epithelial TLR4 signaling to ER stress and ISC apoptosis in intestinal crypts.^17^ The EA data suggest that related protein-folding pressure extends into differentiating absorptive cells. In NEC, increased ER quality control genes may help buffer protein-handling stress during early absorptive differentiation, but persistent ER stress could limit progression toward mature enterocyte function.

Chromatin-regulatory changes in NEC EA cells resembled those seen in ISCs and TA cells, but the implication is different because EA cells have already begun absorptive differentiation (Fig. 9G). In ISCs and TA cells, these changes may support injury-associated flexibility during repair and cell expansion. In EA cells, they suggest that an injury-associated regulatory state persists into cells that should be stabilizing enterocyte identity and activating mature absorptive programs. Higher *DNMT3A* may reflect altered de novo DNA methylation activity, potentially contributing to changes in intestinal epithelial transcriptional regulation.^191,297^ Higher *SMARCA4* and *HDAC1* are compatible with continued expression of chromatin-remodeling and histone-deacetylase machinery.^298–299^ Because *SMARCA4*/SMARCC1 help maintain an immature intestinal epithelial state and restrict premature maturation, this combination is compatible with NEC EA cells failing to fully consolidate a mature enterocyte program.^300^ By contrast, downregulated *SATB2*, *ZBTB20*, *ARID1B*, *DPF3*, *MBD2*, *TET2*, and *BAZ2B* may indicate reduced expression of regulatory genes associated with stable epithelial identity, differentiation, or chromatin state maintenance.^192,194–195,301–303^ These reductions may be important because enterocyte differentiation depends on enhancer programs that maintain epithelial identity.^304^ The pattern is consistent with injury-associated chromatin plasticity seen in neonatal mouse NEC and intestinal epithelial regeneration, suggesting that NEC EA cells remain regulatory flexible but incompletely mature.^157,200^

Reduced expression of mature absorptive-function genes was prominent in NEC EA cells, spanning brush-border digestion, nutrient handling, epithelial surface activity, metabolic flexibility, efflux, and chemical-defense/lipid-mediator metabolism (Fig. 9H). Lower *SI* and *LCT* indicate reduced sucrase-isomaltase and lactase-phlorizin hydrolase expression, supporting weaker brush-border carbohydrate-digestion potential.^305–308^ Lower *CUBN*, *SLC2A5*, *ACE2*, and *MME* point to reduced vitamin B12 uptake, fructose transport, ACE2/B0AT1-associated neutral amino acid transport, and epithelial surface peptidase activity.^309–316^ Lower *PCK1* and *MUC4* are consistent with reduced metabolic flexibility and epithelial surface function.^317–318^ Reduced *CYP3A5*, *CYP2J2*, *CYP4F12*, *SULT1B1*, *SULT2A1*, and *PTGR1* support lower xenobiotic, endogenous lipid-substrate, bile acid/hydroxysteroid, and lipid-mediator metabolic capacity, while reduced *ABCG2*, *ABCC2*, and *ABCB1* indicate lower mucosal ABC-transporter efflux capacity (Fig. 9I).^319–330^ Since dysbiosis can precede NEC and alter luminal metabolite and bile acid exposures, reduced chemical-defense and efflux genes may leave NEC EA cells less able to buffer these stresses during injury or repair.^331–336^ These findings suggest that NEC EA cells begin absorptive differentiation in an injury-stressed state with limited capacity for digestion, transport, detoxification, and luminal defense.

Electrolyte and fluid-transport genes showed an imbalanced pattern in NEC EA cells rather than broad transport loss (Fig. 9J). Upregulation of *SLC9A3* and *CA4* suggests increased apical sodium/proton exchange and membrane-associated carbonic anhydrase activity, potentially as a compensatory response to maintain epithelial pH and sodium absorption during inflammatory/metabolic stress.^337–340^ By contrast, lower *CFTR* and *SLC12A2* indicate reduced chloride-channel and NKCC1-associated chloride/fluid-handling programs.^341–344^ Lower *SLC4A4*, *TRPM6*, and *SLC30A10* further point to reduced bicarbonate-associated, magnesium, and manganese transport capacity, while lower *NR3C2* and *PRKG1* suggest reduced mineralocorticoid-receptor and cGMP-dependent kinase regulatory input into epithelial electrolyte transport.^72,345–351^ Higher *SLC9A3* and *CA4* may reflect a compensatory acid-base response to NEC-associated inflammatory/metabolic stress, supported by studies linking intestinal inflammation, LPS/acidosis, NHE activity, and carbonic anhydrase to epithelial pH regulation.^352–355^

Polarity and barrier-architecture genes suggested that NEC EA cells express lower levels of genes associated with apical-basal polarity and junctional organization, while selected genes linked to attachment of epithelial cells to the underlying basement membrane are higher (Fig. 9K). Reduced *PARD3* and *PARD3B* expression suggests weaker apical-basal polarity scaffolding.^356–358^ Lower *MAGI1*, *MAGI3*, and *TJP1* point to reduced tight junction-associated scaffolding capacity.^359–363^ NEC EA cells also expressed lower levels of *PTPRK* and *NECTIN3*, consistent with reduced epithelial adhesion and junctional organization.^77,364–365^ In contrast, higher *ITGB4*, *ITGA3*, and *PLEC* expression suggests increased basement membrane or hemidesmosome-associated attachment programs.^366–370^ This pattern suggests that NEC EA cells may retain or increase epithelial attachment features while expressing lower levels of genes needed for mature polarity and junctional architecture, potentially increasing vulnerability to barrier failure during NEC initiation, progression, and failed resolution.

Of the 7,305 genes distinguishing fetal from infant control EA cells, 2,410 were also dysregulated in NEC versus infant controls, including 852 with fetal-like expression (35.4% of the 2,410 genes; Fig. 9L,M). This fetal-like set overlapped with the reduced absorptive-function and chemical-defense genes described above, including lower expression of *MUC4*, *CYP3A5*, *ABCG2*, *SLC2A5*, *PCK1*, *MME*, and *CYP4F12* in NEC (Fig. 9H-I, M). *MUC4* supports epithelial surface function, *CYP3A5* and *CYP4F12* support xenobiotic and lipid-substrate metabolism, *ABCG2* supports epithelial efflux capacity, *SLC2A5* supports fructose transport, *PCK1* supports metabolic flexibility, and *MME* supports epithelial surface peptidase activity.^311,313,316–317,319–320,322–326,371^ Because these genes normally increase with postnatal absorptive maturation, their lower expression in NEC EA cells indicates reduced infant-like enterocyte maturation, with lower epithelial surface, nutrient-handling, metabolic, efflux, and chemical-defense capacity. This could increase vulnerability during NEC initiation by limiting postnatal barrier and detoxification maturation, during progression by reducing the ability of differentiating absorptive cells to buffer luminal and inflammatory injury, and during failed resolution by constraining transition into a mature, barrier-competent epithelium.

This fetal-like set also included lower expression of *FRYL*, *DOCK1*, *PALLD*, *AGR3*, *MS4A8*, *KCNJ3*, *MCTP2*, and *PDCD4* in NEC, extending the maturity pattern into epithelial morphogenesis, cytoskeletal organization, surface differentiation, ion/signaling, and translational or inflammatory control.^372–380^ Lower *PDCD4* expression in NEC is notable because the PDCD4 protein restrains eIF4A-dependent translation, and *Pdcd4* deletion in mice worsens experimental colitis through increased IL-6/STAT3 signaling and epithelial proliferation.^381–382^ These data suggest that NEC EA cells show a fetal-like maturity pattern during absorptive differentiation, including lower expression of absorptive, chemical-defense, epithelial-surface, morphogenetic, signaling, and regulatory programs that could weaken barrier formation, amplify injury propagation, and limit epithelial resolution.

TF maturity analysis reinforced this interpretation by identifying fetal-like regulatory genes (Fig. 9N). Higher expression of *JUN*, *JUNB*, and *EGR1* in NEC indicates a fetal-like immediate-early regulatory pattern in NEC EA cells.^383–385^ These factors are compatible with epithelial plasticity and rapid injury response, but sustained expression in differentiating EA cells could maintain a less mature, injury-responsive state that limits absorptive maturation and barrier stabilization. Lower expression of *SMAD3* and *MECOM* in NEC was fetal-like and linked the maturity pattern to altered TGF-β/SMAD-associated regulatory inputs.^386–388^ Altered *RXRA* and *NR3C2* link the maturity pattern to nuclear receptor and electrolyte-regulatory programs that may affect epithelial differentiation and transport physiology.^350,389–390^ Changes in *DPF3*, *MYSM1*, *PRDM2*, *HNF1B*, *MLXIP*, *CHCHD3*, and *TFEB* connect this fetal-like EA expression pattern to chromatin regulation, absorptive maturation, glucose-responsive metabolism, mitochondrial organization, and lysosomal/autophagy adaptation.^157,195,391–398^ These regulatory patterns suggest that NEC EA cells retain injury-responsive plasticity but show reduced postnatal absorptive maturation, increasing vulnerability to factors in the NEC environment.

NEC EA cells enter absorptive differentiation in a stressed state. Initiation may be promoted by early absorptive cells that add inflammatory-defense, oxidant/complement, hypoxia-associated glycolytic, ER protein-folding, inflammasome-associated, and altered chromatin-regulatory signals while expressing lower levels of genes needed for transport, detoxification, bile acid/efflux handling, electrolyte/fluid balance, polarity, junctional organization, epithelial-surface protection, and absorptive maturation. During NEC progression, this state could leave differentiating enterocytes poorly equipped to buffer luminal metabolites, bile acids, microbes, and inflammatory stress while retaining an incompletely matured, injury-responsive regulatory state. Resolution may require these cells to exit this injury-adapted state, restore mature absorptive and barrier architecture, and complete postnatal absorptive maturation. If that transition fails, the epithelium may continue to regenerate cells that remain functionally immature and barrier-vulnerable.

### Mature Absorptive (MA) Enterocytes

MA enterocytes represent the terminal absorptive stage downstream of EA cells, occupying the upper villus and villus-tip epithelium where brush-border digestion, nutrient and solute transport, lipid handling, barrier-facing functions, and xenobiotic metabolism are most fully expressed.^141–143,399^ (Fig. 10A). Whereas EA cells capture the point at which crypt-derived cells first enter absorptive differentiation, MA cells represent the mature lumen-facing epithelial surface exposed to luminal nutrients, microbes, bile acids, and inflammatory injury. Prior human NEC single-cell work reported enterocyte depletion, loss of villus-tip enterocytes, villus blunting, and inflammatory or antimicrobial gene induction in zonated enterocyte populations, and the experimental neonatal mouse multi-omics study discussed above similarly implicated impaired differentiated villus-enterocyte maturation.^18,157^ These studies did not separate EA and MA populations, so the transcriptional state of human MA cells in NEC remained unclear. We therefore analyzed MA cells as a distinct absorptive compartment to define how NEC affects the mature villus-facing epithelium, where defects in absorption, metabolism, detoxification, transport, and barrier support are likely to have direct disease relevance.

**Fig. 10.**
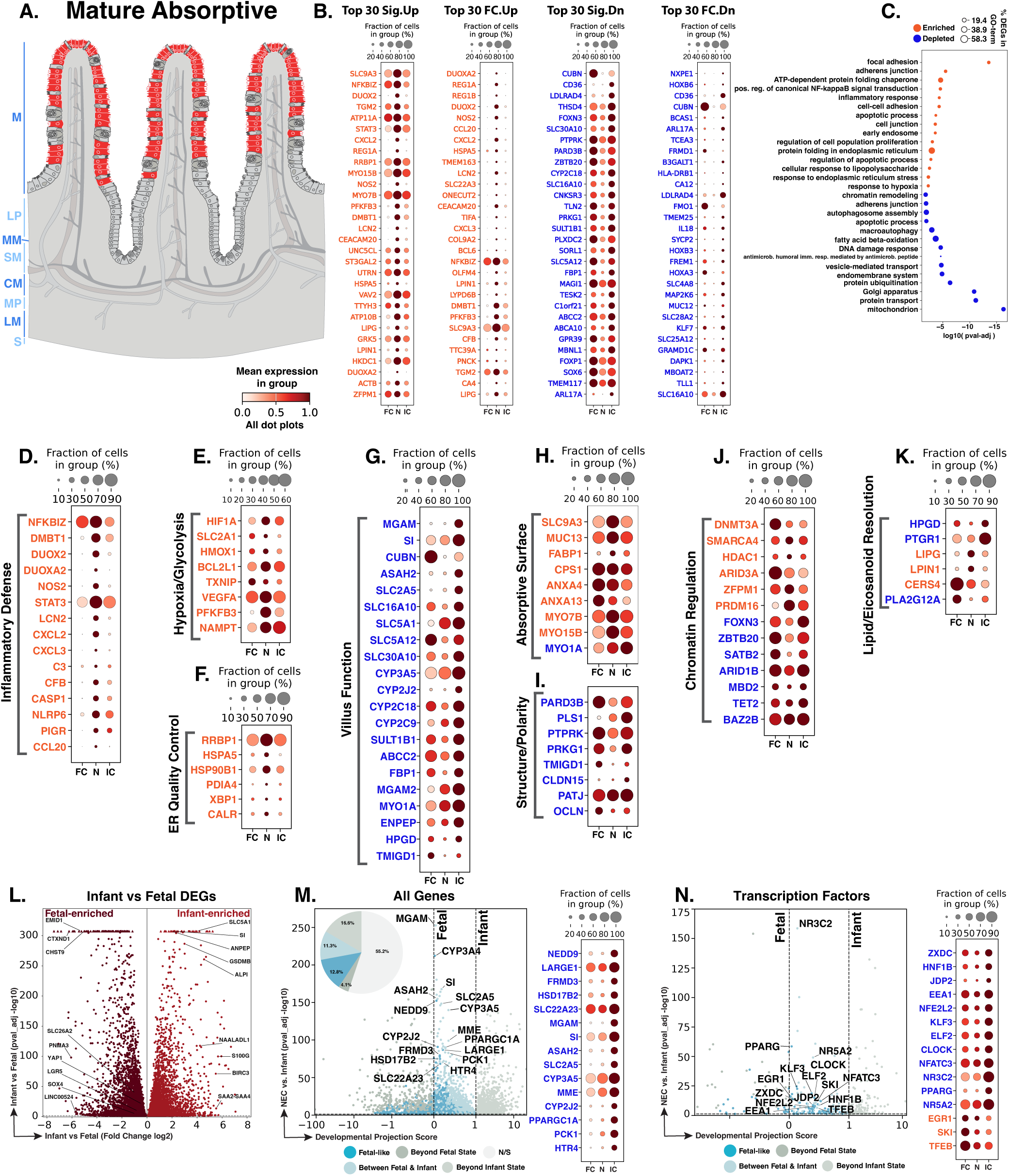
NEC mature absorptive enterocytes show villus-surface inflammatory stress with loss of terminal absorptive function. **A.** Schematic showing the upper villus and villus-tip location of mature absorptive enterocytes (red). Blue text to the left of the Panel A schematic indicates general anatomical regions: M, mucosa; LP, lamina propria; MM, muscularis mucosae; SM, submucosa; CM, circular muscle; MP, myenteric plexus; LM, longitudinal muscle; S, serosa. **B.** Dot plots of top NEC-specific genes ranked by significance and fold change. **C.** GO-term analysis showing enriched and depleted biological processes among NEC-associated DEGs. **D–K.** Curated gene panels showing inflammatory-defense, hypoxia/glycolysis, ER quality-control, villus-function, absorptive-surface, structure/polarity, chromatin-regulatory, and lipid/eicosanoid-resolution programs. In panels **B** and **D–K**, orange gene text indicates upregulation in NEC relative to infant controls, whereas blue gene text indicates downregulation. **L.** Volcano plot of DEGs between fetal and infant controls that define the developmental trajectory for mature absorptive cells. Triangles represent genes with extremely small adjusted P values reported as 0 because of numerical precision limits. Because −log10(0) is undefined, these genes were displayed at the upper y-axis limit with a triangle shape. **M–N.** Gene-level and transcription-factor maturity analyses of fetal-versus-infant DEGs, highlighting the subset also significantly changed in NEC relative to infant controls and positioned relative to fetal and infant reference states, and their corresponding dotplots. The developmental projection score in panels **M–N** is displayed on a symmetric logarithmic (symlog) scale. For expression dot plots, dot size indicates the fraction of nuclei expressing each gene and dot color indicates mean expression. In maturity plots, vertical dotted lines indicate the fetal and infant reference scores. Colored dots classify NEC-versus-infant DEGs as Fetal-like, Between Fetal & Infant, Beyond Fetal State, or Beyond Infant State; gray N/S dots are fetal-versus-infant DEGs without a significant NEC-versus-infant change.

We identified 8,598 MA nuclei across the atlas. In NEC MA cells, differential-expression analysis identified 6,718 genes significantly changed compared to infant controls, including 1,439 upregulated and 5,279 downregulated genes (Fig. 10B). Similar to the EA compartment, the top NEC-associated genes showed increased inflammatory-defense, adhesion/junctional, ER protein-folding, apoptosis-associated priming, hypoxia/glycolytic, and epithelial stress-response programs, together with reduced expression of genes associated with terminal absorptive maturation, lipid and mitochondrial metabolism, vesicle/endomembrane transport, Golgi function, autophagy, ubiquitin biology, and chromatin regulation (Fig. 10B). GO-term analysis supported this split pattern (Fig. 10C). Enriched terms involved focal adhesion, adherens junctions, NF-κB signaling, inflammatory response, cell-cell adhesion, apoptosis, early endosome biology, and ATP-dependent chaperone activity. Depleted terms involved chromatin remodeling, autophagy, fatty acid beta-oxidation, DNA damage response, vesicle and protein transport, protein ubiquitination, Golgi organization, and mitochondrial programs. These findings suggest that NEC MA cells retain villus-lineage identity but shift into an injury-associated state with inflammatory, adhesive, and ER-stress features, while expressing lower levels of genes needed for mature absorptive, metabolic, detoxification, transport, and epithelial-surface function.

The inflammatory-defense response observed in EA cells extended into NEC MA cells, with increased inflammatory, oxidant-defense, complement, chemokine, and inflammasome/pyroptosis-associated priming genes (Fig. 9D; Fig. 10D). The key MA distinction is that this inflammatory-defense state is present in mature absorptive cells, which normally support absorption, detoxification, and luminal barrier-facing functions. MA cells also showed inflammatory and immune-interface genes that were not significant in EA cells, including CXCL8, NFKBIA, TNFAIP3, IL1A, IL1B, IL15RA, IL12RB2, IL34, IFITM1, and IFI6, extending the MA response toward neutrophil recruitment, NF-κB feedback, IL-1/inflammasome-associated inflammatory signaling, immune cell crosstalk, and interferon-associated epithelial stress.^53,114,400–407^ The inflammatory-defense response seen in NEC EA cells is therefore maintained in MA cells and broadened by MA-selective cytokine, interferon, and immune-interface genes. In a population normally responsible for absorption, detoxification, and luminal barrier support, this could increase inflammatory signaling while reducing mature epithelial function.

Hypoxia/glycolytic and protein-folding stress responses also extended from EA into NEC MA cells, indicating that these injury-associated programs span the absorptive lineage. Shared upregulated genes included *VEGFA*, *PFKFB3*, *NAMPT*, *HMOX1*, *RRBP1*, *HSPA5*, *HSP90B1*, *PDIA4*, and *CALR*, supporting low-oxygen/metabolic stress, oxidative-response, and ER protein-handling pressure in both EA and MA cells.^54,62,235–236,241,291,295–296,408–412^ (Fig. 9E-F; Fig. 10E-F). The MA distinction was that *HIF1A*, *SLC2A1*, *BCL2L1*, *TXNIP*, and *XBP1* were significant in MA cells but not EA cells, pointing to a more explicit low-oxygen/glucose-uptake, survival/redox, and unfolded-protein-response signature at the terminal villus region.^101,413–416^ (Fig. 10E-F). This MA-enriched hypoxia/ER-stress response is consistent with prior NEC studies reporting loss of villus-tip/differentiated enterocytes and impaired absorptive maturation.^18,157^ These data suggest that metabolic and ER stress persists from EA into MA cells, with *HIF1A* upregulation marking a stronger low-oxygen response that may further weaken absorptive, detoxification, and barrier-facing functions.

Reduced expression of terminal villus digestive, transport, detoxification, lipid-metabolic, and metabolic-support genes was another prominent feature of NEC MA cells (Fig. 10G). Many of these downregulated genes were also reduced in EA cells, including *MGAM*, *SI*, *CUBN*, *SLC2A5*, *SLC16A10*, *SLC5A1*, *SLC5A12*, *SLC30A10*, *ASAH2*, *FBP1*, *CYP3A5*, *CYP2J2*, *CYP2C18*, *SULT1B1*, and *ABCC2*, indicating that loss of absorptive epithelial function spans both early and mature absorptive compartments (Fig. 9H-I; Fig. 10G). As described for EA cells, these genes support brush-border carbohydrate digestion, nutrient/solute and metal/ion transport, sphingolipid and metabolic support, xenobiotic/eicosanoid metabolism, sulfotransferase activity, and epithelial efflux capacity.^263,305–306,309–311,313,321–322,325,327,349,399,417–421^

Additional MA-selective losses involved genes that were not significant EA DEGs, including *MGAM2*, *MYO1A*, *ENPEP*, *CYP2C9*, *HPGD*, and *TMIGD1* (Fig. 10G). These MA-selective changes identify additional losses of terminal villus functions involving putative MGAM-like carbohydrate digestion, brush-border structure, epithelial peptidase activity, xenobiotic/eicosanoid metabolism, prostaglandin catabolism, and enterocyte adhesion/barrier-associated homeostasis.^263,399,422–426^ These changes could make MA cells less able to digest luminal nutrients, maintain brush-border structure, and buffer microbial or metabolic stress. Reduced carbohydrate-digestion capacity is especially relevant because carbohydrate maldigestion increases NEC incidence and severity in preterm pigs.^427^ Lower xenobiotic/bile acid handling and barrier-support genes may also increase vulnerability to dysbiosis-associated luminal stress.^331,334,428^ Overall, NEC-associated injury appears to reduce specialized MA functions needed for digestion, epithelial organization, detoxification, and protection at the luminal surface.

A mixed apical-surface response was evident in NEC MA cells, with increased selected response genes but loss of key structural genes needed for mature polarity, junctional organization, and brush-border structure. Shared EA/MA changes, including higher *SLC9A3* and lower *PRKG1*, *PARD3B*, and *PTPRK*, suggest persistent alterations in ion/fluid handling, polarity scaffolding, and epithelial contact programs (Fig. 9J-K; Fig. 10H-I).^72,77,337–338,356–358^ Increased *MUC13*, *FABP1*, *CPS1*, *ANXA4*, *ANXA13*, and *MYO7B* suggest activation of apical surface processes involving membrane mucins, lipid handling, metabolic activity, and membrane organization.^429–435^ MA cells also showed reductions that were not significant in EA cells, including *MYO1A*, *TMIGD1*, *CLDN15*, *OCLN*, and *PATJ* (Fig. 10H,I). Lower *MYO1A* and *TMIGD1* suggest weaker brush-border structure, intermicrovillar adhesion, and enterocyte structural maturation.^422,426,436–437^ Lower *CLDN15*, *OCLN*, and *PATJ* suggest impaired paracellular Na⁺ transport, tight junction regulation, and apical junctional organization.^438–440^ These MA-selective losses could weaken the mature absorptive surface, reducing barrier support and increasing exposure to luminal inflammatory stress during NEC.

Chromatin changes in NEC MA cells largely mirrored the EA regulatory state, whereas lipid/eicosanoid alterations revealed additional vulnerabilities of the more terminal villus epithelial zone (Fig. 9G; Fig. 10J-K). Shared chromatin-associated DEGs included increased *DNMT3A*, *SMARCA4*, *HDAC1*, *ARID3A*, *ZFPM1*, and *PRDM16*, together with reduced *FOXN3*, *ZBTB20*, *SATB2*, *ARID1B*, *MBD2*, *TET2*, and *BAZ2B*, supporting altered DNA methylation, chromatin accessibility, histone regulation, and transcriptional-state control across the absorptive lineage.^192,194–195,298–299,301–303^ The lipid/eicosanoid program also contained shared changes in *LIPG*, *LPIN1*, *CERS4*, and *PTGR1*, suggesting altered phospholipid hydrolysis, phosphatidic acid metabolism, sphingolipid biology, and lipid-mediator handling across EA and MA cells.^329–330,441–444^ By contrast, reduced *HPGD* and *PLA2G12A* were MA-selective changes, indicating additional loss of prostaglandin catabolism and epithelial phospholipase-associated lipid-defense capacity at the villus surface.^445–449^ These MA-specific lipid/eicosanoid changes could contribute to NEC progression by reducing epithelial capacity for lipid-mediator processing and maintenance of villus homeostasis. Taken together, these findings suggest that NEC MA cells retain the shared absorptive-lineage regulatory state present in EA cells while developing additional lipid-associated vulnerabilities that may impair epithelial stability and resolution of inflammatory injury.

Of the 11,130 genes distinguishing fetal from infant control MA cells, 4,993 were also dysregulated in NEC versus infant controls, including 1,449 with fetal-like expression (29.0% of the 4,993 genes; Fig. 10L,M). This fetal-like component was associated with lower expression of selected postnatal MA programs while broad villus identity was retained. The fetal-like genes overlapped with the terminal villus-function defects described above. Lower *MGAM* and *SI* support reduced brush-border carbohydrate digestion, lower *ASAH2* suggests altered sphingolipid metabolism, and lower *SLC2A5* suggests reduced fructose transport.^263,305–306,311,313,417–418^ Lower *CYP3A5* and *CYP2J2* point to reduced xenobiotic and endogenous lipid-substrate metabolism, while lower *MME* suggests reduced epithelial surface peptidase activity.^316,319–322^ Lower *PCK1* and *PPARGC1A* suggest reduced metabolic flexibility and mitochondrial support, and lower *HTR4* suggests altered serotonin-responsive epithelial signaling.^211,317,450–451^ Overall, NEC MA cells retain broad terminal villus identity but show lower expression of postnatal digestive, metabolic, detoxification, and signaling genes. Compared with EA cells, where maturity changes suggest incomplete progression toward absorptive identity, MA maturity changes highlight reduced specialized terminal villus functions needed to maintain epithelial physiology during NEC.

TF maturity analysis reinforced that NEC MA cells show fetal-like or intermediate regulatory features (Fig. 10N). Lower *NR3C2*, *PPARG*, and *NR5A2* suggest reduced nuclear receptor input into epithelial ion/fluid control, lipid metabolism, anti-inflammatory signaling, and intestinal differentiation.^256,452–453^ Higher EGR1 and SKI are consistent with stress-responsive immediate-early regulation and modulation of TGF-β/SMAD-dependent transcription.^454–455^ Higher TFEB is consistent with enhanced lysosomal/autophagy regulation, whereas lower NFE2L2 suggests reduced NRF2-mediated antioxidant defense in stressed MA cells.^456–457^ Changes in CLOCK, HNF1B, and NFATC3 extend this pattern to circadian-metabolic regulation, enterocyte gene regulation, and calcium-dependent transcriptional signaling.^394,458–459^ These TF changes suggest that NEC MA cells retain selected adaptive stress responses but may have limited antioxidant protection and reduced regulatory support for mature absorptive villus physiology.

NEC MA cells retain villus-lineage identity but show broad inflammatory, hypoxia/metabolic, ER-stress, and absorptive-function defects. Initiation may be favored when the mature villus surface has reduced digestive, detoxification, lipid-handling, polarity, brush-border, and barrier-support functions, leaving it less able to manage nutrients, microbial products, bile acids, and inflammatory exposure. Progression could be sustained by persistent villus-surface injury, reduced epithelial detoxification, and impaired luminal barrier control. Resolution may fail if MA cells cannot restore terminal villus physiology even when villus identity remains present. This creates a mature epithelial surface that is recognizable by lineage but less capable of protecting the intestine during recovery.

### Goblet Cells

Goblet cells are secretory epithelial cells distributed along the crypt-villus axis (Fig. 11A).^141–142^ In the human intestine, goblet-cell abundance increases along the proximal-to-distal axis, with estimates of approximately 4% of epithelial cells in the duodenum, 6% in the jejunum, 12% in the ileum, and 16% in the distal colon.^460–461^ They produce mucins, including MUC2 in small-intestinal goblet cells, and regulate region-specific mucin glycosylation and sialylation programs that establish the mucus barrier separating luminal microbes from the epithelial surface.^145,462–464^ This compartment is highly relevant to NEC because impaired mucus-barrier function may increase microbial contact with the immature epithelium and amplify mucosal inflammation during injury.^464–465^ Human NEC studies have implicated goblet-cell dysfunction through reduced goblet-cell abundance despite retained MUC2 and TFF3 protein expression in acute NEC tissue and altered *REG4* expression in NEC-associated goblet cells.^466–467^

**Fig. 11.**
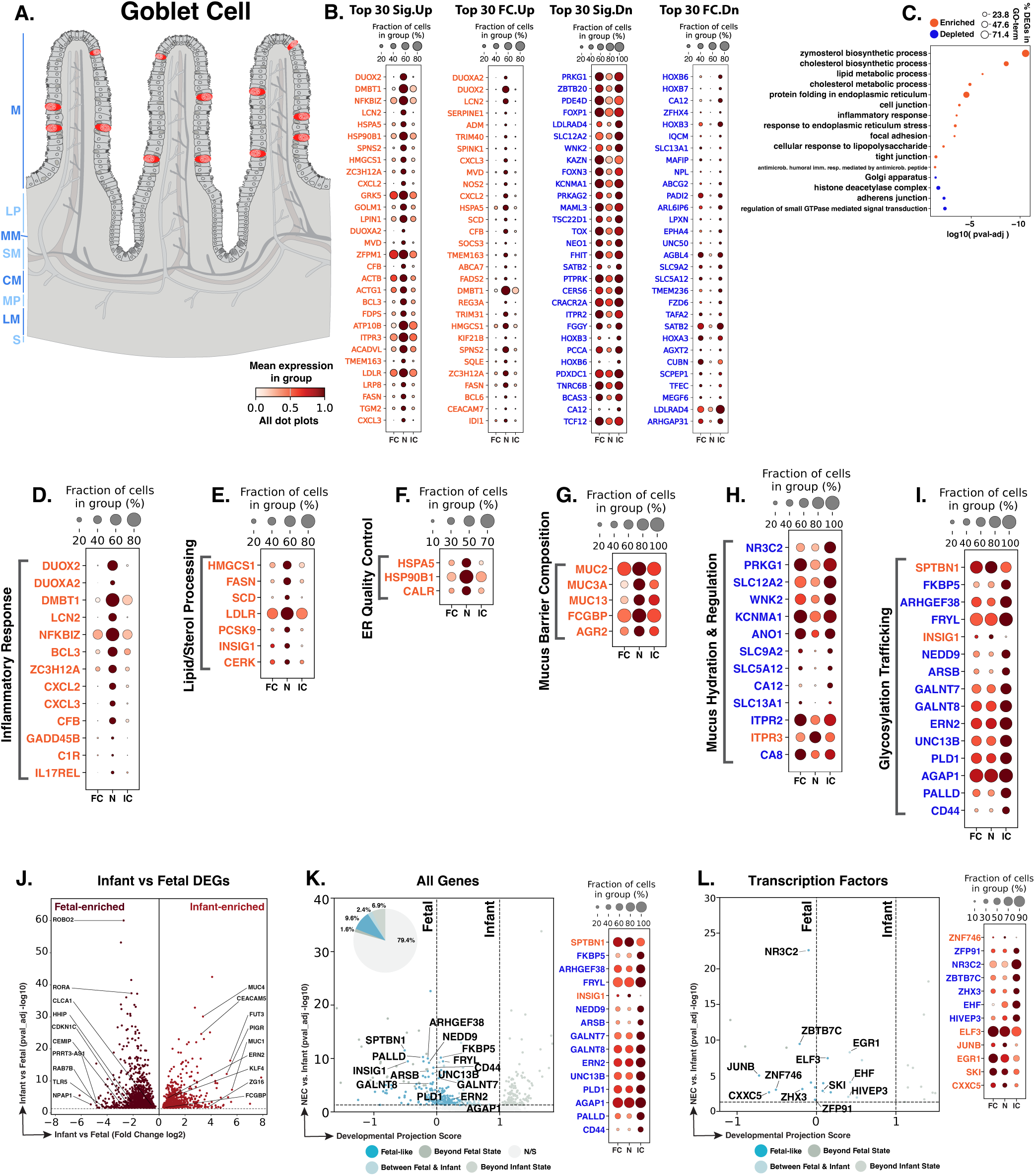
NEC goblet cells show mucus-barrier transcription with impaired secretory support and developmental displacement. **A.** Schematic showing goblet-cell distribution (red) along the crypt-villus axis. Blue text to the left of the Panel A schematic indicates general anatomical regions: M, mucosa; LP, lamina propria; MM, muscularis mucosae; SM, submucosa; CM, circular muscle; MP, myenteric plexus; LM, longitudinal muscle; S, serosa. **B.** Dot plots of top NEC-specific genes ranked by significance and fold change. **C.** GO-term analysis showing enriched and depleted biological processes among NEC-associated DEGs. **D–I.** Curated gene panels showing inflammatory-response, lipid/sterol processing, ER quality-control, mucus-barrier composition, mucus hydration/ion regulation, and glycosylation/trafficking programs. In panels **B** and **D–I**, orange gene text indicates upregulation in NEC relative to infant controls, whereas blue gene text indicates downregulation. **J.** Volcano plot of DEGs between fetal and infant controls that define the developmental trajectory for goblet cells. **K–L.** Gene-level and transcription-factor maturity analyses of fetal-versus-infant DEGs, highlighting the subset also significantly changed in NEC relative to infant controls and positioned relative to fetal and infant reference states, and their corresponding dotplots. The developmental projection score in panels **K–L** is displayed on a symmetric logarithmic (symlog) scale. For expression dot plots, dot size indicates the fraction of nuclei expressing each gene and dot color indicates mean expression. In maturity plots, vertical dotted lines indicate the fetal and infant reference scores. Colored dots classify NEC-versus-infant DEGs as Fetal-like, Between Fetal & Infant, Beyond Fetal State, or Beyond Infant State; gray N/S dots are fetal-versus-infant DEGs without a significant NEC-versus-infant change.

Experimental studies further support goblet-cell vulnerability in NEC, showing that epithelial TLR4 signaling suppresses goblet-cell differentiation and that TNF-alpha/TNFR1-dependent mucus depletion in immature intestine may increase NEC susceptibility.^465,468^ Prior human NEC single-cell work lacked sufficient goblet cells for DEG analysis, leaving this lineage unresolved in human NEC tissue, while experimental neonatal mouse NEC multi-omics identified goblet-cell-associated changes in glycosylation, WNT/beta-catenin, hypoxia, and inflammatory-response programs.^18,157^ We therefore examined human NEC goblet cells to determine whether this lineage maintains mucus-barrier identity or acquires injury-associated programs involving inflammatory defense, secretory function, ER quality control, glycosylation, trafficking, adhesion, ion/fluid regulation, and developmental maturity.

We identified 1,233 goblet-cell nuclei across the atlas. In NEC goblet cells, differential-expression analysis identified 959 genes significantly changed relative to infant controls, including 400 upregulated and 559 downregulated genes (Fig. 11B). The top DEGs pointed to a secretory injury state: inflammatory-response, lipid/sterol biosynthetic, ER quality control, mucus-barrier, and scaffold-associated genes were increased, whereas glycosylation, adhesion, ion/fluid-regulatory, and maturation-associated genes were reduced (Fig. 11B). GO-term enrichment supported this split pattern, with enriched lipid/cholesterol biosynthesis, ER protein-folding/stress, inflammatory and antimicrobial-response, focal adhesion, tight junction, and cell junction terms, and depleted Golgi, adherens junction, histone-deacetylase-complex, and small-GTPase regulatory terms (Fig. 11C). Overall, NEC goblet cells increase inflammatory and secretory-stress responses while showing reduced support for glycosylation, Golgi processing, vesicle trafficking, adhesion, and junctional organization, changes that could affect mucus assembly, hydration, trafficking, and positioning at the injured epithelial surface.

Inflammatory-defense features seen in absorptive lineages were also retained in NEC goblet cells, with increased *DUOX2*, *DUOXA2*, *DMBT1*, *LCN2*, *NFKBIZ*, *BCL3*, *CXCL2*, *CXCL3*, and *CFB* (Fig. 11D). These genes point to oxidant defense, antimicrobial activity, NF-κB-associated inflammatory regulation, chemokine signaling, and complement activity across epithelial compartments.^160,162–163,234,286,469–472^ This overlap is consistent with inflammatory-response enrichment reported in experimental mouse neonatal NEC goblet cells.^157^ Goblet cells also showed lineage-associated increases in *GADD45B*, *C1R*, and *IL17REL*, suggesting stress-survival, complement, and cytokine-regulatory activity in mucus-producing cells.^473–476^ These findings suggest that NEC goblet cells share the broader epithelial inflammatory-defense response while adding goblet-cell-specific regulatory features that could affect mucus-barrier function, microbial interactions, and epithelial immune signaling.

Lipid/sterol biosynthesis genes were also upregulated in NEC goblet cells and overlapped with those observed across epithelial progenitor and absorptive compartments, including *HMGCS1*, *FASN*, *SCD*, *LDLR*, *PCSK9*, and *INSIG1* (Fig. 11E). This shared response points to increased lipid supply, membrane-lipid production, and cholesterol regulation across NEC epithelial lineages.^477–483^ In goblet cells, lipid/sterol upregulation may support membrane supply for mucus secretion and increase capacity for FASN-dependent MUC2 palmitoylation, a modification linked to proper MUC2 secretion and mucus-barrier function.^484–485^ *CERK* was significant in goblet cells but not in other epithelial lineages, adding a goblet-cell-enriched sphingolipid-signaling feature to the NEC response. CERK and its product ceramide-1-phosphate can support inflammatory agonist-responsive prostaglandin synthesis and cPLA2α membrane recruitment.^486–487^ Altogether, lipid/sterol upregulation may support membrane supply for mucus secretion, while *CERK* suggests a goblet-enriched lipid signal that could influence inflammation at the mucus barrier.

Increased ER quality control genes provided another feature of NEC goblet cells, consistent with secretory stress in a mucus-producing lineage. Increased *HSPA5*, *HSP90B1*, and *CALR* were shared across epithelial compartments, suggesting that the ER-stress and protein-folding response seen in absorptive cells also extends into goblet cells (Fig. 11F). These genes encode ER proteins involved in unfolded-protein responses, ER-resident protein folding, calcium buffering, and calreticulin/calnexin-cycle chaperone function.^235–236,488^ This response is especially relevant in goblet cells because mucus-barrier maintenance depends on sustained MUC2 synthesis, ER-dependent folding, and calcium-regulated mucin secretion.^489–491^ These findings suggest that NEC goblet cells increase ER quality control capacity, but persistent MUC2 folding stress could still impair mucus production and weaken mucus-barrier function.^489–490^

Core mucus and epithelial-surface genes were increased in NEC goblet cells, overlapping with mucin and surface-barrier features seen in absorptive compartments. Human intestinal single-cell profiling shows region- and lineage-specific mucin expression, including secreted *MUC2* in goblet cells and transmembrane mucins such as *MUC13* across epithelial populations.^141^ In NEC goblet cells, higher *MUC2* and *FCGBP* suggest increased expression of the MUC2-associated mucus network.^464,492–493^ Higher *MUC3A* and *MUC13* point to increased epithelial-surface mucin expression at the luminal interface, and higher *AGR2*, which encodes an ER-resident mucin-folding protein, suggests greater support for MUC2 production and goblet-cell mucus formation.^429–430,494^ This pattern helps reconcile prior NEC studies reporting reduced goblet-cell abundance or mucus depletion, because goblet cells may be lost or impaired in NEC tissue, while the goblet cells captured here show increased mucus-barrier gene expression.^465–466,468^ Whether this response can restore an effective mucus barrier likely depends on coordinated glycosylation, trafficking, ER secretory capacity, calcium-linked secretion, and ion/fluid regulation (Fig. 11H-I).

Lower expression of ion, fluid, and pH-regulatory genes was another feature of NEC goblet cells, overlapping with transport defects described in absorptive lineages (Fig. 11H). Lower *NR3C2*, *PRKG1*, *SLC12A2*, *SLC9A2*, *SLC5A12*, *CA12*, *SLC13A1*, *KCNMA1*, and *WNK2* suggest reduced mineralocorticoid-associated sodium regulation, cGMP signaling, chloride/fluid transport, sodium/proton exchange, monocarboxylate and sulfate transport, bicarbonate/pH control, and potassium-channel regulation.^72,256,337,495–500^ In goblet cells, these changes are important because small-intestinal mucus release depends on coordinated goblet-cell secretion, calcium/IP3 receptor signaling, epithelial ion transport, and luminal fluid movement.^501^ NEC goblet cells also showed a mixed IP3 receptor pattern, with lower *ITPR2* but higher *ITPR3*; these genes encode ER calcium-release channels that help couple intracellular signaling to mucus secretion.^501^ Lower *CA8* may further affect calcium regulation because CA8 binds IP3R1 and inhibits IP3R1-mediated Ca²⁺ signaling, helping control intracellular calcium homeostasis.^502–503^ Overall, NEC goblet cells increase mucus-barrier transcripts but show lower expression of ion/fluid/pH and calcium-regulatory genes needed to hydrate mucus, coordinate secretion, and organize the mucus barrier. This combination could weaken microbial separation from the epithelium and delay restoration of an organized mucus barrier.

Higher scaffold-associated gene expression, together with lower glycosylation, vesicle-trafficking, adhesion, and restitution-associated genes, extended epithelial structural and trafficking changes seen in absorptive lineages into NEC goblet cells (Fig. 11I). In goblet cells, these changes are important because mucus-barrier function depends not only on mucin transcripts, but also on mucin glycosylation, granule delivery, secretion, and integration with the epithelial surface. Higher *SPTBN1* suggests altered βII-spectrin scaffolding at the apical/secretory surface, while lower *FKBP5* points to reduced HSP90 co-chaperone stress-buffering capacity.^504–505^ Lower *GALNT7* and *GALNT8* suggest reduced mucin-type O-glycosylation capacity; lower *UNC13B* and *PLD1* point to weaker vesicle priming and secretory-vesicle fusion; and lower *NEDD9*, *PALLD*, and *CD44* suggest reduced adhesion, actin-cytoskeletal support, and epithelial restitution.^506–512^ Lower *ARSB* was specific to goblet cells and suggests reduced arylsulfatase B activity, which could increase 4-sulfated glycosaminoglycan features at the mucus interface.^513–514^ This may matter because higher 4-chondroitin-sulfate content strengthens MUC2 binding, and chondroitin-sulfate-MUC2 interactions increase mucus viscoelasticity and reduce mucus permeability.^515^ Overall, NEC goblet cells increase mucus-barrier transcripts but express lower levels of genes needed to glycosylate mucins, deliver secretory vesicles, maintain epithelial attachment, and organize mucus structure. Lower *ARSB* may also make the mucus network more tightly associated with chondroitin sulfate, which could increase mucus stiffness and reduce permeability, potentially making mucus harder to distribute or renew when secretion, hydration, and trafficking genes are also reduced.

Of the 1,714 genes distinguishing fetal from infant control goblet cells, 353 were also dysregulated in NEC versus infant controls, including 165 with fetal-like expression (46.7% of the 353 genes; Fig. 11J,K). This fetal-like component reinforced the patterns described above. Higher *SPTBN1* and lower *FKBP5* suggest persistent cytoskeletal-scaffold features alongside altered stress-response regulation. Lower *GALNT7*, *GALNT8*, *UNC13B*, *PLD1*, *NEDD9*, *PALLD*, and *CD44* point to reduced postnatal mucin glycosylation, secretory trafficking, adhesion, and epithelial repair capacity.^504–512^ Lower *ARSB* adds a mucus-structure implication, because reduced arylsulfatase B could increase 4-sulfated glycosaminoglycan features that strengthen MUC2-chondroitin-sulfate interactions, increase mucus viscoelasticity, and reduce permeability.^513,515^ Taken together, the maturity data suggest that NEC goblet cells retain secretory-lineage identity but show less mature expression of genes needed to build, organize, and restore an effective mucus barrier. This fetal-like expression may reflect persistent developmental immaturity or injury-associated reversion, either of which could limit mucus-barrier maturation and recovery.

The TF maturity panel supported this interpretation, identifying fetal-like regulatory features linked to stimulus-response signaling, TGF-β/SMAD regulation, and epithelial fluid control (Fig. 11L). Higher *EGR1* and *JUNB* suggest persistent immediate-early/AP-1 activity, while higher *SKI* suggests increased inhibition of TGF-β/SMAD signaling that could reduce responsiveness to epithelial repair cues.^277,454–455,516–517^ Lower *NR3C2* suggests weaker mineralocorticoid-linked sodium/fluid regulation, consistent with the reduced ion/fluid gene pattern described above.^256^ These TF changes suggest that NEC goblet cells may retain injury-responsive regulation while expressing less of the regulatory machinery needed for coordinated mucus secretion, hydration, and mucus-barrier recovery.

NEC goblet cells retain secretory identity and increase inflammatory-defense, lipid/sterol, ER quality control, and mucus-barrier transcripts. Initiation may be influenced by reduced glycosylation, trafficking, calcium-linked secretion, ion/fluid regulation, adhesion, and mucus-organization genes that weaken mucus hydration and microbial separation. During NEC progression, goblet cells may continue trying to reinforce the barrier while losing the machinery needed to assemble, deliver, and organize mucus effectively. Resolution may depend on whether mucus production becomes coordinated with glycosylation, secretion, hydration, and epithelial integration. Failed resolution may occur when goblet cells remain secretory but cannot rebuild a mature, hydrated, structurally organized mucus barrier.

### Enteroendocrine Cells (EECs)

EECs are rare hormone-producing epithelial cells distributed throughout the intestinal epithelium, where they are estimated to comprise roughly 1% of epithelial cells and include diverse hormone-secreting subtypes (Fig. 12A).^49,141–142,518^ EECs sense luminal nutrients and can relay sensory information to nerves through neuropod-mediated epithelial-neural circuits, linking this lineage to nutrient sensing, hormone signaling, motility, and epithelial-neural communication in the injured neonatal intestine.^519–521^ NEC-specific EEC literature is limited, but EEC-derived hormone pathways are relevant because intestinal L cells produce GLP-2, GLP-2 promotes epithelial growth, and exogenous GLP-2 was protective in experimental rat NEC while delaying but not preventing NEC onset in preterm pigs.^522–525^ Prior human NEC single-cell work identified EECs but did not resolve a dedicated EEC DEG or hormone-gene response, whereas experimental neonatal mouse NEC multi-omics reported increased EECs with enrichment for peptide transport, synapse transport, axon guidance, and feeding-response regulation.^18,157^ We therefore examined whether human NEC EECs retain hormone-lineage identity while acquiring transcriptional changes that could affect epithelial defense, metabolic adaptation, secretory trafficking, and epithelial-neural communication.

**Fig. 12.**
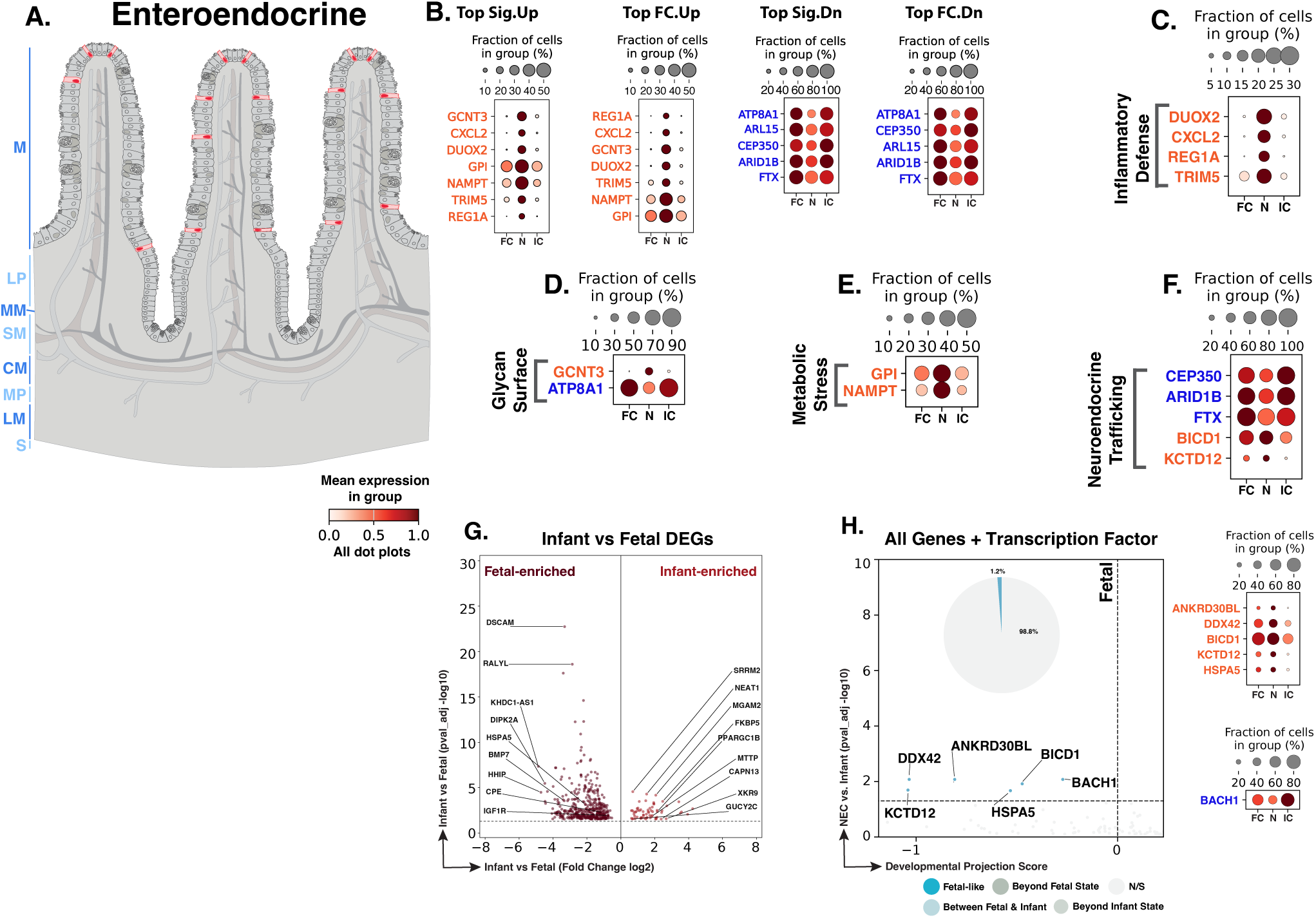
NEC enteroendocrine cells show a restricted injury-associated program without broad hormone-gene loss. **A.** Schematic showing enteroendocrine-cell distribution (red) along the intestinal epithelium. Blue text to the left of the Panel A schematic indicates general anatomical regions: M, mucosa; LP, lamina propria; MM, muscularis mucosae; SM, submucosa; CM, circular muscle; MP, myenteric plexus; LM, longitudinal muscle; S, serosa. **B.** Dot plots of top NEC-specific genes ranked by significance and fold change. **C–F.** Curated gene panels showing inflammatory-defense, glycan/surface, metabolic-stress, and neuroendocrine-trafficking programs. In panels **B–F**, orange gene text indicates upregulation in NEC relative to infant controls, whereas blue gene text indicates downregulation. **G.** Volcano plot of DEGs between fetal and infant controls that define the developmental trajectory for enteroendocrine cells. **H.** Combined gene-level and transcription-factor maturity analysis of fetal-versus-infant DEGs, highlighting the subset also significantly changed in NEC relative to infant controls and factors positioned relative to fetal and infant reference states. The developmental projection score in panel **H** is displayed on a symmetric logarithmic (symlog) scale. Top dot plot (All Genes), Bottom dot plot (Transcription Factors). For expression dot plots, dot size indicates the fraction of nuclei expressing each gene and dot color indicates mean expression. In maturity plots, vertical dotted lines indicate the fetal and infant reference scores. Colored dots classify NEC-versus-infant DEGs as Fetal-like, Between Fetal & Infant, Beyond Fetal State, or Beyond Infant State; gray N/S dots are fetal-versus-infant DEGs without a significant NEC-versus-infant change.

The NEC-associated EEC DEG profile was more restricted than in the major absorptive and goblet populations, consistent with the rarity of this lineage in intestinal single-cell atlases and the smaller number of captured EEC nuclei in this dataset (Fig. 12B).^49,141–142^ GO-term enrichment analysis was performed, but no terms reached significance, likely reflecting the limited number of significant NEC-vs-infant DEGs. We therefore interpreted the EEC response from the significant DEGs using the same curated framework applied to prior epithelial lineages (Fig. 12B). These genes pointed to increased epithelial defense, metabolic stress, and glycan/surface-interface features, together with reduced membrane-organization, cytoskeletal, chromatin-regulatory, and non-coding regulatory inputs. Differences in canonical EEC hormone genes did not reach significance, indicating that the main NEC-associated EEC pattern was not broad loss of hormone-lineage identity but a focused injury-associated response affecting epithelial defense, surface regulation, metabolic adaptation, and epithelial-neural support programs.

We identified 623 EEC nuclei across the atlas. In NEC EECs, differential-expression analysis identified 20 genes significantly changed relative to infant controls, including 13 upregulated and 7 downregulated genes (Fig. 12B). Although this was a smaller DEG set than in major epithelial lineages, the strongest changes pointed to inflammatory defense, surface-glycan regulation, and membrane organization. Higher *DUOX2* and *CXCL2* suggest that EECs may contribute to epithelial oxidant defense and neutrophil-recruitment signaling.^50,160,163^ Higher *REG1A* and *TRIM5* further support an injury-defense and innate-immune response.^526–527^ Higher *GCNT3* suggests altered mucin-type O-glycan branching at the EEC surface, while lower *ATP8A1* points to reduced membrane lipid asymmetry and surface organization.^528–530^ These patterns suggest that NEC EECs share the broader epithelial inflammatory-defense response seen in absorptive and goblet lineages, while also showing surface-glycan and membrane-organization changes that could affect luminal sensing, epithelial polarity, and epithelial-neural communication.

NEC EECs showed metabolic-stress and trafficking changes, similar to those in absorptive and goblet lineages, but with distinct implications for rare sensory epithelial cells. Higher *GPI* and *NAMPT* suggest glycolytic and NAD-salvage support during stress. Lower *CEP350* suggests reduced microtubule anchoring, while higher *BICD1*, also seen in other epithelial compartments, suggests changes in dynein-associated vesicle transport.^54,531–534^ Lower *ARID1B* suggests reduced SWI/SNF/BAF chromatin-regulatory support, which is relevant because EEC and secretory-lineage differentiation depend on chromatin accessibility and enhancer-regulatory programs.^535–537^ Higher *KCTD12*, which encodes a GABA-B receptor auxiliary protein, was detected only in EECs among the epithelial lineages compared, suggesting altered inhibitory neuroactive signaling that could affect EEC responsiveness to neural or epithelial-neural cues.^538^ Overall, NEC EECs retain endocrine identity but show injury-stress, trafficking, chromatin-regulatory, and neuroactive-signaling changes that could affect epithelial-neural communication in the injured neonatal intestine.

Of the 491 genes distinguishing fetal from infant control EECs, 6 were also dysregulated in NEC versus infant controls, including 6 with fetal-like expression (100.0% of the 6 genes; Fig. 12G,H). This limited overlap suggests that NEC EECs largely retain postnatal hormone-lineage identity, but with a small fetal-like set linked to stress handling, vesicle transport, RNA processing, and neuroactive signaling. Higher *BICD1* and *KCTD12* connect this set to dynein-associated vesicle transport and GABA(B)-receptor auxiliary signaling, while higher *HSPA5* and *DDX42* point to increased ER chaperone activity and RNA-processing support during stress.^534,539–544^ Lower *BACH1* may relieve repression of *HMOX1* and permit NRF2-dependent antioxidant transcription during oxidative stress.^545^ Together, these maturity data suggest that NEC EECs retain broad postnatal endocrine identity, while a small fetal-like gene set may alter stress adaptation, vesicle trafficking, antioxidant regulation, and GABA(B)-linked epithelial-neural signaling.

NEC EECs retain broad hormone-producing and sensory identity but show a focused injury-associated response, not widespread hormone-lineage loss. Initiation may be influenced by epithelial-defense, glycan/surface, and metabolic-stress changes that alter how EECs sense and respond to the NEC environment. During progression, altered trafficking and neuroactive-signaling genes could affect epithelial-neural communication, local sensory signaling, and coordination with motility or repair pathways. Resolution may require restoration of polarized sensory and secretory function within the injured epithelium. If EECs remain in a stress-adapted state, they may be less able to support normal epithelial-neural communication during recovery.

### Paneth Cells

Paneth cells are specialized secretory epithelial cells at the small-intestinal crypt base, where they represent an estimated 3%-8% of the small-intestinal epithelium, are enriched within crypts, produce antimicrobial granules, and, in mouse intestine, help support neighboring LGR5+ intestinal stem cells.^150,546–548^ Whether Paneth cells provide the same stem-cell niche support in humans is less clear. Human single-cell profiling shows important species differences, including limited expression of expected mouse Paneth niche factors in human Paneth cells.^141^ Paneth cell disruption is relevant to NEC because immature mouse models show that Paneth cell ablation with dithizone, followed by enteral *Klebsiella pneumoniae*, causes small-intestinal injury and inflammatory activation resembling NEC.^549^ Subsequent work showed that this Paneth cell disruption model requires live bacteria and occurs independently of TLR4 signaling.^550^

Existing human NEC single-cell studies have not resolved Paneth cells as a dedicated population. Experimental neonatal mouse NEC multi-omics reported decreased Paneth cell representation and implicated immature Paneth cells with reduced Wnt secretion in impaired ISC-niche stability and epithelial regeneration.^157^ We therefore examined human NEC Paneth cells to determine whether they retain antimicrobial secretory identity while acquiring injury-associated changes that could affect crypt-base function and epithelial recovery (Fig. 13A).

**Fig. 13.**
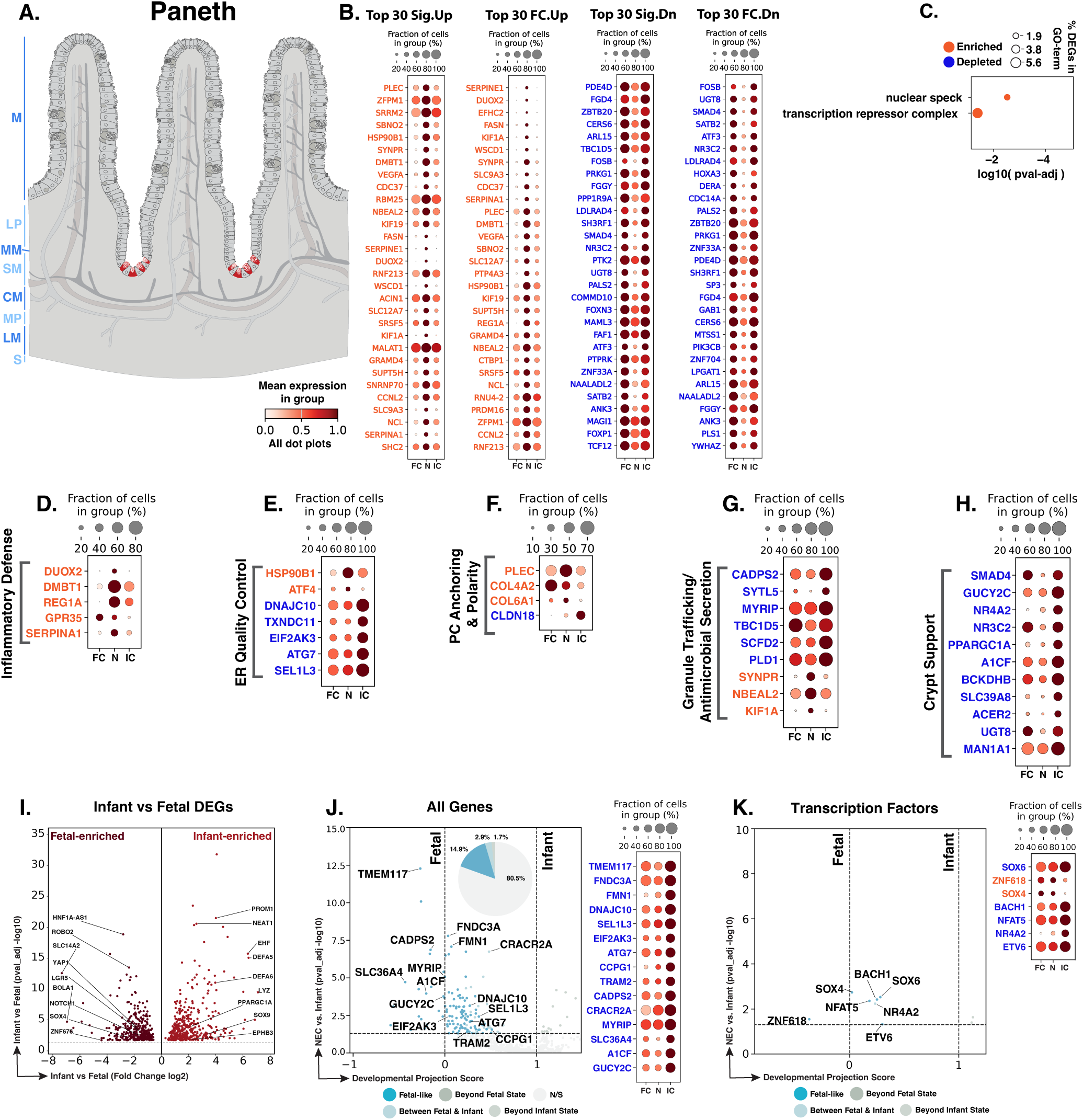
NEC Paneth cells retain crypt-base identity but show secretory stress, granule-routing, and maturity-associated defects. **A.** Schematic showing the crypt-base location of Paneth cells (red). Blue text to the left of the Panel A schematic indicates general anatomical regions: M, mucosa; LP, lamina propria; MM, muscularis mucosae; SM, submucosa; CM, circular muscle; MP, myenteric plexus; LM, longitudinal muscle; S, serosa. **B.** Dot plots of top NEC-specific genes ranked by significance and fold change. **C.** GO-term analysis showing enriched biological processes among NEC-associated DEGs. **D–H.** Curated gene panels showing inflammatory-defense, ER quality-control, Paneth-cell anchoring/polarity, granule trafficking/antimicrobial secretion, and crypt-support programs. In panels **B** and **D–H**, orange gene text indicates upregulation in NEC relative to infant controls, whereas blue gene text indicates downregulation. **I.** Volcano plot of DEGs between fetal and infant controls that define the developmental trajectory for Paneth cells. **J–K.** Gene-level and transcription-factor maturity analyses of fetal-versus-infant DEGs, highlighting the subset also significantly changed in NEC relative to infant controls and positioned relative to fetal and infant reference states, and their corresponding dotplots. The developmental projection score in panels **J–K** is displayed on a symmetric logarithmic (symlog) scale. For expression dot plots, dot size indicates the fraction of nuclei expressing each gene and dot color indicates mean expression. In maturity plots, vertical dotted lines indicate the fetal and infant reference scores. Colored dots classify NEC-versus-infant DEGs as Fetal-like, Between Fetal & Infant, Beyond Fetal State, or Beyond Infant State; gray N/S dots are fetal-versus-infant DEGs without a significant NEC-versus-infant change.

We identified 305 Paneth cell nuclei across the atlas. In NEC Paneth cells, differential-expression analysis identified 295 genes significantly changed relative to infant controls, including 65 upregulated and 230 downregulated genes (Fig. 13B). The top DEGs showed a stress-associated secretory pattern similar to earlier epithelial compartments. Inflammatory-defense, ER quality control, secretory/granule-associated, and crypt-architecture genes were increased, whereas metabolic-regulatory, epithelial-signaling, metal/ion-handling, glycan-processing, and selected maturity-associated genes were reduced (Fig. 13B). GO-term analysis highlighted nuclear speck and transcription-repressor-complex terms (Fig. 13C). Nuclear speckles are nuclear compartments enriched for RNA-processing factors, so this enrichment may reflect altered RNA handling in stressed Paneth cells; together with transcription-repressor-complex enrichment, this could affect how Paneth cells maintain secretory-state genes during NEC.^551^

Paneth cell antimicrobial dysfunction is central to NEC pathobiology. We therefore specifically examined canonical Paneth effectors; *DEFA5*, *DEFA6*, *LYZ*, *PLA2G2A*, and *REG3A* were detected but were not significantly altered in NEC Paneth cells compared to infant controls. We also assessed whether NEC Paneth cells showed evidence of proliferative or ISC-like reversion because mouse injury studies have shown that Paneth cells can acquire stem-like transcriptional programs, re-enter the cell cycle, dedifferentiate after Notch activation, and contribute to epithelial regeneration.^552–554^ However, *MKI67*, *CENPF*, *TOP2A*, *PCNA*, *LGR5*, *OLFM4*, *SOX9*, *ASCL2*, *SMOC2*, *LRIG1*, and *PROM1* did not reach significance, arguing against overt conversion to a proliferative ISC-like state. Together, these findings suggest that NEC Paneth cells retain crypt-base antimicrobial identity while acquiring a stress-associated secretory and regulatory state that could affect antimicrobial delivery, epithelial architecture, and ISC-niche support without evidence of reduced defensin transcript expression or stem-like conversion.

NEC Paneth cells shared inflammatory-defense features seen across absorptive and other secretory lineages (Fig. 13D). Increased *DUOX2* supports epithelial oxidant antimicrobial defense and is consistent with *DUOX2*/*DUOXA2* induction in inflamed intestinal epithelium.^160,469^ Increased *DMBT1* supports mucosal defense, while *REG1A* and *GPR35* suggest epithelial injury-protective and restitution-associated signaling.^162,526,555^ Increased *SERPINA1* suggests antiprotease protection through alpha-1-antitrypsin inhibition of neutrophil elastase.^556^ These data suggest that NEC Paneth cells participate in the broader epithelial inflammatory-defense state; however, this defense program may be less effective if Paneth cell ER quality control or granule handling is impaired.

NEC Paneth cells showed a mixed ER-stress and protein-quality control pattern that overlaps with earlier epithelial lineages but includes Paneth-restricted features relevant to secretory cargo handling (Fig. 13E). Higher *HSP90B1* suggests a shared epithelial ER chaperone response, while higher *ATF4*, Paneth-restricted among epithelial DEGs, points to PERK/eIF2α-linked stress adaptation.^236,557^ In contrast, lower EIF2AK3, DNAJC10, TXNDC11, and ATG7 suggest reduced support for PERK/ISR signaling, ER redox-dependent protein quality control, glycoprotein ER-associated degradation, and autophagy.^557–560^ Lower SEL1L3 also suggests altered ER-stress regulation.^561–562^ Lower *TXNDC11* was also Paneth-restricted among epithelial DEGs, suggesting a Paneth-cell-specific weakness in ER redox and quality control support. Lower *ATG7* is especially relevant because epithelial *Atg7* deletion in mice impairs Paneth cell granule formation and reduces lysozyme, linking autophagy loss to defective antimicrobial granule biology.^560^ Taken together, NEC Paneth cells increase selected stress-response and chaperone genes but express lower levels of ER quality control and autophagy genes needed to handle antimicrobial secretory cargo. This could weaken microbial containment during NEC, consistent with mouse studies showing that Paneth cell disruption plus live bacteria is sufficient to induce NEC-like injury in immature intestine.^549–550^

NEC Paneth cells had altered expression of genes linked to crypt-base anchoring, extracellular matrix, and junctional organization, similar to structural changes seen in absorptive and other secretory lineages (Fig. 13F). Higher *PLEC* suggests increased cytoskeletal anchoring machinery associated with hemidesmosome-like epithelial attachment, while higher *COL4A2* and *COL6A1* point to increased basement membrane and extracellular matrix gene expression at the crypt base.^367,563–566^ In contrast, lower *CLDN18* suggests reduced tight junction-associated barrier support.^567^ Taken together, this pattern suggests that NEC Paneth cells may increase anchoring and matrix-associated gene expression while losing junctional barrier features, potentially weakening crypt-base organization and microbial containment.

Granule-trafficking genes pointed to an imbalanced secretory state (Fig. 13G). Lower *CADPS2*, *SYTL5*, *MYRIP*, *TBC1D5*, and *PLD1* suggest reduced dense-core vesicle exocytosis, Rab27A-linked granule positioning, retromer/Rab7 cargo sorting, and vesicle docking or fusion.^509,568–571^ By contrast, higher NBEAL2 and SYNPR point to granule biogenesis and vesicle-membrane association, while higher KIF1A, a Paneth-cell-specific DEG, suggests altered microtubule-based vesicle transport in NEC Paneth cells.^572–574^ Overall, NEC Paneth cells express lower levels of genes needed to prime, position, sort, and fuse secretory vesicles while retaining selected biogenesis and transport signals. This imbalance could limit timely release of antimicrobial cargo at the crypt base, weakening microbial containment.

A separate group of reduced DEGs implicated Paneth cell maintenance functions rather than inflammatory defense (Fig. 13H). Lower *SMAD4* and *GUCY2C* suggest reduced signaling that normally helps preserve differentiated epithelial identity, restrain WNT-associated dedifferentiation, limit ER stress, and support LGR5+ stem-cell integrity.^575–576^ Lower *PPARGC1A* points to weaker mitochondrial-metabolic support, while lower *SLC39A8* suggests reduced manganese-dependent glycosylation capacity that could affect secretory protein quality.^214,216,450^ Lower *A1CF* suggests reduced APOBEC1-associated RNA-editing support, and lower *ACER2* and *UGT8* point to reduced sphingolipid processing and galactosylceramide production.^577–580^ Together, this pattern suggests reduced capacity to sustain the signaling, metabolic, glycan-processing, RNA-regulatory, and lipid-handling demands of a mature secretory crypt-base cell. Because Paneth cells contribute to intestinal antimicrobial defense, microbial containment, and ISC-niche support, and because Paneth cell disruption can increase susceptibility to NEC-like injury, this loss of maintenance capacity could compromise Paneth cell function during NEC and reduce crypt-base support during recovery.^149–150,549,581^

Of the 770 genes distinguishing fetal from infant control Paneth cells, 150 were also dysregulated in NEC versus infant controls, including 115 with fetal-like expression (76.7% of the 150 genes; Fig. 13I,J). Many of these genes were already part of the functional themes described above, including granule delivery (*CADPS2*, *MYRIP*), ER quality control/autophagy (*DNAJC10*, *SEL1L3*, *EIF2AK3*, *ATG7*), and crypt-base maintenance, metabolism, glycosylation, and RNA regulation (*GUCY2C*, *PPARGC1A*, *SLC39A8*, *A1CF*).^214,450,557,560,568,570,576–577^ Thus, the maturity analysis suggests that NEC Paneth cells express key secretory, metabolic, glycan-processing, and quality control genes closer to fetal levels. This may reflect persistent developmental immaturity or injury-associated reversion, either of which could weaken antimicrobial reserve, microbial containment, and ISC-niche support during NEC.^150,548^

TF maturity analysis added a regulatory layer to the developmental interpretation. NEC Paneth cells showed a fetal-like TF pattern defined by higher *SOX4* and lower *SOX6*, *BACH1*, and *NFAT5* (Fig. 13K). Higher *SOX4*, a Paneth-cell-restricted epithelial DEG, may indicate a fetal-like or intermediate secretory-regulatory state rather than terminal Paneth cell maturation. This is consistent with mouse evidence that *Sox4* marks secretory/progenitor-associated states and promotes *Atoh1*-independent secretory differentiation toward tuft and enteroendocrine fates, whereas mature Paneth cell differentiation is more directly linked to *Sox9*.^582–584^ Lower SOX6 was also fetal-like, supporting a broader SOX-family regulatory pattern, although its specific role in Paneth cells remains less defined. Lower *BACH1* may reduce repression of NRF2/*HMOX1*-linked antioxidant transcription, while lower *NFAT5* suggests reduced osmotic-stress and barrier-support signaling.^545,585^

NEC Paneth cells retain crypt-base antimicrobial identity but adopt an injury-associated secretory and regulatory state. The absence of significant defensin downregulation and lack of proliferative or ISC-marker induction argue against simple defensin loss or overt stem-like conversion. Initiation may instead involve reduced antimicrobial secretory reserve caused by altered ER quality control, autophagy, granule routing, metabolic/glycan support, and crypt-base homeostasis. During NEC progression, these changes could weaken microbial containment and Paneth cell function under inflammatory and ER stress. Resolution may fail if the crypt base cannot restore a mature Paneth cell niche capable of supporting epithelial repair and stable ISC function.

### Stromal compartment

The intestinal stroma, marked by *PDGFRA* and *PTCH1* (Fig. 2), provides structural support and signaling cues to epithelial, vascular, and immune compartments and contributes to epithelial niche maintenance, extracellular matrix organization, and tissue repair.^29,48,586–587^ These stromal functions are highly relevant to NEC, where inflammatory activation, extracellular matrix remodeling, altered niche support, vascular dysfunction, and impaired tissue repair may influence disease progression and recovery.^18,156,588–589^ Prior human NEC single-cell transcriptomic analysis showed increased fibroblast representation and pro-inflammatory gene expression, and experimental neonatal mouse NEC multi-omics identified an expanded fibroblast compartment; however, these studies did not resolve how distinct fibroblast and non-fibroblast stromal populations respond to NEC.^18,157^

To define the stromal populations in our study, heatmap-based annotation resolved nine stromal populations: *ADAMDEC1*+ lamina propria fibroblasts, *RSPO3+/GREM1*+ trophocytes, *PDGFRA*+*/KCNN3*+ SIP fibroblasts, *MYH11*+/*GREM2*+ lamina propria myocytes, *PDGFRA*+/*NRG1*+/*F3*+ subepithelial myofibroblasts, *MKI67*+ proliferative stroma, *PDGFRB*+ pericytes, *ANO1*+ interstitial cells of Cajal, and *WT1*+/*MSLN*+ mesothelial cells (Fig. 14A,B). ICCs and mesothelial cells had insufficient NEC-versus-infant differential gene-expression support for detailed interpretation.

**Fig. 14.**
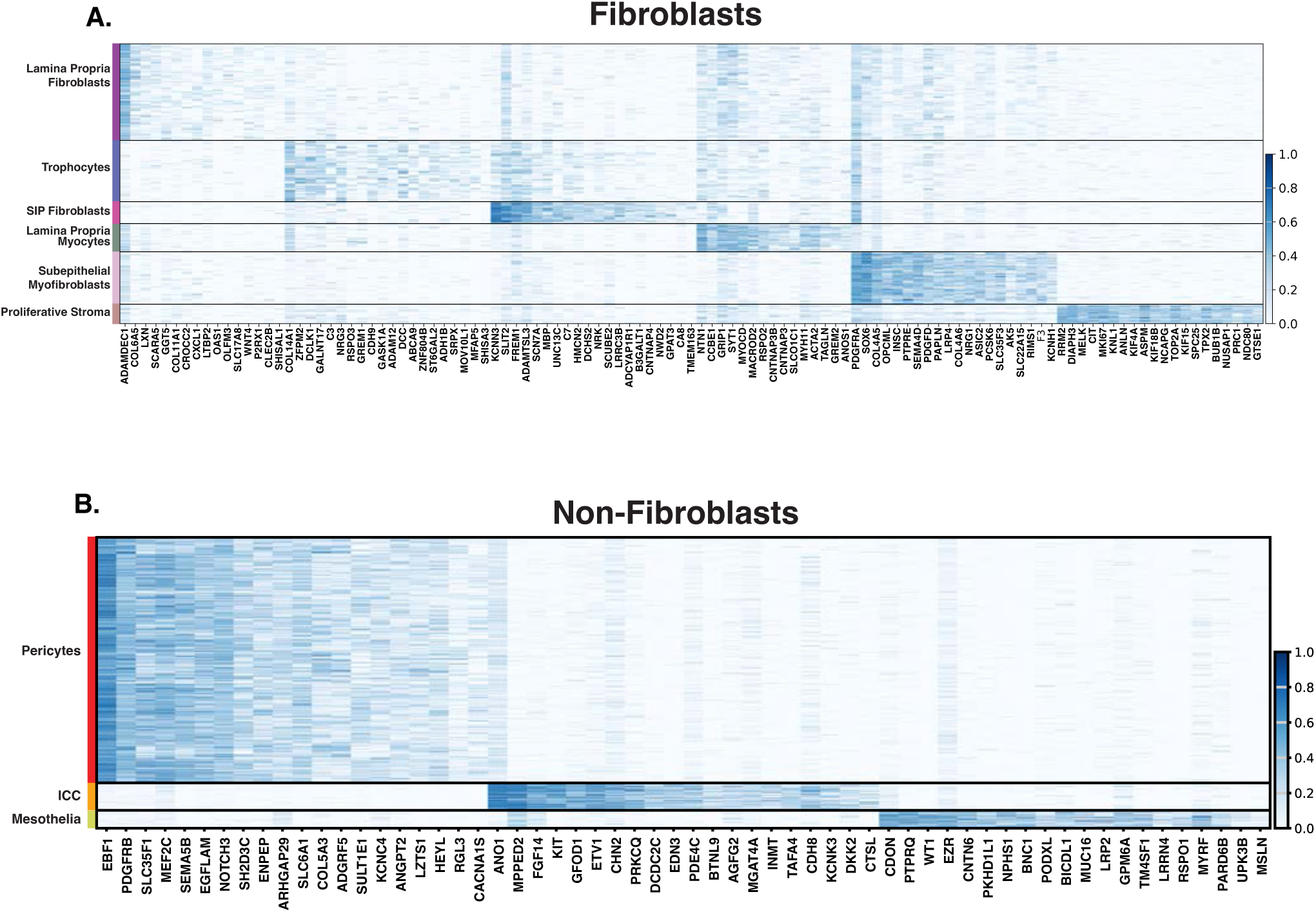
Stromal cell-type signatures define fibroblast and non-fibroblast populations. **A.** Heatmap of marker-gene expression across stromal fibroblast populations, including lamina propria fibroblasts, trophocytes, SIP fibroblasts, lamina propria myocytes, subepithelial myofibroblasts, and proliferative stroma. **B.** Heatmap of marker-gene expression across non-fibroblast stromal populations, including pericytes, interstitial cells of Cajal, and mesothelial cells. Distinct transcriptional signatures support stromal cell-type assignments and provide the framework for the detailed stromal analyses that follow.

The following sections examine how NEC alters stromal populations in different regions of the intestinal wall. Lamina propria fibroblasts and trophocytes capture the mucosal and crypt-associated response, where inflammation, hypoxia/angiogenesis, epithelial-repair signaling, matrix remodeling, and niche-support changes overlap. SIP fibroblasts and lamina propria myocytes extend this view into stromal populations with neuromuscular and contractility-associated features. Subepithelial myofibroblasts and proliferative stroma capture subepithelial support and cycling repair states that appear activated but not fully organized, while pericytes define a vascular-support response involving inflammatory chemokine signaling, vascular tone, endothelial barrier regulation, stress/senescence features, and hypoxia-associated vascular repair. Together, these sections frame NEC stroma as a set of specialized populations that may help coordinate repair, but may also worsen disease when inflammation, matrix remodeling, impaired niche signaling, and incomplete developmental maturation remain unresolved.

### Lamina Propria Fibroblasts

Lamina propria fibroblasts are mucosal fibroblasts positioned beneath the intestinal epithelium within the villus core, where they sense epithelial injury, microbial and immune-derived signals, and coordinate tissue repair and epithelial homeostasis (Fig. 15A).^590–592^ Prior mouse and human intestinal studies define lamina propria fibroblasts as stromal cells marked by low *PDGFRA* and the metalloproteinase *ADAMDEC1*.^592–595^ The prior human and mouse NEC transcriptomic data showing an increase in fibroblasts in NEC together suggest that fibroblasts are important in NEC but pathways and gene changes within specific fibroblast populations like the lamina propria fibroblasts were not defined.^18,157^ Thus, the disease-associated programs adopted by this mucosal fibroblast population near the injured epithelium remain poorly defined in human NEC.

**Fig. 15.**
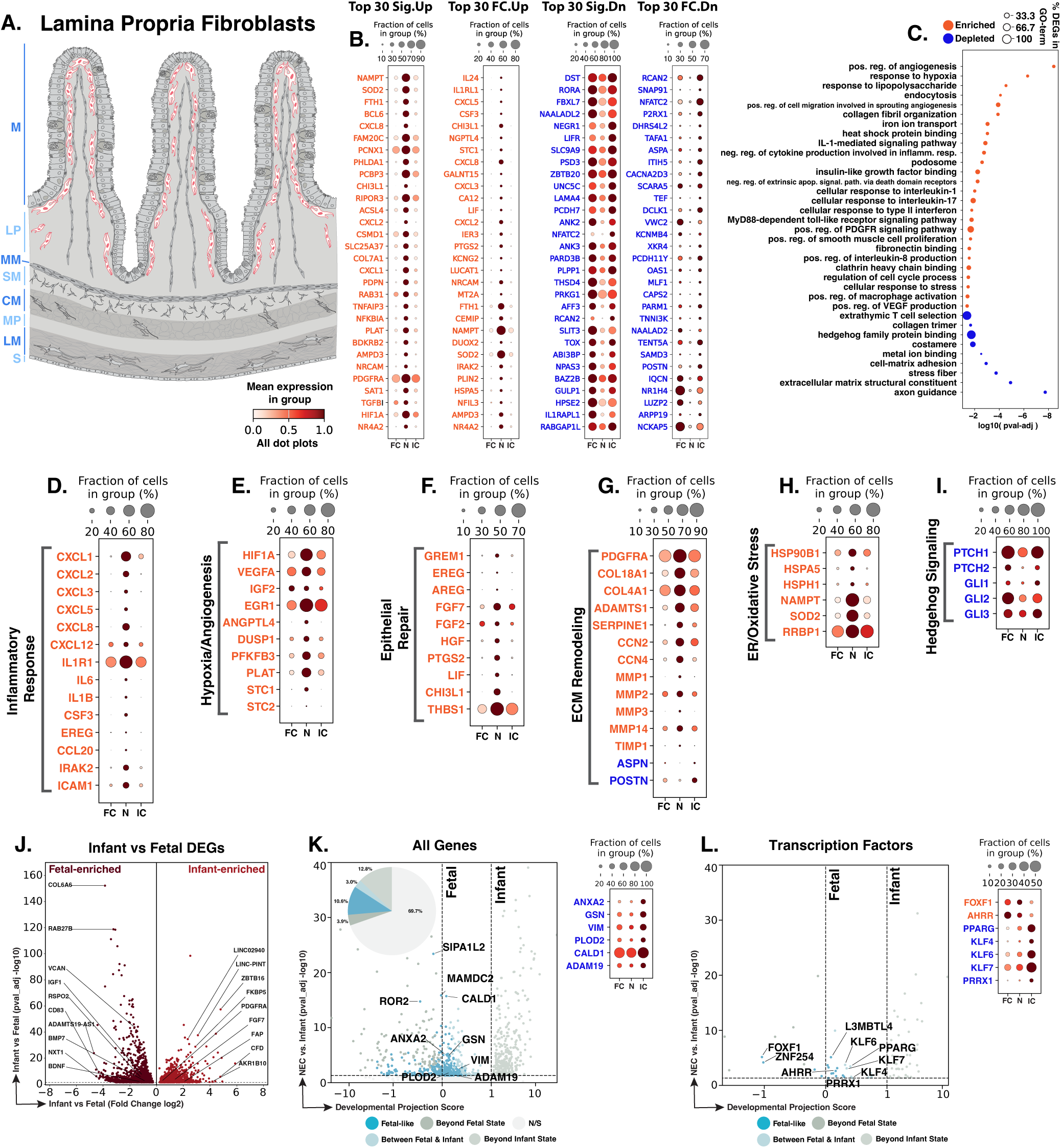
NEC lamina propria fibroblasts show inflammatory, epithelial-repair, extracellular-matrix, Hedgehog-signaling-associated changes. **A.** Schematic showing the mucosal distribution of lamina propria fibroblasts (red). Blue text to the left of the Panel A schematic indicates general anatomical regions: M, mucosa; LP, lamina propria; MM, muscularis mucosae; SM, submucosa; CM, circular muscle; MP, myenteric plexus; LM, longitudinal muscle; S, serosa. **B.** Dot plots of top NEC-specific genes ranked by significance and fold change. **C.** GO-term analysis showing enriched and depleted biological processes among NEC-associated DEGs. **D–I.** Curated gene panels showing inflammatory response, hypoxia/angiogenesis, epithelial repair, extracellular matrix, ER/oxidative stress, and Hedgehog-signaling programs. In panels **B** and **D–I**, orange gene text indicates upregulation in NEC relative to infant controls, whereas blue gene text indicates downregulation. **J.** Volcano plot of DEGs between fetal and infant controls that define the developmental trajectory for lamina propria fibroblasts. **K–L.** Gene-level and transcription-factor maturity analyses of fetal-versus-infant DEGs, highlighting the subset also significantly changed in NEC relative to infant controls and positioned relative to fetal and infant reference states, and their corresponding dotplots. The developmental projection score in panels **K–L** is displayed on a symmetric logarithmic (symlog) scale. For expression dot plots, dot size indicates the fraction of nuclei expressing each gene and dot color indicates mean expression. In maturity plots, vertical dotted lines indicate the fetal and infant reference scores. Colored dots classify NEC-versus-infant DEGs as Fetal-like, Between Fetal & Infant, Beyond Fetal State, or Beyond Infant State; gray N/S dots are fetal-versus-infant DEGs without a significant NEC-versus-infant change.

We identified 3,703 lamina propria fibroblast nuclei across the atlas, with 1,692 genes significantly changed in NEC lamina propria fibroblasts compared to infant controls, including 706 upregulated and 986 downregulated genes (Fig. 15B). GO-term enrichment analysis between NEC and infant control showed that lamina propria fibroblasts in NEC were enriched for angiogenic and hypoxic, innate inflammatory and immune-recruitment, endocytic, and collagen-organization programs, and depleted for adhesion, axon-guidance, and Hedgehog-associated programs (Fig. 15C). These GO-term results were reflected in the inflammatory, hypoxia, epithelial-repair, and Hedgehog signaling curated gene-level analysis below, along with extracellular matrix remodeling and oxidative stress gene changes.

NEC lamina propria fibroblasts showed a prominent inflammatory-response program that could establish innate immune recruitment cues beneath the injured epithelium. *CXCL1*, *CXCL2*, *CXCL5*, and *CXCL8*, CXC chemokines that can be produced by fibroblasts to recruit neutrophils, were significantly upregulated in NEC lamina propria fibroblasts compared to infant control (Fig. 15D).^596–597^ *IL1R1*, the receptor for the pro-inflammatory cytokine IL-1, *IL6*, a fibroblast-derived inflammatory cytokine, and *CCL20*, a chemokine induced by inflammatory cytokine signaling and involved in immune cell recruitment, were also significantly upregulated.^597–599^ This response could support microbial containment after epithelial barrier disruption, but persistent chemokine and cytokine production could contribute to disease progression by sustaining neutrophil-rich inflammation and amplifying mucosal injury.^597,600–602^

Lamina propria fibroblasts in NEC also exhibited significant upregulation of genes associated with hypoxia and angiogenesis. *HIF1A*, the canonical transcription factor induced during hypoxia, and *VEGFA*, an angiogenic factor regulated by HIF1A, were upregulated, supporting a hypoxia-associated vascular response (Fig. 15E).^62,603–604^ *EGR1* is an early-response transcription factor that acts upstream of HIF1A during hypoxia, and *STC1* can be induced by HIF1A under hypoxic conditions, and both are upregulated in NEC lamina propria fibroblasts.^605–607^ *DUSP1*, which can be induced by hypoxia, and the HIF-responsive glycolytic gene *PFKFB3* were also upregulated in NEC, further supporting a cellular response to hypoxia.^291–292,608^ This response could aid in the resolution of NEC if it supports intestinal microvascular integrity, perfusion, and restoration of tissue oxygenation; however, if hypoxia and vascular stress persist, sustained hypoxia-associated angiogenic signaling could instead contribute to ongoing inflammation and disease progression.^609–614^

NEC lamina propria fibroblasts also upregulated epithelial-repair signals, suggesting that inflammatory activation is paired with attempted mucosal regeneration. *GREM1*, a BMP antagonist produced by intestinal mesenchymal populations near the crypt supports ISC homeostasis and epithelial repair.^29,593^ *EREG* and *AREG* are EGF-family ligands implicated in intestinal epithelial regeneration and repair signaling by acting as important niche cues (Fig. 15F).^615–619^ *FGF7*, also known as keratinocyte growth factor, is a fibroblast-derived epithelial mitogen that promotes intestinal epithelial proliferation and repair, while *FGF2* is expressed in mesenchyme surrounding regenerating crypts and supports epithelial stem-cell survival after injury.^620–621^ *LIF* is a fibroblast-produced cytokine that supports intestinal stem-cell homeostasis and epithelial regeneration.^622–623^ Significant upregulation of these genes suggests that lamina propria fibroblasts may attempt to support epithelial repair during NEC, although coupling of this program to inflammatory and hypoxic signals suggests a stressed repair program that may not reach fully coordinated mucosal resolution.

NEC lamina propria fibroblasts showed an altered extracellular matrix program, indicating active but potentially imbalanced matrix turnover. *MMP1*, *MMP2*, *MMP3*, and *MMP14*, matrix metalloproteinases involved in extracellular matrix degradation, pericellular remodeling, and wound-associated tissue remodeling, were significantly upregulated in NEC lamina propria fibroblasts (Fig. 15G).^624–625^ *TIMP1*, an inhibitor of matrix metalloproteinases, was also significantly upregulated, suggesting that matrix degradation and matrix restraint are activated in parallel.^626^ By contrast, *ASPN*, an extracellular matrix protein that can inhibit TGF-beta signaling, was significantly downregulated, suggesting reduced restraint of TGF-beta-linked remodeling.^627–628^ This pattern suggests mixed extracellular matrix turnover, which could support the resolution of NEC if it enables epithelial restitution and stromal reorganization, but could contribute to disease progression if proteolysis, inhibitory feedback, and reduced matrix-regulatory programs destabilize the mucosal matrix.

NEC lamina propria fibroblasts also upregulated genes associated with ER quality control stress and oxidative stress. *HSPA5*, which encodes GRP78/BiP and is a canonical ER stress marker, and *HSP90B1*, which encodes the ER chaperone Grp94 and is induced by ER stress, were upregulated in NEC, supporting increased protein-folding and ER quality control activity (Fig. 15H).^629–631^ *SOD2*, a mitochondrial antioxidant enzyme that can be induced during oxidative stress, was also upregulated, suggesting adaptation to oxidative stress.^632–633^ The upregulation of these genes suggest that NEC lamina propria fibroblasts may activate programs to preserve function during inflammatory and hypoxic stress. Persistent stress responses, however, could mark unresolved injury and contribute to impaired resolution of NEC.

The downregulation of Hedgehog-associated genes in NEC lamina propria fibroblasts suggests disruption of epithelial-mesenchymal crosstalk that normally supports intestinal homeostasis.^634–636^ *PTCH1* and *PTCH2*, canonical Hedgehog receptors, as well as downstream Hedgehog-associated transcription factors *GLI1*, *GLI2*, and *GLI3*, were significantly downregulated in NEC (Fig. 15I).^635,637–638^ Because epithelial-derived Hedgehog signaling normally restrains stromal inflammatory signaling and immune cell recruitment in the intestine, the downregulation of Hedgehog-associated genes in NEC lamina propria fibroblasts is consistent with diminished epithelial regulation of stromal fibroblasts and could contribute to inflammation and enhanced immune cell recruitment within the lamina propria during NEC.^639–640^ This mechanism could further disease progression if the heightened inflammatory response in the lamina propria is left unchecked.

Of the 2,817 genes distinguishing fetal from infant control lamina propria fibroblasts, 854 were also dysregulated in NEC versus infant controls, including 298 with fetal-like expression (34.9% of the 854 genes; Fig. 15K). The developmental contrast included fetal-high *BMP7* and *RSPO2*, consistent with mesenchymal patterning and epithelial-support signals, and infant-high *FGF7*, a fibroblast-derived epithelial growth factor (Fig. 15J).^47,620,641–642^ The fetal-like NEC genes highlighted reduced fibroblast activation, migration, cytoskeletal, and matrix-associated features. Lower *ANXA2*, *GSN*, and *VIM* point to weaker fibroblast migration, actin regulation, activation, proliferation, and wound-healing capacity.^643–646^ Lower *PLOD2* and *CALD1* further suggest reduced collagen crosslinking and actin-associated contractile regulation.^647–649^ Together, these findings suggest that a subset of NEC lamina propria fibroblast genes remains closer to a fetal state, which may reflect persistent immaturity or injury-associated reversion and could limit fibroblast activation, matrix support, and mucosal repair.

At the transcription factor level, NEC lamina propria fibroblasts showed fetal-like regulatory changes linked to migration, fibroblast activation, and extracellular matrix control (Fig. 15L). *FOXF1* and *AHRR* were increased in NEC and remained fetal-like, whereas *PPARG*, *KLF4*, *KLF6*, and *KLF7* were decreased in NEC and closer to the fetal reference. FOXF1/Foxf1 has been linked to extracellular matrix production and fibroblast migration, and AHRR represses AHR signaling, which can reduce collagen synthesis in the gut.^650–655^ Lower *PPARG*, *KLF4*, *KLF6*, and *KLF7* further suggest reduced regulatory input into collagen production, fibroblast activation, proliferation, myofibroblast transition, and inflammatory activity.^656–660^ These TF patterns suggest that NEC lamina propria fibroblasts retain fetal-like regulatory features that could alter matrix remodeling and limit coordinated mucosal repair.

NEC lamina propria fibroblasts occupy a strongly inflammatory, hypoxia- and stress-responsive, epithelial-repair-associated, and matrix-remodeling state beneath the epithelium. Initiation may be influenced by their position at the mucosal interface, where chemokine-mediated immune recruitment, cytokine production, and hypoxic stress responses could amplify early epithelial injury. During progression, persistent inflammatory and matrix-remodeling signals may sustain mucosal damage and disrupt stromal organization. At the same time, increased epithelial-support genes suggest an attempted repair response. Resolution may depend on whether growth factor support and extracellular matrix remodeling become coordinated, or whether stromal repair remains coupled to inflammation and hypoxic stress.

### Trophocytes

Trophocytes are crypt-associated fibroblasts located in the submucosa, where they form an important component of the intestinal stem-cell niche (Fig. 16A).^29,661^ Trophocytes and related crypt-bottom fibroblasts provide WNT- and BMP-regulatory niche signals, including WNT2B, R-spondins, and GREM1, that support intestinal stem-cell maintenance and proliferation, and trophocytes can sustain intestinal organoid growth without additional stromal trophic support.^29,662^ *Rspo3*+/*Grem1*+ crypt-base fibroblasts also act together with neighboring lymphatic endothelial cells to support intestinal stem cells during homeostasis and injury, where following injury, both populations expand and increase *Rspo3* and other growth factors that foster intestinal repair.^594^ Prior human NEC transcriptomic data and experimental neonatal mouse NEC multi-omics showed an increased proportion of fibroblasts in NEC, but did not define trophocyte-specific disease programs.^18,157^ Thus, it remains unknown whether human NEC trophocytes acquire disease-related programs that could shape bowel wall repair.

**Fig. 16.**
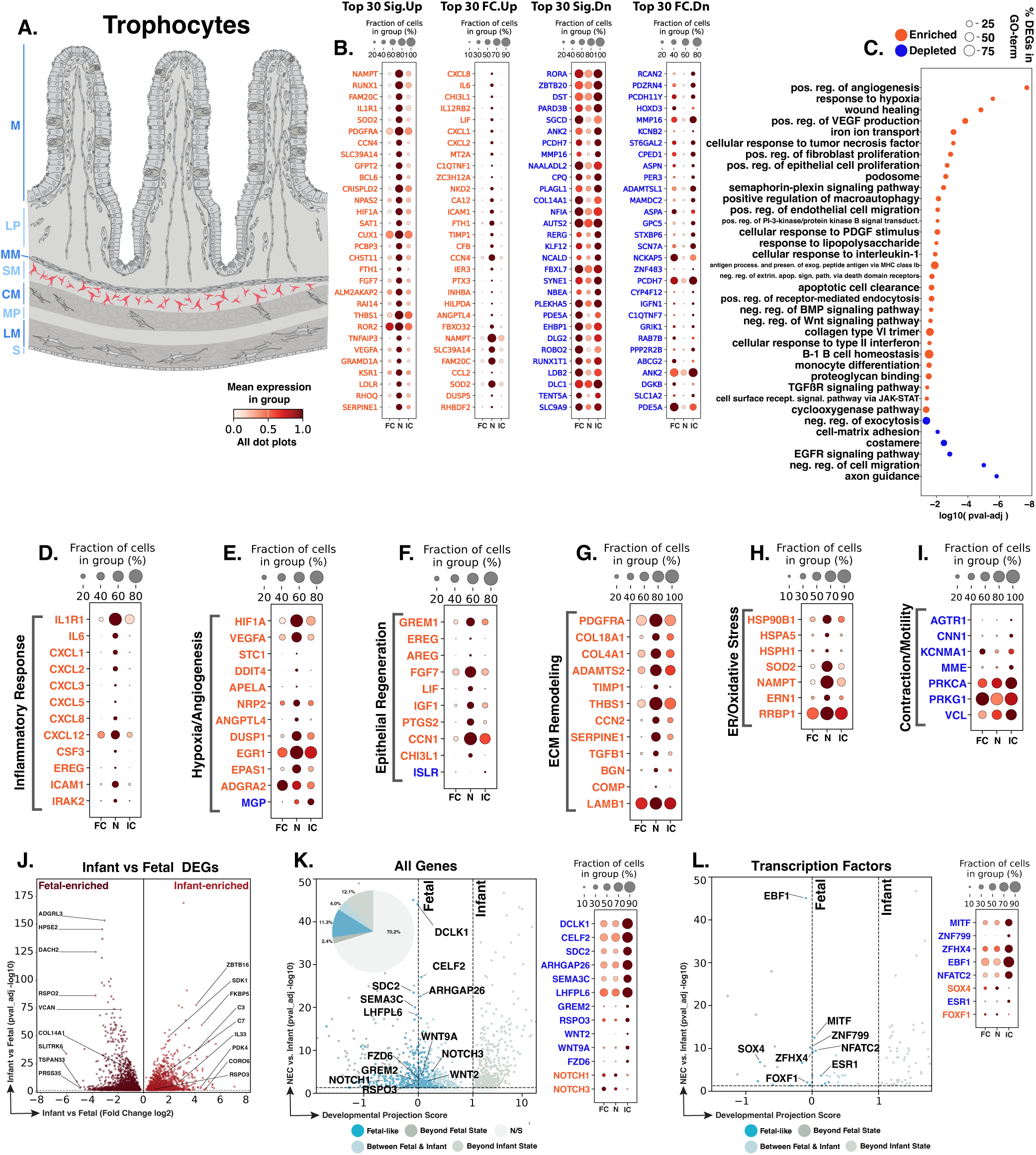
NEC trophocytes show inflammatory, hypoxia/angiogenesis, epithelial-regeneration, and extracellular-matrix-associated changes. **A.** Schematic showing the crypt-associated stromal location of trophocytes (red). Blue text to the left of the Panel A schematic indicates general anatomical regions: M, mucosa; LP, lamina propria; MM, muscularis mucosae; SM, submucosa; CM, circular muscle; MP, myenteric plexus; LM, longitudinal muscle; S, serosa. **B.** Dot plots of top NEC-specific genes ranked by significance and fold change. **C.** GO-term analysis showing enriched and depleted biological processes among NEC-associated DEGs. **D–I**. Curated gene panels showing inflammatory response/neutrophil recruitment, hypoxia/angiogenesis, epithelial regeneration, extracellular matrix remodeling, ER/oxidative stress, and contractile programs. In panels **B** and **D–I**, orange gene text indicates upregulation in NEC relative to infant controls, whereas blue gene text indicates downregulation. **J.** Volcano plot of DEGs between fetal and infant controls that define the developmental trajectory for trophocytes. **K–L.** Gene-level and transcription-factor maturity analyses of fetal-versus-infant DEGs, highlighting the subset also significantly changed in NEC relative to infant controls and positioned relative to fetal and infant reference states, and their corresponding dotplots. The developmental projection score in panels **K–L** is displayed on a symmetric logarithmic (symlog) scale. For expression dot plots, dot size indicates the fraction of nuclei expressing each gene and dot color indicates mean expression. In maturity plots, vertical dotted lines indicate the fetal and infant reference scores. Colored dots classify NEC-versus-infant DEGs as Fetal-like, Between Fetal & Infant, Beyond Fetal State, or Beyond Infant State; gray N/S dots are fetal-versus-infant DEGs without a significant NEC-versus-infant change.

We identified 2,352 trophocyte nuclei across the atlas, with 2,057 genes significantly changed in NEC trophocytes compared to infant controls, including 977 upregulated and 1,080 downregulated genes (Fig. 16B). GO-term enrichment highlighted increased hypoxia/angiogenic, wound-healing/proliferative, inflammatory/cytokine, TGF-β signaling, and iron-transport features in NEC, together with reduced cell-matrix/costamere, EGFR-signaling, migration-regulatory, and axon-guidance GO-terms (Fig. 16C). These GO-term changes were consistent with the inflammatory, hypoxia-angiogenesis, and matrix/wound-healing features described below.

Inflammatory changes in NEC trophocytes could extend immune recruitment into the submucosa around the crypts. *IL1R1*, the receptor for IL-1, was upregulated, consistent with increased responsiveness to IL-1-associated inflammatory fibroblast activation and neutrophil-recruiting stromal programs.^597,663^ IL-1R1 signaling in *Grem1*+ intestinal mesenchymal cells can also increase *Rspo3* and support epithelial regeneration after injury (Fig. 16D).^664^ *IL6*, an inflammatory cytokine produced by activated intestinal fibroblasts, was also upregulated.^663,665^ *CXCL1*, *CXCL2*, *CXCL3*, *CXCL5*, and *CXCL8*, neutrophil-targeting chemokines expressed by inflammatory intestinal fibroblasts, and *CXCL12*, a fibroblast-derived chemokine associated with intestinal immune cell recruitment, were upregulated in NEC trophocytes.^597,640^ This inflammatory response could promote immune cell recruitment to crypts after barrier disruption, but persistent IL-1/IL-6 and chemokine signaling could contribute to NEC progression by increasing inflammatory pressure around the intestinal stem cell niche.

A distinct hypoxia- and angiogenesis-associated response was evident in NEC trophocytes. *HIF1A*, the canonical hypoxia-responsive transcription factor, and *VEGFA*, a HIF-regulated angiogenic factor, were upregulated, supporting a hypoxia-associated vascular response (Fig. 16E).^62,603–604^ *STC1* can be induced by HIF1A under hypoxic conditions, and *APELA*/Elabela promotes post-ischemic angiogenesis and perfusion recovery; both genes were upregulated in NEC trophocytes.^606–607,666^ *NRP2*, a neuropilin receptor involved in angiogenic endothelial cell adhesion and migration, was also upregulated.^667^ By contrast, *MGP*, matrix Gla protein and a suppressor of angiogenic sprouting, was downregulated, suggesting reduced restraint on angiogenic signaling.^668^ This pattern could support NEC resolution if it improves oxygen delivery and vascular repair in the injured bowel wall; however, persistent hypoxia-associated angiogenic signaling could also mark ongoing vascular stress and contribute to disease progression.

Signals that could support epithelial restitution from the crypt-associated submucosa were also prominent. *GREM1*, a mesenchymal-derived BMP antagonist enriched near the crypt that supports ISC homeostasis and epithelial recovery, was upregulated (Fig. 16F).^29,593,669^ *EREG* and *AREG*, EGF-family ligands that serve as important niche cues for intestinal epithelial regeneration, were also upregulated in NEC trophocytes.^615–619^ *FGF7*, also known as keratinocyte growth factor, is a fibroblast-derived epithelial mitogen and was upregulated, suggesting potential support for epithelial proliferation and recovery.^620,670–671^ *LIF*, leukemia inhibitory factor, supports intestinal stem-cell homeostasis and epithelial regeneration.^623^ *IGF1*, an insulin-like growth factor linked to crypt regeneration, was also upregulated.^672^ Together, these genes suggest that NEC trophocytes may provide epithelial growth and regenerative cues during injury that could aid in NEC resolution. However, *ISLR*, a stromal-derived factor known to support epithelial restitution, was downregulated in NEC, pointing to a more complex epithelial-support response.^673^

Extracellular-matrix and wound-healing features formed another major component of the trophocyte response and may help stabilize injured tissue but become maladaptive if sustained. Increased *PDGFRA* and *TIMP1* pointed to enhanced fibroblast activity and reduced matrix degradation (Fig. 16G). *PDGFRA* supports fibroblast proliferation during wound healing, whereas TIMP1 inhibits matrix metalloproteinases and can limit collagen degradation in intestinal myofibroblasts.^674–676^ Increased *THBS1*, *CCN2*, *SERPINE1*, *TGFB1*, *BGN*, and *COMP* further supported a matrix-producing, profibrotic state. These genes have been linked to mucosal wound healing, fibroblast activation, collagen production or preservation, impaired fibrinolysis, and fibrostenotic intestinal tissue.^677–686^ These changes may support tissue stabilization and repair during NEC, but if sustained or poorly coordinated, they could favor excessive matrix accumulation, tissue stiffening, and altered epithelial-stromal signaling.

Cellular stress responses provided an additional feature of the NEC trophocyte state. *HSP90B1*, which is induced by ER stress and encodes the ER chaperone Grp94, and *HSPA5*, which is a canonical ER stress marker and encodes GRP78/BiP, were both upregulated in NEC, consistent with increased ER protein-folding and quality control activity (Fig. 16H).^629–631^ *SOD2*, which can be induced during oxidative stress, and *ERN1*, an ER stress sensor, were also upregulated.^632–633,687^ These changes suggest cellular responses that may help trophocytes maintain function during inflammatory, hypoxic, ER, and oxidative stress. If sustained, however, the same stress-response genes could reflect unresolved cellular stress in NEC.

Genes with known roles in smooth muscle contractile regulation were also downregulated in NEC trophocytes. *AGTR1* encodes the angiotensin II receptor, through which angiotensin II can promote intestinal smooth muscle contraction.^688^ *CNN1* is a contractile regulatory protein that modulates smooth muscle contractility.^689–691^ By contrast, *KCNMA1*-encoded BK channels and *PRKG1*-dependent cGMP signaling participate in pathways that promote smooth muscle relaxation.^692–695^ *MME*/neprilysin degrades substance P, a neuropeptide that promotes intestinal contraction.^696–697^ Thus, decreased expression of genes that regulate contractility suggests disruption of contractile signaling in NEC trophocytes. Experimental NEC shows reduced intestinal motility early in disease, and restoration of motility can reduce disease severity, suggesting that these changes could contribute to NEC initiation or progression through altered intestinal motility.^24^

Of the 3,532 genes distinguishing fetal from infant control trophocytes, 1,052 were also dysregulated in NEC versus infant controls, including 398 with fetal-like expression (37.8% of the 1,052 genes; Fig. 16J,K). Developmentally regulated genes included fetal-high *RSPO2* and infant-high *RSPO3*, stromal R-spondin factors that support intestinal epithelial stem-cell growth and crypt regeneration; *RSPO2* is the dominant R-spondin homolog in human fetal trophocytes.^28,593,642,698^ The fetal-like genes in NEC highlighted changes in ISC-niche signaling (Fig. 16K). *GREM2*, a BMP antagonist expressed by intestinal cryptal fibroblasts and smooth muscle cells, *RSPO3*, a stromal Wnt-signaling potentiator that supports intestinal stem cells and epithelial regeneration, and *WNT2*, a Wnt ligand expressed by crypt-associated fibroblasts that can promote intestinal epithelial proliferation, were all significantly downregulated in NEC and showed fetal-like expression.^28,593,662,698–700^ These changes suggest that NEC trophocytes may provide weaker epithelial-support and ISC-niche signals, potentially limiting epithelial renewal and repair during NEC initiation or progression.

At the transcription factor level, NEC trophocytes showed fetal-like expression of genes associated with fibroblast activation and proliferation. *SOX4*, a transcription factor linked to fibroblast activation in kidney fibrosis models, was upregulated in NEC and aligned with the fetal reference state (Fig. 16L).^701^ In contrast, *EBF1*, which marks ECM-producing and proliferative fibroblast populations, was downregulated and also aligned with the fetal reference state.^702^ *NFATC2*, a calcineurin-responsive transcription factor required for hypoxia-induced fibroblast proliferation, and *ESR1*, a nuclear receptor linked to fibroblast proliferation, were likewise downregulated and aligned with the fetal reference state.^703–704^ This pattern suggests that NEC trophocytes show changes resembling the fetal state in genes linked to fibroblast activation and proliferation, consistent with altered stromal regulation during NEC.

NEC trophocytes acquire inflammatory, hypoxia-angiogenic, epithelial-repair, extracellular matrix, and stress-response features within the crypt-associated stromal niche. Initiation may involve inflammatory and chemokine signals around the ISC niche after barrier disruption. Progression may follow if sustained IL-1/IL-6, hypoxia, and matrix-remodeling signals increase inflammatory pressure near crypts while reducing mature niche coordination. Although epithelial-growth and regeneration cues suggest an attempted repair response, reduced niche-associated and fetal-like stromal-support genes may weaken the ability of trophocytes to sustain a mature ISC-supportive environment. Resolution may therefore depend on whether trophocyte repair signals can be uncoupled from persistent inflammation and restored to a coordinated crypt-niche state.

### SIP Fibroblasts

SIP fibroblasts are the PDGFRA-positive stromal cells positioned within the deeper bowel wall, part of the SIP syncytium, a neuromuscular network composed of Smooth muscle cells, Interstitial cells of Cajal, and these PDGFRA+ cells that coordinates enteric neuromuscular signaling, calcium-dependent signaling, and gastrointestinal motility (Fig. 17A).^705–707^ PDGFRA+ cells in the intestinal muscularis participate in purinergic inhibitory neurotransmission and regulate smooth muscle excitability, while interstitial cells of Cajal provide pacemaker activity and smooth muscle cells provide the contractile component of this network.^707–712^ Prior human NEC single-cell RNA sequencing data and experimental neonatal mouse NEC multi-omics showed fibroblast enrichment in NEC, but neither defined a SIP fibroblast-specific injury program.^18,157^ Thus, it is unknown how human NEC might perturb this PDGFRA+ neuromuscular stromal population.

**Fig. 17.**
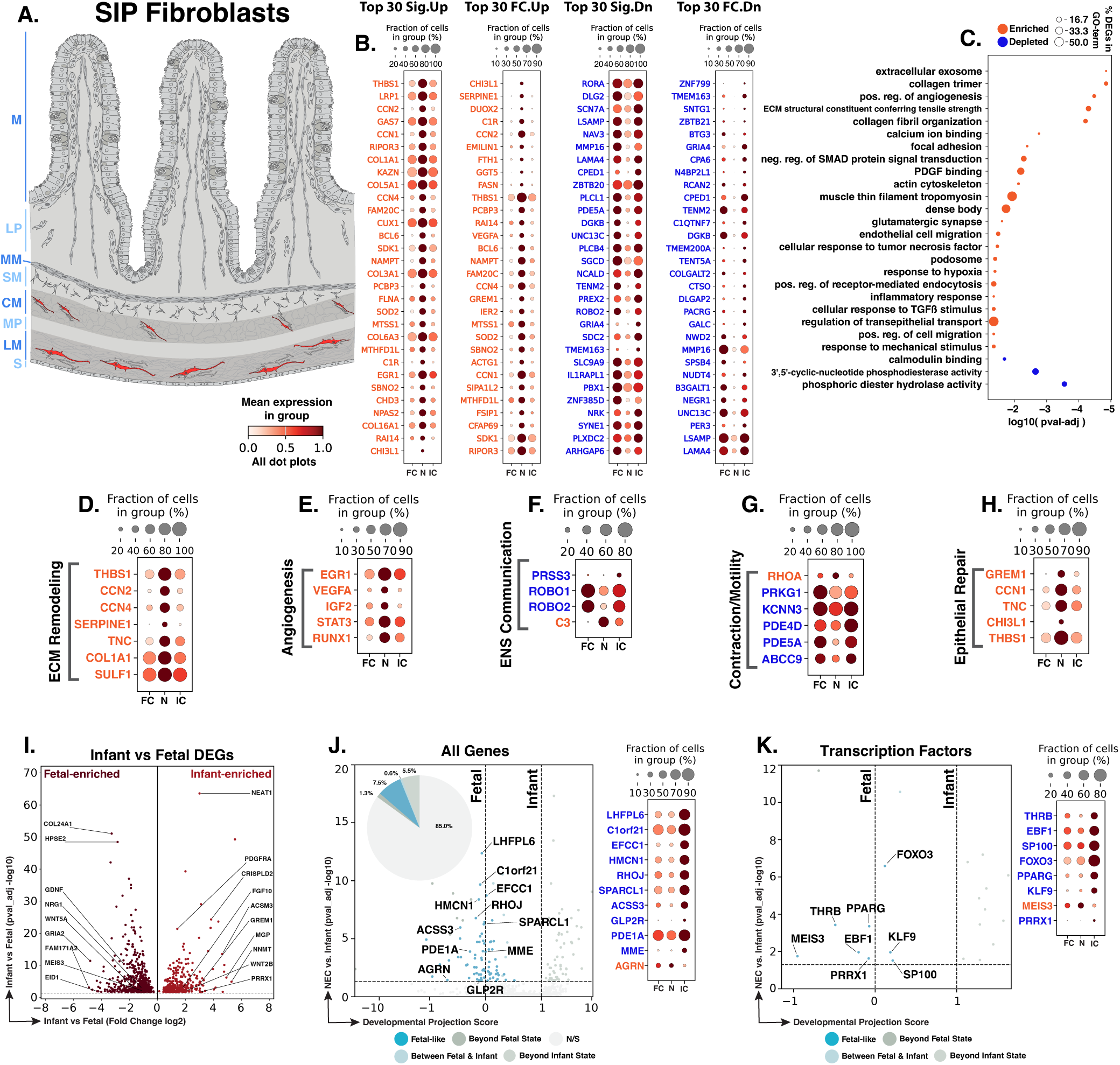
NEC SIP fibroblasts show wound-remodeling, angiogenic, enteric-neuromuscular, and epithelial-repair-associated changes. **A.** Schematic showing the deeper bowel-wall location of SIP fibroblasts (red). Blue text to the left of the Panel A schematic indicates general anatomical regions: M, mucosa; LP, lamina propria; MM, muscularis mucosae; SM, submucosa; CM, circular muscle; MP, myenteric plexus; LM, longitudinal muscle; S, serosa. **B.** Dot plots of top NEC-specific genes ranked by significance and fold change. **C.** GO-term analysis showing enriched and depleted biological processes among NEC-associated DEGs. **D–H.** Curated gene panels showing ECM remodeling, angiogenesis, enteric nervous system-associated signaling, motility/contraction, and epithelial-repair programs. In panels **B** and **D–H**, orange gene text indicates upregulation in NEC relative to infant controls, whereas blue gene text indicates downregulation. **I.** Volcano plot of DEGs between fetal and infant controls that define the developmental trajectory for SIP fibroblasts. **J–K.** Gene-level and transcription-factor maturity analyses of fetal-versus-infant DEGs, highlighting the subset also significantly changed in NEC relative to infant controls and positioned relative to fetal and infant reference states, and their corresponding dotplots. The developmental projection score in panels **J–K** is displayed on a symmetric logarithmic (symlog) scale. For expression dot plots, dot size indicates the fraction of nuclei expressing each gene and dot color indicates mean expression. In maturity plots, vertical dotted lines indicate the fetal and infant reference scores. Colored dots classify NEC-versus-infant DEGs as Fetal-like, Between Fetal & Infant, Beyond Fetal State, or Beyond Infant State; gray N/S dots are fetal-versus-infant DEGs without a significant NEC-versus-infant change.

We identified 839 SIP fibroblast nuclei across the atlas, with 323 genes significantly altered in NEC SIP fibroblasts compared to infant controls, including 85 upregulated and 238 downregulated genes (Fig. 17B). GO-term enrichment analysis showed increases in extracellular matrix and collagen-organization programs, hypoxia-angiogenesis, focal adhesion/actin-cytoskeleton and calcium/contractile signaling in NEC, together with depletion of calmodulin-binding and cyclic nucleotide phosphodiesterase programs (Fig. 17C). These GO-term results were reflected in the extracellular matrix, hypoxia-angiogenesis, enteric neuromuscular, and motility programs described below, while curated gene-level analysis also highlighted an epithelial-repair program.

Extracellular-matrix and wound-healing changes were prominent in NEC SIP fibroblasts and could alter the neuromuscular stromal compartment. The upregulated genes included *THBS1*, *CCN2*, *SERPINE1*, *TNC*, *COL1A1*, and *SULF1* (Fig. 17D). *THBS1* is a matricellular protein linked to wound healing and fibrosis, while *CCN2* promotes collagen production and deposition by fibroblasts.^677–680,713^ *SERPINE1*/PAI-1 can limit fibrinolysis and promote collagen accumulation, whereas *TNC* is induced during injury and contributes to intestinal wound repair.^681–682,714^ Increased *COL1A1*, which encodes the α1 chain of type I collagen, and *SULF1*, an extracellular sulfatase linked to fibrosis, further supported collagen-rich and fibrosis-associated matrix changes.^715^ These changes could help stabilize injured tissue during NEC resolution, but if sustained, they could also stiffen the bowel wall, alter mechanotransduction, and interfere with coordinated motility.^716–718^

A hypoxia- and angiogenesis-associated response was also evident in SIP fibroblasts. *EGR1* is an early-response transcription factor induced during hypoxia and positioned upstream of HIF1A signaling, and *VEGFA* is a HIF1A-linked angiogenic factor that promotes blood-vessel growth (Fig. 17E).^604–605^ *IGF2* can promote angiogenesis, *STAT3* promotes VEGF expression and angiogenesis, and *RUNX1* has been linked to angiogenesis in colorectal cancer systems.^719–722^ Upregulation of these genes suggests that NEC SIP fibroblasts may respond to hypoxic vascular stress and attempt to support revascularization or oxygen delivery. This could aid in the resolution of NEC if it improves tissue oxygenation, but persistent angiogenic signaling with matrix remodeling could also mark ongoing hypoxic stress and disease progression.

Altered enteric neuromuscular and motility-associated signaling was another feature of NEC SIP fibroblasts. *PRSS3*, which encodes trypsin-3 and can mediate signaling from intestinal epithelium to enteric neurons, was downregulated (Fig. 17F). *ROBO1* and *ROBO2*, guidance receptors involved in enteric neural crest migration, were also reduced; *ROBO1* has additionally been linked to enteric nervous system changes in Hirschsprung disease.^723–725^ *C3*, a central complement component, was upregulated. Experimental C3 deficiency alters enteric nervous system function and interstitial cell of Cajal distribution, supporting a role for C3 in enteric neuromuscular homeostasis.^726^ Motility-associated genes showed a mixed pattern (Fig. 17G). *RHOA*, a small GTPase that contributes to intestinal smooth muscle contraction, was upregulated, whereas *PRKG1*, which supports cGMP-dependent smooth muscle relaxation, *KCNN3*, encoding the SK3 potassium channel, *PDE5A*, a cyclic nucleotide phosphodiesterase involved in smooth muscle and motility regulation, and *ABCC9*, encoding the SUR2 KATP-channel subunit, were downregulated.^727–733^ These changes suggest altered neuromuscular control in NEC. If they impair motility, they could contribute to disease progression by promoting intestinal stasis or bacterial overgrowth.^734^

Epithelial-support signals were also evident in NEC SIP fibroblasts. *GREM1*, a BMP antagonist produced by intestinal mesenchyme around crypts, supports ISC homeostasis through local BMP inhibition (Fig. 17H).^29,593^ Additional upregulated genes included *CCN1*, which supports epithelial proliferation during mucosal healing; *TNC*, which contributes to intestinal barrier protection and wound repair; *CHI3L1*, which supports epithelial survival and proliferation during inflammation; and *THBS1*, which has been linked to intestinal wound repair.^679,714,735–736^ Together, these increases suggest that NEC SIP fibroblasts may provide epithelial-support cues that could aid NEC resolution when coordinated with vascular recovery and tissue organization.

Of the 982 genes distinguishing fetal from infant control SIP fibroblasts, 147 were also dysregulated in NEC versus infant controls, including 74 with fetal-like expression (50.3% of the 147 genes; Fig. 17I,J). Developmentally regulated genes included fetal-high *GDNF* and *WNT5A*, mesenchymal factors involved in enteric nervous system development and embryonic intestinal morphogenesis, respectively, and infant-high *PDGFRA*, a defining SIP-cell receptor whose expression increases during postnatal SIP maturation and whose signaling contributes to SIP-cell development and maintenance.^737–740^ The fetal-like genes in NEC highlighted changes in cyclic nucleotide signaling, neuropeptide regulation, epithelial-support signaling, and extracellular matrix organization (Fig. 17J). *PDE1A*, a cGMP phosphodiesterase that can limit intestinal smooth muscle relaxation; *MME*/neprilysin, which degrades substance P, a neuropeptide that promotes intestinal contraction; and *GLP2R*, which is expressed by intestinal subepithelial myofibroblasts and mediates growth factor signaling that supports intestinal growth, were all significantly downregulated in NEC and showed fetal-like expression.^696–697,741–744^ By contrast, fetal-high *AGRN*, an extracellular matrix protein shown to regulate enteric neural crest-cell migration during enteric nervous system development, was upregulated in NEC and also showed fetal-like expression.^745^ These changes suggest that NEC SIP fibroblasts may have altered neuromuscular and extracellular matrix-associated signaling that could affect coordinated intestinal motility and repair.

At the transcription factor level, NEC SIP fibroblasts showed altered expression of genes linked to fibroblast activation, proliferation, and differentiation. *THRB* can inhibit fibroblast proliferation, *PPARG* can suppress myofibroblast differentiation and fibrotic collagen production, and *EBF1* marks extracellular matrix-producing and proliferative fibroblast populations (Fig. 17K).^702,746–749^ *FOXO3* can limit fibrosis through apoptosis-linked control of fibroblast persistence, *KLF9* can restrain abnormal fibroblast proliferation, and *PRRX1* promotes fibroblast activation and myofibroblast differentiation.^750–753^ All of these genes were downregulated in NEC and aligned with fetal controls, suggesting that regulators of fibroblast activation, proliferation, and differentiation show expression resembling the fetal state.

NEC SIP fibroblasts show extracellular matrix remodeling, hypoxia-angiogenesis, epithelial-repair, and altered enteric neuromuscular and contractility-associated features. Initiation may be influenced indirectly if SIP signaling contributes to dysmotility, stasis, or altered epithelial-stromal communication in the injured bowel wall. During progression, matrix remodeling and altered neuromuscular signaling could worsen tissue stiffness, motility disruption, and inflammatory repair imbalance. At the same time, epithelial-support and vascular-repair genes suggest an adaptive role during recovery. Resolution may require SIP fibroblasts to support epithelial and vascular repair without maintaining neuromuscular dysfunction or maladaptive matrix remodeling.

### Lamina Propria Myocytes

Lamina propria myocytes (LPMs) are a distinct intestinal smooth muscle population that extends through the villus lamina propria, separate from the muscularis mucosae beneath the crypts and the external muscularis propria (Fig. 18A).^593^ LPMs share a core smooth muscle transcriptional program with the muscularis mucosae, including expression of *Myh11* and *Acta2*, and both populations arise from early PDGFRA+ mesenchymal precursors.^593^ A study distinguished these closely related mucosal smooth muscle populations in mouse by *Grem2*, which was expressed in muscularis mucosae but not LPMs.^593^ Consistent with this shared smooth muscle identity, the LPM population identified in our dataset showed prominent *MYH11*, *ACTA2*, and *TAGLN* expression, while comparatively high *GREM2* expression suggests that human LPMs in our dataset also retain molecular features shared with the muscularis mucosae.

**Fig. 18.**
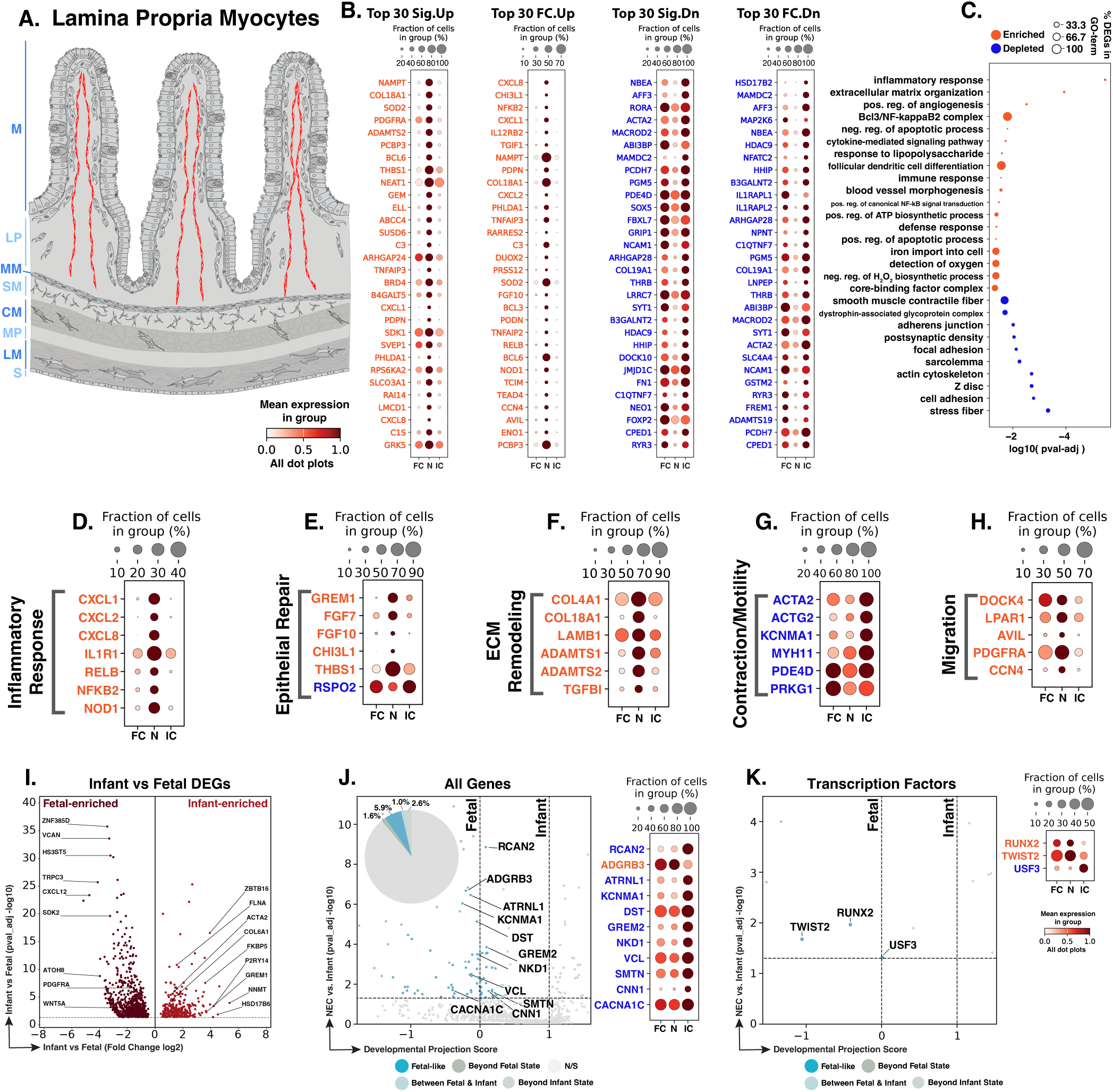
NEC lamina propria myocytes show inflammatory, migratory, extracellular-matrix, epithelial repair, and contractile-associated changes. **A.** Schematic showing the mucosal location of lamina propria myocytes (red). Blue text to the left of the Panel A schematic indicates general anatomical regions: M, mucosa; LP, lamina propria; MM, muscularis mucosae; SM, submucosa; CM, circular muscle; MP, myenteric plexus; LM, longitudinal muscle; S, serosa. **B.** Dot plots of top NEC-specific genes ranked by significance and fold change. **C.** GO-term analysis showing enriched and depleted biological processes among NEC-associated DEGs. **D–H.** Curated gene panels showing inflammatory, epithelial repair, extracellular matrix, contraction/motility, and migration programs. In panels **B** and **D–H**, orange gene text indicates upregulation in NEC relative to infant controls, whereas blue gene text indicates downregulation. **I.** Volcano plot of DEGs between fetal and infant controls that define the developmental trajectory for lamina propria myocytes. **J–K.** Gene-level and transcription-factor maturity analyses of fetal-versus-infant DEGs, highlighting the subset also significantly changed in NEC relative to infant controls and positioned relative to fetal and infant reference states, and their corresponding dotplots. The developmental projection score in panels **J–K** is displayed on a symmetric logarithmic (symlog) scale. For expression dot plots, dot size indicates the fraction of nuclei expressing each gene and dot color indicates mean expression. In maturity plots, vertical dotted lines indicate the fetal and infant reference scores. Colored dots classify NEC-versus-infant DEGs as Fetal-like, Between Fetal & Infant, Beyond Fetal State, or Beyond Infant State; gray N/S dots are fetal-versus-infant DEGs without a significant NEC-versus-infant change.

Despite expressing contractile markers, these cells clustered transcriptionally with our stromal populations rather than with muscularis propria smooth muscle populations, consistent with a specialized mucosal myocyte population that has both stromal and contractile features. Villus smooth muscle cells are closely associated with lacteals and contract to facilitate dietary-fat transport, and this villus musculature develops and is maintained through local mesenchymal progenitor populations.^754–755^ These structural and contractile functions make LPMs relevant to NEC because injury within the villus lamina propria could alter villus smooth muscle organization and its interactions with surrounding mucosal tissue. Prior human NEC single-cell transcriptomic data did not identify a smooth muscle cluster and showed increased fibroblast abundance in NEC, while experimental neonatal mouse NEC multi-omics showed a reduced smooth muscle cell population together with increased fibroblasts; neither study defined LPM-specific injury programs.^18,157^ Thus, it remains unknown whether human NEC LPMs acquire disease-associated changes that could influence mucosal injury and repair.

We identified 1,082 LPM nuclei in our dataset. Differential gene-expression analysis identified 257 genes significantly altered in NEC LPMs compared to infant controls, including 134 upregulated and 123 downregulated genes (Fig. 18B). GO-term enrichment analysis between NEC and infant controls showed that NEC LPMs were enriched for inflammatory signaling, LPS/NF-kappaB response, extracellular matrix organization, and angiogenesis programs and depleted for focal adhesion, actin cytoskeleton, stress fiber, cell adhesion, and smooth muscle contraction terms (Fig. 18C). These GO-term results were reflected in the inflammatory, migration/matrix remodeling, and contraction programs described below, along with an increase in epithelial repair genes.

Inflammatory activation in NEC LPMs could amplify innate immune signaling within injured villus tissue. *CXCL1*, *CXCL2*, and *CXCL8*, CXC chemokines expressed by inflammatory intestinal fibroblasts with neutrophil-chemoattractant activity, were significantly upregulated in NEC LPMs.^597^ *IL1R1*, the receptor for the pro-inflammatory cytokine IL-1, was also significantly upregulated, supporting the potential for increased responsiveness to IL-1 signaling.^597^ Upregulated *RELB* and *NFKB2*, components of non-canonical NF-kappaB signaling, were consistent with an NF-kappaB-associated inflammatory transcriptional state (Fig. 18D).^756–757^ This response could help villus-associated LPMs recruit neutrophils after epithelial barrier disruption, but sustained chemokine- and NF-kappaB-associated signaling could maintain inflammatory pressure within the villus lamina propria and contribute to the progression of NEC.

Epithelial-support factors showed divergent regulation in NEC LPMs. *RSPO2*, an R-spondin ligand that potentiates Wnt signaling and supports Lgr5+ intestinal stem-cell self-renewal, was significantly downregulated (Fig. 18E).^758^ By contrast, *GREM1*, a BMP antagonist produced by crypt-associated intestinal mesenchymal cells, was significantly upregulated. Epithelial BMP signaling restrains Lgr5+ ISC self-renewal, and *Grem1* upregulation has been shown to support adaptive epithelial reprogramming during intestinal regeneration.^29,669,759^ *FGF7* and *FGF10* were also significantly upregulated (Fig. 18E). *FGF7* is a fibroblast-derived epithelial growth factor that can promote intestinal epithelial proliferation and protect against gastrointestinal injury, whereas *FGF10* supports intestinal epithelial progenitor survival and proliferation and promotes small-intestinal ulcer healing.^620,671,760–763^ This pattern suggests that NEC LPMs may alter the balance of epithelial-support cues, with reduced RSPO-associated Wnt potentiation alongside increased BMP-antagonist and FGF signaling potential. These changes could influence epithelial renewal and repair during NEC.

Basement-membrane, extracellular matrix, and stromal-migration genes were increased in NEC LPMs. *COL4A1*, which encodes a type IV collagen chain required for basement membrane stability, *COL18A1*, which encodes collagen XVIII, and *LAMB1*, which encodes a laminin β1 chain found in intestinal basement membrane zones, were significantly upregulated.^764–767^ *ADAMTS2*, a procollagen N-proteinase, and *TGFBI*, a basement membrane-associated glycoprotein linked to tissue repair, were also significantly upregulated (Fig. 18F).^768–769^ Genes associated with stromal movement were increased in parallel. *DOCK4* supports PDGF-dependent fibroblast migration, *LPAR1* mediates fibroblast recruitment after injury, and *AVIL* promotes cell migration.^770–772^ *PDGFRA*, which regulates fibroblast responses after tissue injury, and *CCN4*, which promotes fibroblast migration and tissue repair, were likewise significantly upregulated (Fig. 18H).^674,773^ This pattern suggests that LPMs may become more migratory during NEC, facilitating movement within injured villus tissue. The accompanying basement membrane and extracellular matrix changes could help restore the villus extracellular matrix and epithelial-mesenchymal interface during NEC resolution. If sustained alongside persistent inflammation, however, they could also contribute to disorganized matrix repair, abnormal stromal movement, or disrupted epithelial-stromal organization.

Reduced expression of smooth muscle contraction genes was another feature of NEC LPMs. *ACTA2* is an important component of intestinal smooth muscle contraction, while *ACTG2* contributes to actin organization and function in human intestinal smooth muscle cells (Fig. 18G).^774–775^ *MYH11* similarly encodes a core component of the smooth muscle contractile apparatus and is required for regulated smooth muscle contraction.^776^ Other downregulated genes regulate the balance between contraction and relaxation*. PDE4D* can promote smooth muscle contraction through cAMP-dependent signaling, whereas *KCNMA1*-encoded BK channels and *PRKG1*-dependent cGMP signaling contribute to smooth muscle relaxation.^692–695,777^ Together, reduced expression of contractile machinery genes alongside altered regulators of smooth muscle excitability and relaxation suggests disruption of contractile signaling in NEC LPMs. Because reduced intestinal motility occurs early in experimental NEC and restoring motility can reduce disease severity, these changes could contribute to NEC initiation or progression by altering intestinal transit.^24^

Of the 1,152 genes distinguishing fetal from infant control LPMs, 128 were also dysregulated in NEC versus infant controls, including 68 with fetal-like expression (53.1% of the 128 genes; Fig. 18I,J). Developmentally regulated genes included fetal-high *PDGFRA* and infant-high *ACTA2*, consistent with developmental progression from *PDGFRA*+ subepithelial mesenchymal progenitors toward *ACTA2*+ contractile villus smooth muscle.^754,778^ The fetal-like genes in NEC highlighted changes in BMP/Wnt regulatory signaling and reduced contractile and mechanical features (Fig. 18J). *GREM2*, a BMP antagonist expressed by intestinal mesenchymal and smooth muscle populations, and *NKD1*, a Wnt-inducible negative-feedback regulator, were both significantly downregulated in NEC and showed fetal-like expression.^593,699,779^ *SMTN*, which is required for intestinal smooth muscle contractility; *CNN1*, which modulates smooth muscle contractility; *CACNA1C*/CaV1.2, which is essential for intestinal smooth muscle electromechanical coupling and motility; and *VCL*, which participates in adhesion-cytoskeletal coupling required for smooth muscle tension development, were also significantly downregulated in NEC and showed fetal-like expression.^689–691,780–782^ These developmental changes, together with the NEC-associated changes described above, suggest that LPMs may have altered epithelial-support signaling and reduced mature contractile and mechanical features while expressing increased inflammatory, migratory, and basement membrane repair features.

At the transcription factor level, NEC LPMs showed fetal-like expression of genes associated with fibroblast migration and matrix remodeling (Fig. 18K). *TWIST2* has been shown to support fibroblast migration during fetal wound healing and was significantly upregulated in NEC, with expression similar to fetal control (Fig. 18K).^783^ *RUNX2* was also significantly upregulated in NEC, with fetal-like expression (Fig. 18K). Runx2 has been shown to regulate IGF-associated mesenchymal niche homeostasis in the mouse incisor, and also directly promotes pathological fibroblast formation and fibrotic gene expression in pulmonary fibrosis, raising the possibility that increased *RUNX2* could contribute to fibrotic matrix remodeling in NEC LPMs.^784–785^

NEC lamina propria myocytes show altered epithelial-support signaling and immature contractile features within the villus lamina propria. Initiation may be influenced by reduced RSPO2 and GREM2, which suggest weaker Wnt-potentiating and BMP-antagonist support for epithelial renewal. During disease progression, increased inflammatory, migratory, FGF-associated, and matrix-remodeling genes may sustain villus stromal activation while reduced contractile and mechanical features weaken mucosal support. Repair signals may still aid epithelial recovery, especially through FGF-associated and basement membrane-related changes. Resolution may depend on whether these myocytes regain mature contractile and epithelial-support functions while inflammatory and matrix-remodeling activity subsides.

### Subepithelial Myofibroblasts

Subepithelial myofibroblasts (SEMFs), also called telocytes, are elongated mesenchymal cells located directly beneath the intestinal epithelium along the crypt-villus axis marked by Foxl1 (Fig. 19A).^786–788^ In this position, SEMFs contribute to the epithelial niche by providing signaling cues, including BMP and Wnt pathway ligands, that help coordinate epithelial renewal, crypt maintenance, and epithelial-mesenchymal communication.^29,786–789^ Although the SEMF population in our dataset was not defined solely by *FOXL1* expression, its high expression of *PDGFRA*, *NRG1*, *F3*, *WNT5B*, *BMP5*, and *BMP7* aligns with transcriptional features of SEMFs and telocytes described in mouse and human studies.^29,47–48,592–593,786^ Prior human NEC transcriptomic data and experimental neonatal mouse NEC multi-omics showed an increased proportion of fibroblasts, but did not characterize SEMF-specific changes in epithelial niche signaling in NEC.^18,157^ Thus, it remains unresolved whether SEMFs acquire transcriptional programs that could influence NEC progression or resolution.

**Fig. 19.**
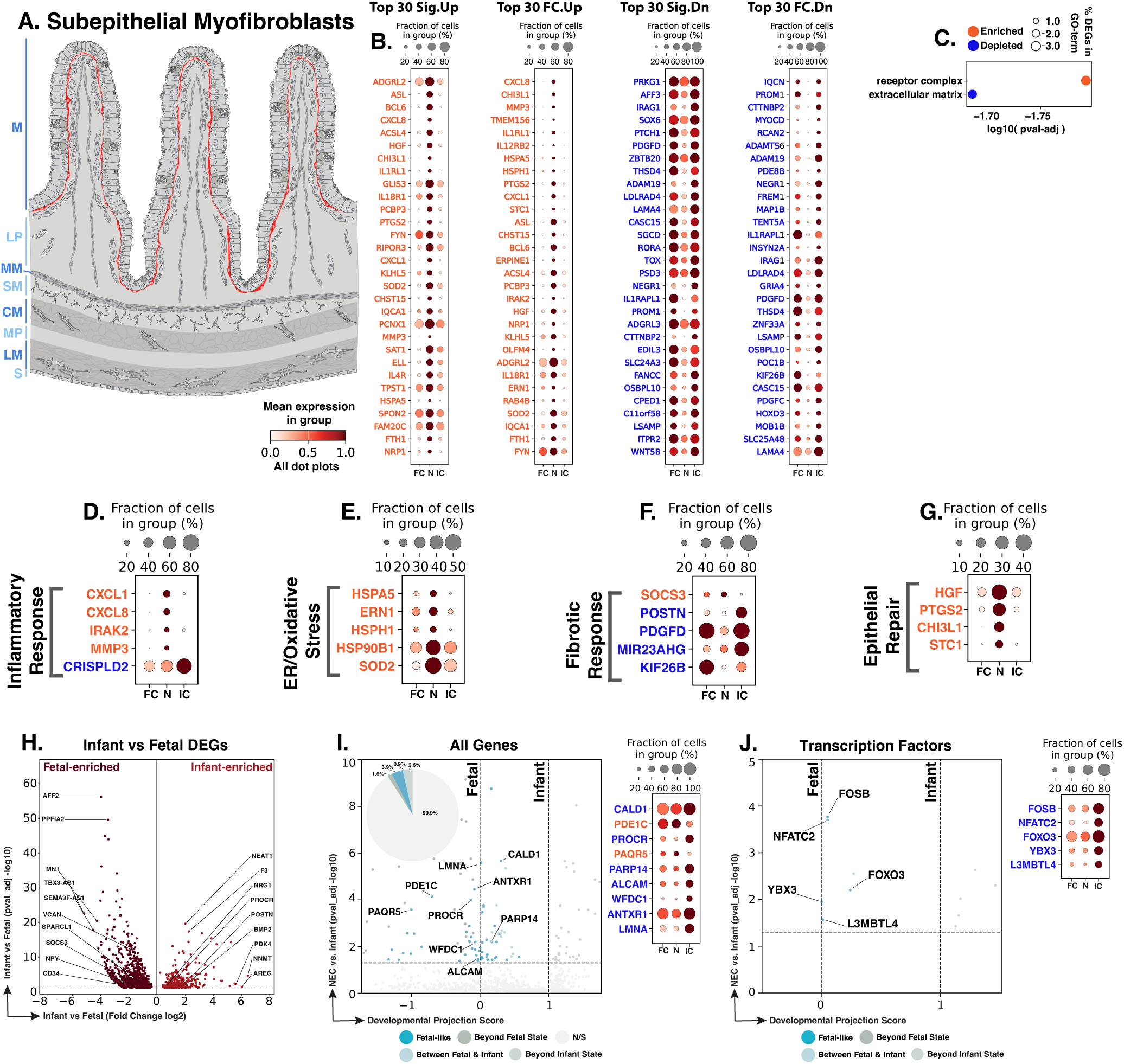
NEC subepithelial myofibroblasts show inflammatory and stress activation with altered fibrotic and epithelial-repair programs. **A.** Schematic showing the subepithelial distribution of subepithelial myofibroblasts (red) along the crypt-villus axis. Blue text to the left of the Panel A schematic indicates general anatomical regions: M, mucosa; LP, lamina propria; MM, muscularis mucosae; SM, submucosa; CM, circular muscle; MP, myenteric plexus; LM, longitudinal muscle; S, serosa. **B.** Dot plots of top NEC-specific genes ranked by significance and fold change. **C.** GO-term analysis showing enriched and depleted biological processes among NEC-associated DEGs. **D–G.** Curated gene panels showing inflammatory-response, stress-response, fibrotic-response, and epithelial-repair programs. In panels **B** and **D–G**, orange gene text indicates upregulation in NEC relative to infant controls, whereas blue gene text indicates downregulation. **H.** Volcano plot of DEGs between fetal and infant controls that define the developmental trajectory for subepithelial myofibroblasts. **I–J.** Gene-level and transcription-factor maturity analyses of fetal-versus-infant DEGs, highlighting the subset also significantly changed in NEC relative to infant controls and positioned relative to fetal and infant reference states, and their corresponding dotplots. The developmental projection score in panels **I–J** is displayed on a symmetric logarithmic (symlog) scale. For expression dot plots, dot size indicates the fraction of nuclei expressing each gene and dot color indicates mean expression. In maturity plots, vertical dotted lines indicate the fetal and infant reference scores. Colored dots classify NEC-versus-infant DEGs as Fetal-like, Between Fetal & Infant, Beyond Fetal State, or Beyond Infant State; gray N/S dots are fetal-versus-infant DEGs without a significant NEC-versus-infant change.

We identified 2,012 subepithelial myofibroblast nuclei in our dataset, and using differential gene expression analysis, identified 211 genes significantly changed in NEC SEMFs compared to infant controls. This included 72 upregulated and 139 downregulated genes (Fig. 19B). Because of the low number of differentially expressed genes, GO-term enrichment was limited, with receptor complex as the only significantly enriched term and extracellular matrix as the only depleted term in NEC (Fig. 19C). Because these GO-term results were sparse, the gene programs below are used to connect significant SEMF DEGs to broader processes that could play a role in NEC pathogenesis or resolution.

Inflammatory changes in NEC SEMFs could amplify immune signaling beneath the injured epithelium. Increased *CXCL1* and *CXCL8* encode CXC chemokines that can promote neutrophil recruitment when expressed by intestinal stromal cells and fibroblasts (Fig. 19D).^596,680,790^ *IRAK2*, an interleukin-1 receptor-associated kinase that can promote nuclear export of selected inflammatory mRNAs after LPS stimulation in macrophages, was also upregulated, suggesting increased responsiveness to innate inflammatory signaling.^791^ Increased *MMP3*, a matrix metalloproteinase that can promote inflammatory signaling, together with reduced *CRISPLD2*, an anti-inflammatory secreted protein, further pointed to increased inflammatory signaling associated with the extracellular matrix.^792–795^ This pattern suggests that SEMFs may contribute to immune recruitment and inflammatory signaling near the epithelial surface. Such a response could help subepithelial SEMFs respond to microbial exposure during epithelial breaches in NEC, but persistent chemokine and matrix-associated inflammatory signaling could sustain inflammation beneath the epithelium and contribute to disease progression.

A separate stress-response pattern in NEC SEMFs was consistent with increased ER and oxidative stress. Higher *HSPA5*, which encodes GRP78/BiP and is a canonical ER-stress marker, was consistent with increased ER protein-folding stress (Fig. 19E).^629^ *ERN1*, an ER-stress sensor, and *HSPH1*, which encodes HSP105 and has been linked to ER-stress responses, were also upregulated.^687,796^ Increased *HSP90B1*, which encodes the ER chaperone GRP94 and is induced by ER stress, further supported increased protein-folding and quality control demand.^630–631^ Higher *SOD2*, a mitochondrial antioxidant enzyme inducible by oxidative stress, suggested additional antioxidant adaptation.^632–633^ These changes suggest that NEC SEMFs may increase stress-buffering responses to preserve stromal function during inflammatory and hypoxic injury. If sustained, however, these protein-folding and oxidative-stress features could reflect unresolved epithelial-stromal injury and impaired NEC resolution.

Fibrosis-associated genes showed a different pattern, with several changes consistent with reduced fibrotic signaling in NEC SEMFs compared with infant controls. *SOCS3*, a suppressor of cytokine signaling whose depletion can promote fibrosis, was upregulated, suggesting increased restraint of fibrotic signaling (Fig. 19F).^797^ In contrast, *POSTN*, a matricellular protein that promotes fibrosis, and *PDGFD*, a driver of fibrogenesis, were downregulated.^798–799^ *KIF26B*, a kinesin family protein linked to fibroblast fibrosis in nonintestinal systems, was also reduced.^800^ This pattern is consistent with less fibrotic signaling in NEC SEMFs despite their inflammatory activation. Reduced fibrotic signaling could limit maladaptive matrix accumulation and aid NEC resolution, but it could also weaken subepithelial matrix repair if SEMFs fail to rebuild an organized extracellular matrix scaffold after epithelial injury.

Epithelial-support signals were also evident in NEC SEMFs. Increased *HGF*, a fibroblast-derived epithelial mitogen that promotes intestinal epithelial proliferation, suggested greater epithelial support (Fig. 19G).^801^ *PTGS2*, which encodes cyclooxygenase-2, was also upregulated and marks prostaglandin-producing stromal cells that promote intestinal epithelial proliferation during injury.^802^ Higher *CHI3L1*, a glycoprotein that promotes epithelial survival and proliferation during intestinal inflammation and can be secreted by fibroblasts, added another potential epithelial-protective signal.^736,803^ *STC1*, an oxidative-stress-responsive mesenchymal factor that can promote re-epithelialization, was likewise increased.^804–805^ These genes suggest that SEMFs may provide epithelial survival, proliferation, and restitution cues during NEC. This response could support disease resolution, although its effectiveness may depend on whether epithelial support is accompanied by restoration of tissue organization.

Of the 1,300 genes distinguishing fetal from infant control SEMFs, 120 were also dysregulated in NEC versus infant controls, including 51 with fetal-like expression (42.5% of the 120 genes; Fig. 19H,I). Developmentally regulated genes included fetal-high *CD34* and infant-high *BMP2* and *NRG1*, highlighting mesenchymal factors that contribute to intestinal niche organization and epithelial differentiation.^47,806^ The fetal-like genes in NEC highlighted reduced epithelial support and cytoskeletal/contractile features together with increased senescence-associated features (Fig. 19I). *PROCR*/CD201, an infant-high mesenchymal marker linked to intestinal morphogenesis, epithelial proliferation, and epithelial differentiation, and *CALD1*, an actin-binding regulator of fibroblast contractility, focal adhesions, and cytoskeletal organization, were both significantly downregulated in NEC and showed fetal-like expression.^565,648,807^ *ANTXR1*, a stromal receptor whose deficiency promotes fibroblast senescence and disrupts nuclear architecture and actin dynamics, and *LMNA*, whose loss in fibroblasts activates DNA damage and senescence responses, were also downregulated and showed fetal-like expression.^808–809^ These changes suggest that NEC SEMFs have reduced epithelial-support features together with increased senescence-associated characteristics, which could impair epithelial-stromal repair and contribute to disease progression.

At the transcription factor level, NEC SEMFs showed fetal-like expression of genes linked to fibroblast proliferation, activation, and fibrotic regulation (Fig. 19J). *FOSB*, an AP-1 family immediate-early transcription factor induced during fibroblast matrix contraction and linked to fibroblast cell-cycle progression, was downregulated in NEC and aligned with the fetal reference state.^810–811^ *NFATC2*, which contributes to hypoxia-induced fibroblast proliferation, was also downregulated and showed fetal-like expression.^703^ Reduced *FOXO3*, a forkhead transcription factor that can restrain fibroblast activation and fibrosis and promote collagen-matrix-induced apoptosis in fibrotic fibroblasts, likewise aligned with the fetal reference state.^750–751^ *YBX3*, a Y-box binding regulator expressed by activated fibroblasts, was similarly downregulated and showed fetal-like expression.^812^ These changes suggest that NEC SEMFs may incompletely acquire a mature regulatory state associated with fibroblast proliferation, activation, and fibrotic restraint, which could affect epithelial-stromal repair.

NEC SEMFs acquire inflammatory, stress-response, and epithelial-repair features while showing reduced fibrotic matrix signals and fetal-like changes in support, cytoskeletal, and senescence-associated genes. Initiation may involve subepithelial chemokine and innate-inflammatory signaling that responds to epithelial injury or microbial exposure. During progression, persistent stress and inflammatory signaling beneath the epithelium could maintain epithelial-stromal injury. Upregulated epithelial-support genes such as HGF, PTGS2, CHI3L1, and STC1 suggest a potential repair role. Resolution may fail if these epithelial-support signals are not matched by mature stromal organization, matrix recovery, and stable epithelial-stromal integration.

### Proliferative Stroma

Proliferative stromal cells are cycling mesenchymal cells found across intestinal stromal compartments, with proliferation being particularly prominent during intestinal development as mesenchymal populations expand and mature alongside the growing intestine (Fig. 20A).^47–49^ This population likely reflects a proliferative state within the broader stromal compartment, rather than a terminal stromal lineage.^49^ Although these cells may reflect normal intestinal growth and maturation, they could also respond to inflammatory cues, hypoxic stress, and epithelial-mesenchymal signaling during NEC, potentially contributing to mucosal repair, epithelial restitution, and angiogenic adaptation.^587,813–814^ Previous human neonatal NEC transcriptomic and mouse neonatal NEC multi-omics studies did not specifically characterize proliferating stromal populations.^18,157^ Thus, it remains unclear whether proliferative stromal cells in NEC reflect ongoing intestinal development and maturation alone or also acquire additional transcriptional programs associated with inflammatory stress and tissue repair.

**Fig. 20.**
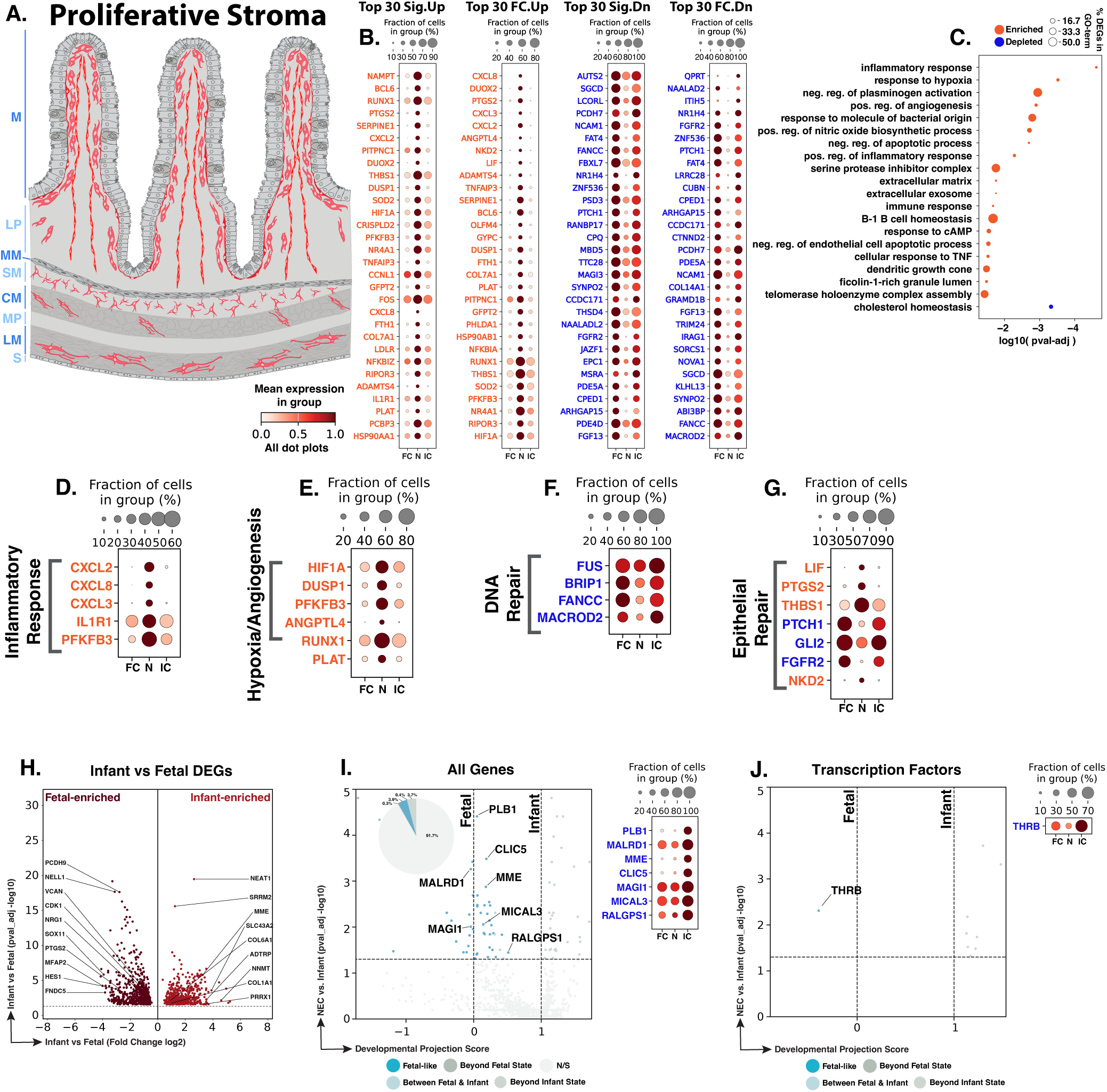
NEC proliferative stroma show inflammatory, hypoxia/angiogenesis, DNA-repair, and epithelial-repair-associated changes. **A.** Schematic showing the distribution of proliferative stromal cells (red) across the intestinal stroma. Blue text to the left of the Panel A schematic indicates general anatomical regions: M, mucosa; LP, lamina propria; MM, muscularis mucosae; SM, submucosa; CM, circular muscle; MP, myenteric plexus; LM, longitudinal muscle; S, serosa. **B.** Dot plots of top NEC-specific genes ranked by significance and fold change. **C**. GO-term analysis showing enriched and depleted biological processes among NEC-associated DEGs. **D–G.** Curated gene panels showing inflammatory-response/neutrophil-recruitment, hypoxia/angiogenesis, DNA-repair, and epithelial-repair programs. In panels **B** and **D–G**, orange gene text indicates upregulation in NEC relative to infant controls, whereas blue gene text indicates downregulation. **H.** Volcano plot of DEGs between fetal and infant controls that define the developmental trajectory for proliferative stromal cells. **I–J.** Gene-level and transcription-factor maturity analyses of fetal-versus-infant DEGs, highlighting the subset also significantly changed in NEC relative to infant controls and positioned relative to fetal and infant reference states, and their corresponding dotplots. The developmental projection score in panels **I–J** is displayed on a symmetric logarithmic (symlog) scale. For expression dot plots, dot size indicates the fraction of nuclei expressing each gene and dot color indicates mean expression. In maturity plots, vertical dotted lines indicate the fetal and infant reference scores. Colored dots classify NEC-versus-infant DEGs as Fetal-like, Between Fetal & Infant, Beyond Fetal State, or Beyond Infant State; gray N/S dots are fetal-versus-infant DEGs without a significant NEC-versus-infant change.

We identified 749 proliferative stromal nuclei across the atlas, with 165 genes significantly altered in NEC proliferative stromal cells compared to infant controls, including 46 upregulated and 119 downregulated genes (Fig. 20B). GO-term enrichment analysis showed NEC enrichment for inflammatory and innate-defense, hypoxia-angiogenesis and nitric-oxide, extracellular matrix, and cell-survival programs, together with depletion of cholesterol homeostasis in NEC (Fig. 20C). These GO-term results were reflected in the inflammatory and hypoxia-angiogenesis programs described below, while curated gene-level analysis also highlighted DNA repair and epithelial-repair programs.

Inflammatory and neutrophil-recruitment signals were prominent in NEC proliferative stromal cells and could establish innate immune cues near injured mucosal compartments. *CXCL2*, *CXCL3*, and *CXCL8* encode chemokines that can be released by fibroblasts to recruit neutrophils, consistent with a stromal contribution to neutrophil-rich inflammation in NEC (Fig. 20D).^596,815–816^ *IL1R1*, the receptor for IL-1, was also upregulated, suggesting increased responsiveness to IL-1-family inflammatory signaling.^817–818^ Increased *PFKFB3*, a glycolytic regulator induced by inflammatory and hypoxic cues, further supported this response; higher stromal *PFKFB3* expression has also been linked to intestinal inflammation.^291–292,819^ These changes could support microbial containment after epithelial barrier disruption. However, persistent chemokine and IL-1-responsive signaling could sustain neutrophil recruitment and inflammatory amplification, contributing to NEC progression.^600,663,815^

A distinct hypoxia- and angiogenesis-associated response was also evident in NEC proliferative stroma. *HIF1A*, a central regulator of cellular hypoxia responses, was upregulated, while increased *DUSP1* and *PFKFB3* provided additional evidence of a hypoxia-responsive state (Fig. 20E).^241,291–292,608^ *ANGPTL4*, *RUNX1*, and *PLAT* were likewise upregulated and have been linked to endothelial angiogenesis or angiogenic responses in other contexts.^722,820–821^ This response could support NEC resolution if it improves perfusion and oxygen delivery. Because it occurs alongside inflammatory gene upregulation, however, it may also reflect persistent hypoxic injury without fully effective vascular recovery.

Reduced DNA repair-associated gene expression pointed to a potential vulnerability in this cycling stromal population. *FUS*, *BRIP1*, *FANCC*, and *MACROD2* were significantly downregulated in NEC (Fig. 20F). *FUS* has been linked to repair of oxidative DNA breaks, *BRIP1* and *FANCC* participate in DNA repair and Fanconi-anemia pathways, and *MACROD2* regulates PARP1 activity and DNA-break repair.^822–826^ Because these cells are proliferative, reduced expression of DNA repair genes could be especially important in the inflammatory and hypoxic NEC environment. These findings suggest that stromal expansion may occur during injury while genome-protective gene expression is incompletely maintained.

Epithelial-support signals in NEC proliferative stromal cells were mixed. *PTGS2* marks an injury-associated prostaglandin response, and *Ptgs2*-expressing stromal cells can promote intestinal epithelial proliferation during injury (Fig. 20G).^802^ *THBS1*, which can be secreted by fibroblasts and has been linked to intestinal wound repair, and *LIF*, a fibroblast-derived cytokine that supports epithelial regeneration and ISC homeostasis, were also significantly upregulated.^622–623,679,827^ In contrast, *PTCH1* and *GLI2*, components of Hedgehog-responsive mesenchymal signaling downstream of epithelial Hedgehog ligands, were reduced.^634,638,828^ Because mesenchymal Hedgehog signaling regulates intestinal stem-cell niche cues and epithelial proliferation and differentiation, reduced *PTCH1* and *GLI2* could weaken coordinated epithelial regeneration.^638,828^ Lower *FGFR2* may similarly indicate reduced responsiveness to epithelial FGF signals that regulate intestinal mesenchymal proliferation, differentiation, and reciprocal epithelial-mesenchymal signaling.^829–830^ Increased *NKD2*, a negative regulator of WNT signaling, could further restrain WNT-associated regenerative crosstalk.^831–834^ Overall, proliferative stroma expresses epithelial-support signals alongside reduced Hedgehog/FGF responsiveness and potential restraint of WNT-associated regenerative signaling.

Of the 1,049 genes distinguishing fetal from infant control proliferative stromal cells, 87 were also dysregulated in NEC versus infant controls, including 41 with fetal-like expression (47.1% of the 87 genes; Fig. 20H,I). Developmentally regulated genes included fetal-high *HES1* and *CDK1*, which have been linked to human intestinal mesenchymal development and cell-cycle progression, respectively, and infant-high *PRRX1*, a transcription factor that promotes fibroblast progression toward a myofibroblast state.^753,835–836^

The fetal-like genes in NEC highlighted changes in focal adhesion, cytoskeletal organization, and proliferation (Fig. 20I). *MAGI1*, a focal adhesion regulator whose loss impairs focal adhesion maturation and actin stress-fiber formation; *MICAL3*, an actin-disassembly factor involved in F-actin reorganization and cytokinesis; and *CLIC5*, which associates with the cortical actin cytoskeleton and supports F-actin-based structures, were all significantly downregulated in NEC and showed fetal-like expression.^837–840^ *THRB*, a nuclear receptor shown to inhibit fibroblast proliferation through regulation of G1/S cell-cycle genes, was the only transcription factor that showed fetal-like expression in NEC and was also significantly downregulated relative to infant controls (Fig. 20J).^746–747^ These changes suggest that NEC proliferative stromal cells retain fetal-like focal adhesion and cytoskeletal features together with reduced anti-proliferative regulation, which could impair coordinated stromal repair.

NEC proliferative stromal cells represent a conflicted stromal repair state. Initiation may involve chemokine and IL-1-responsive signals from cycling stromal cells that support early microbial containment after barrier disruption. Progression may occur if stromal proliferation remains linked to inflammatory amplification, hypoxia, reduced DNA repair genes, altered Hedgehog/FGF responsiveness, and potential WNT restraint. These features suggest that stromal expansion occurs under injury conditions but may not fully mature into organized mucosal rebuilding. NEC resolution may require proliferative stroma to transition from inflammatory cycling toward stable matrix repair, epithelial support, and genome-protective capacity.

### Pericytes

Pericytes are mural stromal cells embedded along capillaries and small vessels, where they wrap endothelial tubes and help regulate vascular stability, endothelial barrier integrity, vessel tone and tissue perfusion, and angiogenic sprouting (Fig. 21A).^841–845^ In the intestine, pericytes are positioned along the capillary network in the small-intestinal villus stroma, placing them at the interface between circulating inflammatory mediators, endothelial transport, perfusion, and epithelial injury.^846–847^ Prior NEC studies support reduced intestinal VEGF signaling and defective VEGFR2-dependent microvascular development as vascular contributors to disease susceptibility, but no pericyte-specific role.^609,848^ In addition, human neonatal NEC single-cell RNA sequencing and mouse neonatal NEC multi-omics data contain fibroblast populations and vascular/lymphatic endothelium but did not report a pericyte population.^18,157^ Although these studies have emphasized epithelial injury, immune activation, and endothelial dysfunction in NEC, the pericyte-specific response remains less defined. Thus, it remains unresolved whether human NEC pericytes preserve vascular support functions or acquire associated features that could influence intestinal perfusion, vascular leakage, and repair during disease.

**Fig. 21.**
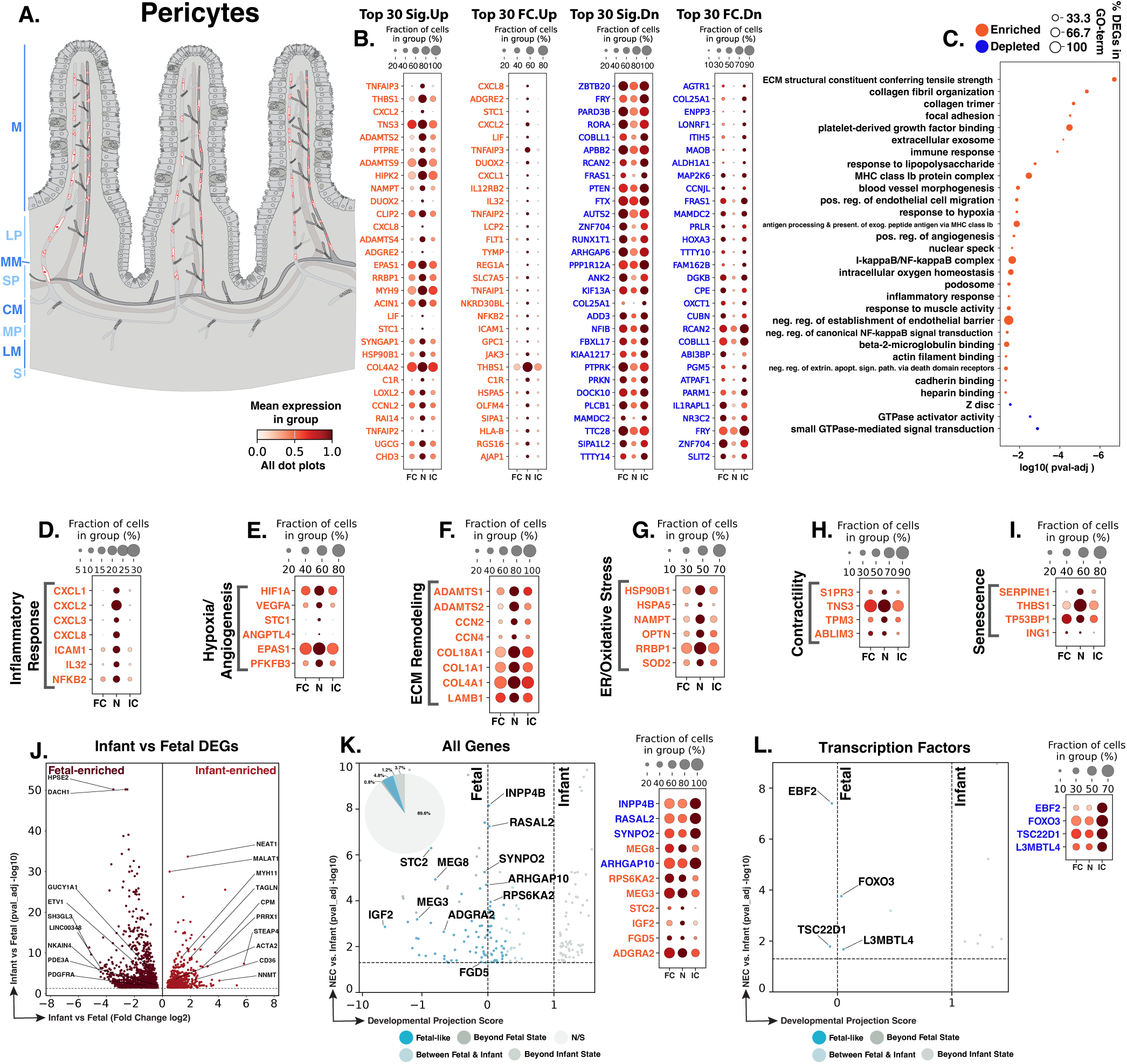
NEC pericytes show inflammatory, hypoxia/angiogenesis, ECM remodeling, contractility, and senescence-associated changes. **A.** Schematic showing the vascular-associated distribution of intestinal pericytes (red). Blue text to the left of the Panel A schematic indicates general anatomical regions: M, mucosa; LP, lamina propria; MM, muscularis mucosae; SP, submucosal plexus; CM, circular muscle; MP, myenteric plexus; LM, longitudinal muscle; S, serosa. **B.** Dot plots of top NEC-specific genes ranked by significance and fold change. **C.** GO-term analysis showing enriched and depleted biological processes among NEC-associated DEGs. **D–I.** Curated gene panels showing inflammatory-response, hypoxia/angiogenesis, extracellular matrix, ER/oxidative stress, contractility, and senescence-associated programs. In panels **B** and **D–I**, orange gene text indicates upregulation in NEC relative to infant controls, whereas blue gene text indicates downregulation. **J.** Volcano plot of DEGs between fetal and infant controls that define the developmental trajectory for pericytes. **K– L.** Gene-level and transcription-factor maturity analyses of fetal-versus-infant DEGs, highlighting the subset also significantly changed in NEC relative to infant controls and positioned relative to fetal and infant reference states, and their corresponding dotplots. The developmental projection score in panels **K–L** is displayed on a symmetric logarithmic (symlog) scale. For expression dot plots, dot size indicates the fraction of nuclei expressing each gene and dot color indicates mean expression. In maturity plots, vertical dotted lines indicate the fetal and infant reference scores. Colored dots classify NEC-versus-infant DEGs as Fetal-like, Between Fetal & Infant, Beyond Fetal State, or Beyond Infant State; gray N/S dots are fetal-versus-infant DEGs without a significant NEC-versus-infant change.

We identified 1,748 pericyte nuclei across the atlas. Differential expression analysis identified 441 genes significantly changed in NEC pericytes compared to infant controls, including 206 upregulated and 235 downregulated genes (Fig. 21B). GO-term enrichment analysis showed enrichment of extracellular matrix/collagen and focal adhesion programs, inflammatory/antigen presentation and NF-κB programs, vascular morphogenesis/hypoxia/oxygen-homeostasis and angiogenesis programs, and junctional functions in NEC, together with depletion of Z-disc and small-GTPase regulatory/signaling programs (Fig. 21C). Enrichment for negative regulation of endothelial barrier establishment and cadherin binding may indicate altered endothelial barrier support, as pericyte-endothelial N-cadherin signaling can directly restrict endothelial permeability.^842^ This may be particularly relevant in NEC, where vascular barrier dysfunction has been observed in experimental NEC.^849^ Consistent with these patterns, NEC pericytes showed gene-expression programs associated with inflammation, hypoxia/angiogenesis, ECM remodeling, contractility, and stress/senescence.

Inflammatory changes in NEC pericytes could promote leukocyte recruitment around injured intestinal vessels. Increased *CXCL1*, *CXCL2*, *CXCL3*, and *CXCL8* support a neutrophil-recruiting chemokine response, while increased *ICAM1* supports leukocyte adhesion and pericyte-associated extravasation into inflamed tissue (Fig. 21D).^850–852^ Consistent with this pattern, LPS-stimulated pericytes have been shown to increase *CXCL1*, *CXCL2*, *CXCL3*, *CXCL8*, and *ICAM1* expression together with increased nuclear NF-κB localization.^850^ Increased *IL32*, a pro-inflammatory cytokine linked to NF-κB activation, provided additional evidence of inflammatory signaling.^853^ Persistent chemokine and adhesion signaling could contribute to NEC progression by promoting vascular-associated neutrophil entry and increasing local inflammatory pressure.^851–852^

A distinct hypoxia- and angiogenesis-associated response was also evident in NEC pericytes. Increased *HIF1A* and *VEGFA* were consistent with HIF1A-linked angiogenic signaling during hypoxia (Fig. 21E).^604^ Higher *STC1* may reflect a response to ischemic injury, while increased *ANGPTL4* and *EPAS1* add angiogenesis-associated signaling, with EPAS1 linked to *VEGFA*/FLT1 regulation.^820,854–855^ These changes suggest that NEC pericytes may respond to hypoxic vascular stress by increasing angiogenesis-associated signals that could support vascular recovery. Persistent hypoxia-associated signaling, however, may also reflect ongoing vascular stress.

Basement-membrane and extracellular matrix genes were increased in NEC pericytes, suggesting changes in perivascular tissue organization. *ADAMTS2*, a procollagen N-proteinase, *COL18A1*, which encodes the basement membrane proteoglycan collagen XVIII, and *COL4A1*, which encodes a type IV collagen chain required for basement membrane stability, were significantly upregulated (Fig. 21F).^764–766,768^ *CCN4*, which can promote cell migration and wound healing, and *LAMB1*, which encodes a laminin β1 chain found in intestinal basement membrane zones, were also increased.^767,773^ These changes could help reconstruct and stabilize the perivascular basement membrane after injury. If excessive or poorly coordinated, however, they could alter pericyte-endothelial interactions, vessel structure, and effective vascular repair.

Cellular stress responses provided another feature of the NEC pericyte state. *HSP90B1*, which encodes the ER chaperone GRP94 and is induced by ER stress, and *HSPA5*, which encodes GRP78/BiP and is a canonical ER-stress marker, were significantly upregulated, consistent with increased ER protein-folding stress (Fig. 21G).^629–631^ *SOD2*, a mitochondrial antioxidant enzyme inducible by oxidative stress, was also increased, suggesting greater antioxidant defense.^632–633^ These responses may help pericytes maintain perivascular function during inflammatory, hypoxic, and oxidative injury. If unresolved, however, persistent cellular stress could compromise pericyte function and endothelial support, potentially limiting restoration of a stable intestinal microvasculature during NEC resolution.

Contractility-associated genes were also increased in NEC pericytes and could influence vascular tone. Higher *S1PR3* supports sphingosine-1-phosphate receptor signaling involved in vascular contraction (Fig. 21H).^856^ Increased *TNS3* may strengthen focal adhesion links between the cytoskeleton and extracellular matrix, which are important for force transmission in pericytes and vascular smooth muscle cells.^857–859^ Increased *TPM3* suggests greater tropomyosin-associated actin contractile machinery, while increased *ABLIM3* may support coupling of adherens junctions to circumferential actin bundles.^860–862^ These changes suggest a more contractile vascular-support state in NEC pericytes. Because increased vasoconstrictor responsiveness and reduced microvascular perfusion have been observed in experimental NEC, greater pericyte contractility could contribute to disease progression by further limiting perfusion in already hypoxic tissue.^863–864^

Senescence-associated genes added another potential vulnerability to the NEC pericyte response. Increased *SERPINE1*, which encodes PAI-1, supports senescence-associated signaling, while increased *THBS1* has been linked to endothelial cell senescence (Fig. 21I).^865–867^ Increased *TP53BP1* and *ING1* further supported DNA damage and senescence-associated cell state regulation.^868–869^ Pericyte senescence has been reported to impair vascular barrier function in the brain, suggesting that a similar stress state in NEC could weaken microvascular stability or repair capacity.^870^ Thus, NEC pericytes combine inflammatory and angiogenesis-associated responses with senescence-associated features that could reduce their ability to maintain a stable vascular niche during resolution.

Of the 1,694 genes distinguishing fetal from infant control pericytes, 177 were also dysregulated in NEC versus infant controls, including 81 with fetal-like expression (45.8% of the 177 genes; Fig. 21J,K). Developmentally regulated genes included fetal-high *PDGFRA*, which has been associated with primitive mesenchymal mural cell progenitors, and infant-high *ACTA2* and *TAGLN*, which are associated with mural-cell differentiation and maturation.^871–873^ The fetal-like genes in NEC highlighted angiogenic and pericyte-migration features (Fig. 21K). *ADGRA2*/GPR124, which promotes pericyte polarization and directional migration through regulation of filopodia formation, and *IGF2*, an angiogenic growth factor that promotes endothelial migration and capillary-like network formation, were both significantly upregulated in NEC and showed fetal-like expression.^719,874^ *STC2*, which has been shown to regulate endothelial angiogenesis, and *FGD5*, which supports VEGFA signaling and endothelial migration, were also upregulated in NEC and showed expression similar to fetal controls.^875–876^ These changes suggest that NEC pericytes have fetal-like angiogenic features that could affect microvascular stabilization and repair and potentially contribute to disease pathogenesis.

At the transcription factor level, fetal-like genes in NEC pericytes were associated with vascular maturation and proliferation (Fig. 21L). Reduced *FOXO3* may influence vessel formation, maturation, and endothelial cell apoptosis.^877–878^ Reduced *TSC22D1* may indicate altered vascular signaling because TSC22D1 contributes to TGF-β- and PDGF-dependent signaling in vascular smooth muscle cells and has been implicated in endothelial dysfunction and microvascularization.^879–880^ Reduced *L3MBTL4* may similarly suggest lower vascular smooth muscle or mural-cell proliferative and migratory potential.^881^ The reduced expression of these genes in NEC, together with their similarity to fetal controls, suggests that NEC pericytes retain fetal-like regulatory features associated with vascular maturation and proliferation.

NEC pericytes show inflammatory chemokine signaling, hypoxia/angiogenesis, matrix remodeling, cellular stress, senescence-associated features, and increased contractility-associated genes. Initiation may be influenced by vascular-associated leukocyte entry and altered microvascular tone near injured epithelium. During progression, pericyte inflammation, contractility, and senescence could worsen perfusion defects, endothelial barrier instability, and local vascular injury. Hypoxia-angiogenic and matrix-remodeling genes suggest an attempted vascular repair response, but this may be maladaptive if it does not restore stable pericyte-endothelial support. Resolution may depend on recovery of mature pericyte function, endothelial stabilization, and effective microvascular perfusion.

### Endothelial Compartment

The intestinal endothelial compartment, marked by *PECAM1* and *VWF* (Fig. 2), contains blood vascular and lymphatic endothelial cells that regulate tissue perfusion, vascular exchange, immune cell trafficking, and fluid drainage. Endothelial dysfunction is mechanistically implicated in NEC at several levels. Intestinal VEGF protein is reduced in human and experimental NEC, and impaired VEGFR2/KDR signaling disrupts neonatal intestinal microvascular development and increases susceptibility to NEC-like injury.^609,848^ Endothelial-specific TLR4 signaling impairs intestinal microcirculatory perfusion, while developmental insufficiency of intestinal arginine-nitric-oxide signaling limits postprandial microvascular blood flow in the immature intestine.^611–612^ Prenatal inflammatory exposure further reduces VEGF/VEGFR2 expression, endothelial proliferation, and microvascular density. Conversely, macrophage-derived IGF-1 preserves intestinal VEGF expression, endothelial proliferation, and microvascular development and protects neonatal mice from experimental NEC.^610,882^ Prior human NEC transcriptomic analysis also identified endothelial activation, including increased endothelial *CCL2*, *CXCL1*, *CXCL2*, *CXCL3*, and *ICAM1*.^18^

To define the endothelial populations in our study, heatmap-based annotation of marker-gene expression resolved four endothelial populations, including *DLL4*+/*GJA5*+/*SEMA3G*+ arterial endothelial cells, *PLVAP*+/*SPARCL1*+/*EBF3*+ venous endothelial cells, *MKI67*+/*MELK*+/*KNL1*+ proliferative vascular endothelial cells, and *PROX1*+/*LYVE1*+ lymphatic endothelial cells (Fig. 22). This organization extends the vascular and lymphatic populations identified in prior human NEC single-cell work by distinguishing arterial and venous identities and a proliferative vascular state.^18^

**Fig. 22.**
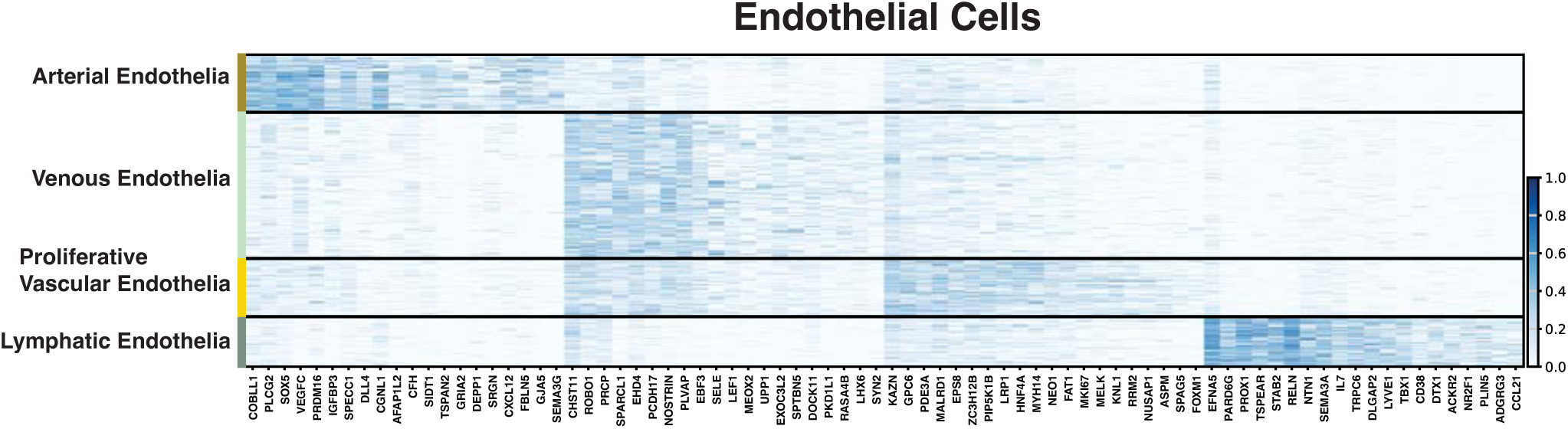
Endothelial cell-type signatures define arterial, venous, proliferative vascular, and lymphatic endothelial populations. Heatmap showing marker-gene expression across arterial, venous, proliferative vascular, and lymphatic endothelial cells. Distinct transcriptional signatures support endothelial cell-type assignments and provide the framework for the detailed endothelial analyses that follow.

The following sections examine NEC-associated transcriptional changes across these four endothelial populations. Arterial endothelial cells showed changes in hypoxia- and perfusion-associated pathways, angiogenic signaling, arterial specification, and genes supporting vessel stability. Venous endothelial cells concentrated inflammatory adhesion and chemokine signals together with procoagulant, antifibrinolytic, and barrier-associated changes. Lymphatic endothelial cells combined inflammatory signaling with altered expression of genes involved in scavenging, transport, mechanoregulation, and immune cell trafficking. Proliferative vascular endothelial cells retained a strong mitotic program alongside angiogenic and matrix-associated responses consistent with endothelial repair. Together, these sections frame NEC endothelium as an injury-responsive vascular system in which inflammatory entry and repair signaling increase, but vessel stabilization, lymphatic clearance, and functional recovery may remain incomplete.

### Arterial Endothelium

Arterial endothelial cells line intestinal arteries and arterioles and regulate tissue and vascular tone. In the intestine, endothelial vasodilation contributes to absorptive hyperemia, while endothelial growth and repair sustain the vascular network required for oxygen delivery and epithelial regeneration (Fig. 23A).^883–884^ This vascular support is impaired in NEC: intestinal VEGF protein is reduced in human and experimental disease, and disruption of VEGFR2/KDR signaling compromises neonatal intestinal microvascular development and increases susceptibility to NEC-like injury.^609,848^ Human single-cell analysis also identified an inflamed vascular-endothelial population expressing adhesion molecules and neutrophil-recruiting chemokines, but arterial and venous endothelial cells were analyzed together, leaving the arterial contribution unresolved.^18^

**Fig. 23.**
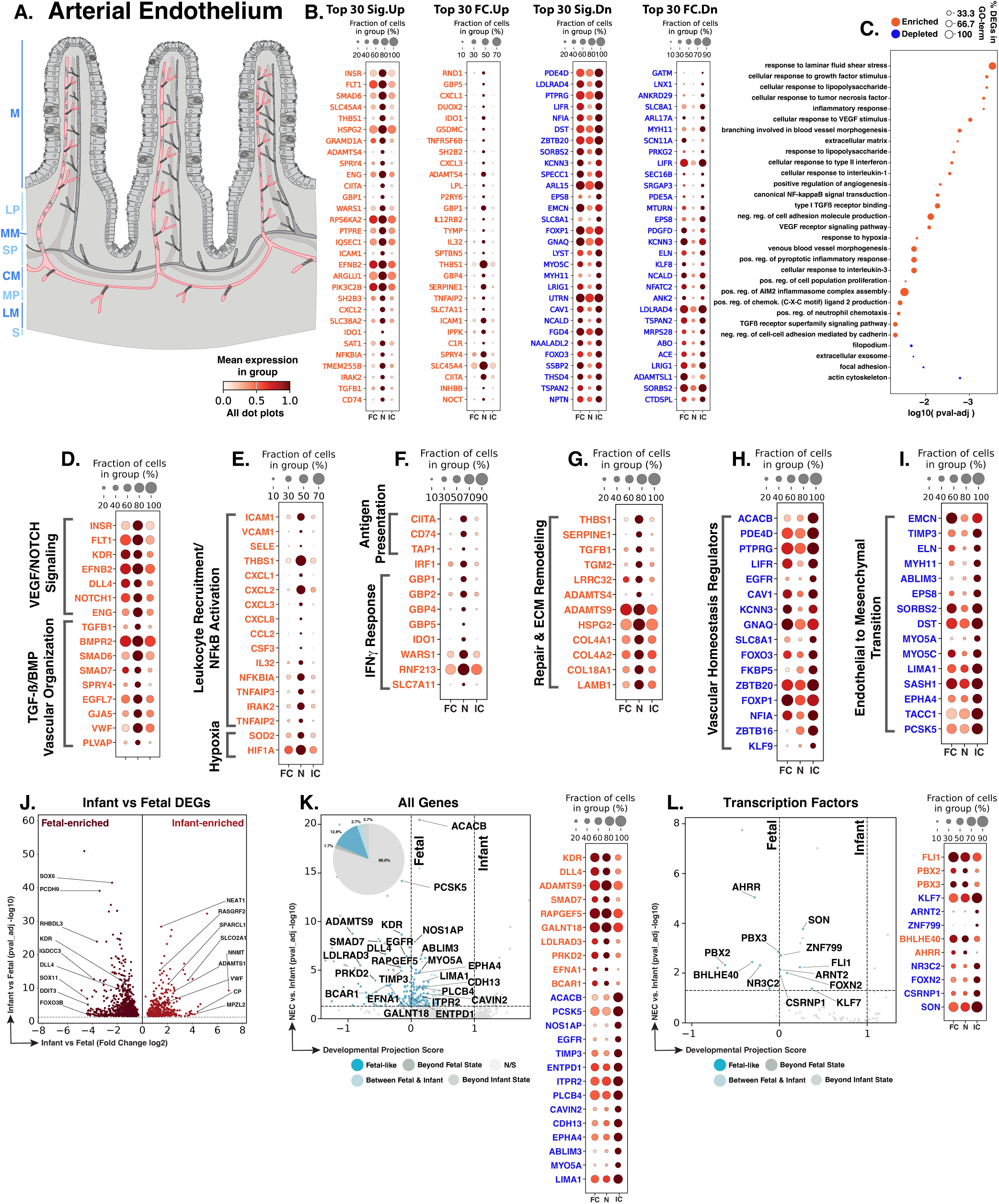
NEC arterial endothelial cells show inflammatory, angiogenic, matrix-remodeling, vascular-homeostatic, and maturity-associated changes. **A.** Schematic showing the arterial endothelial distribution (red) within the intestinal vasculature. Blue text to the left of the Panel A schematic indicates general anatomical regions: M, mucosa; LP, lamina propria; MM, muscularis mucosae; SP, submucosal plexus; CM, circular muscle; MP, myenteric plexus; LM, longitudinal muscle; S, serosa. **B.** Dot plots of top NEC-specific genes ranked by significance and fold change. **C.** GO-term analysis showing enriched and depleted biological processes among NEC-associated DEGs. **D–I.** Curated gene panels showing angiogenesis/receptor-ligand and TGF-β/BMP arterial remodeling, inflammatory-response/hypoxia, antigen-presentation and IFNγ-response, repair/matrix remodeling, vascular-homeostasis, and endothelial-to-mesenchymal-transition-associated programs. In panels **B** and **D–I**, orange gene text indicates upregulation in NEC relative to infant controls, whereas blue gene text indicates downregulation. **J.** Volcano plot of DEGs between fetal and infant controls that define the developmental trajectory for arterial endothelial cells. **K–L.** Gene-level and transcription-factor maturity analyses of fetal-versus-infant DEGs, highlighting the subset also significantly changed in NEC relative to infant controls and positioned relative to fetal and infant reference states, and their corresponding dotplots. The developmental projection score in panels **K–L** is displayed on a symmetric logarithmic (symlog) scale. For expression dot plots, dot size indicates the fraction of nuclei expressing each gene and dot color indicates mean expression. In maturity plots, vertical dotted lines indicate the fetal and infant reference scores. Colored dots classify NEC-versus-infant DEGs as Fetal-like, Between Fetal & Infant, Beyond Fetal State, or Beyond Infant State; gray N/S dots are fetal-versus-infant DEGs without a significant NEC-versus-infant change.

We identified 1,040 arterial endothelial nuclei across the atlas. Differential-expression analysis identified 534 genes significantly changed in NEC compared to infant controls, including 287 upregulated and 247 downregulated genes (Fig. 23B). GO-term enrichment connected the upregulated response to laminar-shear and growth factor sensing, VEGF signaling, branching-vessel morphogenesis, inflammatory responses to LPS, TNF, and IL-1, canonical NF-κB signaling, neutrophil chemotaxis, hypoxia, TGF-β-superfamily signaling, and extracellular matrix remodeling. These responses were accompanied by depletion of filopodial, focal adhesion, actin-cytoskeletal, and extracellular-vesicle programs (Fig. 23C). The arterial response therefore combines inflammatory and angiogenic activation with reduced representation of structural and homeostatic programs needed to stabilize the remodeling vessel wall.

NEC arterial endothelial cells showed increased expression of genes associated with VEGF, Notch, and TGFβ/BMP signaling. Increased *KDR*/VEGFR2, a receptor required for postnatal intestinal microvascular development, occurred with increased *FLT1*/VEGFR1, which modulates the strength and spatial distribution of VEGFR2 signaling during vessel formation (Fig. 23D).^609,885^ Because intestinal VEGF availability is reduced in NEC, increased receptor expression may represent an attempt by surviving arterial endothelial cells to recover growth factor responsiveness rather than evidence that effective VEGF signaling has been restored. Within this receptor-responsive state, increased *DLL4* and *NOTCH1* align with endothelial sprouting and arterial specification, while increased *EFNB2* reinforces arterial identity, and increased *GJA5* is consistent with an arteriogenic response.^886–889^ These changes indicate reinforcement of arterial identity-associated gene expression associated with an active remodeling state within the NEC arterial endothelial population.

This VEGF/Notch response was integrated with TGF-β/BMP signaling. Increased *ENG* and *TGFB1* support endothelial TGF-β-pathway engagement: endoglin regulates endothelial activation and quiescence, while TGF-β1 can reshape VEGF-dependent endothelial growth and survival responses.^890–891^ Increased *BMPR2* is likewise consistent with signaling that supports endothelial survival and monolayer integrity.^892^ Simultaneous induction of the inhibitory SMADs SMAD6 and SMAD7 suggests that this pathway is accompanied by feedback restraint, limiting unrestricted TGF-β/BMP activation.^893–894^ A similar balance is suggested by increased *SPRY4*, which constrains endothelial migration through integrin-associated signaling, together with EGFL7-dependent vascular-tube organization, VWF-linked angiogenic control, and PLVAP-associated fenestral exchange.^895–898^ Collectively, these genes associated with the GO-term demonstrate a VEGF- and TGF-β-responsive program in surviving arterial endothelial cells, consistent with an angiogenic repair response to NEC intestinal injury.

The reparative program developed within a strongly inflammatory arterial surface. Increased *ICAM1* and *VCAM1* support firm leukocyte adhesion and transendothelial recruitment, while increased *SELE* supports the preceding leukocyte-rolling step (Fig. 23E).^899–900^ The accompanying increase in *CXCL1*, *CXCL2*, and *CXCL3* establishes a GRO-family neutrophil-directed chemokine program, reinforced by increased *CXCL8*.^52,901^ Increased *CCL2* extends this recruitment axis toward monocytes, whereas *CSF3* may reinforce neutrophil production and mobilization.^902–903^ These findings extend previous human NEC observations by resolving this inflammatory response specifically within arterial endothelial cells.^18^

Inflammatory pathway activation was accompanied by induction of endogenous NF-κB regulatory mechanisms. Increased *IRAK2* supports TLR-dependent inflammatory signaling, whereas *NFKBIA*, *TNFAIP2*, and *TNFAIP3* represent inducible negative-feedback regulators activated during NF-κB signaling.^400^ Increased *SOD2* and *HIF1A* further indicated antioxidant and hypoxia-responsive transcription.^241,904^ Within this inflammatory and NF-κB response, enrichment of GO terms related to AIM2 inflammasome assembly and pyroptotic inflammatory responses, together with increased *GSDMC* expression suggests potential sensitivity to pyroptosis. GSDMC can mediate caspase-8-dependent membrane permeabilization, and microvascular endothelial cells can undergo NLRP3-linked pyroptosis under inflammatory stimulation.^113,905^ Additional validation would be required to determine if pyroptosis is ongoing in the arterial endothelium.

Arterial endothelial cells also acquired an interferon-associated immune-interaction state. Increased *CIITA* and *CD74* supported greater MHC class II processing and presentation capacity, consistent with the ability of human intestinal microvascular endothelial cells to present complex protein antigens through MHC class II, while increased *TAP1* extended this response to cytokine-inducible MHC class I peptide loading (Fig. 23F).^906–908^ Increased *IRF1* and coordinated upregulation of *GBP1*, *GBP2*, *GBP4*, and *GBP5* further supported an interferon-responsive vascular state; GBP1 and GBP5 are also inducible in activated human endothelium.^909–911^ Increased *IDO1* and *WARS1* linked this response to tryptophan metabolism, while *RNF213* added an interferon-responsive vascular-stress and angiogenic component.^912–914^ Increased *SLC7A11* may further support cystine uptake and glutathione-dependent antioxidant buffering.^915^ Overall, NEC arterial endothelial cells show an interferon-responsive state with increased antigen-processing machinery and stress-buffering features that could enhance endothelial-immune interactions at the vascular interface.

NEC arterial endothelial cells also increased expression of extracellular matrix and vascular basement membrane genes (Fig. 23G). *THBS1* was increased; its encoded protein can activate latent TGFβ and contributes to inflammatory and wound-associated matrix regulation.^916–917^ Increased *SERPINE1*, encoding PAI-1, implicated regulation of fibrinolysis,^918^ while increased *TGM2*, *LRRC32*, and *ADAMTS4* further implicated matrix crosslinking, TGFβ regulation, and extracellular proteolysis. ADAMTS9 was also increased and is known to be strongly inducible by IL-1β and TNFα in experimental inflammatory settings, with NF-κB and MAPK signaling contributing to its transcriptional induction.^919–921^ Its increased expression is therefore consistent with the strong IL-1/TNF-associated inflammatory program identified in NEC arterial endothelium. Importantly, ADAMTS9 also functions as a cell-autonomous inhibitor of angiogenesis in microvascular endothelial cells, where its loss increases endothelial migration and tube formation.^922^ *HSPG2*, *COL4A1*, *COL4A2*, *COL18A1*, and *LAMB1* were concurrently increased, indicating extensive vascular basement membrane remodeling. Thus, matrix remodeling occurs together with induction of ADAMTS9, which may provide an endogenous restraint on the concurrent angiogenic response during inflammatory endothelial activation.

NEC arterial endothelial cells showed reduced expression of genes involved in endothelial signaling and vascular homeostasis (Fig. 23H). *CAV1* was reduced, potentially affecting caveolar organization, endothelial nitric oxide signaling, permeability, and vascular responsiveness.^923^ *FOXO3* was reduced, consistent with diminished oxidative-stress resistance and vascular homeostasis. *FOXP1* was also reduced. Endothelial FOXP1 promotes proliferation, migration, and vascular sprouting while also restraining inflammatory and inflammasome-associated transcription.^924–925^ Its reduction in NEC may indicate loss of regulatory control needed to coordinate endothelial growth, migration, and inflammatory signaling. Reductions in *PDE4D*, *GNAQ*, *SLC8A1*, and *KCNN3* further involved cyclic nucleotide regulation, calcium signaling, ion handling, and endothelial membrane-potential control. *EGFR* was also reduced, together with *FKBP5* and several transcriptional regulators including *ZBTB20*, *NFIA*, *ZBTB16*, and *KLF9*. Together with increased VEGF/Notch-associated and extracellular matrix remodeling programs, these changes indicate that angiogenic activation in NEC occurs alongside reduced expression of genes supporting endothelial homeostasis, signal integration, and vascular stability.

This functional imbalance extended into the endothelial cytoskeleton and junctional architecture. Reduced *EMCN* may alter endothelial surface organization and VEGFR2-dependent migration and morphogenesis, whereas reduced *TIMP3* may weaken extracellular matrix and angiogenic restraint (Fig. 23I).^926–928^ Reduced *ABLIM3*, *EPS8*, *LIMA1*, and *SORBS2* converge more directly on the actin-junction interface and vascular-lumen architecture: ABLIM3 participates in anchoring adherens junctions to actin; EPS8 regulates VE-cadherin-associated junctional remodeling; LIMA1/EPLIN stabilizes the linkage between endothelial junctions and the actin cytoskeleton; and SORBS2/ArgBP2 participates in actomyosin control during vascular-lumen formation.^862,929–931^ Reduced *SASH1* adds loss of an endothelial scaffold that couples TLR4 signaling to inflammatory activation and migration.^932^ Reduced *MYO5A* and *MYO5C* further indicate weaker actin-based vesicle transport and secretory trafficking.^933–934^ Most junctional, cytoskeletal, and trafficking genes shown in the panel were reduced, along with *TACC1* and *PCSK5*. *ELN* and *MYH11*—genes associated with vascular elastic architecture and smooth muscle contractile function—were also reduced.^695,935^ This transcriptional profile combines inflammatory and growth-associated gene induction with reduced expression of genes involved in junction-actin coupling, intracellular trafficking, and vascular-wall organization.

Of the 1,261 genes distinguishing fetal from infant control arterial endothelial cells, 252 were also dysregulated in NEC versus infant controls, including 162 with fetal-like expression (64.3% of the 252 genes; Fig. 23J,K). Developmentally regulated genes included fetal-high *KDR* and *DLL4*, associated with angiogenic and arterial-patterning functions, and infant-high *SPARCL1*, *SLCO2A1*, and *VWF*, associated with endothelial quiescence, prostaglandin transport, and hemostatic function.^609,887,936–938^ The fetal-like genes in NEC highlighted increased angiogenic and arterial-patterning signals along with reduced postnatal vessel-stabilizing features (Fig. 23K). *KDR*, *DLL4*, *ADAMTS9*, and *SMAD7* were increased in NEC and showed fetal-like expression, consistent with greater angiogenic, arterial-patterning, matrix-regulatory, and TGF-β-feedback signaling.^609,887,894,922^ By contrast, *EGFR*, *TIMP3*, *ENTPD1*, *CAVIN2*, and *LIMA1*, which normally increased from fetal to infant, were reduced in NEC and also showed fetal-like expression. These changes suggest weaker postnatal EGFR-dependent growth regulation, matrix restraint, purinergic signaling, caveolar organization, and junction-actin coupling.^927,930,939–941^

At the transcription factor level, *FLI1* and *BHLHE40* also showed fetal-like expression in NEC (Fig. 23L). FLI1 may support endothelial-lineage identity and vascular homeostatic regulation during injury, whereas BHLHE40 may support hypoxia-associated proliferation and angiogenesis.^942–943^ Taken together, the maturity analysis suggests that NEC arterial endothelial cells show fetal-like angiogenic features while expressing lower levels of genes associated with postnatal vessel stabilization. This combination may sustain endothelial growth and repair while providing less of the signaling and structural support needed to stabilize the arterial wall.

NEC arterial endothelium shows strong inflammatory and angiogenic remodeling but reduced vascular-stabilizing capacity. Initiation may be influenced by altered perfusion, leukocyte-adhesion signals, chemokine production, and reduced homeostatic endothelial signaling. During NEC progression, persistent VEGF/Notch and TGF-β/BMP responsiveness, interferon-associated immune interaction, basement membrane production, and inflammatory surface activation may sustain vascular remodeling without restoring stable flow. At the same time, reductions in caveolar, cyclic nucleotide, calcium/potassium, junction-actin, vesicle-trafficking, matrix-restraint, and antioxidant genes suggest weaker vessel stabilization. Resolution may fail if angiogenic repair does not mature into a quiescent, well-perfused arterial wall.

### Venous Endothelium

Venous endothelial cells line veins and venules that return blood from intestinal tissue, while postcapillary venules provide the principal site of leukocyte adhesion and transendothelial migration during inflammation (Fig. 24A).^944–945^ In the human intestine, MADCAM1 is constitutively expressed by lamina propria venules and binds α4β7 integrin to support mucosal lymphocyte homing.^946–947^ Previous human NEC single-cell analysis identified increased *SELE*, *ICAM1*, *CCL2*, *CXCL1*, *CXCL2*, and *CXCL3* expression and predicted enhancement of MADCAM1-ITGB7 interactions within a broad vascular-endothelial population.^18^ These findings place venous endothelium at a major blood-tissue gateway for inflammatory cell recruitment in NEC, although its specific transcriptional response has not previously been separated from the arterial endothelium.

**Fig. 24.**
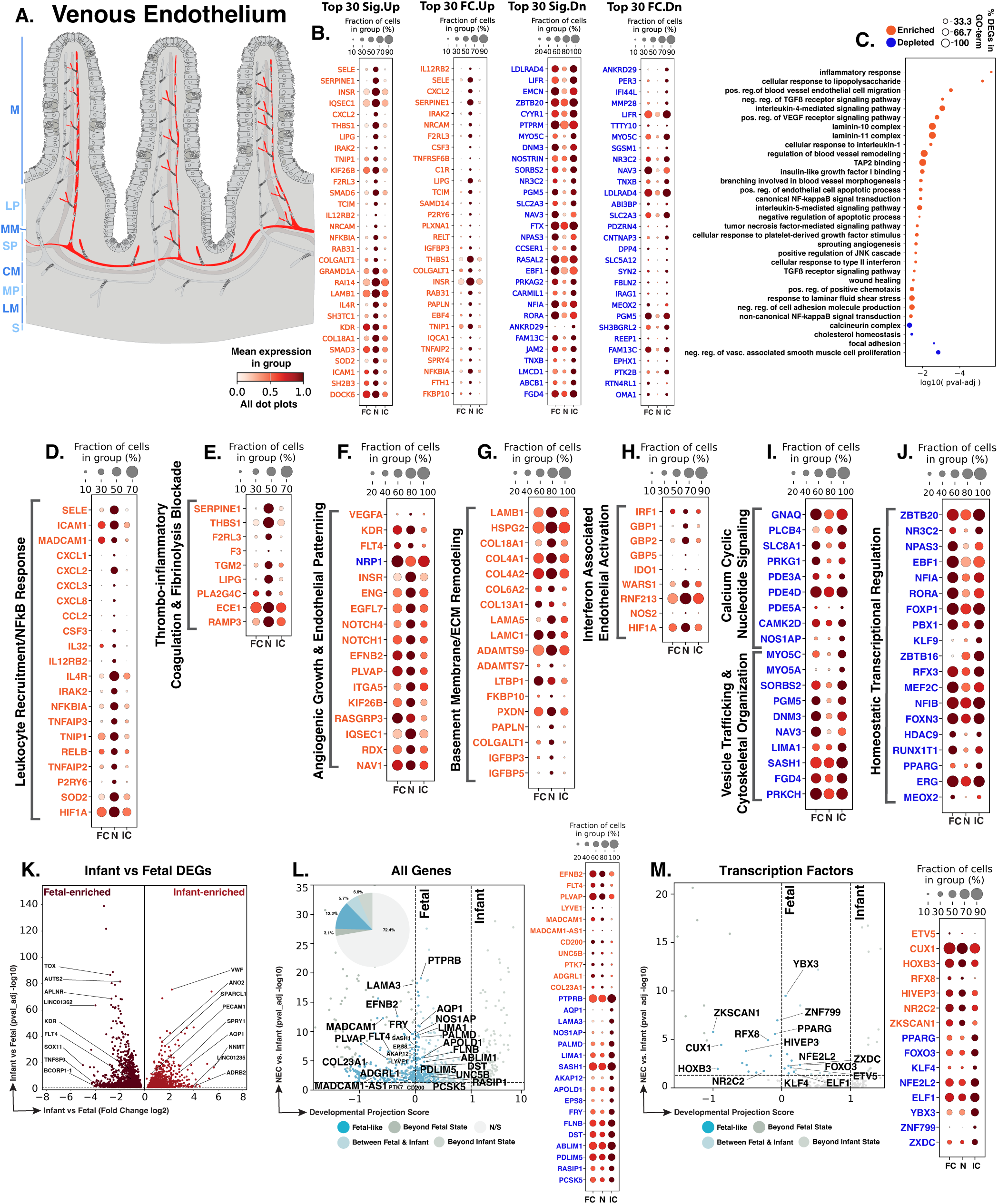
NEC venous endothelial cells show inflammatory and thrombo-inflammatory activation with angiogenic, matrix, vascular-regulatory, and maturity-associated changes. **A.** Schematic showing the venous endothelial distribution (red) within the intestinal vasculature. Blue text to the left of the Panel A schematic indicates general anatomical regions: M, mucosa; LP, lamina propria; MM, muscularis mucosae; SP, submucosal plexus; CM, circular muscle; MP, myenteric plexus; LM, longitudinal muscle; S, serosa. **B.** Dot plots of top NEC-specific genes ranked by significance and fold change. **C.** GO-term analysis showing enriched and depleted biological processes among NEC-associated DEGs. **D–J.** Curated gene panels showing inflammatory-response/leukocyte-recruitment, thrombo-inflammatory/coagulation and fibrinolysis-blockade, angiogenic/endothelial patterning, basement-membrane/ECM, interferon-associated endothelial activation, calcium/cyclic-nucleotide and vesicle/cytoskeletal regulation, and homeostatic transcriptional-regulation programs. In panels **B** and **D–J**, orange gene text indicates upregulation in NEC relative to infant controls, whereas blue gene text indicates downregulation. **K.** Volcano plot of DEGs between fetal and infant controls that define the developmental trajectory for venous endothelial cells. **L–M.** Gene-level and transcription-factor maturity analyses of fetal-versus-infant DEGs, highlighting the subset also significantly changed in NEC relative to infant controls and positioned relative to fetal and infant reference states, and their corresponding dotplots. The developmental projection score in panels **L–M** is displayed on a symmetric logarithmic (symlog) scale. For expression dot plots, dot size indicates the fraction of nuclei expressing each gene and dot color indicates mean expression. In maturity plots, vertical dotted lines indicate the fetal and infant reference scores. Colored dots classify NEC-versus-infant DEGs as Fetal-like, Between Fetal & Infant, Beyond Fetal State, or Beyond Infant State; gray N/S dots are fetal-versus-infant DEGs without a significant NEC-versus-infant change.

We identified 2,635 venous endothelial nuclei across the atlas. Differential-expression analysis identified 1,555 NEC-versus-infant DEGs, including 807 upregulated and 748 downregulated genes (Fig. 24B). GO-term enrichment identified LPS-, TNF-, IL-1-, NF-κB-, interferon-, and hypoxia-associated responses together with VEGF signaling, endothelial migration, angiogenesis, TGFβ regulation, and basement membrane organization. Focal-adhesion and vascular-homeostatic pathways were depleted (Fig. 24C). Thus, venous endothelial cells shared inflammatory, stress, angiogenic, and matrix-associated responses with arterial endothelium, while showing prominent changes associated with leukocyte trafficking, coagulation, and endothelial barrier organization.

NEC venous endothelial cells showed increased expression of adhesion molecules and chemokines involved in leukocyte recruitment (Fig. 24D). SELE and ICAM1 mediate leukocyte capture and firm adhesion to the endothelium, respectively, and expression of *SELE*, *ICAM1*, and *MADCAM1* can be induced in stressed human intestinal microvascular endothelial cells to promote leukocyte recruitment in the injured zone.^948^ Increased *MADCAM1* provides an intestine-specific adhesion pathway for α4β7-expressing lymphocytes.^946–947^ *CXCL1*, *CXCL2*, and *CXCL3* can promote leukocyte arrest under flow, while *CXCL8* and *CCL2* recruit monocytes.^949–951^ *CSF3*, which promotes neutrophil production and mobilization, was also increased.^952^ This profile is consistent with inflammatory activation of venous endothelium, with increased expression of adhesion molecules and chemokines involved in leukocyte recruitment.

NEC venous endothelial cells showed increased expression of inflammatory signaling genes together with inducible negative-feedback regulators. Increased *IRAK2* supports TLR-dependent inflammatory signaling, while increased *RELB* is consistent with engagement of NF-κB-associated transcription.^953–954^ *NFKBIA*, *TNFAIP3*, and *TNIP1* were also increased, representing well-established feedback regulators induced during NF-κB activation.^955–957^ *TNFAIP2* was concurrently increased as an inflammation-responsive NF-κB target.^958^ P2RY6 was also increased and is known to be induced by TNF-α, IL-1, and LPS and responds to extracellular UDP released during tissue injury, amplifying NF-κB-dependent chemokine and adhesion-molecule expression.^959^ Increased *SOD2* and *HIF1A* further indicated antioxidant and hypoxia-responsive signaling.^960–961^ Taken all together with sustained expression of adhesion molecules and chemokines, this pattern is consistent with ongoing endothelial inflammatory activation accompanied by induction of endogenous feedback mechanisms that normally limit NF-κB signaling.

NEC venous endothelial cells also showed increased expression of genes associated with coagulation and inhibition of fibrinolysis (Fig. 24E). *F3* and *SERPINE1*, which were previously reported in the vascular-endothelial response to human NEC, promote tissue-factor-dependent coagulation and inhibit plasminogen-activator-dependent fibrinolysis, respectively.^18,962–963^ Increased *THBS1* can promote latent TGFβ activation and matrix organization, while *F2RL3*/PAR4 provides a receptor for coagulation-protease signaling in endothelial cells.^964^ Together, these changes indicate a procoagulant and antifibrinolytic endothelial response during NEC.

Venous endothelial cells also increased angiogenic and vascular-repair genes seen in arterial endothelium, although inhibitory and organizational changes complicated this response (Fig. 24F). Increased *VEGFA*, *KDR*/VEGFR2, and *FLT4*/VEGFR3 suggested attempted activation of VEGF-responsive growth and survival pathways despite prior reports of reduced intestinal VEGF protein and impaired VEGFR2-dependent microvascular development in NEC.^609,848^ Increased *ENG* and *EGFL7* further supported angiogenic signaling, endothelial sprouting, and vascular integrity, while *NOTCH1* and *NOTCH4* linked this response to angiogenic patterning and VEGFR3 regulation.^965–970^ *KIF26B* and *RASGRP3* added features related to endothelial polarity and VEGF/DAG-responsive vascular signaling.^971–972^ However, reduced *NRP1*, an endothelial VEGF coreceptor and adhesion regulator, suggests that increased ligand and receptor expression may not produce fully coordinated VEGF signaling.^973^ Increased *PLVAP*, which contributes to fenestrated microvascular diaphragms and vascular permeability, further points to altered organization of the specialized venular exchange interface.^898^ Overall, this profile is consistent with attempted vascular repair that remains insufficiently organized to restore normal venous architecture.

Multiple vascular basement membrane and extracellular matrix genes were also increased (Fig. 24G). *HSPG2* can modulate matrix-associated VEGF signaling, while *COL4A1, COL4A2, LAMA5, LAMC1,* and *COL18A1* represent structural components or regulators of the vascular basement membrane and vascular permeability.^974–977^ Matrix reorganization may directly affect immune cell passage because leukocytes preferentially cross permissive regions of the venular basement membrane, and endothelial laminin composition can regulate transendothelial migration.^978–979^ *TGM2* may couple fibronectin deposition to matrix-bound VEGF signaling, while *LTBP1* supports extracellular storage and organization of latent TGFβ complexes.^980–981^ *ADAMTS9* adds an antiangiogenic metalloprotease signal within the same matrix environment.^922^ Thus, NEC alters the extracellular environment at the venular interface where angiogenic signaling, barrier function, and leukocyte extravasation converge.

Consistent with previous reports and the arterial endothelial findings, venous endothelial cells showed a strong interferon-associated transcriptional response (Fig. 24H).^18^ *GBP1* is directly induced by inflammatory cytokines in endothelial cells and marks activated endothelium in inflammatory disease.^910^ *GBP2*, *GBP5*, and *WARS1* broadened the IFNγ-responsive endothelial program, while *IDO1* introduced immunoregulatory tryptophan metabolism.^911,913,982^ *RNF213* linked interferon signaling to altered endothelial inflammatory and angiogenic behavior, whereas *HIF1A* linked this response to hypoxia-responsive transcription.^914,961,983^ This combination may shape immune cell interactions and vascular adaptation at the venular wall through interferon-responsive, immunoregulatory, and hypoxia-associated signals.

NEC venous endothelial cells showed reduced expression of genes involved in calcium and cyclic nucleotide signaling (Fig. 24I). Lower *GNAQ* may impair VEGF-dependent calcium mobilization, endothelial permeability, and angiogenic signaling, while reduced *PLCB4*, *SLC8A1*, and *CAMK2D* point to altered phospholipase-C signaling, calcium homeostasis, and calcium-responsive endothelial regulation.^984–987^ *PRKG1*, a major effector of cGMP-dependent barrier protection, was also reduced, together with *PDE3A*, *PDE4D*, and *PDE5A*, suggesting altered local cAMP- and cGMP-dependent control of permeability, junctional responses, and NOS3 activity.^988–991^

Secretory trafficking, junctional organization, and mechanoadaptation were also reduced. Lower *MYO5A* and *MYO5C* may impair Weibel-Palade body trafficking and calcium-evoked von Willebrand factor externalization.^992–993^ Reduced *LIMA1* and *SORBS2* point to weaker junction-actin coupling, cytoskeletal contractility, and vessel-lumen integrity.^930–931^ Lower *SASH1*, an endothelial TLR4 scaffold, further suggests altered inflammatory sensing alongside barrier and secretory defects.^932^ Overall, inflammatory and adhesive features remain elevated despite reduced molecular support for coordinated calcium signaling, junctional adaptation, secretion, and structural recovery.

Multiple transcriptional regulators of endothelial homeostasis and angiogenesis were reduced in NEC venous endothelial cells (Fig. 24J). *ERG* downregulation may remove a central regulator of endothelial quiescence and NF-κB restraint, including repression of *ICAM1*, whereas *PPARG* downregulation may weaken protection against IL-1β-associated oxidative and endothelial dysfunction.^994–995^ *FOXP1* and *MEF2C* reductions suggest impaired coordination of endothelial migration, proliferation, and VEGF-responsive transcription.^924,996^ At the same time, reductions in the proangiogenic regulators *PBX1* and *HDAC9* occurred together with loss of *ZBTB16* and *MEOX2*, which can restrain angiogenesis or endothelial inflammatory activation.^997–1000^ The combined reductions in these key transcription factors important for homeostasis of the endothelium indicate an erosion of the transcriptional balance that normally couples endothelial quiescence, inflammatory restraint, and organized repair.

Of the 2,732 genes distinguishing fetal from infant control venous endothelial cells, 755 were also dysregulated in NEC versus infant controls, including 334 with fetal-like expression (44.2% of the 755 genes; Fig. 24K,L). Developmentally regulated genes included fetal-high *KDR* and *FLT4*, associated with vascular-growth signaling, and infant-high *SPARCL1*, *AQP1*, and *VWF*, associated with endothelial quiescence, water transport, and hemostatic function.^609,936,938,1001–1002^

The fetal-like genes in NEC highlighted increased mucosal homing and vascular guidance features, along with reduced postnatal barrier and transport functions (Fig. 24L). *MADCAM1* and *PLVAP* were increased and showed fetal-like expression, consistent with greater mucosal homing capacity and fenestrated-barrier specialization.^946,1003^ *FLT4*, *LYVE1*, and *EFNB2* also showed fetal-like expression, adding lymphatic- and arterial-associated transcripts.^1001,1004–1005^ *UNC5B* similarly showed fetal-like expression; its encoded receptor participates in vascular guidance and has also been shown to maintain blood-brain barrier integrity through Wnt/β-catenin signaling in mice.^1006–1007^ By contrast, *PTPRB*, *AQP1*, *LIMA1*, *AKAP12*, *APOLD1*, and *RASIP1*, which normally increased from fetal to infant, were reduced in NEC and showed fetal-like expression, suggesting weaker postnatal junctional stability, transport, cytoskeletal organization, and permeability control.^930,1002,1008–1011^

At the transcription factor level, *KLF4*, *PPARG*, *FOXO3*, and *NFE2L2*/NRF2 were also reduced and showed fetal-like expression, adding weaker inflammatory restraint, barrier maintenance, and antioxidant protection to this pattern (Fig. 24M).^995,1012–1015^ Overall, these findings suggest that NEC venous endothelial cells retain fetal-like vascular-guidance and mucosal homing features while expressing lower levels of genes associated with postnatal barrier, transport, and stress-control functions.

NEC venous endothelium shares inflammatory, interferon-associated, angiogenic, and matrix-producing features with arterial endothelium, but its main pathobiologic role centers on leukocyte entry and barrier control. Initiation may involve enhanced mucosal homing, adhesion, chemokine signaling, and permeability at the blood-tissue interface. During progression, venous procoagulant and antifibrinolytic genes, plasma leakage, leukocyte extravasation, and reduced calcium/junctional regulation could sustain inflammatory tissue injury. Evidence of vascular repair is present, but reduced antioxidant protection, postnatal regulatory control, and barrier-stabilizing signals suggest incomplete recovery. Resolution may require return of venous selectivity, fibrinolytic balance, and endothelial homeostasis.

### Lymphatic Endothelium

Complementing this vascular route of immune cell entry, lymphatic endothelial cells form intestinal lacteals and deeper lymphatic vessels that drain interstitial fluid, carry immune cells and microbial antigens to lymph nodes, and absorb dietary lipids. VEGF-C-dependent maintenance of intestinal lymphatics is required for normal lipid absorption, while continuous lacteal regeneration supports dietary-fat transport (Fig. 25A).^846,1016^ Experimental intestinal studies show that enhancing VEGF-C/VEGFR3-dependent drainage promotes inflammatory cell mobilization and bacterial-antigen clearance, whereas impaired lymphatic function aggravates intestinal edema and inflammation.^1017–1019^ NEC-specific evidence remains limited, but lymphatic ligation in neonatal piglets produced pneumatosis and, in the smallest animals, the complete histopathological spectrum of NEC.^1020^ A prior human NEC single-cell study identified inflammatory activation and enrichment of apoptotic programs in lymphatic endothelial cells, supported by increased TUNEL staining in LYVE1-positive endothelial structures.^18^ Changes in lymphatic growth, scavenging, transport, and vessel maintenance remain incompletely defined and could affect edema, inflammatory-material clearance, and resolution.

**Fig. 25.**
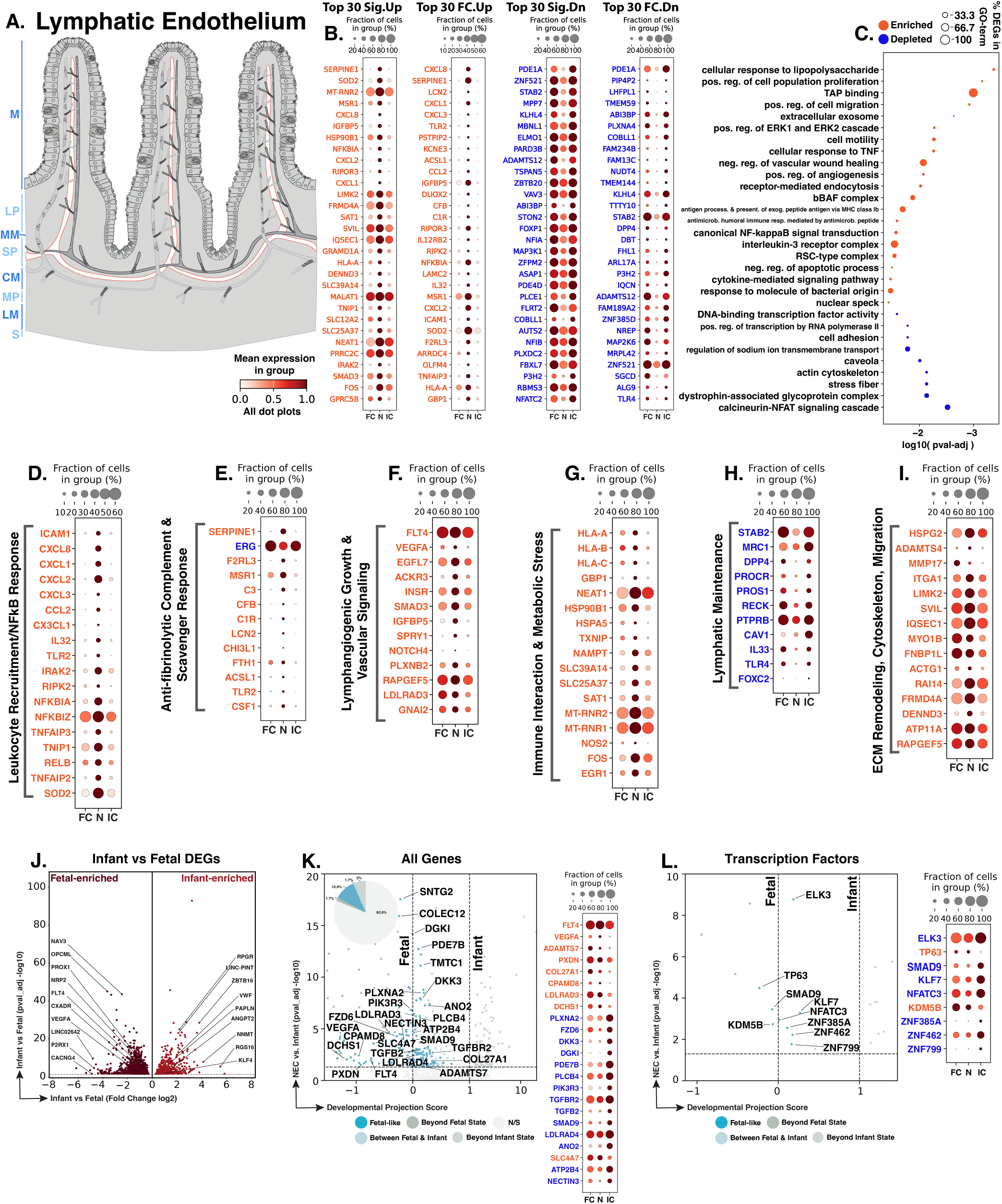
NEC lymphatic endothelial cells show inflammatory activation with altered fibrinolytic, lymphangiogenic, immune-stress, lymphatic-maintenance, matrix-remodeling, and maturity-associated programs. **A.** Schematic showing the lymphatic endothelial distribution (red) within the intestinal wall. Blue text to the left of the Panel A schematic indicates general anatomical regions: M, mucosa; LP, lamina propria; MM, muscularis mucosae; SP, submucosal plexus; CM, circular muscle; MP, myenteric plexus; LM, longitudinal muscle; S, serosa. **B.** Dot plots of top NEC-specific genes ranked by significance and fold change. **C.** GO-term analysis showing enriched and depleted biological processes among NEC-associated DEGs. **D–I.** Curated gene panels showing inflammatory-response, fibrinolysis/scavenger-debris-interface, lymphangiogenic/vascular-remodeling, immune and metabolic-stress, lymphatic-maintenance, and ECM-remodeling/cytoskeletal-migration programs. In panels **B** and **D–I**, orange gene text indicates upregulation in NEC relative to infant controls, whereas blue gene text indicates downregulation. **J.** Volcano plot of DEGs between fetal and infant controls that define the developmental trajectory for lymphatic endothelial cells. **K–L.** Gene-level and transcription-factor of fetal-versus-infant DEGs, highlighting the subset also significantly changed in NEC relative to infant controls and positioned relative to fetal and infant reference states, and their corresponding dotplots. The developmental projection score in panels **K–L** is displayed on a symmetric logarithmic (symlog) scale. For expression dot plots, dot size indicates the fraction of nuclei expressing each gene and dot color indicates mean expression. In maturity plots, vertical dotted lines indicate the fetal and infant reference scores. Colored dots classify NEC-versus-infant DEGs as Fetal-like, Between Fetal & Infant, Beyond Fetal State, or Beyond Infant State; gray N/S dots are fetal-versus-infant DEGs without a significant NEC-versus-infant change.

We identified 900 lymphatic endothelial nuclei across the atlas. Differential-expression analysis detected 707 NEC-versus-infant DEGs, including 214 upregulated and 493 downregulated genes (Fig. 25B). The DEG burden was intermediate between arterial and venous endothelium, but its distribution was distinct: nearly 70% of lymphatic DEGs were downregulated, whereas arterial and venous DEGs were more evenly distributed. As in blood vascular endothelium, GO-term enrichment encompassed LPS- and TNF-responsive inflammation, canonical NF-κB signaling, proliferation, ERK signaling, migration, and vascular growth. Enrichment of these pathways coincided with depletion of adhesion, caveolar, actin-cytoskeletal, stress-fiber, transcriptional, and calcineurin-NFAT functions (Fig. 25C). Lymphatic endothelium shares inflammatory and vascular-growth features with blood endothelium but shows a more pronounced loss of lymphatic structural and regulatory functions.

At the inflammatory interface, increased *ICAM1* may facilitate leukocyte and dendritic-cell transit across activated lymphatic endothelium, while increased *CXCL8*, *CXCL1*, *CXCL2*, and *CXCL3* provide a strong neutrophil-directed chemokine signal (Fig. 25D).^52,901,1021^ *CCL2* and *CX3CL1* broaden this leukocyte-recruiting program through monocyte chemotaxis and adhesive chemokine signaling.^1022–1023^ The *ICAM1*- and chemokine-rich state closely reproduces the lymphatic-endothelial signature previously identified in human NEC.^18^ *TLR2* and *IRAK2* were increased, linking endothelial microbial sensing to IL-1/TLR-associated NF-κB activation, while *RIPK2* introduces a parallel NOD1/2-responsive pathway.^1024–1026^ Concurrent induction of *NFKBIA*, *NFKBIZ*, *TNFAIP3*, *TNIP1*, *RELB*, and *TNFAIP2* indicates that this activation is accompanied by extensive NF-κB-responsive feedback regulation.^1027–1032^ *IL32* may amplify vascular inflammation, whereas *SOD2* suggests compensatory mitochondrial antioxidant activity during inflammatory stress.^960,1033^ These genes position lymphatic endothelium as an active site of microbial sensing and inflammatory cell trafficking.

Inflammatory activation was coupled to altered fibrinolytic, complement, and injury-clearance machinery (Fig. 25E). *SERPINE1*/PAI-1 was increased, suggesting greater antifibrinolytic pressure, while *F2RL3*/PAR4 introduces a candidate thrombin- and protease-responsive pathway.^964,1034^ In lymphatic endothelial cell (LEC)-specific experiments, combined loss of *Erg* and *Fli1* induces inflammatory and prothrombotic transcription, impaired drainage, and severe lymphatic failure.^1035^ The marked reduction of *ERG* in NEC is consistent with loss of a transcriptional regulator required for lymphatic endothelial stability during inflammatory stress. Concurrent increases in *C3*, *CFB*, and *C1R* further identified a complement-associated inflammatory response involving central, alternative, and classical complement components.^1036–1038^ *MSR1*, *LCN2*, *CHI3L1*, and *FTH1* extend this response to debris recognition, antibacterial iron sequestration, endothelial activation and immune recruitment, and antioxidant iron storage.^229,1039–1041^ The resulting profile is compatible with exposure to tissue-derived inflammatory material and acquisition of an antifibrinolytic state that could impede its clearance.

Immune-interaction and cellular-stress genes may further influence lymphatic transport (Fig. 25G). *HLA-A*, *HLA-B*, and *HLA-C* were increased, consistent with greater MHC-I antigen presentation capacity, a function that lymphatic endothelial cells can use to shape CD8 T-cell responses.^1042–1043^ *GBP1* and *NEAT1* connect interferon and TNF signaling to endothelial activation, with *NEAT1* directly promoting TNF-induced *ICAM1*, *VCAM1*, *CXCL8*, and *CCL2* expression in endothelial cells.^910,1044^ *NOS2* is especially relevant to lymphatic function because inflammatory induction of LEC-derived iNOS can inhibit collecting-vessel contraction and drainage.^1045^ *HSPA5*, *TXNIP*, and *NAMPT* connect this state to ER-protein-folding demand, oxidative stress, and NAD-dependent redox adaptation.^1046–1048^ *FOS* and *EGR1* mediate a rapid immediate-early response that can be induced by mechanical stress in endothelial cells.^1049^ Immune communication occurs alongside metabolic and mechanical adaptation; *NOS2* induction offers a direct mechanism by which inflammation could compromise lymphatic pumping.

NEC lymphatic endothelial cells showed increased expression of genes involved in lymphangiogenic growth and migration (Fig. 25F). Higher *FLT4*/VEGFR3 expression supports LEC growth, survival, migration, and lymphangiogenesis, while VEGFA and EGFL7 support lymphatic endothelial growth and endothelial migration.^1050–1052^ *INSR* induction may reinforce VEGFA-ERK sprouting signals, and RAPGEF5-associated Rap signaling can regulate endothelial chemotaxis and cytoskeletal responses.^1053–1054^ IGFBP5, SPRY1, and NOTCH4 introduce antiangiogenic or feedback constraints on endothelial growth, ERK activity, and matrix adhesion.^1055–1057^ This suggests an attempted regenerative response in which growth and migratory activity remain constrained by inflammatory stress and inhibitory feedback.

NEC lymphatic endothelium concurrently lost genes required for stable lymphatic specialization and transport (Fig. 25H). *FOXC2* was reduced; this factor coordinates collecting-vessel maturation, valve formation, mechanotransduction, and adult intestinal lymphatic-barrier function.^1058–1060^ Reduced *FOXC2* expression and reduced representation of calcineurin-NFAT-associated GO terms are consistent with disruption of flow-responsive lymphatic maturation and maintenance. Reductions in *STAB2*, *MRC1*, and *DPP4* point to diminished scavenging, lymphocyte interaction, adhesion, migration, and vessel organization.^1061–1063^ *PTPRB* and *CAV1* were reduced, suggesting altered junctional stabilization and mechanotransduction; lower *PROCR* and *RECK* extended this pattern to protein-C-associated endothelial support and control of matrix proteolysis.^1008,1064–1067^ The opposing *TLR2* and *TLR4* changes indicate altered receptor balance in microbial sensing; TLR4 signaling in LECs can promote inflammation-associated lymphangiogenesis.^1068^ Growth and inflammatory activation coexist with reduced scavenging, mechanosensory, barrier, and transport machinery characteristic of mature lymphatic endothelium.

Extracellular matrix turnover and cytoskeletal regulation were also altered in NEC lymphatic endothelium (Fig. 25I). *HSPG2*/perlecan was increased, consistent with basement membrane organization during lymphatic regeneration, whereas *ADAMTS4* and *MMP17* indicate active extracellular matrix proteolysis.^1069–1071^ Increased *LIMK2* and *RAPGEF5* expressions connect these changes to endothelial actin regulation and chemotactic cytoskeletal responses.^1054,1072^ The simultaneous expression of matrix components, proteases, and cytoskeletal regulators indicates active extracellular matrix turnover and cellular reorganization accompanying the lymphangiogenic response.

Of the 1,514 genes distinguishing fetal from infant control lymphatic endothelial cells, 289 were also dysregulated in NEC versus infant controls, including 163 with fetal-like expression (56.4% of the 289 genes; Fig. 25J,K). Developmentally regulated genes included fetal-high *PROX1*, *FLT4*, and *VEGFA*, associated with lymphatic identity and growth, and infant-high *KLF4* and *VWF*, associated with endothelial homeostatic and hemostatic functions.^938,1012,1050–1051,1073^. The fetal-like genes in NEC highlighted continued lymphangiogenic signaling together with reduced postnatal regulatory features (Fig. 25K). *FLT4* and *VEGFA* were increased and showed fetal-like expression, consistent with persistent or re-engaged lymphatic growth.^1050–1051^ By contrast, *TGFBR2*, *TGFB2*, and *PIK3R3*, which normally increased from fetal to infant, were reduced in NEC and showed fetal-like expression, suggesting weaker postnatal TGF-β- and PI3K-associated lymphatic regulation and repair signaling.^1074–1075^

At the transcription factor level, *ELK3* and *SMAD9* were also reduced and showed fetal-like expression, adding weaker ETS- and BMP-associated regulation to this pattern (Fig. 25L).^1076–1077^ Overall, NEC lymphatic endothelial cells retain fetal-like lymphangiogenic features while expressing lower levels of genes associated with postnatal TGF-β, PI3K, ETS, and BMP regulation. This combination could support lymphatic growth without fully restoring vessel maturation, transport, or drainage.

NEC lymphatic endothelium shares inflammatory NF-κB, chemokine, growth, and matrix activity with blood vascular endothelium but is distinguished by reduced lymphatic specialization, mechanoregulation, scavenging, and transport. During NEC initiation, these alterations may impair the handling of microbial antigens, tissue fluid, and inflammatory material in the injured intestine. Increased leukocyte recruitment and vascular permeability without corresponding lymphatic drainage could promote edema and retention of inflammatory debris, leading to the progression of NEC. Although lymphangiogenic gene induction suggests a compensatory repair response, reduced mature lymphatic programs may limit restoration of transport and clearance, potentially prolonging intestinal inflammation despite ongoing lymphangiogenic signaling.

### Proliferative Vascular Endothelium

Proliferative vascular endothelial cells represent a cycling state superimposed on arterial, venous, or other blood-vascular identities. Their expression of arterial-patterning, angiogenic, and permeability-associated genes is consistent with contributions from multiple vascular compartments (Fig. 26A). Endothelial proliferation is required for postnatal intestinal microvascular growth and depends on VEGFA-VEGFR2 signaling. Experimental NEC reduces endothelial proliferation and microvascular density, while human NEC tissue contains fewer proliferating CD31⁺Ki67⁺ villous endothelial cells.^609–610^ Previous single-cell analysis nevertheless found an increased proportion of *MKI67*-expressing vascular and lymphatic endothelial cells in NEC alongside endothelial apoptotic programs.^18^ Because endothelial expansion is one component of vascular repair, we examined cell-cycle programs alongside genes involved in inflammation, junctional organization, and vascular regulation. These comparisons characterize the molecular state accompanying endothelial cycling in NEC and identify which responses coexist within the proliferative endothelial population.

**Fig. 26.**
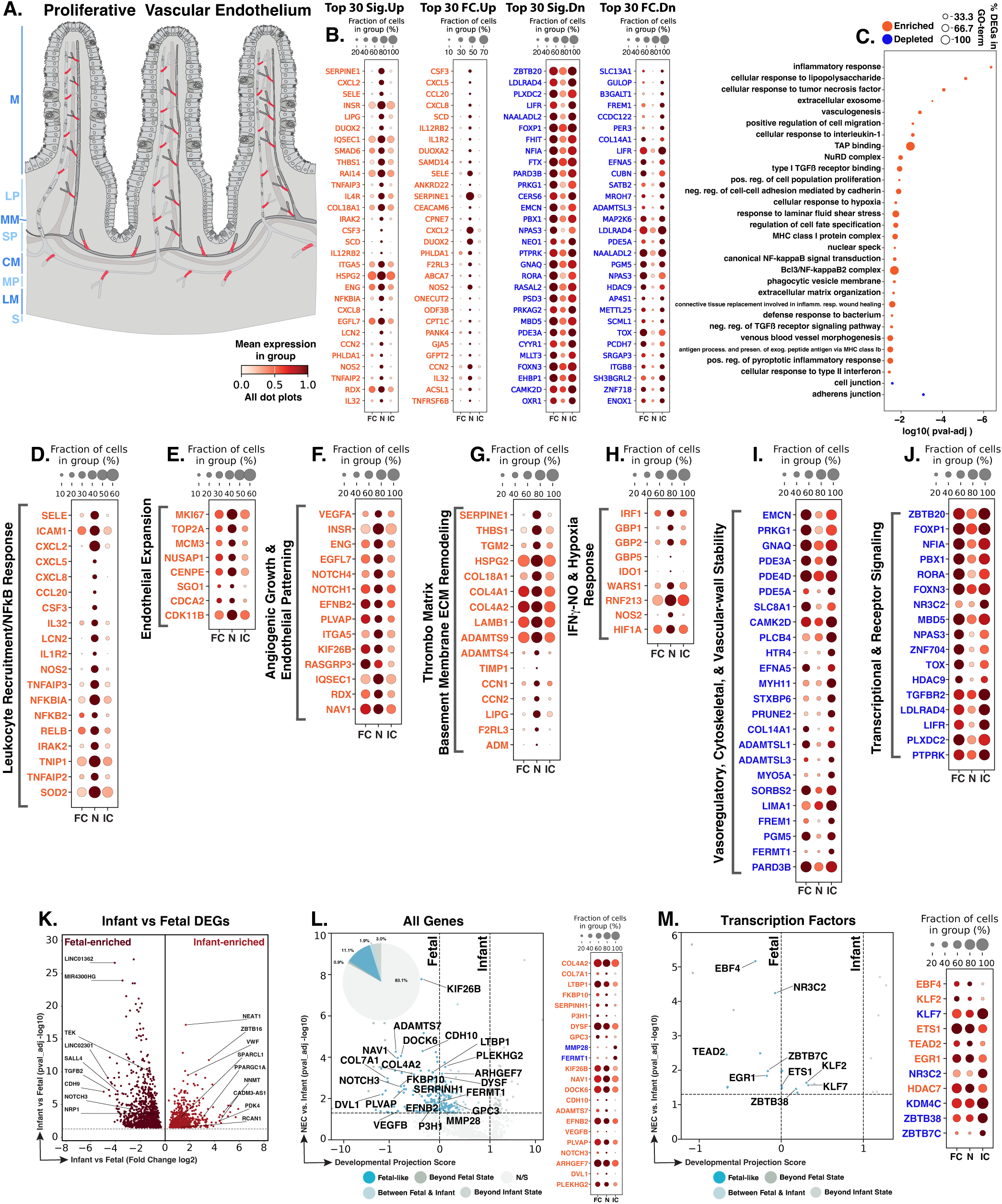
NEC proliferative vascular endothelial cells show inflammatory activation and endothelial expansion with angiogenic, matrix, immunometabolic, vascular-stability, and maturity-associated changes. **A.** Schematic showing the distribution of proliferative vascular endothelial cells (red) within the intestinal vasculature. Blue text to the left of the Panel A schematic indicates general anatomical regions: M, mucosa; LP, lamina propria; MM, muscularis mucosae; SP, submucosal plexus; CM, circular muscle; MP, myenteric plexus; LM, longitudinal muscle; S, serosa. **B.** Dot plots of top NEC-specific genes ranked by significance and fold change. **C.** GO-term analysis showing enriched and depleted biological processes among NEC-associated DEGs. **D–J.** Curated gene panels showing inflammatory endothelial activation, endothelial expansion, angiogenic/endothelial remodeling, thrombo-matrix/basement-membrane remodeling, IFNγ–NO immunometabolic stress, vasoregulatory/cytoskeletal/vascular-wall stability, and transcriptional and receptor regulators. In panels **B** and **D–J**, orange gene text indicates upregulation in NEC relative to infant controls, whereas blue gene text indicates downregulation. **K.** Volcano plot of DEGs between fetal and infant controls that define the developmental trajectory for proliferative vascular endothelial cells. **L–M.** Gene-level and transcription-factor maturity analyses of fetal-versus-infant DEGs, highlighting the subset also significantly changed in NEC relative to infant controls and positioned relative to fetal and infant reference states, and their corresponding dotplots. The developmental projection score in panels **L–M** is displayed on a symmetric logarithmic (symlog) scale. For expression dot plots, dot size indicates the fraction of nuclei expressing each gene and dot color indicates mean expression. In maturity plots, vertical dotted lines indicate the fetal and infant reference scores. Colored dots classify NEC-versus-infant DEGs as Fetal-like, Between Fetal & Infant, Beyond Fetal State, or Beyond Infant State; gray N/S dots are fetal-versus-infant DEGs without a significant NEC-versus-infant change.

We identified 1,049 proliferative vascular endothelial nuclei across the atlas. Differential-expression analysis identified 693 genes significantly changed in NEC compared to infant controls, including 363 upregulated and 330 downregulated genes (Fig. 26B). GO-term enrichment connected endothelial proliferation with LPS-, TNF-, and IL-1-responsive inflammation, canonical NF-κB and interferon signaling, hypoxia, vascular growth and migration, extracellular matrix organization, and responses to laminar shear stress. These increases were accompanied by depletion of cell junction and adherens junction programs (Fig. 26C). Thus, NEC proliferative endothelial cells combine inflammatory and remodeling-associated gene induction with depletion of junction-associated transcriptional programs.

The inflammatory response was categorized by leukocyte recruitment with extensive regulation of cytokine signaling (Fig. 26D). Higher *SELE* and *ICAM1* support leukocyte rolling, adhesion, and transendothelial migration across activated endothelium.^1078–1079^ Upregulation of *CXCL2*, *CXCL5*, and *CXCL8* suggests a strong neutrophil-directed chemokine signal, while *CCL20* extends this response toward mucosal lymphocyte recruitment and *CSF3* provides granulopoietic and neutrophil-supportive activity.^231,901,1080–1082^ Higher *IL32* may further support vascular inflammatory signaling, whereas *IRAK2* links IL-1 signaling to endothelial NF-κB activation.^1025,1033^ *NFKB2*, *RELB*, and *TNFAIP2* further support TNF-, IL-1-, and LPS-responsive inflammatory signaling.^954,1032^ At the same time, higher *NFKBIA*, *TNFAIP3*, *TNIP1*, and *IL1R2* indicate negative-feedback and decoy-receptor mechanisms that may restrain NF-κB and IL-1 signaling.^1029–1030,1083–1084^ Increased LCN2 expression, which can be induced through NF-κB activation, may contribute to antimicrobial defense through iron sequestration and may also support endothelial barrier integrity by restoring membrane localization of junctional proteins under inflammatory conditions.^229,1085^ Higher *SOD2* adds a mitochondrial antioxidant response to TNF-associated oxidative stress.^960^ Overall, these findings suggest that cycling endothelial cells share leukocyte-recruiting and NF-κB-responsive features observed in arterial and venous endothelium while also expressing regulatory mechanisms that may limit inflammatory signaling.

Within that inflammatory state, a coherent mitotic program confirms active endothelial expansion (Fig. 26E). Increased *MKI67* marks cycling cells, while *MCM3* and *TOP2A* support DNA replication and mitotic chromosome organization.^180,1086–1087^ Increased *NUSAP1* and *CENPE* connect spindle organization with chromosome congression, whereas *SGO1* protects centromeric cohesion during mitosis.^1088–1090^ *CDCA2* coordinates mitotic chromatin regulation, and *CDK11B* contributes to centrosome maturation and bipolar-spindle formation.^1091–1092^ These genes suggest that the cells are actively dividing. However, because this proliferation occurs alongside inflammatory, hypoxic, interferon, and thrombotic/matrix-associated responses, NEC-associated cycling may reflect a vascular growth state that differs from uncomplicated developmental proliferation.

The angiogenic response combined endothelial growth with migration, vascular patterning, and permeability-associated features (Fig. 26F). Higher *VEGFA*, *ENG*, and *EGFL7* were consistent with angiogenic growth and endothelial migration, while increased *ITGA5* and *KIF26B* added features associated with vascular organization, polarity, and directional migration.^965,971,1052,1093–1094^ *RASGRP3* and *RDX* further linked this response to angiogenic signaling, permeability, and actin-membrane organization.^972,1095^ Higher *NOTCH1* and *NOTCH4* were consistent with regulation of vascular-network organization and endothelial sprouting.^1096–1097^ *EFNB2* added an arterial-patterning feature, while increased *PLVAP* suggested fenestral-diaphragm and permeability-associated specialization.^898,1098–1100^ Overall, these changes suggest active but heterogeneous vascular reconstruction involving growth, migration, patterning, and reorganization of the endothelial exchange interface.

The extracellular matrix also showed broad changes (Fig. 26G). Higher *SERPINE1*/PAI-1, *THBS1*, and *TGM2* were consistent with increased antifibrinolytic activity, latent TGF-β activation, and matrix crosslinking.^916,1034,1101^ Increased *HSPG2*, *COL18A1*, *COL4A1*, *COL4A2*, and *LAMB1* suggested a vascular basement membrane response with potential effects on growth factor availability, sprouting, permeability, and vessel stabilization.^974,977,1102–1104^ Higher *ADAMTS9*, *ADAMTS4*, and *TIMP1* were consistent with signals that can restrain angiogenesis or endothelial migration, whereas *CCN1* and *CCN2* added features associated with angiogenic activity, pericyte adhesion, and basement membrane formation.^922,1070,1105–1107^ Increased *LIPG*, *F2RL3*, and *ADM* further suggested changes in lipid processing, thrombin-responsive signaling, and vascular-tone regulation.^964,1108–1109^ Together, these findings suggest vascular matrix rebuilding accompanied by both proangiogenic and angiogenesis-limiting signals.

Interferon, inflammatory, and hypoxia-associated genes further distinguished cycling endothelial cells from stable endothelium (Fig. 26H). Higher *IRF1*, *GBP1*, *GBP2*, and *GBP5* were consistent with an IFNγ-responsive endothelial state, while increased *IDO1* and *WARS1* added immunometabolic and translational features.^910–911,913,1110–1112^ *RNF213* linked this response to inflammatory and angiogenic regulation, while *NOS2* suggested cytokine-responsive nitric oxide signaling.^914,1113–1114^ Higher *HIF1A* further linked this state to hypoxia-associated transcription.^241,904^ This pattern suggests that endothelial proliferation in NEC occurs within an interferon-rich, inflammatory, and hypoxic environment rather than a fully resolved regenerative state.

Genes involved in vascular tone, permeability, calcium signaling, and junctional organization were reduced at the same time (Fig. 26I). Lower *EMCN* suggested weaker VEGFR2-dependent migration and proliferation, while reduced *PRKG1*, *PDE3A*, *PDE4D*, and *PDE5A* were consistent with altered cAMP- and cGMP-dependent regulation of permeability, junctions, and NOS3 signaling.^989–991,1115–1116^ Lower *GNAQ*, *SLC8A1*, and *CAMK2D* further suggested weaker VEGF-, calcium-, and mechanosensitive signaling, whereas reduced *MYO5A* and *LIMA1* pointed to lower support for Weibel-Palade-body trafficking and junction-actin coupling.^930,984,992,1117–1119^ Lower *HTR4* and *EFNA5* indicated potential reductions in angiogenic and ephrin-associated signaling.^1120–1121^ These changes suggest that mitotic activity persists despite reduced expression of genes needed to coordinate endothelial signaling, secretion, and junctional maturation.

Additional transcriptional and receptor-signaling changes reinforced this separation between proliferation and maturation (Fig. 26J). Reduced *FOXP1*, *PBX1*, and *HDAC9* were consistent with altered regulation of endothelial proliferation, migration, sprouting, and angiogenesis.^924,997–998^ Concurrent reduction of *TGFBR2* and the TGF-β inhibitor *LDLRAD4* suggested reorganization of TGF-β signaling rather than simple pathway suppression.^1122–1123^ Lower *LIFR* and *PTPRK* may further affect endothelial survival, growth, adhesion, tubule formation, and migration.^1124–1125^ Together, these changes suggest that NEC-associated endothelial proliferation occurs alongside altered regulatory signals that normally coordinate angiogenesis with endothelial maturation.

Of the 1,673 genes distinguishing fetal from infant control proliferative endothelial cells, 282 were also dysregulated in NEC versus infant controls, including 185 with fetal-like expression (65.6% of the 282 genes; Fig. 26K,L). Developmentally regulated genes included fetal-high *NOTCH3* and *NRP1*, associated with vascular maturation and VEGF-dependent angiogenesis, and infant-high *SPARCL1* and *VWF*, associated with endothelial quiescence and hemostatic function.^936,938,1126–1127^ The fetal-like genes in NEC highlighted basement membrane production, collagen maturation, cytoskeletal organization, and vascular patterning (Fig. 26L). *COL4A2*, *COL7A1*, and *LTBP1*, together with the collagen-processing genes *FKBP10*, *SERPINH1*, and *P3H1*, were increased in NEC and showed fetal-like expression.^1103,1128–1132^ These changes suggest reuse of developmental features associated with basement membrane formation and collagen processing during NEC vascular repair.

Additional fetal-like genes extended this pattern to endothelial polarity, cytoskeletal regulation, and vascular specialization. *KIF26B*, *DOCK6*, *PLEKHG2*, *ARHGEF7*, and *DVL1* were increased in NEC and showed fetal-like expression, consistent with changes in endothelial polarity, Rho-family GTPase signaling, cytoskeletal organization, and junctional regulation.^971,1133–1136^ Increased *EFNB2*, *NOTCH3*, and *PLVAP* added arterial-patterning, vessel-maturation, and fenestral-exchange features.^898,1098,1126^ At the transcription factor level, *KLF2* also showed fetal-like expression and may provide counter-regulatory support for endothelial quiescence, nitric-oxide production, and antithrombotic function within the cycling population (Fig. 26M).^1137–1138^ *ETS1*, *TEAD2*, *EGR1*, and *HDAC7* added regulatory features associated with endothelial migration, proliferation, injury responses, and growth control.^1139–1145^ Overall, the developmental comparison suggests that NEC proliferative endothelial cells retain fetal-like features associated with matrix production, polarity, permeability, and vascular patterning, while most developmentally regulated genes remain infant-like.

NEC proliferative vascular endothelium represents active mitotic repair within an unresolved vascular-injury environment. Initiation may involve compensatory endothelial cycling in response to hypoxia, inflammatory injury, and impaired vascular integrity. During progression, cell-cycle activity remains embedded within leukocyte recruitment, interferon, hypoxia, angiogenic patterning, antifibrinolytic, and basement membrane signals, suggesting repair occurs in a still-inflamed vascular bed. Reduced junctional organization, mechanosensory and cyclic nucleotide signaling, endothelial secretion, and regulatory integration may prevent new endothelial cells from forming a stable perfused network. Resolution may depend on whether proliferative vascular endothelium can transition from inflammatory expansion to mature microvascular reconstruction.

### Immune compartment

The intestinal immune compartment, marked by *PTPRC*/CD45 (Fig. 2), contains resident and recruited innate cells together with adaptive lymphocytes that maintain tolerance to commensal and dietary antigens while responding to microbial invasion and tissue injury. Macrophages survey the lamina propria, phagocytose microbes and dying cells, and coordinate inflammation and repair; T cells provide regulatory, helper, and cytotoxic functions; B cells generate mucosal antibody responses; and dendritic cells sample antigens and direct T-cell priming.^1146–1149^ Human NEC studies have documented recruitment of inflammatory CD16+CD163+ monocytes/macrophages, increased inflammatory and cytotoxic T-cell activity with TCRβ clonal expansion, and depletion of circulating B cells with relative plasmablast enrichment, although total mucosal B-cell abundance was unchanged.^18,33,1150^

To define the immune populations in our study, marker-based annotation resolved four immune populations: *BLK*/*PAX5*/*MS4A1*-expressing B cells, *CD3E*/*CD3G*/*TRBC1*-expressing T cells, *CSF1R*/*CD163*/*MRC1*-expressing macrophages, and a *CLEC9A*/*BATF3*/*XCR1*-enriched dendritic-cell population. (Fig. 27;^18,49^) Dendritic cells were retained in the atlas-level immune cell map, but the limited number of nuclei and absence of significant NEC-versus-infant DEGs prevented detailed disease interpretation. The subsequent sections therefore focus on macrophages, T cells, and B cells.

**Fig. 27.**
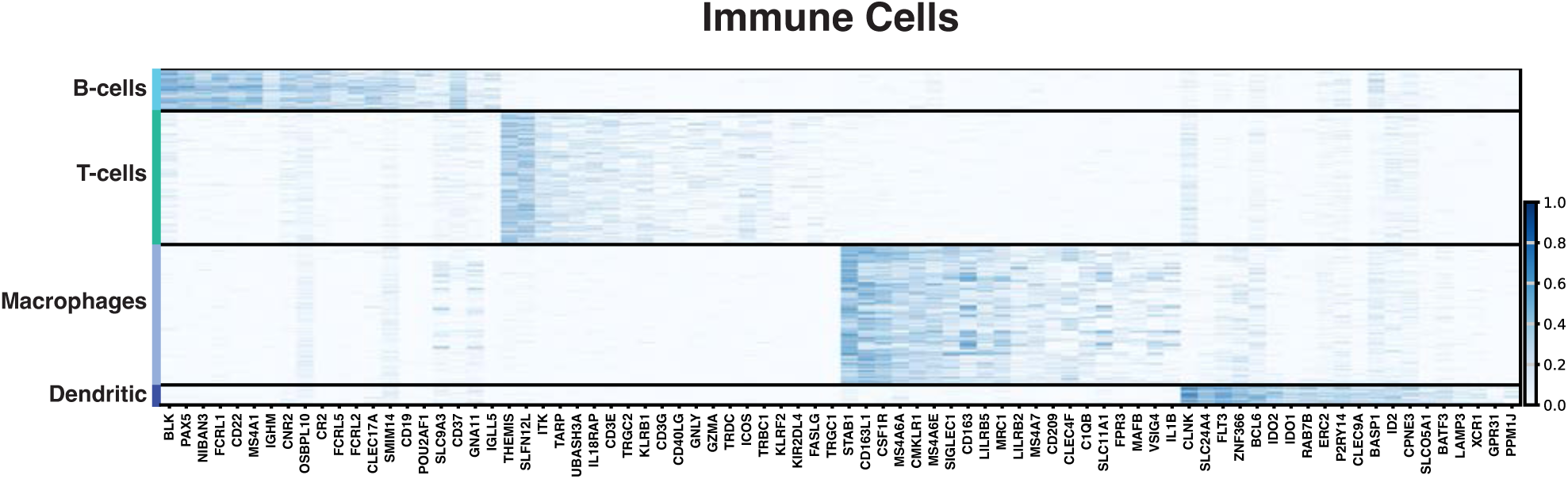
Immune cell-type signatures define B-cell, T-cell, macrophage, and dendritic-cell populations. Heatmap showing marker-gene expression across B cells, T cells, macrophages, and dendritic cells. Distinct transcriptional signatures support immune cell-type assignments and provide the framework for the detailed immune-cell analyses that follow.

The following sections examine how NEC alters innate and adaptive immune populations in the intestinal wall. Macrophages drive the dominant immune response, in which inflammatory amplification, tissue clearance, antigen presentation, angiogenic support, and repair capacity are most strongly altered. T cells capture a smaller but biologically coherent adaptive response involving stress adaptation, proliferation, tissue interaction, inflammatory effector activity, and regulatory control. B cells provide a contrasting response, with limited transcriptional activation and reduced antigen-receptor, cytokine-sensing, and differentiation-associated signals. Together, these sections frame NEC immune activity as intense but incompletely coordinated, with macrophage-driven inflammation occurring alongside reduced clearance, antigen presentation, and adaptive immune signal integration with a concurrent disruption of receptor-coupled signaling in T and B cells, which may further weaken coordination between innate and adaptive immunity.

### Macrophages

Intestinal macrophages are innate immune phagocytes that form a dense surveillance network throughout the intestinal wall, especially within the lamina propria beneath the epithelium. This population includes long-lived tissue-resident macrophages and cells continually replenished from recruited circulating monocytes.^1146,1151–1152^ Under homeostatic conditions, intestinal macrophages monitor the local environment, phagocytose microbes, present antigens, and clear apoptotic cells while limiting unnecessary inflammation. During infection or injury, activated macrophages and recruited monocytes coordinate leukocyte recruitment and help contain the insult. Efferocytosis can limit secondary tissue damage while supporting inflammatory resolution, epithelial regeneration, angiogenesis, and microvascular repair (Fig. 28A).^1152–1154^ Prior human NEC single-cell studies identified inflammatory macrophage states and changes in antigen presentation, developmental features, and pyroptosis, but did not define how these relate to core tissue-support functions.^18,34–35,1155^ Macrophages can also provide protective functions; in experimental NEC, macrophage-derived IGF-1 supports intestinal VEGF expression, endothelial proliferation, and postnatal microvascular development.^610^ Human NEC studies also identified recruited inflammatory CD16⁺CD163⁺ monocytes/macrophages and states enriched for *IL1A*, *IL1B*, *IL6*, *CXCL8*, *CXCL2*, *CCL20*, and NF-κB-associated genes.^18,33^ How this inflammatory activation might relate to apoptotic-cell clearance, antigen presentation, and trophic support in human NEC remains unresolved.

**Fig. 28.**
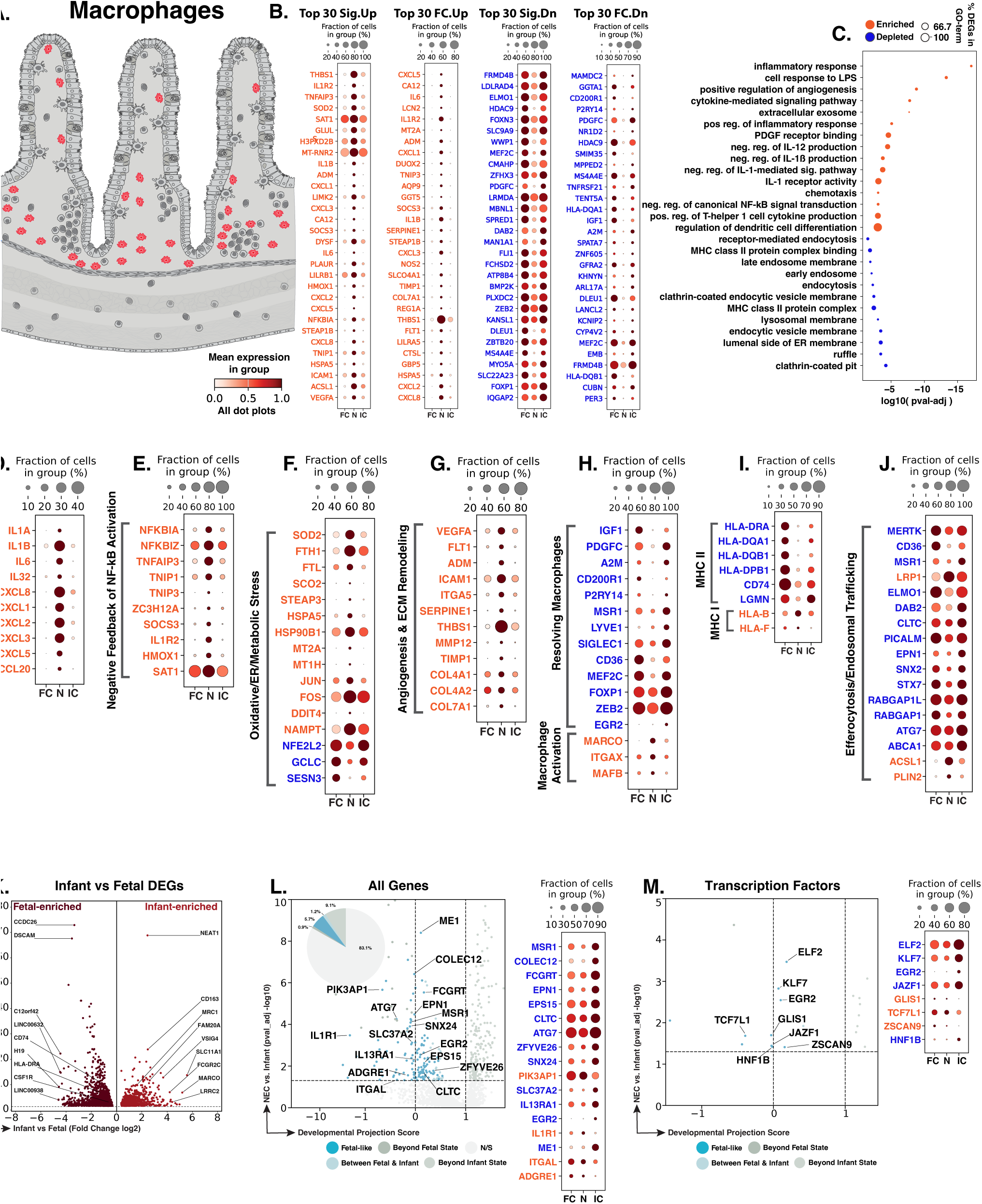
NEC macrophages show inflammatory and NF-κB activation with oxidative/metabolic stress, angiogenic/tissue remodeling, macrophage-state, antigen-presentation, trafficking, and maturity-associated changes. **A.** Schematic showing macrophage distribution (red) throughout the intestinal wall. Blue text to the left of the Panel A schematic indicates general anatomical regions: M, mucosa; LP, lamina propria; MM, muscularis mucosae; SP, submucosal plexus; CM, circular muscle; MP, myenteric plexus; LM, longitudinal muscle; S, serosa. **B.** Dot plots of top NEC-specific genes ranked by significance and fold change. **C.** GO-term analysis showing enriched and depleted biological processes among NEC-associated DEGs. **D–J.** Curated gene panels showing inflammatory-response, NF-κB activation, oxidative/ER/metabolic-stress, angiogenesis/tissue-remodeling, resolving-versus-inflammatory macrophage programs, MHC-I/MHC-II antigen-presentation, and efferocytosis/endosomal-trafficking programs. In panels **B** and **D–J**, orange gene text indicates upregulation in NEC relative to infant controls, whereas blue gene text indicates downregulation. **K.** Volcano plot of DEGs between fetal and infant controls that define the developmental trajectory for macrophages. **L–M.** Gene-level and transcription-factor maturity analyses of fetal-versus-infant DEGs, highlighting the subset also significantly changed in NEC relative to infant controls and positioned relative to fetal and infant reference states, and their corresponding dotplots. The developmental projection score in panels **L–M** is displayed on a symmetric logarithmic (symlog) scale. For expression dot plots, dot size indicates the fraction of nuclei expressing each gene and dot color indicates mean expression. In maturity plots, vertical dotted lines indicate the fetal and infant reference scores. Colored dots classify NEC-versus-infant DEGs as Fetal-like, Between Fetal & Infant, Beyond Fetal State, or Beyond Infant State; gray N/S dots are fetal-versus-infant DEGs without a significant NEC-versus-infant change.

We identified 1,398 macrophage nuclei across the atlas, making macrophages the largest sampled immune population. Differential-expression analysis identified 860 genes significantly altered in NEC macrophages compared to infant controls, including 310 upregulated and 550 downregulated genes: the largest NEC-associated DEG burden within the immune compartment (Fig. 28B). GO-term enrichment connected the upregulated response to inflammation, LPS and cytokine signaling, chemotaxis and angiogenesis, whereas depleted programs converged on MHC-II biology, receptor-mediated endocytosis, endosomal and lysosomal membranes, and clathrin-coated vesicles (Fig. 28C).

Cytokine and chemokine induction dominated the upregulated genes. Higher *IL1A*, *IL1B*, *IL6*, *IL32*, *CXCL8*, *CXCL1*, *CXCL2*, *CXCL3*, *CXCL5*, and *CCL20* defined a macrophage state consistent with amplified local inflammation and leukocyte recruitment (Fig. 28D). This closely recapitulates prior human NEC findings in which *IL1A*, *IL1B*, *IL6*, *CXCL8*, *CXCL2*, and *CCL20* were increased in inflammatory macrophages.^18^ Together with reported accumulation of CD16⁺CD163⁺ inflammatory monocytes/macrophages in NEC tissue, these findings support inflammatory myeloid recruitment and macrophage activation.^33^ This response was accompanied by NF-κB-linked feedback. Increased *NFKBIZ* supported secondary Toll-like-receptor- and IL-1-responsive transcription, while higher *NFKBIA*, *TNFAIP3*, *TNIP1*, *TNIP3*, *ZC3H12A*, *SOCS3*, and *IL1R2* reflected compensatory restraints on NF-κB, Toll-like-receptor, cytokine, and IL-1 signaling (Fig. 28E).^1084,1156–1162^ Increased *HMOX1* further suggested heme and oxidative stress, while higher *SAT1* has been associated with ferroptosis and inflammatory responses in macrophages.^1163^ Macrophage cytokine induction thus occurs alongside NF-κB signaling response and oxidative-stress-associated genes, suggesting an inflammatory state under simultaneous regulatory and metabolic pressure.

The NEC macrophage transcriptome also showed features of oxidative, iron, proteostatic, and metabolic adaptation. Higher *SOD2* and the ferritin genes *FTH1* and *FTL* were consistent with mitochondrial antioxidant defense and iron sequestration, while *STEAP3* links iron handling to macrophage inflammatory signaling (Fig. 28F).^1164–1166^ Increased *SCO2*, *HSPA5*, and *HSP90B1* pointed to mitochondrial respiratory support, ER protein folding, and Toll-like-receptor quality control.^1167–1169^ Higher *MT2A* and *MT1H* suggested metal and oxidative-stress adaptation, with *JUN* and *FOS* adding immediate-early inflammatory signaling.^1170–1171^ Increased *DDIT4* and *NAMPT* were consistent with mTOR restraint, NAD metabolism, and macrophage survival in metabolically demanding environments.^1172–1173^ This adaptation appeared incomplete because lower *NFE2L2* and *GCLC* suggested weaker antioxidant support. Lower *SESN3* indicated altered immune regulation; in a mouse colitis model, *SESN3* promoted macrophage inflammatory cytokine production and generation of Th1 and Th17 cells.^1174–1176^ Thus, NEC macrophages appear to increase stress-adaptive responses while expressing lower levels of some genes needed for baseline antioxidant protection.

Angiogenesis- and matrix-associated genes were increased in NEC macrophages. Higher *VEGFA* and *FLT1* indicated altered VEGF signaling with potentially opposing effects: *VEGFA* promotes angiogenesis, whereas macrophage *FLT1* can restrain *VEGFA*-driven vessel growth. Increased *ITGA5* could couple matrix sensing to VEGFA production and angiogenic macrophage behavior (Fig. 28G).^1177–1179^ Increased *ADM* added a vasoactive and inflammation-responsive signal; higher *ICAM1* expression raised the adhesive potential of activated macrophages.^1180^ Angiogenic activation coincided with altered matrix regulation. Higher *SERPINE1* and *THBS1* expression implicated fibrinolytic and matricellular control; *MMP12* and *TIMP1* combined protease activity with restraint; and *COL4A1*, *COL4A2*, and *COL7A1* marked collagen deposition.^916,1181–1185^ These genes are consistent with ongoing vascular and matrix injury accompanied by angiogenic, adhesive, proteolytic, and collagen-deposition activity.

Conversely, trophic and tissue-adapted macrophage functions were broadly reduced. Lower *IGF1* was particularly notable because macrophage-derived IGF-1 supports neonatal intestinal microvascular development and protection from NEC (Fig. 28H).^610^ Reduced *PDGFC* may indicate weaker macrophage-fibroblast communication, as macrophage-derived PDGF-C can stimulate PDGFRα-expressing fibroblasts and, in other injury settings, promote fibroblast VEGF-C production and lymphatic repair.^1186–1187^ Lower *A2M* may likewise reduce extracellular protease control and A2M-associated antimicrobial and pro-resolving functions.^1188–1189^ Reduced *CD200R1*, *P2RY14*, *MSR1*, *LYVE1*, *SIGLEC1*, and *CD36* pointed to weaker immune regulation, recruitment sensing, scavenging, lipid handling, and tissue-associated macrophage identity.^1152,1190–1194^ Lower *ZEB2* and *EGR2* were also consistent with reduced tissue-specific identity and repair-associated macrophage differentiation.^1195–1196^ *FOXP1* downregulation can facilitate monocyte-to-macrophage differentiation, whereas MEF2C can sustain inflammatory macrophage activity, suggesting that some differentiation-associated features remain intact.^1197–1198^ Increased *MAFB* and *MARCO* may reflect a limited compensatory response that preserves macrophage differentiation, efferocytic capacity, and phagocytic clearance despite broader loss of trophic and tissue-adapted features.^1199–1201^ Overall, resident and trophic features were reduced as inflammatory and injury-adapted macrophage responses increased.

MHC-II antigen presentation genes were broadly reduced. Reduced *HLA-DRA*, *HLA-DQA1*, *HLA-DQB1*, *HLA-DPB1*, *CD74*, and *LGMN* indicated impaired MHC-II antigen processing and presentation to T cells (Fig. 28I).^1202–1203^ This agrees with previous human NEC studies reporting reduced macrophage MHC-II programs, diminished MHC-II-CD4 T-cell interactions, and expansion of HLA-DR-low inflammatory myeloid states.^18,1155^ By contrast, *HLA-B* and *HLA-F* increased; HLA-F can engage receptors expressed by natural killer (NK) cells and CD8+ T cells.^1204^ Opposing MHC-II and MHC-I expression may shift macrophage-lymphocyte communication away from MHC-II-dependent CD4⁺ T-cell interactions while preserving greater potential for HLA-I-mediated interactions with cytotoxic CD8⁺ T cells and NK cells.^1205–1206^

Reduced clearance-related machinery suggested weaker handling of injured tissue. *MERTK*, *CD36*, and *ELMO1*, which contribute to apoptotic cell recognition and engulfment, were reduced in NEC macrophages (Fig. 28J).^1194,1207–1208^ In contrast, *LRP1* was increased. Because macrophage LRP1 supports survival and apoptotic-cell clearance, its elevation may represent a compensatory response to ongoing injury.^1209–1210^ Lower *MSR1* added a scavenger-receptor deficit, while reduced *DAB2*, *CLTC*, *PICALM*, *EPN1*, *SNX2*, and *STX7* pointed to changes in clathrin-mediated uptake, endosomal sorting, and vesicle maturation.^1211–1214^ *RABGAP1* and *RABGAP1L* provide additional candidate links to Rab-dependent vesicle regulation, although their roles in intestinal macrophages are less established. Reduced *ATG7* and *ABCA1* suggested weaker autophagy-related maturation and cholesterol export, whereas increased *ACSL1* and *PLIN2* were consistent with inflammatory fatty acid activation and lipid-droplet storage.^1215–1218^ These changes are consistent with reduced apoptotic cell uptake and intracellular cargo handling, accompanied by greater inflammatory lipid metabolism and storage.

Of the 2,174 genes distinguishing fetal from infant control macrophages, 368 were also dysregulated in NEC versus infant controls, including 124 with fetal-like expression (33.7% of the 368 genes; Fig. 28K,L). Developmentally regulated genes included *MRC1* and *CSF1R*, along with the scavenging and antimicrobial genes *CD163*, *SLC11A1*, and *MARCO*, consistent with postnatal macrophage differentiation and host-defense functions.^49,1219–1222^ Among the fetal-like genes, reduced *MSR1* and *COLEC12* pointed to weaker scavenger and tissue-associated functions, while lower *FCGRT*, *EPN1*, *EPS15*, and *CLTC* involved immune-complex and endocytic trafficking.^1152,1212,1214,1223–1224^ Reduced *ATG7*, *SLC37A2*, *IL13RA1*, and *EGR2* extended this pattern to autophagy, metabolic control of inflammation, IL-13-responsive repair, and tissue-adapted differentiation.^1196,1215,1225–1226^ Conversely, fetal-high *PIK3AP1* and *IL1R1* also showed fetal-like expression, suggesting altered innate-signaling sensitivity; *PIK3AP1* links Toll-like receptors to PI3K-dependent inflammatory regulation.^1227^ The transcription factors *KLF7* and *JAZF1* further implicated regulation of NF-κB activity and inflammatory restraint (Fig. 28M).^1228–1229^ Thus, most developmental genes remained infant-like, but a subset showed fetal-like expression involving scavenging, endocytic trafficking, autophagy, metabolic control, and repair-associated macrophage functions.

NEC macrophages show the largest immune transcriptional response and combine strong inflammatory activity with reduced tissue-supportive functions. Initiation may involve IL-1, chemokine, NF-κB, oxidative/iron-stress, and metabolic-stress features that amplify local inflammation and recruit additional immune cells. During progression, macrophages may sustain tissue injury through inflammatory feedback, matrix remodeling, and altered angiogenic signaling while losing resident identity, macrophage-derived IGF1 support, MHC-II antigen presentation, efferocytosis, scavenging, and endosomal processing. This creates a state that is inflammatory and repair-associated but poorly equipped for clearance and coordination. Resolution may fail if macrophages cannot remove damaged cells, process antigen appropriately, and support epithelial and vascular recovery.

### T cells

Intestinal T cells form distinct immune populations within the epithelium and lamina propria. Intraepithelial T cells provide rapid surveillance at the epithelial surface, while lamina propria CD4⁺ and CD8⁺ effector T cells coordinate antigen-specific defense and FOXP3⁺ regulatory T cells limit excessive inflammation and maintain mucosal tolerance.^1230–1231^ During infection or tissue injury, T-cell recruitment, expansion, and effector differentiation strengthen host defense, while regulatory populations limit collateral tissue damage. This balance is important during neonatal intestinal injury: adoptive transfer of FOXP3⁺ regulatory T cells reduced antigen-presenting-cell and T-cell activation and attenuated experimental NEC (Fig. 29A).^31^ Human NEC tissue shows disruption of this balance, with reduced ratios of mucosal FOXP3⁺ regulatory T cells to CD4⁺ and CD8⁺ effector T cells.^1232^ Experimental NEC also links epithelial TLR4-CCL25 signaling to CD4⁺ T-cell recruitment and STAT3-dependent polarization toward a pathogenic Th17 phenotype.^32^ Human single-cell studies further identified increased TCRβ clonal expansion and inflammatory activation, reduced regulatory T-cell suppressive features, and enrichment of mTOR-, TNF-, and MYC-associated activity.^18,34^ These studies establish strong inflammatory and proliferative T-cell responses in NEC, but whether T cells retain coordinated TCR signaling, antigen recognition, tissue adaptation, and immune regulation remains unresolved.

**Fig. 29.**
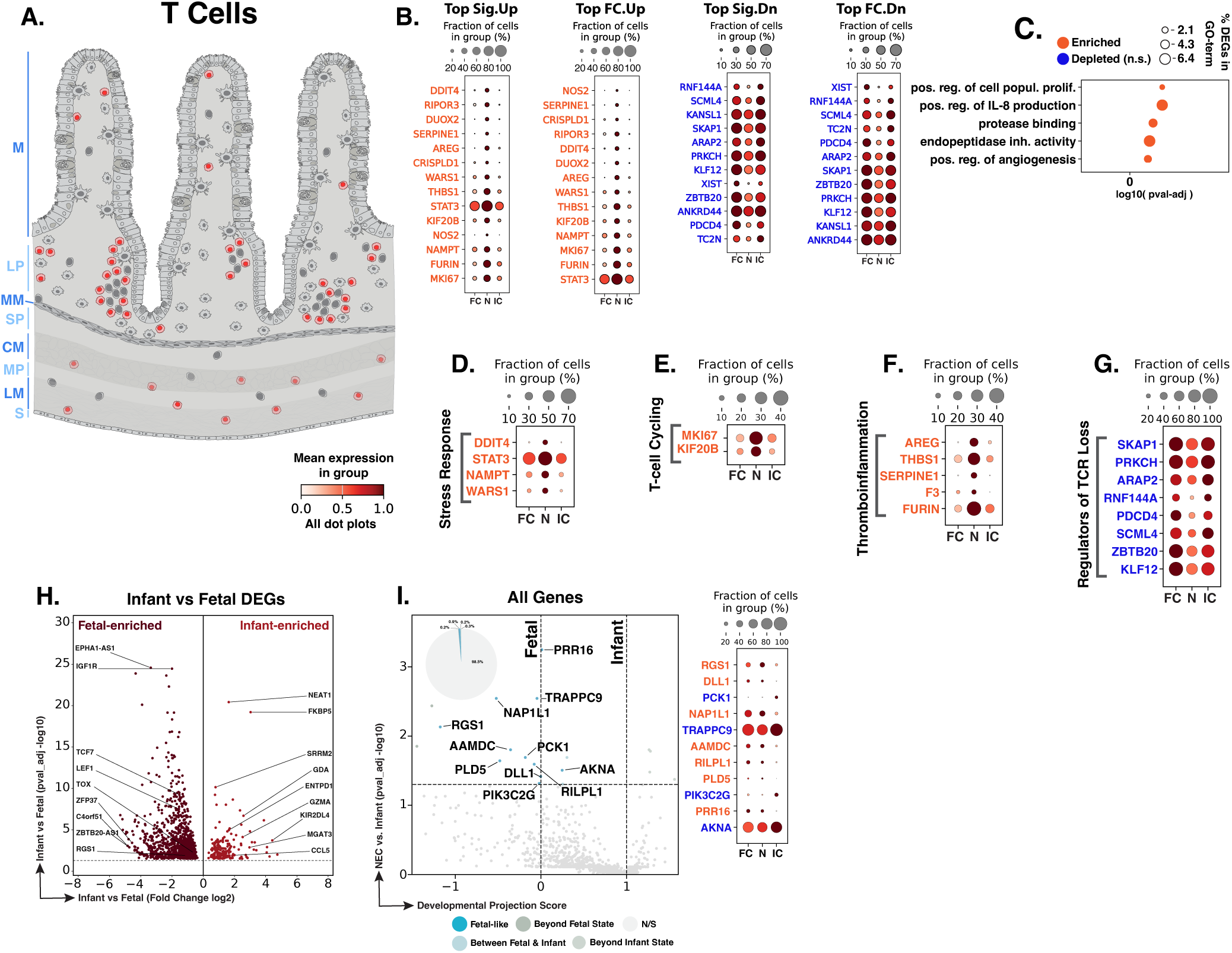
NEC T cells show stress-response and proliferative activation with tissue-repair and T-cell-receptor-associated changes and developmental displacement. **A.** Schematic showing T-cell distribution (red) throughout the intestinal wall. Blue text to the left of the Panel A schematic indicates general anatomical regions: M, mucosa; LP, lamina propria; MM, muscularis mucosae; SP, submucosal plexus; CM, circular muscle; MP, myenteric plexus; LM, longitudinal muscle; S, serosa. **B.** Dot plots of top NEC-specific genes ranked by significance and fold change. **C.** GO-term analysis showing enriched biological processes among NEC-associated DEGs. **D–G.** Curated gene panels showing stress-response, T-cell cycling, tissue-interaction/thrombo-inflammatory, and T-cell-receptor-regulatory programs. In panels **B** and **D–G**, orange gene text indicates upregulation in NEC relative to infant controls, whereas blue gene text indicates downregulation. **H.** Volcano plot of DEGs between fetal and infant controls that define the developmental trajectory for T cells. **I.** Gene-level maturity analysis of fetal-versus-infant DEGs, highlighting the subset also significantly changed in NEC relative to infant controls and positioned relative to fetal and infant reference states, and their corresponding dotplots. The developmental projection score in panel I is displayed on a symmetric logarithmic (symlog) scale. For expression dot plots, dot size indicates the fraction of nuclei expressing each gene and dot color indicates mean expression. In maturity plots, vertical dotted lines indicate the fetal and infant reference scores. Colored dots classify NEC-versus-infant DEGs as Fetal-like, Between Fetal & Infant, Beyond Fetal State, or Beyond Infant State; gray N/S dots are fetal-versus-infant DEGs without a significant NEC-versus-infant change.

We identified 1,330 T-cell nuclei across the atlas. Compared with macrophages, T cells showed a smaller transcriptional response to NEC, with 39 genes significantly altered in NEC compared to infant controls, including 25 upregulated and 14 downregulated genes (Fig. 29B). GO-term enrichment linked the upregulated genes to cell-population proliferation, IL-8 production, protease binding, endopeptidase inhibitor activity, and angiogenesis, whereas no depleted GO-terms reached significance (Fig. 29C).

NEC T cells also showed features of cellular stress and cytokine exposure (Fig. 29D). Increased *DDIT4* was consistent with stress-responsive restraint of mTOR signaling, while higher *STAT3* linked NEC T cells to the Th17-polarizing pathway implicated in experimental NEC.^32,1233^ Increased *NAMPT* suggested greater reliance on NAD-salvage metabolism, which supports NAD⁺ availability and survival in activated T cells.^1234^ Higher *WARS1* expression was consistent with interferon exposure because it is induced by type I and type II interferons.^1235^ Together, these changes suggest that NEC T cells are responding to metabolic stress, inflammatory signaling, and interferon exposure while engaging stress-limiting feedback.

Stress adaptation was accompanied by a proliferative state (Fig. 29E). Increased *MKI67* marks nuclei engaged in active cell-cycle progression, and increased *KIF20B* supports completion of cytokinesis through its organization of the central spindle and midbody.^180,1236^ Their increased expression indicates a greater representation of cycling T cells in NEC, as previously reported.^18^ The concurrent induction of *STAT3*, *NAMPT*, and *DDIT4* places this proliferation in a cytokine-exposed, metabolically demanding environment.^18^

A third group of genes linked T-cell activation with tissue interactions, repair-associated signaling, and thrombo-inflammatory activity in NEC (Fig. 29F). Increased *AREG* may provide a tissue-supportive signal because amphiregulin from tissue-associated regulatory T cells can act on local progenitor cells during repair, while *THBS1* can influence T-cell migration through extracellular matrix interactions involving calreticulin and CD47.^1237–1238^ These potentially adaptive features occurred alongside increased *SERPINE1*, which can promote Th17 differentiation and inflammatory activity, and increased *F3*, adding a coagulation-associated feature.^1239^ Tissue-factor-expressing CD4⁺ T cells have also been identified in pediatric inflammatory bowel disease and experimental colitis, where they generate tissue-factor-dependent thrombin activity.^1240^ Higher *FURIN* further linked this response to T-cell activation, TCR-regulated transcription, cytokine processing, and immune tolerance.^1241–1242^ Overall, NEC T cells retain tissue-repair and migratory signals through *AREG* and *THBS1*, while *SERPINE1* and *F3* add Th17-associated and procoagulant features and *FURIN* supports ongoing activation and cytokine processing.

Alongside proliferation and tissue-interaction changes, NEC T cells showed lower expression of genes involved in immune-receptor coupling and intracellular regulation (Fig. 29G). Reduced *SKAP1* and *PRKCH* may weaken TCR-dependent activation: *SKAP1* links TCR stimulation to integrin activation, ICAM1 adhesion, and stable interactions with antigen-presenting cells, while *PRKCH*/PKCη supports TCR signaling and homeostatic proliferation of mature CD8⁺ T cells.^1243–1244^ Lower *ARAP2* and *RNF144A* extended these changes to cytokine-signaling control. *ARAP2* can support IFN-γ signaling by limiting SOCS1 inhibition of the JAK-STAT pathway, whereas *RNF144A* helps maintain IL-2R-STAT5 signaling and CD8⁺ T-cell effector function.^1245^ Increased *WARS1* together with lower *ARAP2* is consistent with interferon exposure accompanied by reduced intracellular signaling control, while lower *RNF144A* may weaken IL-2-responsive signaling.

Additional reductions involved translational, metabolic, and memory-associated regulation. *PDCD4* contributes to activation-linked translational and metabolic control in CD8⁺ T cells, while *SCML4* supports the metabolic fitness and function of tissue-resident memory CD8⁺ T cells.^1246–1247^ *ZBTB20* regulates CD8⁺ T-cell metabolism and memory differentiation, so its reduction may alter differentiation control.^66^ Lower *KLF12* expression pointed to additional transcriptional regulation, although its specific role in intestinal T cells remains unclear; KLF12 is an established CtBP-associated transcriptional repressor.^1248^ All together, these changes suggest reduced TCR-to-adhesion coupling, cytokine-signaling control, and metabolic and memory-associated regulation in NEC T cells.

Of the 1,273 genes distinguishing fetal from infant control T cells, 19 were also dysregulated in NEC versus infant controls, including 11 with fetal-like expression (57.9% of the 19 genes; Fig. 29H,I). Developmentally regulated genes included fetal-high *TCF7* and *LEF1*, established regulators of T-cell development, and fetal-high *RGS1*, which regulates intestinal T-cell trafficking.^1249–1250^ Infant-high *CCL5*, *GZMA*, and *ENTPD1* reflected postnatal increases in effector, cytotoxic, and purinergic-regulatory functions.^1251–1253^ Among the genes altered in NEC, *RGS1* was higher than in both reference groups, whereas the infant-high genes *PCK1* and *AKNA* were reduced toward fetal levels. *PCK1* supports glycogen-dependent antioxidant metabolism required for CD8⁺ memory T-cell formation and maintenance, while *AKNA*, the only transcription factor with fetal-like expression, regulates *CD40*, *CD40LG*, and activation-associated genes including *IL2*.^1254–1256^ Overall, NEC T cells largely retained an infant-like developmental profile, with limited fetal-like changes involving intestinal trafficking, memory-associated metabolism, and costimulatory regulation.

NEC T cells show a smaller but coherent response involving stress adaptation, metabolism, interferon exposure, proliferation, tissue interaction, Th17-associated activity, and procoagulant features. Initiation may involve recruitment and early expansion of T cells in a cytokine-rich environment, consistent with prior NEC evidence for T-cell involvement. During progression, proliferative and inflammatory signals may persist while reduced TCR-to-adhesion coupling, cytokine-signaling control, translational regulation, and memory-associated differentiation limit effective immune coordination. When viewed alongside reduced macrophage MHC-II expression, these findings suggest weakened antigen-dependent macrophage-T-cell communication despite ongoing inflammation. Resolution may depend on restoring regulatory restraint, antigen recognition, and tissue-adapted T-cell function.

### B cells

Intestinal B cells are found in Peyer’s patches, isolated lymphoid follicles, and the lamina propria, where antigen recognition combines with T-cell and innate signals to support IgA class switching and differentiation into antibody-secreting plasmablasts and plasma cells.^1257^ Secretory IgA helps contain luminal microbes by coating and aggregating bacteria and limiting their contact with intestinal tissues (Fig. 30A).^1258^ In premature infants, mucosal antibody production remains immature, making maternal milk IgA an important source of intestinal antibody during early postnatal life and protective in experimental NEC.^12^ Human mass-cytometry studies found depletion of circulating total and pre-switched B cells with increased plasmablasts, while overall mucosal B-cell abundance was unchanged.^1150^ However, the transcriptional state of intestinal B cells in NEC, including BCR signaling, cytokine sensing, and antibody-secreting differentiation, remains poorly defined.

**Fig. 30.**
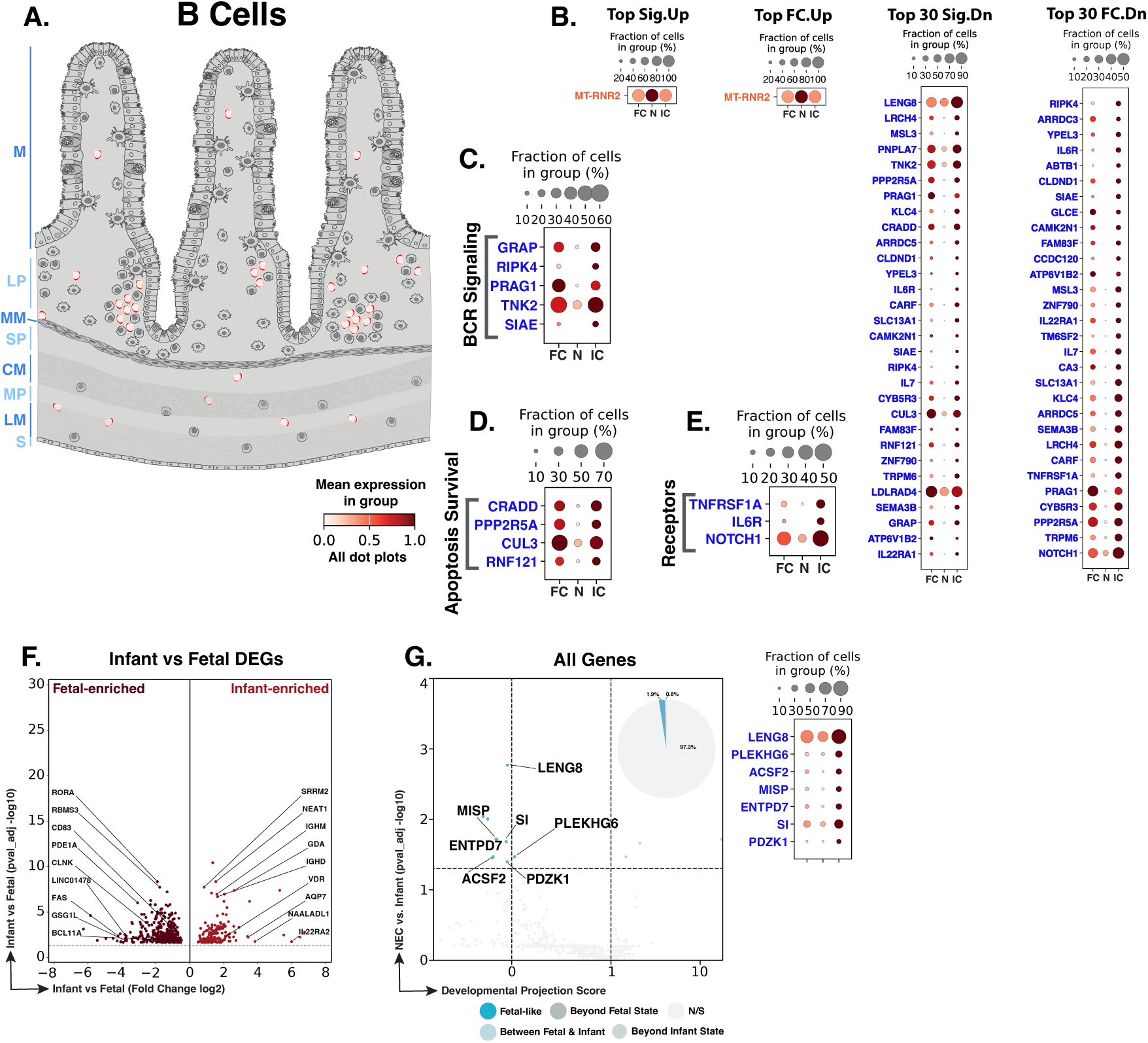
NEC B cells show a restricted transcriptional response characterized by reduced B-cell-receptor/NF-κB signaling, cytokine-receptor and apoptosis-survival programs, and developmental displacement. **A.** Schematic showing B-cell distribution (red) throughout the intestinal wall. Blue text to the left of the Panel A schematic indicates general anatomical regions: M, mucosa; LP, lamina propria; MM, muscularis mucosae; SP, submucosal plexus; CM, circular muscle; MP, myenteric plexus; LM, longitudinal muscle; S, serosa. **B.** Dot plots of top NEC-specific genes ranked by significance and fold change. **C–E.** Curated gene panels showing B-cell-receptor/NF-κB signaling, apoptosis-survival, and receptor-associated programs. In panels **B–E**, orange gene text indicates upregulation in NEC relative to infant controls, whereas blue gene text indicates downregulation. **F.** Volcano plot of DEGs between fetal and infant controls that define the developmental trajectory for B cells. **G.** Gene-level maturity analysis of fetal-versus-infant DEGs, highlighting the subset also significantly changed in NEC relative to infant controls and positioned relative to fetal and infant reference states, and their corresponding dotplots. The developmental projection score in panel G is displayed on a symmetric logarithmic (symlog) scale. For expression dot plots, dot size indicates the fraction of nuclei expressing each gene and dot color indicates mean expression. In maturity plots, vertical dotted lines indicate the fetal and infant reference scores. Colored dots classify NEC-versus-infant DEGs as Fetal-like, Between Fetal & Infant, Beyond Fetal State, or Beyond Infant State; gray N/S dots are fetal-versus-infant DEGs without a significant NEC-versus-infant change.

We identified 404 B-cell nuclei across the atlas. Differential-expression analysis identified 49 genes significantly changed in NEC relative to infant controls, including one upregulated and 48 downregulated genes (Fig. 30B). No GO-terms met statistical significance, likely in part because of the relatively small number of B-cell nuclei available for analysis. The only increased transcript was *MT-RNR2*. Although this mitochondrial 16S rRNA locus contains the coding sequence for humanin, higher *MT-RNR2* expression does not establish humanin peptide production or cytoprotection.^1259^ The 48 downregulated genes included regulators of antigen-receptor signaling, cytokine and differentiation sensing, cell fate, protein turnover, and intracellular signaling. In contrast to the broad inflammatory response in macrophages and the proliferative and stress-associated features in T cells, NEC B cells showed no broad inflammatory, proliferative, or antibody-secreting signature.

The downregulated genes centered on BCR-signal transmission and calibration (Fig. 30C). *GRAP* connects BCR engagement to ERK activation in human B cells, whereas *RIPK4*/PKK supports BCR-induced NF-κB activation, proliferation, survival, and maintenance of mature recirculating B cells.^1260–1261^ Lower *GRAP* and *RIPK4* are consistent with weaker transmission of antigen-receptor signals into proliferative and survival pathways. However, the inhibitory regulator *SIAE* was reduced simultaneously. SIAE controls BCR-signal strength through sialic-acid-dependent regulation of CD22-associated inhibitory signaling and contributes to peripheral B-cell tolerance.^1262^ This imbalance may blunt antigen-receptor transmission and diminish the precision of its inhibitory calibration. Reduced *PRAG1*/PEAK2 and *TNK2*/ACK1 extended the deficit to scaffold- and kinase-dependent signaling. *PRAG1* forms a signaling complex with CSK that regulates SRC-family kinase activity, adhesion, and cell morphology, whereas *TNK2* can couple activated receptors to AKT signaling.^1263–1264^ Together, reduced expression of activating, inhibitory, scaffold, and kinase components suggests less reliable transmission and calibration of BCR signals in NEC B cells.

Genes governing cell fate, receptor trafficking, ubiquitin-dependent processing, and phosphorylation were also reduced (Fig. 30D). *CRADD* recruits caspase-2 into apoptotic signaling complexes formed during cellular stress, whereas *CUL3* supports pro-B-cell proliferation and regulates the surface abundance and internalization of the inhibitory B-cell receptor CD22.^1265–1266^ *RNF121* can promote stimulus-responsive NF-κB activation through ubiquitin-dependent mechanisms, whereas *PPP2R5A* encodes a PP2A regulatory subunit that restrains AKT and NF-κB signaling.^1267–1268^ Their simultaneous loss may impair coordination of stress-induced cell fate with receptor turnover, ubiquitination, and phosphorylation-dependent signaling.

Receptor downregulation pointed to weaker sensing of inflammatory and differentiation cues (Fig. 30E). *TNFRSF1A* encodes TNFR1, which transmits TNF signals into inflammatory and survival pathways through NF-κB or, under different signaling conditions, cell death pathways.^1269–1270^ *TNFRSF1A* downregulation may diminish reception of TNF signals. *IL6R* downregulation may similarly limit IL-6-dependent differentiation of activated B cells into antibody-secreting cells; reduced *NOTCH1* may limit Notch-supported differentiation of activated mature B cells toward antibody-secreting states.^1271–1272^ Despite the cytokine-rich NEC environment generated partly by inflammatory macrophages, B cells appear less equipped to integrate TNF, IL-6, antigenic, and differentiation cues. This limited signal-integration capacity may contribute to the absence of broad inflammatory or plasma-cell-like activation seen in NEC.

Of the 521 genes distinguishing fetal from infant control B cells, 14 were also dysregulated in NEC versus infant controls, including 10 with fetal-like expression (71.4% of the 14 genes; Fig. 30F,G). Developmentally regulated genes included fetal-high *BCL11A*, an established regulator of lymphoid development, and infant-high *IGHM* and *IGHD*, which mark IgM- and IgD-bearing receptors characteristic of mature naïve B cells.^1273–1275^ The small set of NEC-associated developmental genes was functionally diverse and did not define a clear B-cell maturation pattern. *LENG8* is involved in nuclear RNA quality control, whereas *ACSF2* encodes a mitochondrial acyl-CoA synthetase involved in fatty acid activation.^1276–1277^ Other genes, including *ENTPD7*, *MISP*, *SI*, and *PDZK1*, are more closely associated with intestinal epithelial functions such as luminal ATP regulation, microvillar organization, carbohydrate digestion, and brush-border transport.^1278–1281^ Overall, NEC B cells largely retained the infant developmental profile, with only a small and heterogeneous set of developmental genes altered.

NEC B cells show a small, predominantly downregulated response while retaining a largely postnatal developmental profile. Initiation does not appear to be driven by strong local B-cell inflammatory activation in this atlas. During progression, reduced BCR signaling, cytokine sensing, and differentiation-associated genes may limit how B cells respond to antigenic and inflammatory cues in the injured intestine. This muted response contrasts with macrophage inflammatory activation and T-cell stress/proliferation. Failed resolution may reflect broader disruption of coordinated innate and adaptive immunity, where reduced macrophage MHC-II, altered T-cell receptor coupling, and weak B-cell antigen/cytokine integration limit effective immune adaptation and mucosal antibody-associated recovery.

### Enteric nervous system compartment

The enteric nervous system, marked by *UCHL1* and *ELAVL3* (Fig. 2), comprises interconnected neuronal and glial networks organized principally within the myenteric and submucosal plexuses. The myenteric, or Auerbach’s, plexus lies between the circular and longitudinal muscle layers and predominantly regulates motility. Submucosal plexuses, including Meissner’s plexus, lie within the submucosa and regulate mucosal functions, including secretion and local blood flow. Enteric motor neurons coordinate muscle contraction and relaxation, secretomotor and vasodilator neurons regulate epithelial secretion and circulation, and sensory neurons and interneurons organize local reflex circuits. Enteric glia participate in communication among these circuits, the epithelium, and immune cells.^1282–1285^ Human NEC studies have documented neuronal and glial loss, enteric neural apoptosis, and persistent abnormalities of nitrergic and VIP-associated innervation.^1286–1287^

To define the cell populations belonging to the enteric nervous system in our study, marker-based annotation resolved glia and four neuronal populations (Fig. 31). Aside from *S100B* and *SOX10* (Fig. 2), glia were marked by *QKI*, *PTPRZ1*, and *CDH19*, cholinergic excitatory neurons by *SLC5A7*, *CHAT*, and *COLQ*, and inhibitory neurons by enriched *NOS1* and *VIP* expression.^1288^;^49,1289–1290^) Two additional neuronal populations contained low and variable markers consistent with putative interneuron and sensory/IPAN-like features, but these groups could not be confidently resolved in this dataset and were therefore excluded from the detailed NEC disease sections.^49,1291^

**Fig. 31.**
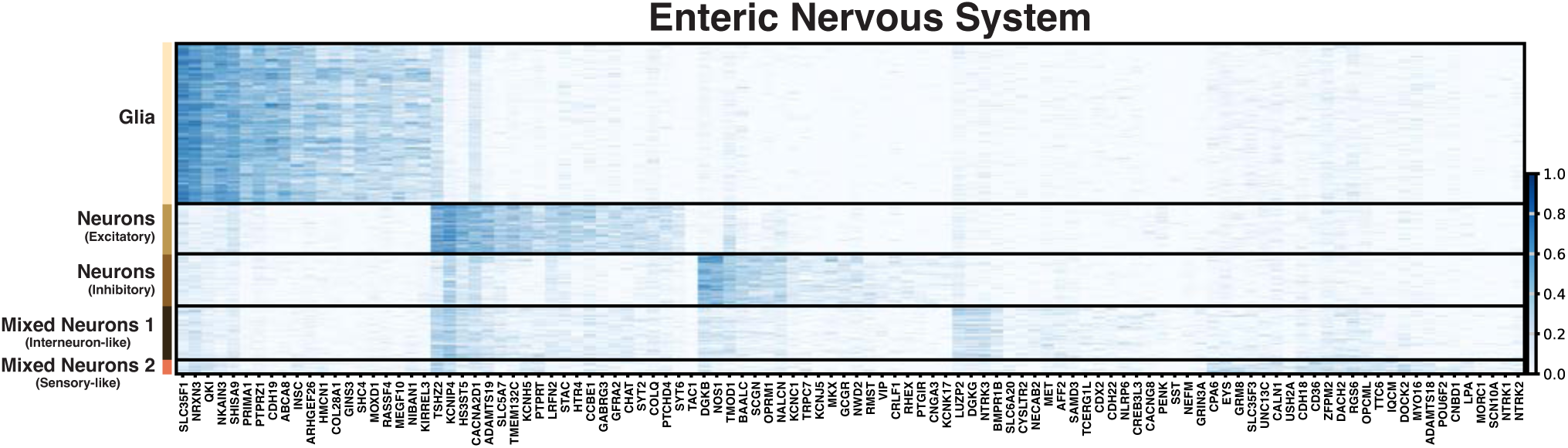
Enteric neural cell-type signatures define glial and neuronal populations. Heatmap showing marker-gene expression across glia, excitatory neurons, inhibitory neurons, and two mixed neuronal populations with unresolved identities. Distinct transcriptional signatures support neural cell-type assignments and provide the framework for the detailed enteric neural analyses that follow.

The following sections examine how NEC alters enteric glia and transcriptionally resolved neuronal populations. Glia capture the neuroglial response at the interface of epithelial support, immune communication, neuronal maintenance, and motility. Inhibitory neurons capture the NOS1/VIP-associated relaxation arm of the enteric motor system, providing a population-resolved extension of previous human NEC studies implicating inhibitory-neural injury. In contrast, excitatory neurons showed minimal NEC-associated transcriptional changes despite sampling comparable to inhibitory neurons, indicating broad preservation of the contractile arm of enteric reflexes. Together, these findings localize the principal enteric nervous system abnormalities to glia and inhibitory neurons, where impaired neuroglial support and inhibitory neuromuscular signaling may compromise relaxation, reflex coordination, and intestinal motility.

### Glia

Enteric glia are glial cells of the enteric nervous system found in the intestinal mucosa, submucosal and myenteric plexuses, interganglionic nerve-fiber bundles, and along nerve fibers in the circular and longitudinal muscle layers.^1292^ Within enteric ganglia, connexin-43-dependent glial communication and calcium signaling help regulate neuronal circuits and motor responses, while mucosal glia predominantly influence epithelial cell secretion and microbial ecology (Fig. 32A).^23,1293–1294^ Human transcriptomic single cell studies have identified substantial molecular heterogeneity within enteric glial cells and Schwann cell lineages.^49,1295^ Human NEC tissue also shows heterogeneous glial injury: 11 of 20 specimens had disrupted myenteric architecture, reduced *GFAP* expression, and loss of submucosal and mucosal innervation, while others retained more preserved neural structure; *CCL20* was increased in the myenteric plexus and strongly expressed by enteric glia.^1296^ In experimental NEC, TLR4-dependent glial loss preceded inflammatory injury and impaired motility, whereas glial *Tlr4* deletion preserved glial viability and motility and reduced disease severity which suggests a strong implication of enteric glia-gut microbiota functional interactions in NEC development.^24^ Human-milk oligosaccharides likewise preserved enteric glia and motility in experimental NEC.^1297^ These studies establish enteric glial cell injury in NEC but do not define the molecular characteristics retained by glia that remain in diseased tissue.

**Fig. 32.**
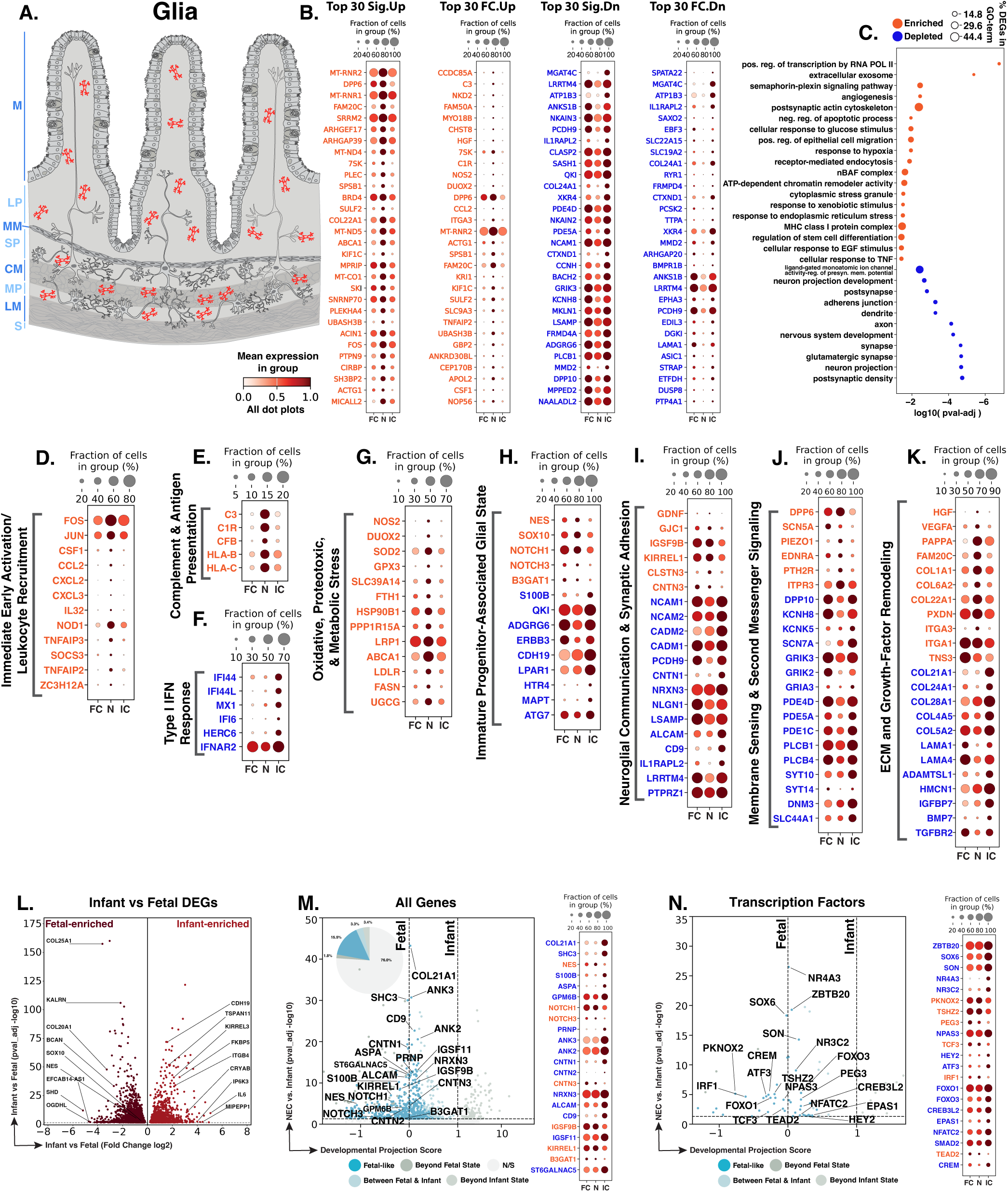
NEC enteric glia show inflammatory and stress activation with altered glial-state, neuroglial-support, membrane-sensing, extracellular-matrix, and maturity-associated programs. **A.** Schematic showing enteric glia distribution (red) throughout the intestinal wall. Blue labels indicate anatomical regions: M, mucosa; LP, lamina propria; MM, muscularis mucosae; SP, submucosal plexus; CM, circular muscle; MP, myenteric plexus; LM, longitudinal muscle; S, serosa. **B.** Dot plots of top NEC-specific genes ranked by significance or fold change. **C.** GO-term enrichment of NEC-associated DEGs. **D–K.** Curated gene panels showing immediate-early/leukocyte-recruitment, complement/antigen-presentation, type I IFN, oxidative/proteotoxic/metabolic-stress, immature glial-state, neuroglial-support, membrane/second-messenger signaling, and ECM/growth-factor remodeling programs. In panels **B** and **D–K**, orange gene text indicates upregulation in NEC relative to infant controls, and blue indicates downregulation. **L.** Volcano plot of fetal-versus-infant DEGs defining the glial developmental trajectory. **M–N.** Gene-level and transcription-factor maturity analyses of fetal-versus-infant DEGs, highlighting the subset also significantly changed in NEC relative to infant controls and positioned relative to fetal and infant reference states, and their corresponding dotplots. Developmental projection scores in panels M-N are shown on a symlog scale; vertical dotted lines mark fetal and infant reference scores. Colored dots classify NEC-versus-infant DEGs as Fetal-like, Between Fetal & Infant, Beyond Fetal State, or Beyond Infant State; gray N/S dots are fetal-versus-infant DEGs without a significant NEC-versus-infant change.

We identified 2,154 glial nuclei across the atlas, including 1,127 fetal, 480 NEC, and 547 infant control nuclei. Differential-expression analysis detected 1,394 NEC-versus-infant glial DEGs (Fig. 32B). Enriched GO-terms involved transcriptional and chromatin regulation, inflammatory and TNF-associated signaling, hypoxia and ER-stress adaptation, immune interaction, angiogenic and epithelial-migration-associated signaling, and stress-granule biology (Fig. 32C). Depleted terms included ligand-gated ion-channel activity, neuronal projection development, adherens junction organization, axonal and dendritic structure, glutamatergic signaling, synaptic organization, and postsynaptic-density biology. NEC glia therefore displayed inflammatory, stress-adaptive, trophic, and repair-associated activity alongside depletion of genes supporting mature neuroglial communication.

Inflammatory activation combined immediate-early transcription with myeloid recruitment. Increased *FOS* and *JUN* marked AP-1 activity, while higher *CCL2*, *CXCL2*, and *CXCL3* supported recruitment of monocytes and other CXCR2-dependent myeloid cells (Fig. 32D).^951,1298–1299^ Increased *CSF1* could further support macrophage survival and differentiation. Previous studies have shown that enteric glial CCL2 and CSF1 contribute to monocyte accumulation and macrophage responses after intestinal injury.^1300–1301^ Higher *IL32* and *NOD1* added inflammatory and microbial-pattern sensing.^1302^ At the same time, increased *TNFAIP3*, *SOCS3*, *ZC3H12A*, and *TNFAIP2* reflected feedback mechanisms that can restrain inflammatory signaling.^1161,1303–1305^ Thus, NEC glia may contribute to myeloid recruitment while simultaneously engaging mechanisms that limit prolonged inflammation.

Complement and antigen presentation genes broadened this immune response. Increased *C3*, *C1R*, and *CFB* represented multiple components of the complement system, although higher transcript abundance does not establish complement activation or deposition (Fig. 32E).^1306^ Higher *HLA-B* and *HLA-C* suggested greater MHC-I-associated antigen presentation capacity, consistent with the ability of enteric glia to perform MHC-I presentation and cross-presentation.^1307^ In contrast, lower *IFI44*, *IFI44L*, *MX1*, *IFI6*, *HERC6*, and *IFNAR2* pointed to reduced representation of interferon-responsive genes (Fig. 32F).^407,1308–1310^ NEC glia therefore combined increased complement and MHC-I-associated expression with reduced expression of interferon-responsive genes.

Immune activation also occurred alongside oxidative, proteotoxic, and metabolic stress (Fig. 32G). Increased *NOS2* and *DUOX2* suggested greater nitric-oxide- and hydrogen-peroxide-generating capacity, while higher *SOD2* and *GPX3* reflected concurrent antioxidant responses.^50,1311–1313^ Higher *SLC39A14* and *FTH1* linked this response to metal uptake and iron storage, while *HSP90B1* and *PPP1R15A* pointed to ER protein quality control and integrated stress signaling.^1314–1317^ Increased *LRP1*, *LDLR*, and *ABCA1* implicated extracellular cargo and lipid handling, while higher *FASN* and *UGCG* pointed to fatty acid and glycosphingolipid synthesis.^1318–1323^ This profile is consistent with NEC glia experiencing oxidative and proteotoxic stress while increasing antioxidant, iron-buffering, endocytic, and lipid-metabolic responses.

NEC glia showed increased developmental and progenitor-associated expression alongside reduced expression of genes associated with mature glial functions. Increased *NES*, *NOTCH1*, *NOTCH3*, and *B3GAT1*/HNK-1 reintroduced developmental and progenitor-associated features (Fig. 32H).^1324–1326^ Increased *SOX10* was distinct because SOX10 is retained in adult enteric glia and regulates glial identity.^1327–1328^ At the same time, lower *S100B*, *QKI*, *ADGRG6*, *ERBB3*, and *CDH19* pointed to reduced expression of genes associated with differentiated glial functions and Schwann-lineage maintenance.^1329–1333^ Reduced *LPAR1* was particularly notable because LPAR1 regulates glial calcium signaling, neural activity, and intestinal motility.^1334^ Lower *HTR4*, *MAPT*, and *ATG7* further suggested changes in ENS support, cytoskeletal organization, and autophagy, although direct enteric-glial mechanisms are less established.^1335–1337^ Together, NEC glia showed developmental features alongside reduced expression of genes associated with mature glial functions.

Several trophic and cell-cell communication genes increased in NEC glia, but a broader set of neuronal-glial interface/communication genes was reduced. Increased GDNF could support enteric-neuron survival and epithelial barrier repair, while higher GJC1, IGSF9B, KIRREL1, CLSTN3, and CNTN3 suggested altered gap-junctional communication, cell adhesion, and neuron-glia interface functions (Fig. 32I).^1338–1345^ These increases occurred alongside lower *NCAM1*, *NCAM2*, *CADM1*, *CADM2*, *CNTN1*, *PTPRZ1*, *NRXN3*, and *NLGN1*, indicating reduced expression of multiple adhesion and synapse-associated genes.^1346–1352^ Lower *PCDH9*, *LSAMP*, *ALCAM*, *CD9*, *IL1RAPL2*, and *LRRTM4* extended this loss to additional cell-surface organizers.^1353–1358^ Increased *GDNF* may therefore represent a trophic response occurring alongside reduced machinery for stable neural interactions. The *GDNF*-high/*NCAM1*-low pattern is particularly interesting because GDNF-NCAM1 signaling can support neurogenesis in enteric glia, although this has been demonstrated in Hirschsprung disease rather than NEC.^1359^

Changes in membrane sensing and second-messenger genes further suggested altered neuron-glia communication (Fig. 32J). Opposing *DPP6* and *DPP10* expression indicated differential regulation of Kv4 potassium channels, while higher *SCN5A*, *PIEZO1*, *EDNRA*, *PTH2R*, and *ITPR3* implicated sodium-channel, mechanosensory, receptor, and calcium-release functions.^1360–1366^ In contrast, *KCNH8*, *KCNK5*, *SCN7A*, *GRIK2*, *GRIK3*, and *GRIA3* were reduced, pointing to changes in ion-channel and glutamate-receptor expression.^1367–1372^ Lower *PDE4D*, *PDE5A*, *PDE1C*, *PLCB1*, and *PLCB4* further suggested altered cAMP, cGMP, phosphoinositide, and calcium signaling.^70–71,1373–1375^ Reduced *SYT10*, *SYT14*, *DNM3*, and *SLC44A1* added changes in vesicle-associated signaling, endocytosis, and choline transport, although their roles in enteric glia remain less clear.^1376–1379^ Thus, NEC glia did not show a uniform increase or decrease in excitability; mechanosensory gains occurred alongside reduced neurotransmitter sensing and second-messenger regulation.

Extracellular-matrix genes also showed mixed changes (Fig. 32K). Increased *HGF* and *VEGFA* could support epithelial repair and angiogenesis, while *PAPPA*, *FAM20C*, and *PXDN* could influence IGF availability, extracellular-protein phosphorylation, and collagen-IV crosslinking.^1380–1384^ Higher *COL1A1*, *COL6A2*, *COL22A1*, *TNS3*, *ITGA1*, and *ITGA3* suggested increased production of selected matrix components and altered cell-matrix attachment.^1385–1390^ In contrast, *COL21A1*, *COL24A1*, *COL28A1*, *COL4A5*, *COL5A2*, *LAMA1*, *LAMA4*, *ADAMTSL1*, and *HMCN1* were reduced, indicating loss of other fibrillar, basement membrane, and extracellular glycoprotein components.^1391–1399^ Lower *IGFBP7*, *BMP7*, and *TGFBR2* further pointed to changes in growth factor availability and BMP/TGF-β signaling.^1400–1402^ This pattern may reflect a repair-associated but compositionally altered glial matrix; protein-level studies will be needed to determine whether these transcriptional changes alter matrix deposition or tissue mechanics.

Of the 3,361 genes distinguishing fetal from infant control glia, 806 were also dysregulated in NEC versus infant controls, including 520 with fetal-like expression (64.5% of the 806 genes; Fig. 32L,M). Developmentally regulated genes included fetal-associated *NES* and *SOX10*, reflecting ENS progenitor and enteric glial lineage specification, and infant-associated *CDH19* and *ITGB4*, reflecting peripheral glial identity, cell adhesion, and interactions with the basement membrane.^1324,1327,1403–1404^ Fetal-like increases in *NES*, *NOTCH1*, and *NOTCH3* pointed to developmental and progenitor-associated signaling, while *CNTN3*, *IGSF9B*, *KIRREL1*, and *B3GAT1* highlighted neural-contact and adhesion functions.^1324–1326,1341–1342,1344^ In contrast, the infant-high genes *S100B*, *ASPA*, *GPM6B*, and *PRNP* were reduced toward fetal levels, suggesting lower expression of genes involved in glial signaling, metabolic support, membrane organization, and axon-glia maintenance.^1329,1405–1407^ Reduced *ANK2*, *ANK3*, *CNTN1*, *CNTN2*, *NRXN3*, *ALCAM*, *CD9*, and *IGSF11* extended this pattern to membrane scaffolding, adhesion, axon-glia interactions, and intercellular coupling.^1351,1355–1356,1408–1411^ Lower *COL21A1*, *SHC3*, and *ST6GALNAC5* further implicated extracellular matrix, neurotrophin-associated signaling, and cell-surface sialylation.^1391,1412–1413^ These changes suggest that the fetal-like component of NEC glia combines progenitor-associated signaling with reduced transcriptional support for mature axon-glia interactions.

At the transcription factor level, fetal-high *SOX6* and *ZBTB20* remained at or near fetal-associated expression in NEC, extending this pattern to regulators of glial differentiation (Fig. 32N). Experimental studies link SOX6 to restraint of glial differentiation and ZBTB20 to astroglial specification.^1414–1415^ Altered *ATF3* and *FOXO3* expression also implicated regulators linked experimentally to injury responses in peripheral glia and antioxidant defense in microglia, respectively (Fig. 32N).^1416–1417^ Overall, a substantial subset showed fetal-like expression involving glial metabolism, membrane organization, axon-glia adhesion, and neurotrophin-associated signaling.

NEC glia show broad transcriptional changes involving immediate-early activity, myeloid recruitment, complement and MHC-I expression, oxidative/proteotoxic stress adaptation, trophic-factor production, lipid handling, cargo trafficking, and matrix remodeling. Initiation may involve glial responses to epithelial injury, microbial signals, and tissue inflammation at neuroimmune interfaces. During NEC progression, glia may help organize inflammatory and repair responses but may provide altered structural and signaling support for enteric neurons, because S100B- and LPAR1-associated functions, neurotransmitter/second-messenger sensing, and neuroglial adhesion are reduced, although higher GDNF may support enteric-neuron survival. Resolution may fail if glia remain stress-adapted but cannot stabilize neural circuits that regulate gastrointestinal functions.

### Inhibitory Neurons

Enteric neurons form interconnected myenteric and submucosal plexuses within the intestinal wall. Inhibitory motor neurons, located mainly in the myenteric plexus, work with excitatory motor neurons to coordinate intestinal smooth muscle activity. By releasing nitric oxide, purines, and inhibitory neuropeptides, they promote muscle relaxation and descending inhibition ahead of advancing luminal contents. This balance between contraction and relaxation is essential for peristalsis, propulsion, and intestinal transit (Fig. 33A).^1418–1420^ Human single-nucleus studies have revealed substantial enteric-neuronal diversity, including multiple *NOS1*-expressing populations with putative inhibitory motor or interneuron identities.^1289^ Prior human NEC studies showed broad ENS injury with prominent loss of nitrergic and VIP-associated innervation. One study reported fewer HuC/D⁺ myenteric neurons and a lower proportion of nNOS⁺ neurons, while another identified reduced VIP- and NOS-immunoreactive structures in the submucosal plexus and circular muscle.^1286–1287^ These studies support impaired inhibitory innervation in NEC but did not resolve the affected neurons into transcriptomic populations.

**Fig. 33.**
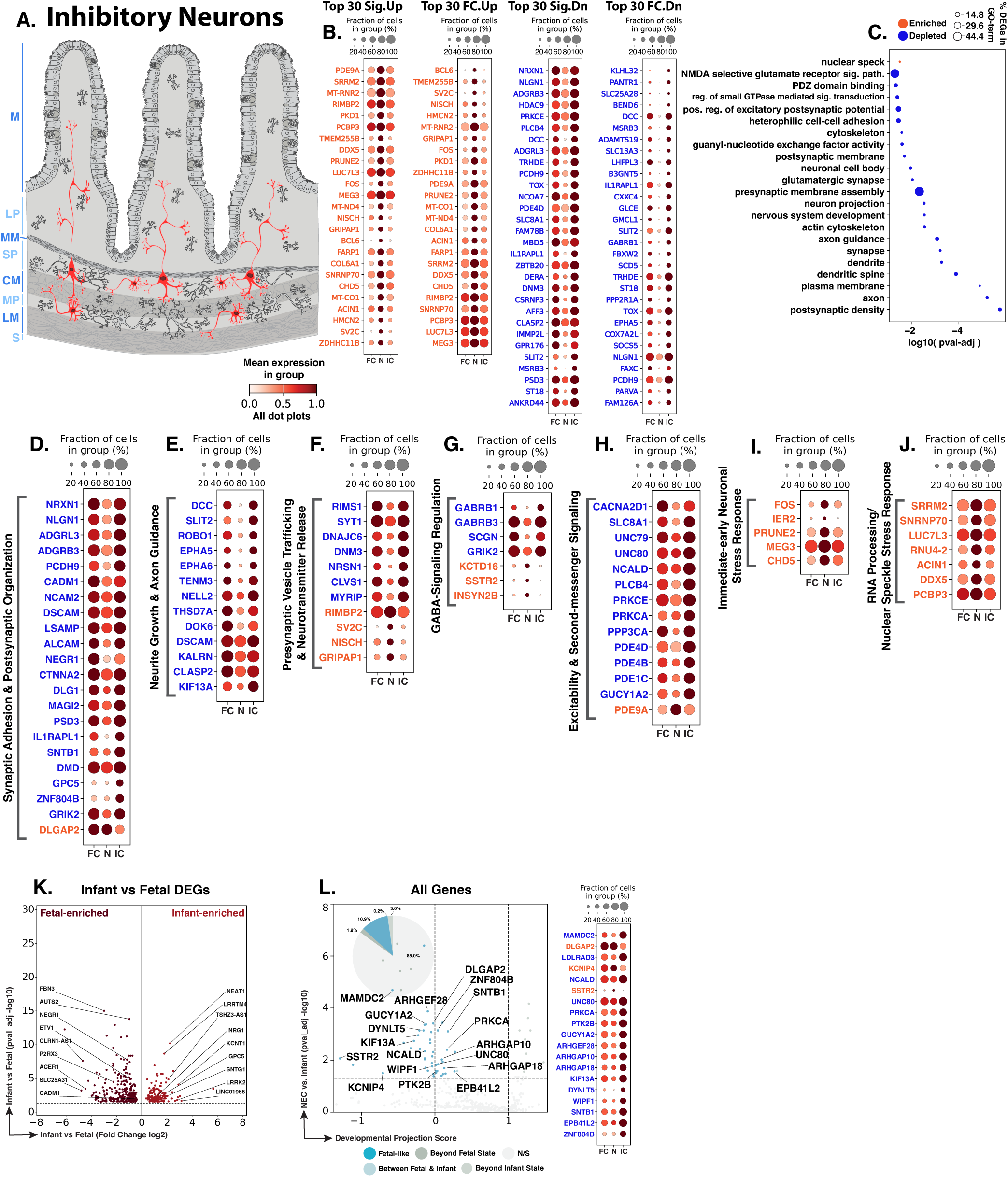
NEC inhibitory enteric neurons show broad loss of synaptic and neuronal organization with altered vesicle trafficking, GABA signaling, excitability, stress-response, RNA-processing, and maturity-associated programs. **A.** Schematic showing the distribution of inhibitory enteric neurons (red) within the intestinal wall. Blue text to the left of the Panel A schematic indicates general anatomical regions: M, mucosa; LP, lamina propria; MM, muscularis mucosae; SP, submucosal plexus; CM, circular muscle; MP, myenteric plexus; LM, longitudinal muscle; S, serosa. **B.** Dot plots of top NEC-specific genes ranked by significance and fold change. **C.** GO-term analysis showing predominantly depleted neuronal and synaptic biological processes among NEC-associated DEGs. **D–J.** Curated gene panels showing synaptic-adhesion/postsynaptic-organization, neurite-growth/axon-guidance, presynaptic-vesicle-trafficking and neurotransmitter-release, GABA-signaling regulation, excitability/second-messenger signaling, immediate-early neuronal stress response, and RNA-processing/nuclear-speck-associated programs. In panels **B** and **D–J**, orange gene text indicates upregulation in NEC relative to infant controls, whereas blue gene text indicates downregulation. **K.** Volcano plot of DEGs between fetal and infant controls that define the developmental trajectory for inhibitory neurons. **L.** Gene-level maturity analysis of fetal-versus-infant DEGs, highlighting the subset also significantly changed in NEC relative to infant controls and positioned relative to fetal and infant reference states, and their corresponding dotplots. The developmental projection score in panel L is displayed on a symmetric logarithmic (symlog) scale. For expression dot plots, dot size indicates the fraction of nuclei expressing each gene and dot color indicates mean expression. In maturity plots, vertical dotted lines indicate the fetal and infant reference scores. Colored dots classify NEC-versus-infant DEGs as Fetal-like, Between Fetal & Infant, Beyond Fetal State, or Beyond Infant State; gray N/S dots are fetal-versus-infant DEGs without a significant NEC-versus-infant change.

Across the atlas, we identified 702 inhibitory neuronal nuclei by expression of canonical inhibitory enteric-neuron markers NOS1, GAL, and VIP, comprising 501 fetal, 109 NEC, and 92 infant control nuclei. Differential-expression analysis detected 276 NEC-versus-infant DEGs in inhibitory neurons, including 35 upregulated and 241 downregulated genes (Fig. 33B). GO-term enrichment highlighted nuclear-speck-associated processes, whereas depleted terms included synaptic organization, postsynaptic density and membrane, presynaptic assembly, neuronal projections, axon guidance, dendrites and dendritic spines, cell adhesion, actin-cytoskeletal organization, and regulation of postsynaptic signaling (Fig. 33C). The predominance of downregulation shows broad loss of neuronal connectivity and communication systems, accompanied by a smaller immediate-early and nuclear RNA-processing response.

Changes in NEC inhibitory neurons are associated with genes involved in neuronal recognition and synaptic organization. *NRXN1* and *NLGN1* were among the most strongly reduced genes, suggesting weaker trans-synaptic adhesion; both are expressed in the enteric nervous system and are reduced in affected Hirschsprung intestine (Fig. 33D).^1421^ Reduced *ADGRL3*, *ADGRB3*, *PCDH9*, *CADM1*, *NCAM2*, *ALCAM*, *NEGR1*, and *LSAMP* extended this pattern across adhesion systems involved in synapse formation, dendritic organization, neuronal recognition, and neurite growth.^1422–1429^ Lower *CTNNA2*, *DLG1*, *MAGI2*, *IL1RAPL1*, and *DMD* further implicated dendritic-spine stabilization, receptor anchoring, postsynaptic organization, and inhibitory-synapse scaffolding.^1430–1434^ Increased *DLGAP2* was a focal exception but did not offset the broader loss of synaptic-organization genes.^1435^ These changes may reduce the ability of NEC inhibitory neurons to establish and maintain stable contacts within enteric circuits.

Genes supporting neuronal projections were also broadly reduced. Lower *SLIT2* and *ROBO1* pointed to weaker axon guidance, branching, and neuronal positioning (Fig. 33E).^1436^ *SLIT2*-*ROBO1* downregulation has also been reported in affected Hirschsprung intestine, although the aganglionic composition of that tissue limits cell-intrinsic interpretation.^725^ Reduced *DCC*, *DSCAM*, *EPHA5*, *EPHA6*, and *TENM3* extended this loss across several axon guidance systems.^1437–1440^ Lower *NELL2*, *THSD7A*, *DOK6*, *KALRN*, and *CLASP2* implicated an altered state of axon extension, neurite branching, dendritic architecture, and microtubule organization, while reduced *KIF13A* could affect microtubule-dependent transport of receptors and membrane cargo into neuronal processes.^1441–1446^ Thus, molecular support for maintaining neuronal processes within the injured intestinal wall was broadly reduced.

Presynaptic changes centered on machinery needed for neurotransmitter release. Reduced *RIMS1* and *SYT1* expression pointed to potential defects in active-zone organization, vesicle priming, calcium-dependent exocytosis, and vesicle retrieval (Fig. 33F).^1447–1448^ Lower *DNAJC6* and *DNM3* further suggested impaired clathrin-coat removal and synaptic-vesicle recycling, while reduced *NRSN1* and *MYRIP* implicated transport-vesicle localization and secretory trafficking.^1378,1449–1451^ In contrast, *RIMBP2*, *SV2C*, and *GRIPAP1* were increased, affecting active-zone, synaptic-vesicle, and recycling-endosome machinery.^1452–1454^ These increases do not establish preserved release capacity because core genes required for vesicle priming, exocytosis, and recycling were simultaneously reduced. The overall pattern is consistent with less reliable inhibitory neurotransmission in NEC, with a compromised coordination of vesicle release and recycling.

NEC inhibitory neurons also showed altered receptor and neuromodulatory gene expression. Reduced *GABRB1* and *GABRB3* indicated lower representation of GABA_A_-receptor β subunits that are normally expressed across enteric neuronal populations (Fig. 33G).^1455^ Because GABA responses depend on receptor composition and circuit context, these changes may alter inhibitory-neuron responsiveness to GABA.^1456^ *SCGN* was also reduced, indicating loss of a calcium-sensitive neurosecretory feature. Secretagogin is expressed in subsets of intestinal enteric neurons and has been linked experimentally to SNAP-25-associated regulated exocytosis.^1457–1459^ In contrast, increased *KCTD16* and *SSTR2* suggested greater GABA_B_-receptor-associated and somatostatin-dependent modulation.^1460–1461^ Thus, these changes indicate a reorganization of inhibitory-neuron signaling in NEC, with reduced GABA_A_-receptor and neurosecretory features occurring alongside increased GABA_B_- and somatostatin-associated modulation.

Genes that translate membrane stimulation into intracellular signaling and synaptic transmission were also reduced (Fig. 33H). Lower *CACNA2D1* and *SLC8A1* suggested altered voltage-gated calcium-channel support and sodium-calcium exchange, potentially affecting calcium entry and recovery from calcium transients.^1462–1463^ Reduced *UNC79* and *UNC80* pointed to weaker support for the NALCN channel complex, which contributes to background sodium conductance, resting membrane potential, and neuronal excitability.^1464^ Lower *NCALD* further implicated calcium sensing, cytoskeletal regulation, and membrane trafficking.^1465^ Reductions in *PLCB4*, *PRKCE*, *PRKCA*, and *PPP3CA* affected phosphoinositide, protein kinase C, and calcium-calcineurin pathways that regulate neuronal signaling and synaptic plasticity.^1466–1469^ Cyclic-nucleotide regulation was also altered, with reduced *PDE4D*, *PDE4B*, *PDE1C*, and *GUCY1A2*, while *PDE9A* was increased. This pattern is consistent with broad disruption of cyclic nucleotide control in NEC, with the combination of reduced *GUCY1A2* and increased *PDE9A* particularly supporting diminished cGMP signaling. Together, the coordinated reduction of calcium-handling, NALCN-complex, phosphoinositide, PKC/calcineurin, and cyclic nucleotide signaling genes indicates a broad weakening of the molecular machinery that couples membrane excitability to intracellular signaling and synaptic transmission in NEC inhibitory neurons.

These neurons also showed evidence of an injury-associated stress response. Increased *FOS* and *IER2* marked immediate-early transcription consistent with neuronal stimulation or tissue injury (Fig. 33I). FOS is rapidly induced after experimental neuronal injury and ischemia, although it does not identify the specific NEC stimulus.^1470–1471^ *IER2* is similarly induced by external stimuli, although its role in enteric neurons remains unclear.^1472^ Increased *PRUNE2*, *MEG3*, and *CHD5* extended this response to neuronal morphology, injury-associated long non-coding RNA, and chromatin organization.^1473–1475^ These changes are consistent with active nuclear adaptation to the injured environment of NEC tissues. The enriched nuclear-speck program provided an additional layer of nuclear adaptation. Increased *SRRM2*, a structural organizer of nuclear speckles and splicing-associated condensates, was accompanied by higher *SNRNP70*, *LUC7L3*, *RNU4-2*, *ACIN1*, and *DDX5*, spanning splice-site recognition and selection, tri-snRNP function, alternative-splicing regulation, and RNA-helicase activity (Fig. 33J).^1476–1481^ Increased *PCBP3* extended this pattern to poly(C)-binding activity.^1482^ These findings are consistent with stress-associated regulation of RNA processing. Experimental work demonstrates that cellular stress can reorganize splicing regulators and modify splicing control, including through heat-induced nuclear stress bodies.^1483^

Of the 433 genes distinguishing fetal from infant control inhibitory neurons, 69 were also dysregulated in NEC versus infant controls, including 47 with fetal-like expression (68.1% of the 69 genes; Fig. 33K,L). Developmentally regulated genes included fetal-high *NEGR1* and *CADM1*, associated with neurite growth and synaptic adhesion, and infant-high *NRG1*, *KCNT1*, and *LRRTM4*, associated with postnatal synaptic development, membrane excitability, and synapse organization.^1425,1428,1484–1486^ Most of the overlapping genes were reduced in NEC, with 63 of 65 expressed at lower levels than in infant controls. *NCALD* and *UNC80*, which normally increased from fetal to infant, were reduced to fetal-like levels in NEC, consistent with lower calcium-sensing and NALCN-associated excitability.^1464–1465^ *PRKCA*, *GUCY1A2*, and *KIF13A* showed a similar fetal-like reduction, suggesting weaker PKC-dependent synaptic regulation, NO-cGMP signaling, and microtubule-dependent endosomal transport.^1446,1468,1487^ In contrast, fetal-high *DLGAP2* and *SSTR2* increased to fetal-like levels in NEC, indicating a shift toward fetal-associated postsynaptic and somatostatin-receptor expression. *SSTR2* is developmentally regulated in fetal neural tissues, where it appears early and changes across gestation, supporting a potential role in neural development and circuit maturation.^1488–1489^

NEC inhibitory neurons show immediate-early and nuclear RNA-processing responses alongside broad loss of genes supporting neuronal recognition, synaptic scaffolding, projection maintenance, vesicle cycling, receptor composition, excitability, calcium handling, NO-cGMP signaling, and intracellular signal integration. Initiation may be influenced by early disruption of inhibitory neuromuscular signaling, which could alter relaxation, transit, and bowel-wall coordination. During NEC progression, weakened inhibitory neurotransmission may promote dysmotility, stasis, and impaired reflex control in an already inflamed intestine. Developmental analysis reinforces this deficit, with several circuit-supporting genes at fetal-like levels or reduced beyond control states. Resolution may require restoration of inhibitory neuromuscular signaling, not only survival of neuronal structures.

### Excitatory Neurons

Following the broad inhibitory-neuronal disruption described above, we next examined excitatory enteric neurons. Excitatory enteric neurons participate in the contractile arm of enteric reflexes; excitatory motor neurons deliver cholinergic and tachykininergic signals to intestinal smooth muscle and coordinate propulsive contraction (Fig. 34A).^1490–1491^ Prior human NEC studies did not isolate excitatory neurons as a distinct population. Acute NEC reduced *CHAT* mRNA in bulk myenteric-plexus preparations alongside general neuronal loss, but *CHAT* recovered at stoma closure, whereas nNOS remained reduced.^1286^ These bulk-tissue, plexus-level, and histological studies did not resolve excitatory neurons as a distinct transcriptional population, leaving their specific transcriptional response undefined.

**Fig. 34.**
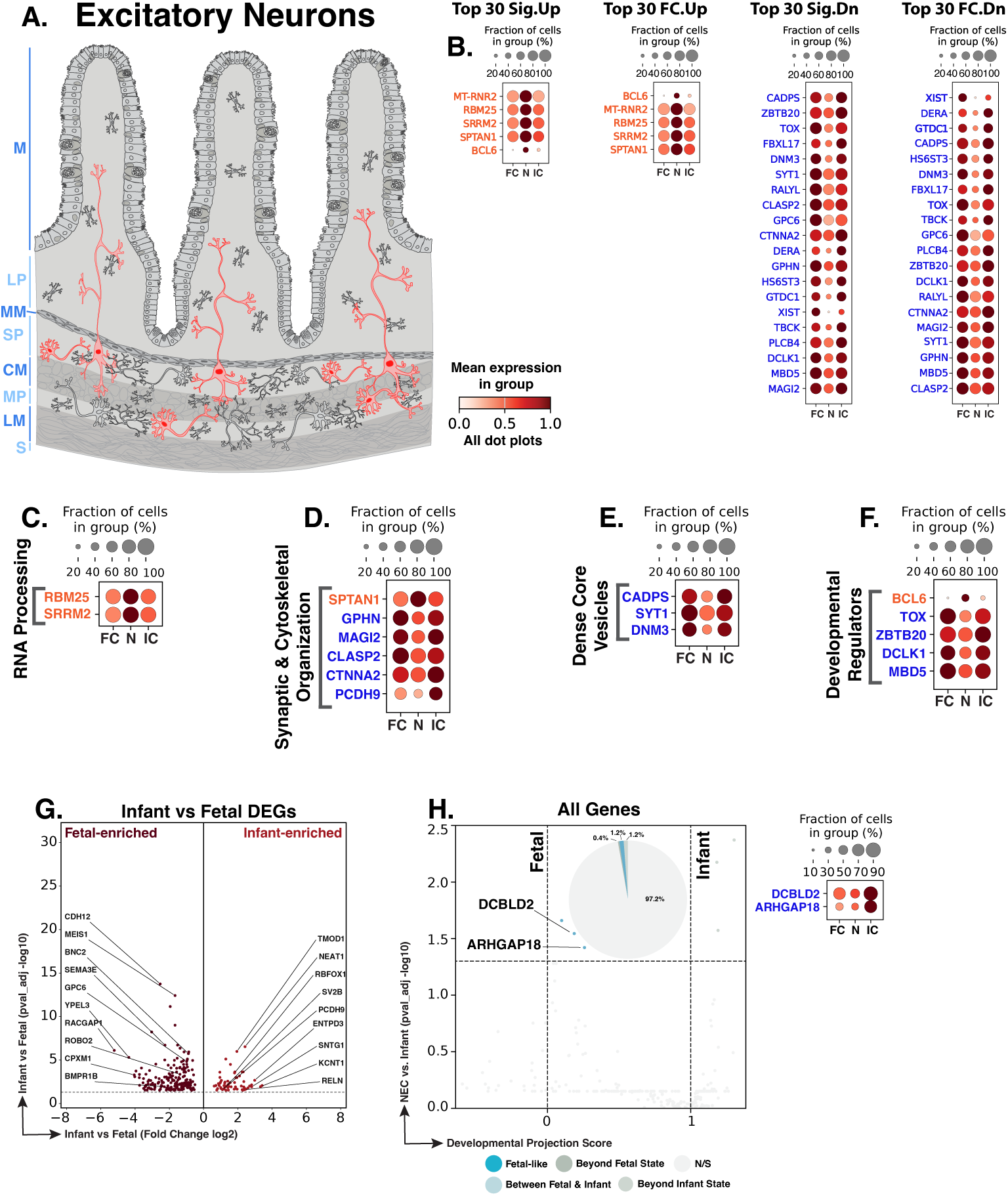
NEC excitatory enteric neurons show RNA-processing activation with reduced synaptic/cytoskeletal, dense-core-vesicle, developmental-regulatory, and maturity-associated programs. **A.** Schematic showing the distribution of excitatory enteric neurons (red) within the intestinal wall. Blue text to the left of the Panel A schematic indicates general anatomical regions: M, mucosa; LP, lamina propria; MM, muscularis mucosae; SP, submucosal plexus; CM, circular muscle; MP, myenteric plexus; LM, longitudinal muscle; S, serosa. **B.** Dot plots of top NEC-specific genes ranked by significance and fold change. **C–F.** Curated gene panels showing RNA-processing, synaptic/cytoskeletal remodeling, dense-core-vesicle, and developmental-regulatory programs. In panels **B–F**, orange gene text indicates upregulation in NEC relative to infant controls, whereas blue gene text indicates downregulation. **G.** Volcano plot of DEGs between fetal and infant controls that define the developmental trajectory for excitatory neurons. **H.** Gene-level maturity analysis of fetal-versus-infant DEGs, highlighting the subset also significantly changed in NEC relative to infant controls and positioned relative to fetal and infant reference states, and their corresponding dotplots. The developmental projection score in panel H is displayed on a symmetric logarithmic (symlog) scale. For expression dot plots, dot size indicates the fraction of nuclei expressing each gene and dot color indicates mean expression. In maturity plots, vertical dotted lines indicate the fetal and infant reference scores. Colored dots classify NEC-versus-infant DEGs as Fetal-like, Between Fetal & Infant, Beyond Fetal State, or Beyond Infant State; gray N/S dots are fetal-versus-infant DEGs without a significant NEC-versus-infant change.

We identified 677 excitatory neuronal nuclei across the atlas, including 480 fetal, 118 NEC, and 79 infant control nuclei. Differential-expression analysis detected 31 NEC-versus-infant DEGs in excitatory neurons, nearly ninefold fewer than the 276 DEGs in inhibitory neurons despite comparable NEC-versus-infant representation of the two neuronal populations (Fig. 34B). This difference reflects a substantially narrower transcriptional disturbance in excitatory neurons. Within this limited response, the affected genes were concentrated in RNA processing and synaptic structure, with postsynaptic density representing the only significantly depleted GO-term, consistent with a small but focused neuronal reaction to the NEC environment.

Among the few genes upregulated in NEC excitatory neurons, *RBM25* and *SRRM2* both participate in RNA processing. Their increased expression is consistent with greater engagement of alternative-splicing machinery: RBM25 promotes the inclusion of alternatively spliced exons, whereas SRRM2 organizes nuclear-speck-associated splicing condensates and regulates alternative splicing (Fig. 34C).^1476,1492^ These increases indicate a limited shift in transcript processing, contrasting with the broader activation of inflammatory and cellular-stress pathways in inhibitory neurons.

The strongest reductions involved genes that organize and stabilize postsynaptic sites (Fig. 34D). Lower *GPHN* may weaken inhibitory synapses onto excitatory neurons because gephyrin controls postsynaptic clustering and membrane dynamics of GABA_A_ receptors, while reduced *MAGI2* may destabilize receptor- and adhesion-associated postsynaptic complexes.^1493–1494^ Other reductions affected synaptic structure: *CTNNA2* links cadherin adhesion to the actin cytoskeleton and stabilizes dendritic spines, *PCDH9* contributes to glutamatergic-synapse organization, and *CLASP2* supports microtubule organization, neuronal polarity, neurite growth, and synaptic function.^1424,1430,1445^ In contrast, increased *SPTAN1* may help preserve cytoskeletal integrity because αII-spectrin supports axonal and dendritic organization, neuronal polarity, and synaptogenesis.^1495^ Overall, this pattern suggests partial structural compensation despite weaker postsynaptic organization and reduced integration of excitatory neurons within enteric circuits.

Molecular support for presynaptic vesicle secretion and recycling was also reduced (Fig. 34E). *CADPS*/CAPS1 promotes dense-core-vesicle exocytosis at a calcium-dependent prefusion step, and *SYT1* couples calcium sensing to synaptic-vesicle exocytosis and endocytosis.^1448,1496^ Decreased *DNM3* completes this sequence at the vesicle-retrieval stage because dynamin 3 has a major presynaptic role and cooperates with other dynamin isoforms during activity-dependent synaptic-vesicle endocytosis.^1378^ Coordinated reduction of *CADPS*, *SYT1*, and *DNM3* could hinder vesicle priming, calcium-triggered release, and membrane recycling. These vesicle-cycle deficits occurred together with reduced postsynaptic scaffolding and adhesion genes, weakening the molecular basis for stable synaptic communication.

Transcription-factor expression was directionally mixed across regulators of neural differentiation and neurite maintenance (Fig. 34F). Increased *BCL6*, which promotes neuronal differentiation through repression of the progenitor-maintaining Notch target *Hes5*, contrasted with decreased *TOX* and *ZBTB20*, which support neural-progenitor expansion and neurite growth or branching.^1497–1499^ Reduced *DCLK1* links these genes to microtubule-dependent axon elongation through phosphorylation of MAP7D1; *MBD5* was likewise reduced and regulates the balance between neural proliferation and differentiation.^1500–1501^ The opposing behavior of these genes indicates uneven regulation of neuronal differentiation and neurite maintenance.

Of the 252 genes distinguishing fetal from infant control excitatory neurons, 7 were also dysregulated in NEC versus infant controls, including 3 with fetal-like expression (42.9% of the 7 genes; Fig. 34G,H). Developmentally regulated genes included fetal-high *GPC6* and infant-high *PCDH9* and *KCNT1*, associated with excitatory synapse formation, synaptic organization, and membrane excitability.^1424,1485,1502^ All seven NEC-associated developmental genes were downregulated. Among the genes reduced to fetal-like levels, *DCBLD2* encodes a receptor-like transmembrane protein linked to human developmental phenotypes, while *ARHGAP18* regulates RhoA-dependent actin organization, cell shape, and motility.^1503–1504^ Overall, NEC excitatory neurons largely retained an infant-like developmental profile, with only a small set of changes affecting RhoA-actin regulation, dendritic organization, and synapse formation or maintenance.

NEC excitatory neurons show a narrower transcriptional disturbance than inhibitory neurons. Initiation and progression may still be influenced by reduced postsynaptic organization, adhesion, microtubule stability, and vesicle release/recycling, which could weaken propulsion and circuit reliability. However, the limited DEG burden and broad preservation of the postnatal profile suggest that excitatory-neuron disruption is less extensive than the inhibitory-neuron deficit. This asymmetry supports a model in which loss of relaxation and inhibitory control is the dominant neuronal abnormality, with a marginal excitatory synaptic deficit adding to dysmotility. Resolution may depend on recovery of balanced inhibitory and excitatory signaling across the enteric motor circuit.

### Smooth Muscle Compartment

The intestinal smooth muscle compartment, marked by *MYH11*, *ACTA2*/α-SMA, and *TAGLN* (Fig. 2), generates the contractile force required for intestinal propulsion and mixing and serves as the contractile effector downstream of enteric neural and pacemaker signaling.^1505–1506^ These functions are particularly relevant to NEC, where intestinal dysmotility can precede overt disease; in experimental NEC, intestinal motility is reduced before the onset of inflammation, and pharmacologic restoration of motility reduces disease severity.^24^ Human data also support involvement of intestinal muscle layers in severe NEC, with circulating smooth muscle actin detected in infants with severe surgical NEC.^1507^ Despite this, cell-type-resolved smooth muscle responses in NEC remain poorly defined. A published human NEC single-cell atlas did not report smooth muscle as a distinct compartment, while experimental neonatal mouse NEC multi-omics identified a reduced smooth muscle cell compartment and substantial NEC-associated changes in smooth muscle chromatin accessibility but did not resolve anatomically distinct muscularis populations.^18,157^

To define the smooth muscle populations in our study, heatmap-based annotation resolved three anatomically distinct smooth muscle populations: *RSPO2*+/*NPNT*+ muscularis mucosae, *MYOCD*+ inner muscularis, and *CAPN3*+ outer muscularis (Fig. 35). These populations separate the epithelial-adjacent muscularis mucosae from the deeper circular and longitudinal muscle layers of the muscularis externa. This organization allowed NEC-associated smooth muscle changes to be examined according to anatomical position, preserving regional distinctions that would be obscured by treating smooth muscle as a single broad compartment.

**Fig. 35.**
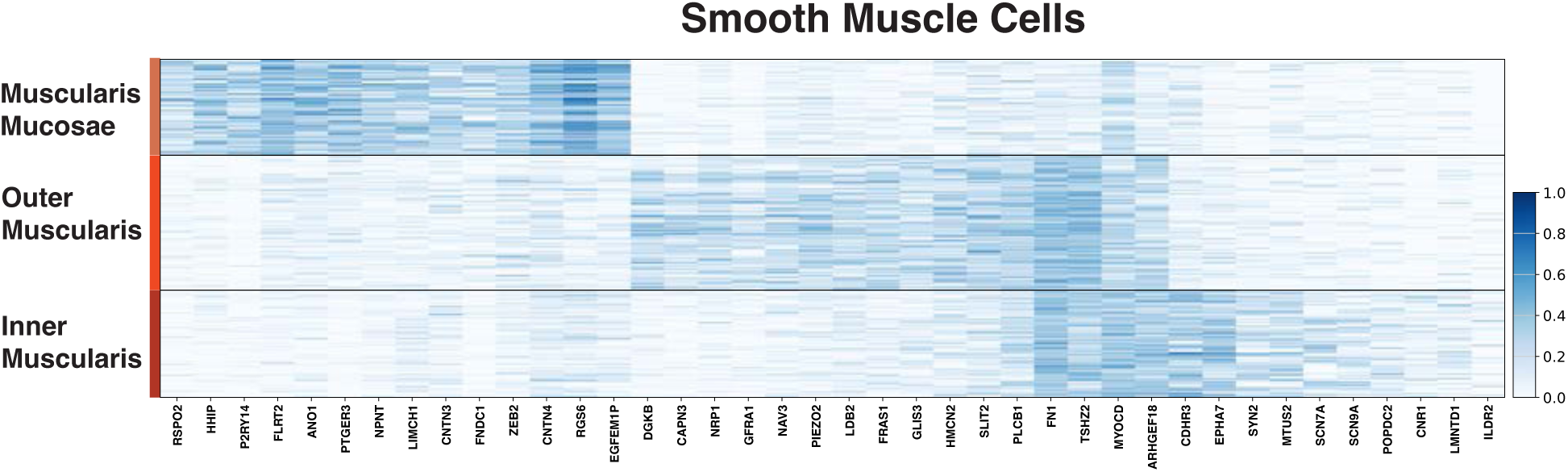
Smooth-muscle cell-type signatures define muscularis mucosae, outer muscularis, and inner muscularis populations. Heatmap showing marker-gene expression across muscularis mucosae, outer muscularis, and inner muscularis smooth-muscle populations. Distinct transcriptional signatures support smooth-muscle cell-type assignments and provide the framework for the detailed smooth-muscle analyses that follow.

The following sections examine how NEC alters smooth muscle populations across the intestinal wall. The muscularis mucosae captures the epithelial-adjacent smooth muscle response, where repair and matrix signals may influence mucosal recovery and epithelial support. Inner muscularis captures the circular muscle layer, where inflammatory, hypoxia/angiogenesis, matrix-remodeling, and contractility-associated changes could affect constriction, mixing, and propulsion. Outer muscularis captures the longitudinal muscle layer and allows deeper bowel-wall remodeling and contractile identity to be evaluated within the muscularis externa. Together, these sections frame NEC smooth muscle as a contractile system that may participate in epithelial and vascular repair, but may also contribute to dysmotility when inflammation, matrix remodeling, stress responses, and immature contractile regulation disrupt coordinated contraction-relaxation function.

### Muscularis Mucosae

The muscularis mucosae is an epithelial-adjacent smooth muscle layer at the base of the mucosa, beneath the lamina propria, where it is positioned to influence crypt architecture and epithelial repair (Fig. 36A).^593,1508^ This layer has been implicated in intestinal stem-cell homeostasis and crypt repair, supporting the possibility that it provides epithelial-supportive signals after mucosal injury.^593,1508^ Direct NEC literature focused specifically on the muscularis mucosae is limited. A published human NEC transcriptomic atlas did not identify a smooth muscle population, and experimental neonatal mouse NEC multi-omics found a reduced smooth muscle cell compartment without resolving the muscularis mucosae as a separate smooth muscle population.^18,157^ Thus, it is unknown whether human NEC affects this epithelial-adjacent smooth muscle layer in ways that could influence mucosal repair and bowel-wall function.

**Fig. 36.**
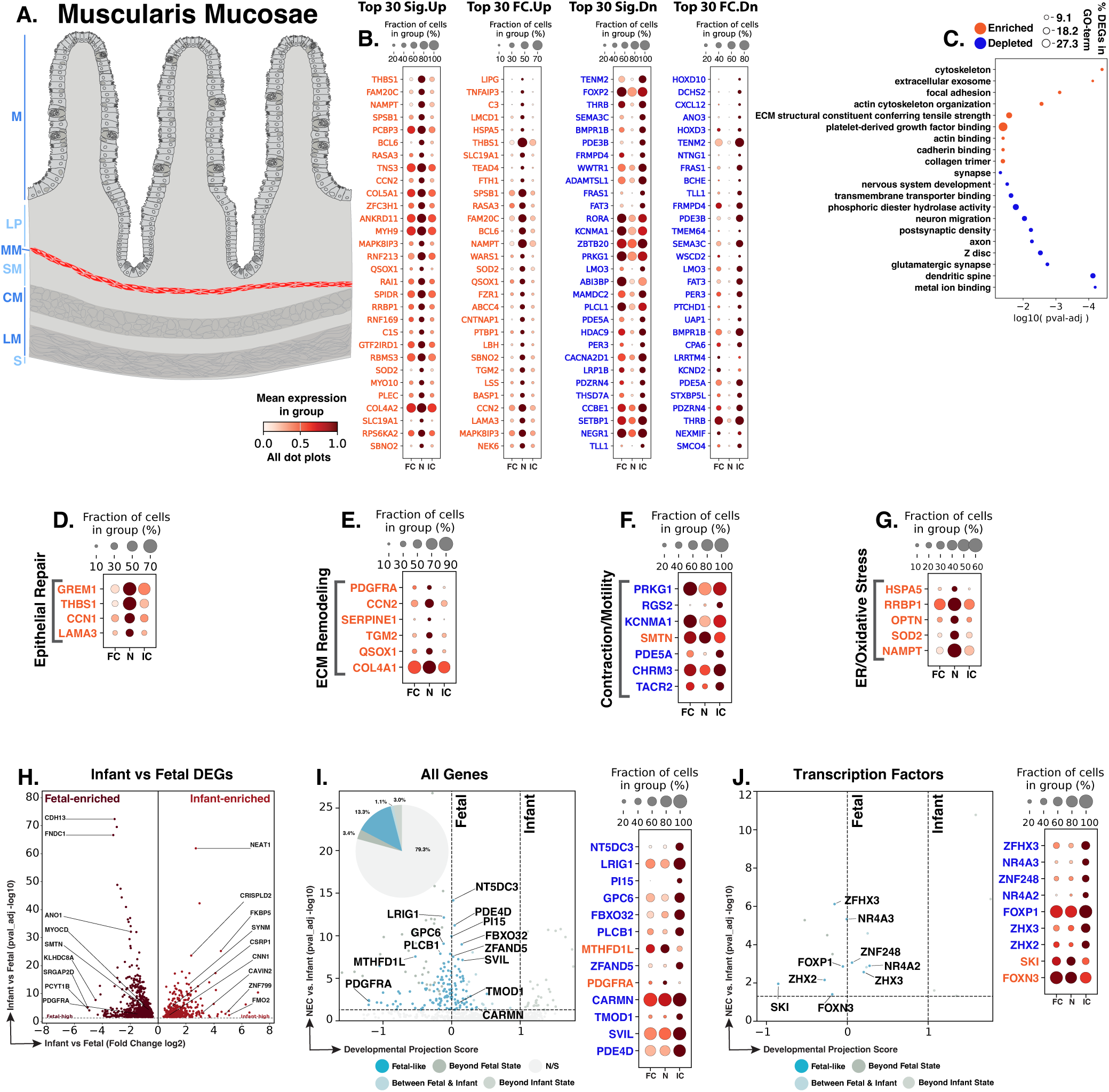
NEC muscularis mucosae shows epithelial-repair and matrix-remodeling activation with altered contractility, and oxidative/ER stress. **A.** Schematic showing the epithelial-adjacent location of the muscularis mucosae (red). Blue text to the left of the Panel A schematic indicates general anatomical regions: M, mucosa; LP, lamina propria; MM, muscularis mucosae; SM, submucosa; CM, circular muscle; MP, myenteric plexus; LM, longitudinal muscle; S, serosa. **B.** Dot plots of top NEC-specific genes ranked by significance and fold change. **C.** GO-term analysis showing enriched and depleted biological processes among NEC-associated DEGs. **D–G.** Curated gene panels showing epithelial-repair, ECM-remodeling/fibrosis, contractility/mobility, and ER/oxidative-stress programs. In panels **B** and **D–G**, orange gene text indicates upregulation in NEC relative to infant controls, whereas blue gene text indicates downregulation. **H.** Volcano plot of DEGs between fetal and infant controls that define the developmental trajectory for muscularis mucosae cells. **I–J.** Gene-level and transcription-factor maturity analyses of fetal-versus-infant DEGs, highlighting the subset also significantly changed in NEC relative to infant controls and positioned relative to fetal and infant reference states, and their corresponding dotplots. The developmental projection score in panels **I–J** is displayed on a symmetric logarithmic (symlog) scale. For expression dot plots, dot size indicates the fraction of nuclei expressing each gene and dot color indicates mean expression. In maturity plots, vertical dotted lines indicate the fetal and infant reference scores. Colored dots classify NEC-versus-infant DEGs as Fetal-like, Between Fetal & Infant, Beyond Fetal State, or Beyond Infant State; gray N/S dots are fetal-versus-infant DEGs without a significant NEC-versus-infant change.

We identified 1,051 muscularis mucosae nuclei across the atlas, with 446 genes significantly altered in NEC muscularis mucosae compared to infant controls, including 139 upregulated and 307 downregulated genes (Fig. 36B). GO-term enrichment analysis showed significant enrichment for platelet-derived growth factor binding, and terms related to extracellular matrix and cytoskeletal remodeling in NEC, along with depletion of phosphoric diester hydrolase activity, neuron migration, and nervous-system development terms (Fig. 36C). PDGF signaling can promote smooth muscle proliferation, and smooth muscle proliferation can alter the contractile phenotype.^1509–1511^ In addition, phosphodiesterase activity contributes to intestinal smooth muscle contractile regulation through cyclic nucleotide signaling.^1512^ Thus, these GO-term changes are consistent with altered contractile regulation in muscularis mucosae during NEC. These results were reflected in the extracellular matrix and contractility programs described below, while curated gene-level analysis also highlighted epithelial-repair and stress-adaptation programs.

NEC muscularis mucosae showed increased expression of genes associated with epithelial support and tissue repair (Fig. 36D). Higher *GREM1* is notable because GREM1 is a BMP antagonist expressed in the intestinal crypt niche, and smooth muscle in the muscularis mucosae contributes BMP antagonists that support intestinal stem-cell homeostasis.^593,1508^ *GREM1* expression has also been reported in human colonic muscularis mucosae, supporting its relevance to this epithelial-adjacent smooth muscle population.^699^ Increased *THBS1* and *CCN1* further pointed to mucosal wound repair and epithelial proliferation, while higher *LAMA3* could support epithelial adhesion and migration during repair.^679,735,1513^ Together, these changes suggest that NEC muscularis mucosae may contribute epithelial-supportive and repair-associated signals during mucosal injury.

Matrix-remodeling and fibrosis-associated genes were also increased in NEC muscularis mucosae (Fig. 36E). Higher *PDGFRA* may reflect greater PDGF-associated mesenchymal activation because PDGFRA supports fibroblast proliferation during wound healing.^674^ Increased *CCN2* can promote collagen deposition, while *SERPINE1*/PAI-1 can limit fibrinolytic matrix degradation and contribute to intestinal fibrosis.^681–682,713,1514^ Higher *TGM2* and *QSOX1* further supported extracellular matrix crosslinking, assembly, and remodeling.^1515–1518^ Overall, these changes suggest repair-associated matrix remodeling that could contribute to fibrosis if sustained or excessive.

Contractility-associated genes showed mixed changes in NEC muscularis mucosae (Fig. 36F). *PRKG1* and *RGS2*, which support smooth muscle relaxation, were reduced.^694–695,1519–1520^ *KCNMA1*, encoding the BK potassium channel, and *PDE5A*, which regulates cGMP-dependent relaxation, were also reduced.^692–693,732^ In contrast, *SMTN*, which encodes smoothelin A and supports intestinal smooth muscle contractility, was increased and may reflect compensatory contractile adaptation.^780^ Together, these changes suggest an altered balance between contraction and relaxation that could affect intestinal transit. Because impaired motility can contribute to NEC, altered muscularis mucosae contractility may influence disease progression.^24^

NEC muscularis mucosae also showed increased genes associated with protein-folding, oxidative, and metabolic stress (Fig. 36G). Higher *HSPA5*, *RRBP1*, and *OPTN* were consistent with ER protein folding, stress adaptation, and protection from ER-stress-associated injury.^410,1521–1523^ Increased *SOD2* supported mitochondrial antioxidant defense, while higher *NAMPT* pointed to NAD⁺-salvage metabolism during oxidative stress.^633,1524–1525^ These changes suggest that NEC muscularis mucosae engages adaptive survival responses under protein-folding, oxidative, and metabolic stress. Such responses may help preserve smooth muscle function during injury, although persistent stress could impair motility or epithelial-niche support.

Of the 1,042 genes distinguishing fetal from infant control muscularis mucosae, 216 were also dysregulated in NEC versus infant controls, including 139 with fetal-like expression (64.4% of the 216 genes; Fig. 36H,I). Developmentally regulated genes included fetal-high *PDGFRA*, a marker of digestive mesenchymal progenitors that give rise to muscularis mucosae smooth muscle after birth, and infant-high *CNN1* and *SYNM*, associated with differentiated smooth muscle identity.^593,1526–1528^ This pattern indicates that most developmental genes remained infant-like, while a substantial subset showed fetal-like expression in NEC.

The fetal-like genes highlighted reduced contractile and cytoskeletal programs (Fig. 36I). The mesenchymal progenitor marker *PDGFRA* was significantly upregulated in NEC and had fetal-like expression.^593,1526^ By contrast, *CARMN*, a smooth muscle-enriched lncRNA required for gastrointestinal smooth muscle contractile gene expression and function; *TMOD1*, an actin-filament pointed-end capping protein found in intestinal smooth muscle cytoskeletal structures; and *SVIL*, a cytoskeletal protein linked to maintenance of the contractile smooth muscle phenotype, were significantly downregulated in NEC and had fetal-like expression.^1529–1533^ These changes suggest that NEC muscularis mucosae cells might retain progenitor-associated features while losing postnatal contractile and cytoskeletal features, which could impair mucosal mechanical support.

At the transcription factor level, NEC muscularis mucosae showed mixed regulation of proliferation and contractile maturation. *NR4A3* deficiency can preserve the vascular smooth muscle contractile phenotype during calcification and was downregulated in NEC with fetal-like expression (Fig. 36J).^1534^ *SKI* inhibits vascular smooth muscle cell proliferation and was upregulated in NEC with fetal-like expression.^1535^ *FOXP1* modulates vascular smooth muscle cell proliferation, and *ZHX2* can inhibit portal-vein smooth muscle cell proliferation; these two genes were downregulated in NEC with expression similar to fetal control.^924,1536^ The expression of these transcription factors suggests altered regulation of genes involved in smooth muscle proliferation and contractile regulation.

NEC muscularis mucosae occupies a mixed epithelial-supportive, matrix-remodeling, contractility-altered, and stress-adapted state. Initiation may be influenced by altered epithelial-adjacent mechanical support and changes in contraction-relaxation signaling near the mucosa. During progression, persistent matrix remodeling, ER/oxidative stress, and immature contractile features could disrupt mucosal support, crypt architecture, and local motility. At the same time, increased epithelial-niche and wound-repair genes suggest that this layer may contribute to mucosal recovery. NEC resolution may depend on whether muscularis mucosae repair signals restore epithelial support without maintaining maladaptive matrix remodeling or immature contractile regulation.

### Inner Muscularis

The inner muscularis is the inner circular smooth muscle layer of the muscularis externa, where circumferential contraction contributes to intestinal constriction and propulsion and mixing (Fig. 37A).^1537–1538^ Human intestinal single-cell and spatial analyses have identified transcriptionally distinct inner muscularis populations, including inner-muscularis expression of *PNCK*, *MORN5*, and *AHNAK2*.^48^ Consistent with this annotation, our inner muscularis cluster shows higher *MYOCD* and a greater fraction of cells expressing *PNCK*, *MORN5*, and *AHNAK2*, together with canonical contractile smooth muscle genes including *ACTG2*, *MYH11*, *TAGLN*, and *DES*.^48,593,1539–1540^

**Fig. 37.**
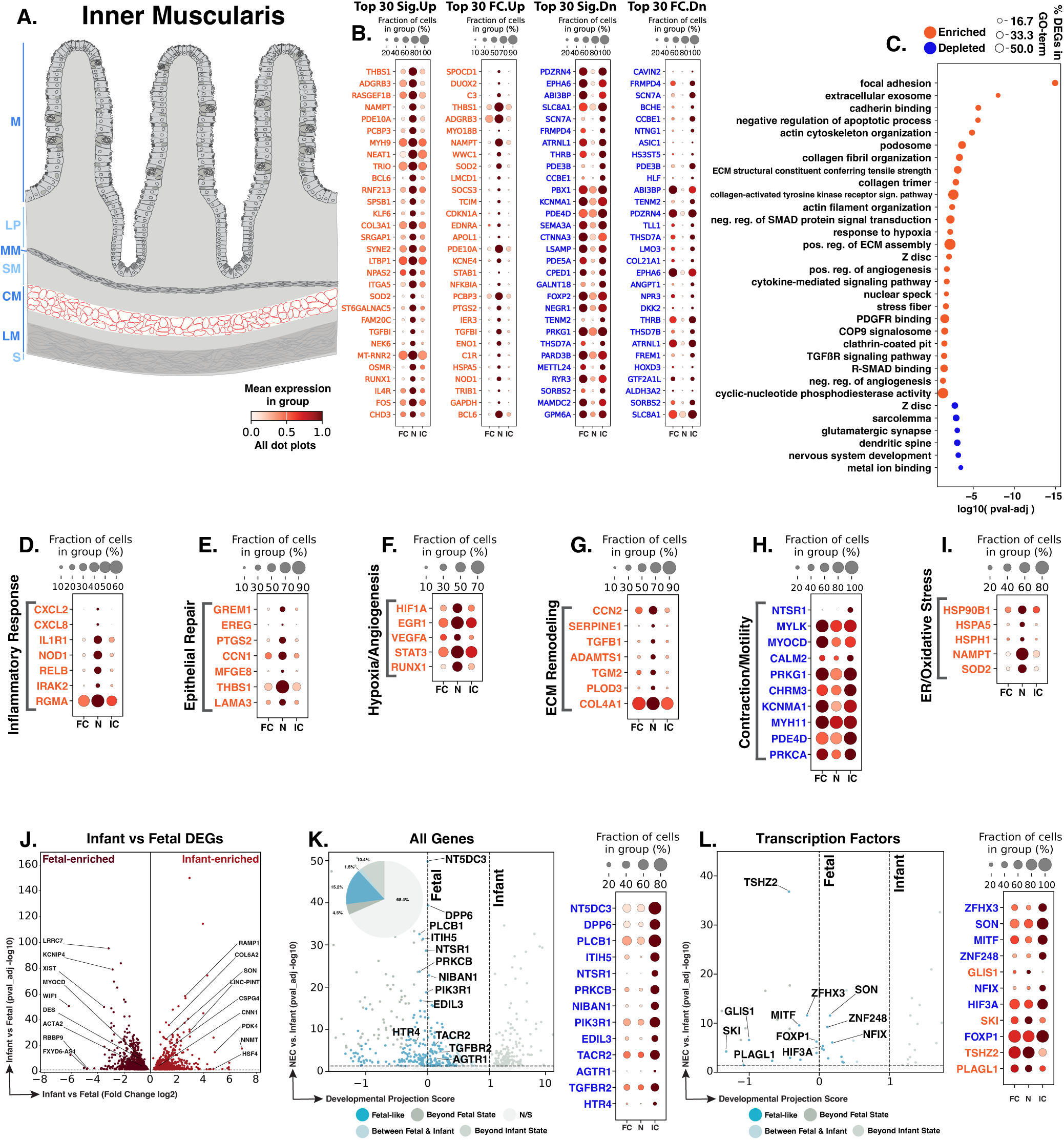
NEC inner muscularis shows inflammatory, epithelial-repair, hypoxia/angiogenesis, extracellular-matrix, and neuromuscular-associated changes. **A.** Schematic showing the location of the inner muscularis (red) within the intestinal muscular wall. Blue text to the left of the Panel A schematic indicates general anatomical regions: M, mucosa; LP, lamina propria; MM, muscularis mucosae; SM, submucosa; CM, circular muscle; MP, myenteric plexus; LM, longitudinal muscle; S, serosa. **B.** Dot plots of top NEC-specific genes ranked by significance and fold change. **C.** GO-term analysis showing enriched and depleted biological processes among NEC-associated DEGs. **D–I.** Curated gene panels showing inflammatory, epithelial-repair, hypoxia/angiogenesis, extracellular-matrix, neuromuscular/contractile-signaling, and ER/oxidative-stress programs. In panels **B** and **D–I**, orange gene text indicates upregulation in NEC relative to infant controls, whereas blue gene text indicates downregulation. **J.** Volcano plot of DEGs between fetal and infant controls that define the developmental trajectory for inner muscularis cells. **K–L.** Gene-level and transcription-factor maturity analyses of fetal-versus-infant DEGs, highlighting the subset also significantly changed in NEC relative to infant controls and positioned relative to fetal and infant reference states, and their corresponding dotplots. The developmental projection score in panels **K–L** is displayed on a symmetric logarithmic (symlog) scale. For expression dot plots, dot size indicates the fraction of nuclei expressing each gene and dot color indicates mean expression. In maturity plots, vertical dotted lines indicate the fetal and infant reference scores. Colored dots classify NEC-versus-infant DEGs as Fetal-like, Between Fetal & Infant, Beyond Fetal State, or Beyond Infant State; gray N/S dots are fetal-versus-infant DEGs without a significant NEC-versus-infant change.

The inner circular smooth muscle layer functions within neuromuscular networks that include interstitial cells of Cajal, PDGFRA+ cells, and enteric neurons to coordinate intestinal motility, and intestinal smooth muscle has also been shown to contribute to enteric nervous system patterning during intestinal development.^709,1541–1543^ Direct NEC literature focused specifically on the inner muscularis is limited; however, a published human NEC single-cell transcriptomic atlas did not resolve smooth muscle-specific injury programs, while experimental neonatal mouse NEC multi-omics found a reduced smooth muscle cell compartment but did not examine the inner muscularis as a separate mechanistic population.^18,157^ Thus, it is unknown how NEC might affect this human inner circular smooth muscle population in a way that could affect motility and bowel-wall repair.

We identified 1,175 inner muscularis nuclei across the atlas. Differential-expression analysis identified 1,302 genes significantly changed in NEC relative to infant controls, including 478 upregulated and 824 downregulated genes (Fig. 37B). GO-term enrichment showed increased inflammatory/cytokine signaling, hypoxia-angiogenesis, TGF-β signaling, extracellular matrix remodeling, and cytoskeletal/focal adhesion programs, with reduced contractile programs (Fig. 37C). The COP9 signalosome was also enriched; its subunit *CSN6* can promote PDGF-induced smooth muscle cell proliferation.^1544^ Collagen-activated tyrosine kinase receptor signaling was likewise enriched, and *DDR1* signaling has been shown to promote smooth muscle migration and collagen remodeling.^1545–1546^ Overall, these GO-terms suggest increased proliferative, migratory, and matrix-remodeling activity in NEC inner muscularis. These patterns were reflected in the inflammatory, hypoxia-angiogenesis, matrix, and contractility programs described below, while gene-level analysis also identified an epithelial-repair response not captured by the summarized GO-terms.

NEC inner muscularis showed increased inflammatory and innate immune signaling that could extend inflammatory activity into the muscularis externa (Fig. 37D). Higher *CXCL2* and *CXCL8* supported neutrophil recruitment; notably, intestinal smooth muscle cells from Crohn’s disease tissue can express and secrete CXCL8/IL-8.^1547–1549^ Increased *IL1R1* suggested greater responsiveness to IL-1, while higher *NOD1* and *RELB* pointed to innate immune and NF-κB-associated signaling.^597,1550–1552^ Higher *RGMA* may also reflect inflammatory phenotypic change, as RGMa has been linked to macrophage-like features in vascular smooth muscle.^1553^ Taken together, these changes suggest that NEC inner muscularis can respond to microbial or injury-associated inflammatory signals and may contribute to inflammatory activity within contractile tissue.

NEC inner muscularis also showed increased epithelial-support and repair-associated genes (Fig. 37E). Higher *GREM1* is notable because smooth muscle-derived BMP antagonists support intestinal epithelial repair and the stem-cell niche.^593,1508^ Increased *EREG*, *PTGS2*, and *CCN1* further pointed to epithelial proliferation and mucosal wound repair, while higher *MFGE8* and *THBS1* have also been linked to mucosal healing.^615,618,679,735,802,1554–1555^ These changes suggest that NEC inner muscularis may provide epithelial-supportive signals during attempted mucosal repair.

Hypoxia- and angiogenesis-associated genes were also increased in NEC inner muscularis (Fig. 37F). Higher *HIF1A* and *EGR1* were consistent with a hypoxic response, while increased *VEGFA*, *STAT3*, and *RUNX1* pointed to angiogenic signaling.^62,241,605,721–722,1556–1557^ Together, these increases suggest that NEC inner muscularis responds to hypoxic and vascular stress. This response could support vascular repair and reoxygenation, although persistent activation may indicate unresolved tissue hypoxia.

Extracellular-matrix and fibrosis-associated genes were also increased (Fig. 37G). Higher *CCN2*, *SERPINE1*, and *TGFB1* supported collagen accumulation and profibrotic signaling, while *ADAMTS1*, *TGM2*, and *PLOD3* implicated matrix modification, crosslinking, and collagen processing.^681–682,713,1514,1558–1564^ This matrix response may help stabilize injured tissue during repair, but if sustained could contribute to tissue stiffening and impaired motility.

Contractility-associated genes were reduced in NEC inner muscularis (Fig. 37H). Lower *NTSR1*, *MYLK*, *MYOCD*, and *CALM2* suggested weaker neurotensin-responsive, calcium-dependent, and smooth muscle contractile signaling.^1505,1565–1569^ These changes suggest reduced contractile and motility-associated capacity. Because impaired intestinal motility has been linked to NEC, lower contractile signaling could contribute to altered intestinal transit and mechanical function.^24^

Cellular-stress genes were also increased in NEC inner muscularis (Fig. 37I). Higher *HSP90B1* and *HSPA5* were consistent with ER protein-folding stress, while increased *SOD2* supported a mitochondrial antioxidant response.^629–633^ These changes suggest that inner muscularis cells engage adaptive stress responses during inflammatory, hypoxic, and oxidative injury. Persistent stress, however, could impair smooth muscle function and intestinal motility.

Of the 1,572 genes distinguishing fetal from infant control inner muscularis, 496 were also dysregulated in NEC versus infant controls, including 239 with fetal-like expression (48.2% of the 496 genes; Fig. 37J,K). Developmentally regulated genes included fetal-high *ACTA2* and infant-high *CNN1*, both associated with smooth muscle differentiation.^1570–1571^ Fetal-like genes highlighted neurohumoral and contractile regulation. *TACR2* and *HTR4*, which mediate tachykinin-induced contraction and serotonin-dependent relaxation in human colonic smooth muscle, were both reduced to fetal-like levels in NEC.^1572–1573^ *TGFBR2*, which supports smooth muscle contractile gene expression and postnatal homeostasis, was also reduced to fetal-like expression.^1574–1575^ These changes suggest that NEC inner muscularis retains fetal-like features in pathways that regulate circular-muscle contraction and relaxation.

At the transcription factor level, NEC inner muscularis also showed fetal-like changes in regulators of smooth muscle proliferation (Fig. 37L). *SKI*, *PLAGL1*, and *TSHZ2*, which have been associated with restraint of cell proliferation, were increased with fetal-like expression, whereas *FOXP1*, which can promote vascular smooth muscle proliferation, was reduced to fetal-like expression.^924,1535,1576–1577^ This pattern suggests that NEC inner muscularis shows fetal-like transcriptional regulation of smooth muscle proliferative control.

NEC inner muscularis acquires inflammatory, epithelial-repair, hypoxia-angiogenic, cellular-stress, and extracellular matrix remodeling features while showing reduced contractility-associated and neurohumoral-response signaling. Initiation may be influenced by impaired circular-muscle coordination, altered constriction, and disrupted transit. As the disease progresses, inflammatory and matrix-remodeling activity may extend into the muscularis externa while reduced contractile and neurohumoral signaling compromises mixing, propulsion, and mechanical coordination. Epithelial repair and angiogenic genes may support recovery by restoring mucosal and vascular function. Resolution may fail if repair signals remain coupled to inflammation, matrix remodeling, and impaired circular-muscle contractility.

### Outer Muscularis

The outer muscularis is the longitudinal smooth muscle layer of the muscularis externa, located outside the inner circular layer and contributing to intestinal propulsion and motility (Fig. 38A).^1537^ Human single-cell and spatial studies have identified a distinct outer muscularis population marked by *BCHE*, *CAPN3*, and *IGFBP5*.^48^ Consistent with this annotation, our outer muscularis cluster expressed *BCHE* and *CAPN3* together with canonical smooth muscle genes including *ACTG2*, *MYH11*, *TAGLN*, and *DES*.^48,593,1539–1540,1578^ The muscularis externa works with interstitial cells of Cajal, PDGFRA⁺ cells, and the enteric nervous system to coordinate smooth muscle contraction, intestinal propulsion, and bowel-wall mechanics.^709,1539,1541^ Direct NEC studies of the outer muscularis are limited. Prior human NEC single-cell work did not resolve smooth muscle injury programs, and experimental neonatal mouse NEC multi-omics identified a reduced smooth muscle compartment without examining the outer muscularis separately.^18,157^ Thus, how human NEC affects this outer contractile compartment and whether these changes contribute to dysmotility remain unclear.

**Fig. 38.**
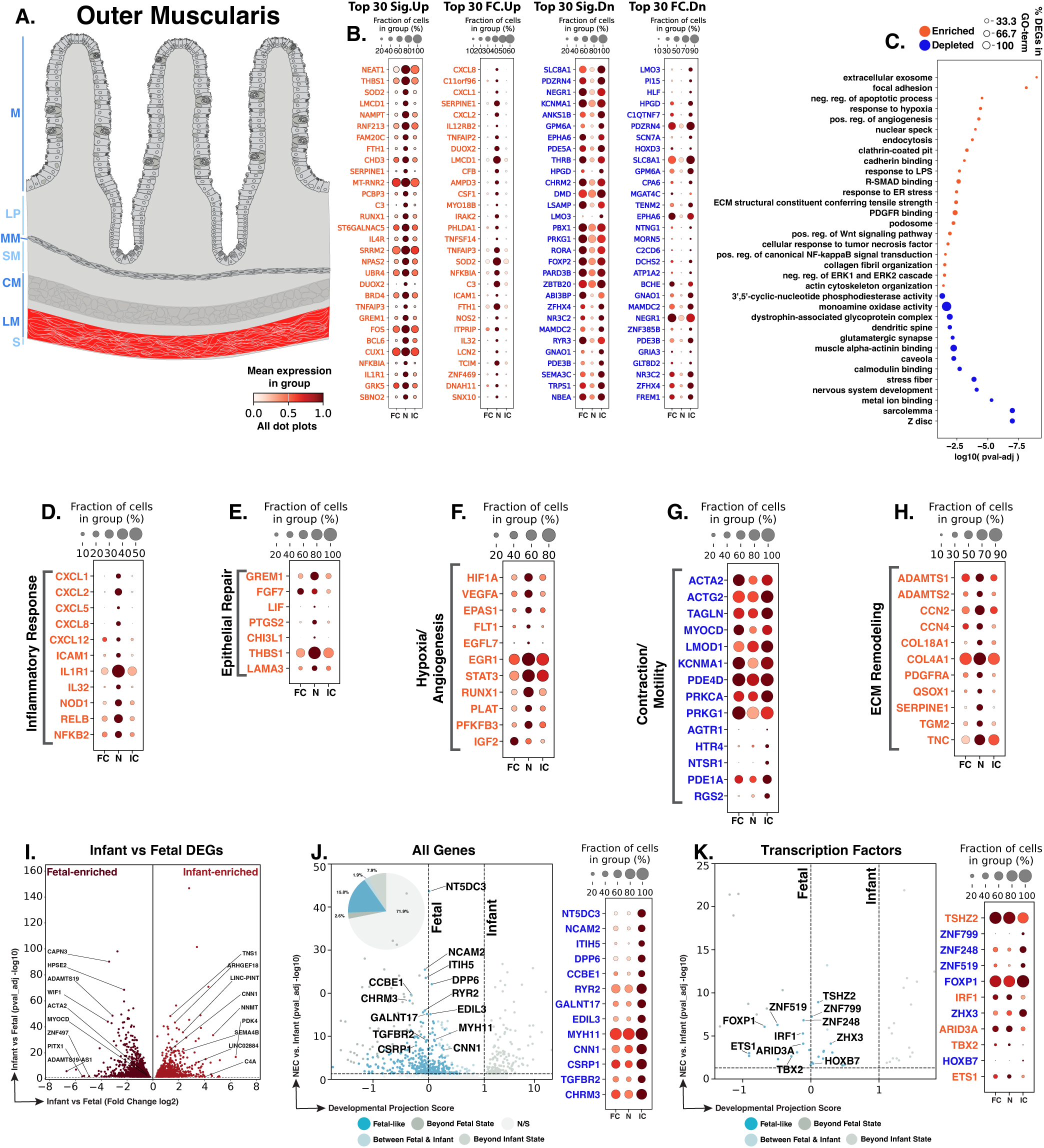
NEC outer muscularis cells show inflammatory and repair-associated activation with hypoxia/angiogenesis and reduced contractile programs. **A.** Schematic showing the location of the outer muscularis (red) within the intestinal muscular wall. Blue text to the left of the Panel A schematic indicates general anatomical regions: M, mucosa; LP, lamina propria; MM, muscularis mucosae; SM, submucosa; CM, circular muscle; MP, myenteric plexus; LM, longitudinal muscle; S, serosa. **B.** Dot plots of top NEC-specific genes ranked by significance and fold change. **C.** GO-term analysis showing enriched and depleted biological processes among NEC-associated DEGs. **D–H.** Curated gene panels showing inflammatory-response, epithelial-repair, hypoxia/angiogenesis, contraction/motility, and extracellular-matrix programs. In panels **B** and **D–H**, orange gene text indicates upregulation in NEC relative to infant controls, whereas blue gene text indicates downregulation. **I.** Volcano plot of DEGs between fetal and infant controls that define the developmental trajectory for outer muscularis cells. **J–K.** Gene-level and transcription-factor maturity analyses of fetal-versus-infant DEGs, highlighting the subset also significantly changed in NEC relative to infant controls and positioned relative to fetal and infant reference states, and their corresponding dotplots. The developmental projection score in panels **J–K** is displayed on a symmetric logarithmic (symlog) scale. For expression dot plots, dot size indicates the fraction of nuclei expressing each gene and dot color indicates mean expression. In maturity plots, vertical dotted lines indicate the fetal and infant reference scores. Colored dots classify NEC-versus-infant DEGs as Fetal-like, Between Fetal & Infant, Beyond Fetal State, or Beyond Infant State; gray N/S dots are fetal-versus-infant DEGs without a significant NEC-versus-infant change.

We identified 1,478 outer muscularis nuclei across the atlas. Differential-expression analysis identified 1,681 genes significantly changed in NEC relative to infant controls, including 611 upregulated and 1,070 downregulated genes (Fig. 38B). GO-term enrichment showed increased inflammatory/LPS-NF-κB, hypoxia-angiogenesis, focal adhesion/extracellular matrix, and cytoskeletal programs, with reduced calmodulin/cyclic nucleotide and α-actinin/Z-disc-associated contractile programs (Fig. 38C). Calmodulin-dependent MLCK signaling and α-actinin both support smooth muscle contraction, suggesting altered contractile capacity in NEC outer muscularis.^712,1505,1579–1580^ These GO-term patterns were reflected in the inflammatory, hypoxia-angiogenesis, matrix, and contractility responses described below, while gene-level analysis also identified an epithelial-repair response not fully captured by the summarized GO-terms.

Inflammatory and NF-κB-associated genes were increased in NEC outer muscularis (Fig. 38D). Higher *CXCL1*, *CXCL2*, *CXCL5*, and *CXCL8* supported neutrophil recruitment, while *CXCL12* implicated lymphocyte chemotaxis.^1581–1584^ Intestinal smooth muscle cells from Crohn’s disease tissue can also secrete CXCL8/IL-8, supporting a direct role for smooth muscle in inflammatory chemokine production.^1549^ Increased *ICAM1* further supported leukocyte adhesion and trafficking, while higher *IL1R1*, *IL32*, and *NOD1* pointed to greater responsiveness to inflammatory and microbial signals.^597,853,899,1585–1586^ Overall, NEC outer muscularis may contribute inflammatory signaling and immune cell recruitment in addition to its contractile role.

Epithelial-support and repair-associated genes were also increased (Fig. 38E). Higher *GREM1* is notable because smooth muscle-derived BMP antagonists contribute to intestinal stem-cell maintenance and epithelial repair.^593^ Increased *FGF7* further suggested mesenchymal support of epithelial proliferation, while higher *LIF*, *PTGS2*, and *CHI3L1* implicated stem-cell maintenance, epithelial regeneration, and survival during injury.^620,623,736,761,802,1587–1588^ Increased *THBS1* added another mucosal wound-repair signal.^679,1589^ These changes suggest that NEC outer muscularis may provide epithelial-supportive signals during attempted mucosal repair, even as its contractile programs are reduced.

Hypoxia- and angiogenesis-associated genes were also increased in NEC outer muscularis (Fig. 38F). Higher *HIF1A* reflected a hypoxic response, while increased *VEGFA*, *EPAS1*, and *EGFL7* supported angiogenic signaling and endothelial growth.^62,603,855,1052,1556–1557,1590–1591^ This pattern is consistent with hypoxic or vascular stress and may support vascular repair and reoxygenation, although persistent activation could reflect unresolved tissue hypoxia.

Contractile smooth muscle genes were broadly reduced in NEC outer muscularis (Fig. 38G). Lower *ACTA2* and *ACTG2* pointed to reduced actin-based contractile machinery, while decreased *TAGLN*, *MYLK*, *MYOCD*, and *LMOD1* implicated smooth muscle differentiation, myosin activation, contractile gene regulation, and force generation.^774–775,1505,1566,1578,1592–1593^ These changes suggest reduced contractile capacity in NEC outer muscularis. Because impaired intestinal motility has been implicated experimentally in NEC, loss of contractile machinery could contribute to altered intestinal transit.^24^

Extracellular-matrix and basement membrane genes were also increased (Fig. 38H). Higher *ADAMTS2*, *COL18A1*, and *COL4A1* pointed to collagen processing and basement membrane organization.^764–766,768^ Increased *CCN4* and *PDGFRA* further implicated cell-matrix adhesion, migration, and smooth muscle proliferation.^1594–1595^ These changes suggest active matrix remodeling that could help restore bowel-wall structure after injury but, if persistent, could disrupt smooth muscle organization and contribute to maladaptive remodeling.

Of the 2,158 genes distinguishing fetal from infant control outer muscularis, 607 were also dysregulated in NEC versus infant controls, including 341 with fetal-like expression (56.2% of the 607 genes; Fig. 38I,J). Developmentally regulated genes included fetal-high *ACTA2* and infant-high *CNN1*, both associated with human mesenchymal-to-smooth muscle differentiation.^1570^ Fetal-like genes highlighted reduced contractile and regulatory functions. *MYH11*, *CNN1*, and *TGFBR2*, which support smooth muscle contraction, contractile regulation, differentiation, and postnatal homeostasis, were all reduced to fetal-like levels in NEC.^689–691,776,1575,1596^ *CHRM3*, which mediates contraction of ileal longitudinal smooth muscle, was also reduced with fetal-like expression.^1597–1598^ Overall, NEC outer muscularis retained substantial fetal-like features involving contractile and smooth muscle regulatory genes that could contribute to impaired intestinal motility.^24^

At the transcription factor level, NEC outer muscularis showed fetal-like changes in regulators of smooth muscle differentiation (Fig. 38K). Experimental studies link *TBX2* and *Tbx3* to smooth muscle differentiation, while *HOXB7* can promote a smooth muscle-like phenotype and *ETS1* can repress smooth muscle differentiation genes.^1599–1601^ Fetal-like expression of *TBX2*, *HOXB7*, and *ETS1* therefore suggests altered transcriptional regulation of smooth muscle differentiation in NEC.

NEC outer muscularis shows inflammatory, epithelial-repair, hypoxia-angiogenic, and matrix-remodeling features together with reduced contractile machinery resembling an immature state. Initiation may be influenced by weakened longitudinal-muscle function and altered bowel-wall mechanics. During progression, inflammatory chemokine signaling, matrix remodeling, and hypoxia-associated vascular stress could extend injury through the muscularis externa while reduced smooth muscle identity impairs propulsion and coordinated motility. Repair and angiogenic signals may support recovery if they restore bowel-wall structure and vascular supply. Failed resolution may occur if inflammation and matrix remodeling persist while contractile programs remain immature or suppressed.

## Discussion

This study provides a full-intestine, cell-type-resolved view of human NEC. The atlas defines NEC not as a single pathway or injured cell type, but as a coordinated multicellular disease state linking epithelial maturation failure, regulated cell-death susceptibility, stromal-vascular repair responses, impaired immune clearance, enteric neuroglial disruption, and altered smooth muscle contractile signaling. Across regulated cell-death pathways, NEC-associated transcription reflected concurrent priming for death and protection, suggesting that cell fate may depend on whether death-promoting or protective programs predominate. These programs may therefore offer therapeutic targets for shifting the balance toward cell survival. Across cell types, NEC showed tissue injury coupled to incomplete compensatory responses. Epithelial proliferation, mucus-barrier reinforcement, angiogenic signaling, matrix repair, trophic support, and inflammatory feedback suggest that many compartments attempt to adapt to injury, yet these responses remain coupled to incomplete maturation, persistent stress, or loss of specialized function. The central implication is that NEC progression and recovery may depend on whether epithelial repair, vascular repair, immune clearance, and restoration of intestinal motility proceed in a coordinated manner, or whether persistent inflammation, developmental immaturity, and cellular stress sustain dysfunction across these compartments.

Several limitations should guide interpretation. NEC tissue obtained at resection captures advanced disease and cannot establish where each sampled region falls relative to fully necrotic tissue, less affected margins, or adjacent transitional zones. It also cannot determine when each transcriptional program began or whether it contributed to injury, arose in response to injury, or reflected incomplete recovery. While single-nucleus RNA sequencing enabled analysis of rare frozen transmural NEC specimens, RNA data do not directly measure protein abundance, pathway activity, secretion, spatial relationships, or cellular function. Therefore, the biological interpretation of these inferred transcriptional programs will require spatial, protein-level, and functional validation. Rare populations such as BEST4+ cells, tuft cells, dendritic cells, and interstitial cells of Cajal, and heterogeneous groups such as T cells, macrophage, glia, and mixed neuronal populations, will require deeper sampling or targeted enrichment to resolve subtype-specific mechanisms. Together, these limitations underscore that the timing and causal roles proposed in this study remain hypotheses, but the atlas provides a roadmap for empirical testing by prioritizing the cell types, gene programs, and tissue interactions most likely to influence NEC initiation, progression, and resolution.

The resource also opens several opportunities for future investigation. First, the recurrent cross-lineage genes identified here could be tested in stool, blood, or other biospecimens as candidate multigene markers of evolving NEC risk, disease activity, or recovery. Second, this transcriptomic resource can guide spatial and protein-level validation to determine whether predicted programs occur in the same tissue neighborhoods and whether key proteins, glycans, matrix features, and pathway effectors are present. Third, ligand-receptor and cell-cell interaction analyses can use this atlas to prioritize candidate signaling axes that connect specific injured or adaptive cell states. This would allow future studies to ask, for example, whether epithelial stress programs are linked to stromal repair signals, whether macrophage inflammatory and clearance states influence vascular recovery, whether lymphatic dysfunction reflects altered endothelial-immune communication, and whether glial or neuronal injury signals converge on smooth muscle contractile programs. Several NEC-relevant pathways already support this type of cross-compartment testing, including VEGF/VEGFR2-dependent postnatal microvascular development, macrophage-derived IGF1 support of neonatal intestinal vasculature, GLP-2-dependent epithelial protection, and glia-dependent preservation of motility in experimental NEC.^24,524–525,609–610^ These examples illustrate how cell-type-resolved signals from the atlas could be tested as preventive or therapeutic targets that stabilize vulnerable compartments, limit disease amplification, or improve recovery after injury.

NEC may be best understood as a breakdown in coordinated recovery across the premature intestine. This NEC atlas shows that many cell types activate responses that could support defense or repair, but these responses often occur alongside stress, incomplete maturation, and loss of normal cellular function. This creates a framework for testing why some injury responses help restore tissue homeostasis, whereas others prolong inflammation, weaken barrier and vascular recovery, impair immune clearance, disrupt neuromuscular function, or prevent the intestine from rebuilding after injury.

## Methods

### Human subjects and cohort selection

Human intestinal tissue procurement, storage, and use were conducted in accordance with protocols approved by the Institutional Review Boards (IRBs) of the University of Pittsburgh (PRO14070508 and PRO14100537), Washington University in St. Louis (201701032, 201706182, and 201802101), and the University of North Carolina at Chapel Hill (22-3289 and 21-3134). The latter IRB protocol (21-3134) serves as the single IRB for the NEC Biorepository with reliance agreements established with Baylor College of Medicine, Indiana University, Oregon Health and Science University, Weill Cornell Medicine, and University of California at Davis.

Infants were enrolled in the NEC Biorepository at participating neonatal intensive care units (NICUs), following informed consent by a parent or legal guardian. NEC was diagnosed based on clinical and radiographic findings consistent with modified Bell’s staging criteria, with surgical NEC defined as Bell’s stage III requiring operative intervention. The diagnosis of NEC was further confirmed by histopathological examination of the resected intestinal tissue by a clinical Pathologist.

Non-NEC infant surgical control specimens were obtained from infants undergoing intestinal surgery for conditions unrelated to NEC, including non-inflammatory conditions such as gastroschisis, atresia, spontaneous intestinal perforation, ostomy takedown, intestinal obstruction, and imperforate anus. Infant control specimens were included when there was no clinical, operative, or pathologic evidence of NEC or other intestinal inflammatory disease. Archived human fetal intestinal specimens (<24 weeks’ gestation) were obtained from the Health Sciences Tissue Bank at Magee-Womens Hospital, University of Pittsburgh, as part of previously established tissue collections (IRB PRO14100537).

In some cases, de-identified intestinal samples were obtained under IRB-approved waivers of consent from the University of Pittsburgh IRB (PRO14070508 and PRO14100537) and Washington University (201701032). For these specimens, available clinical information was limited to the anatomical location of the resected intestine and the surgical indication. Intestinal specimens were flash-frozen and stored at −80°C until analysis.

Clinical variables collected, when available, included gestational age at birth, postnatal age at tissue collection, sex, intestinal region, and demographic characteristics. Postmenstrual age (PMA) at tissue collection was calculated as gestational age at birth plus postnatal age at the time of surgery. Clinical and demographic characteristics are summarized (Fig. 39). Missing clinical information was recorded as not available and was not imputed for descriptive analyses.

**Fig. 39.**
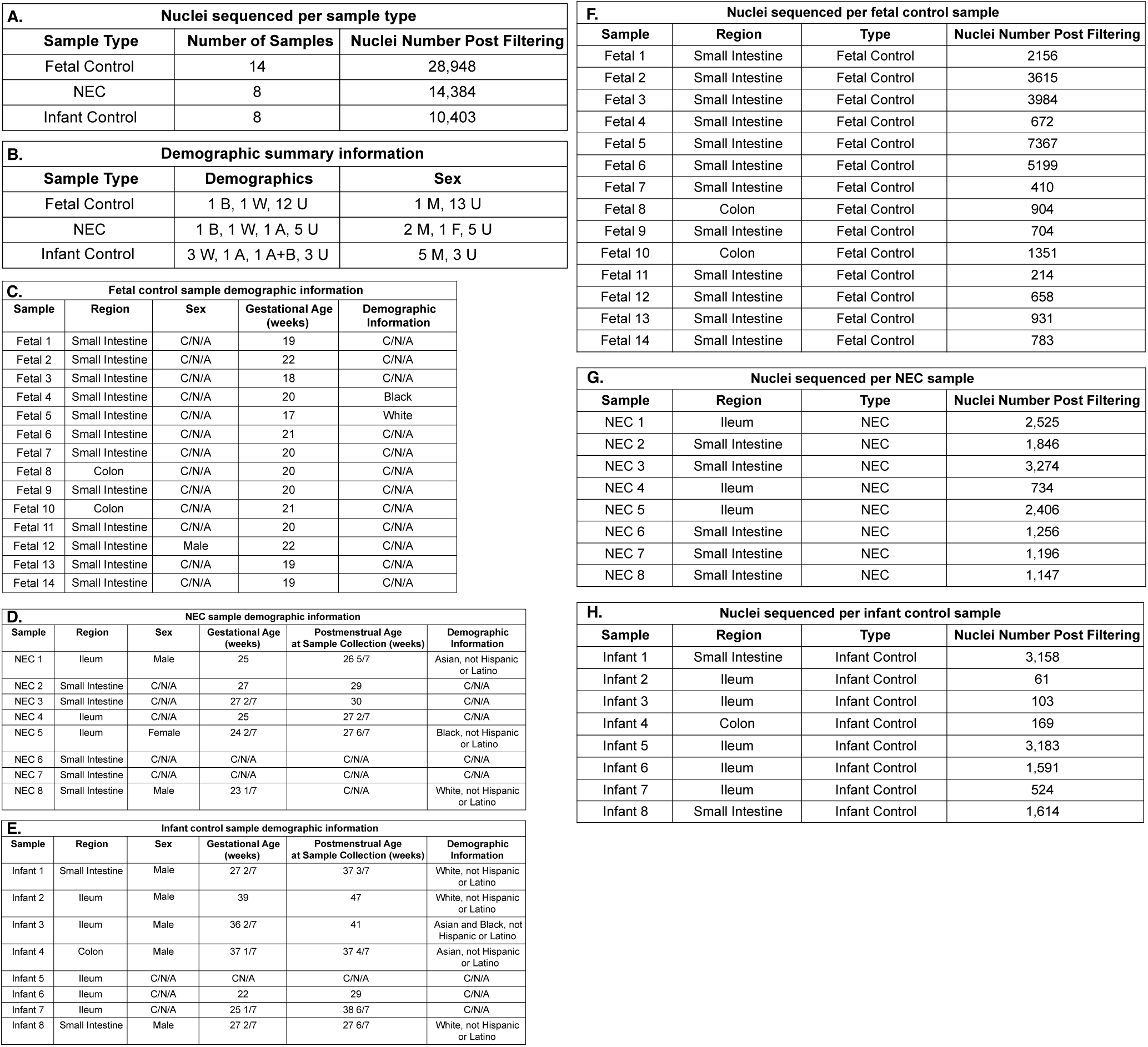
Clinical characteristics and sequencing yield of the study cohort. **A.** Summary of the number of fetal control, NEC, and infant control samples and the number of nuclei retained after filtering for each group. **B.** Summary of available demographic and sex information for the three study groups. **C.** Sample-level region, sex, gestational age, and available demographic information for fetal controls. **D.** Sample-level region, sex, birth age, postmenstrual age at sample collection, and available demographic information for NEC samples. **E.** Sample-level region, sex, birth age, postmenstrual age at sample collection, and available demographic information for infant controls. Postmenstrual age (in weeks) was calculated as gestational age at birth plus postnatal age (time elapsed since birth). **F–H.** Numbers of nuclei retained after filtering for each individual fetal control, NEC, and infant control sample, respectively. A, Asian; B, Black; F, female; M, male; U, unknown or unavailable; W, White. HN, Healed-NEC. C/N/A – clinical information not available.

Full-thickness intestinal specimens were obtained through the multicenter NEC Biorepository, established to facilitate collaborative translational research in NEC.^36–37^ A total of 192 flash-frozen intestinal specimens were available for nuclear isolation, of which 130 specimens underwent sequencing. Following application of pre-specified quality control criteria, 30 specimens met criteria for downstream analysis. The final cohort included 14 fetal control specimens, 8 surgical NEC specimens, and 8 infant control (non-NEC) specimens.

### Nuclei isolation from flash-frozen intestinal biopsies

Nuclei were isolated using the Minute Detergent-Free Single Nuclei Isolation Kit (Invent Biotechnologies, Inc.; NI-024) adapted from the manufacturer’s protocol. In brief, forceps and razor blades were kept on dry ice with a 10 cm Petri dish prior to obtaining samples from the −80°C freezer. Flash-frozen intestinal biopsies (approximate weight: 30 - 100 mg) were placed on the Petri dish on dry ice and cut into 4-5 tissue pieces (each piece approximately 2 mm^3^) with the pre-chilled razor blade, using pre-chilled forceps to steady. These pieces were then loaded onto a filter column in a collection tube on ice. 200 µL of cold buffer A + RNase inhibitor (Millipore; 3335399001) was added to the filter column, and a sterilized plastic rod (supplied in kit) was used to gently grind the tissue onto the filter column for 2 minutes. 300 µL more of cold buffer A + RNase inhibitor was added to the same filter column and incubated on ice for 5-10 minutes with the cap to the collection tube open, then repeated for each sample. After each sample was ground up and cold buffer A + RNase inhibitor was added, the collection tubes with a filter column were capped and inverted a few times to homogenize the mixture, then centrifuged at 15,000 x g for 30 seconds at room temperature. The filter columns were discarded, and the pellet in the flow-through was resuspended with a P1000. Resuspended flow-through fractions were capped and centrifuged at 500 x g for 2-3 minutes at 4°C. The supernatant was discarded, and the pellet was resuspended in 500 µL of cold buffer B + RNase inhibitor, then centrifuged at 600 x g for 8-10 minutes at 4°C. The isolated nuclei in the pellet were fixed using the Evercode Nuclei Fixation v2 (Parse Biosciences - ECF2103) or v3 (Parse Biosciences - ECFN3300) kits, according to the manufacturer’s instructions, and stored at −80°C for less than a month before library preparation.

### Nuclei imaging for correlation scoring

Fixed nuclei were recovered during the Evercode™ Nuclei Fixation v2 or v3 protocol (Parse Biosciences) immediately after section 2.1 step 16/17 (permeabilization stop/neutralization). From the 520 µL fixed nuclei suspension, 50 µL was transferred to a flat-bottom 96-well plate containing 150 µL PBS + DAPI (1:1000 DAPI:PBS) and allowed to settle for 10 min at room temperature before imaging. For each condition, three representative fields were acquired at 40× magnification in matched phase contrast and DAPI channels, using the Keyence BZ-X810 All-in-one Fluorescence Microscope. Nuclear morphology was assessed by direct comparison of the paired images from each field. Individual objects were annotated as: a, intact nuclei; b, nuclei with membrane protrusions; c, non-spherical nuclei; d, debris; and e, objects lacking detectable DAPI signal. Intact nuclei were defined as discrete DAPI-positive nuclei with smooth, compact, approximately round or oval morphology. Nuclei with membrane protrusions were defined as DAPI-positive nuclei containing one or more focal bleb-like extensions from an otherwise contiguous nuclear body. Non-spherical nuclei were defined as DAPI-positive nuclei with globally distorted morphology, including elongated, bent, pinched, lobulated, or otherwise non-ovoid shapes. Debris was defined as irregular non-nuclear material lacking the size, boundary definition, or DAPI staining pattern expected for an intact nucleus, and DAPI-negative objects were defined as phase-visible structures without a corresponding discrete DAPI-positive nucleus (Fig. 40A,B). For each field, the percentage of intact nuclei was calculated relative to all annotated objects, and fields were assigned a nuclei quality score of 0, 1, or 2, corresponding to 0%-15%, 16%-50%, or 51%-100% intact nuclei, respectively. The three field-level scores for each sample were averaged to generate a single sample-level nuclei quality score, which was then used for correlation with transcript recovery metrics. Mean transcripts per nucleus were calculated before filtering. The association between nuclei quality and mean transcript recovery was evaluated in an initial test set of 28 samples, and the nuclei score was positively associated with mean transcripts/nucleus (R² = 0.5404, P value < 0.0001) (Fig. 40C). Simple linear regression was performed using GraphPad Prism, with the reported P value obtained from an F test of whether the regression slope differed from 0. The experimental set comprised 63 samples, with 28 samples used in both the test set and experimental set. A reference of 1,000 mean transcripts per nucleus was used to summarize sequencing yield. Among the 63 experimental preparations, none of the score 0 preparations exceeded 1,000 mean transcripts per nucleus, compared with 50% of score 1 and 71% of score 2 preparations (Fig. 40D). Samples with a nuclei score of 1 or 2 were advanced to library preparation and sequencing. Sequenced samples with fewer than 500 recovered nuclei were excluded from downstream analysis.

**Fig. 40.**
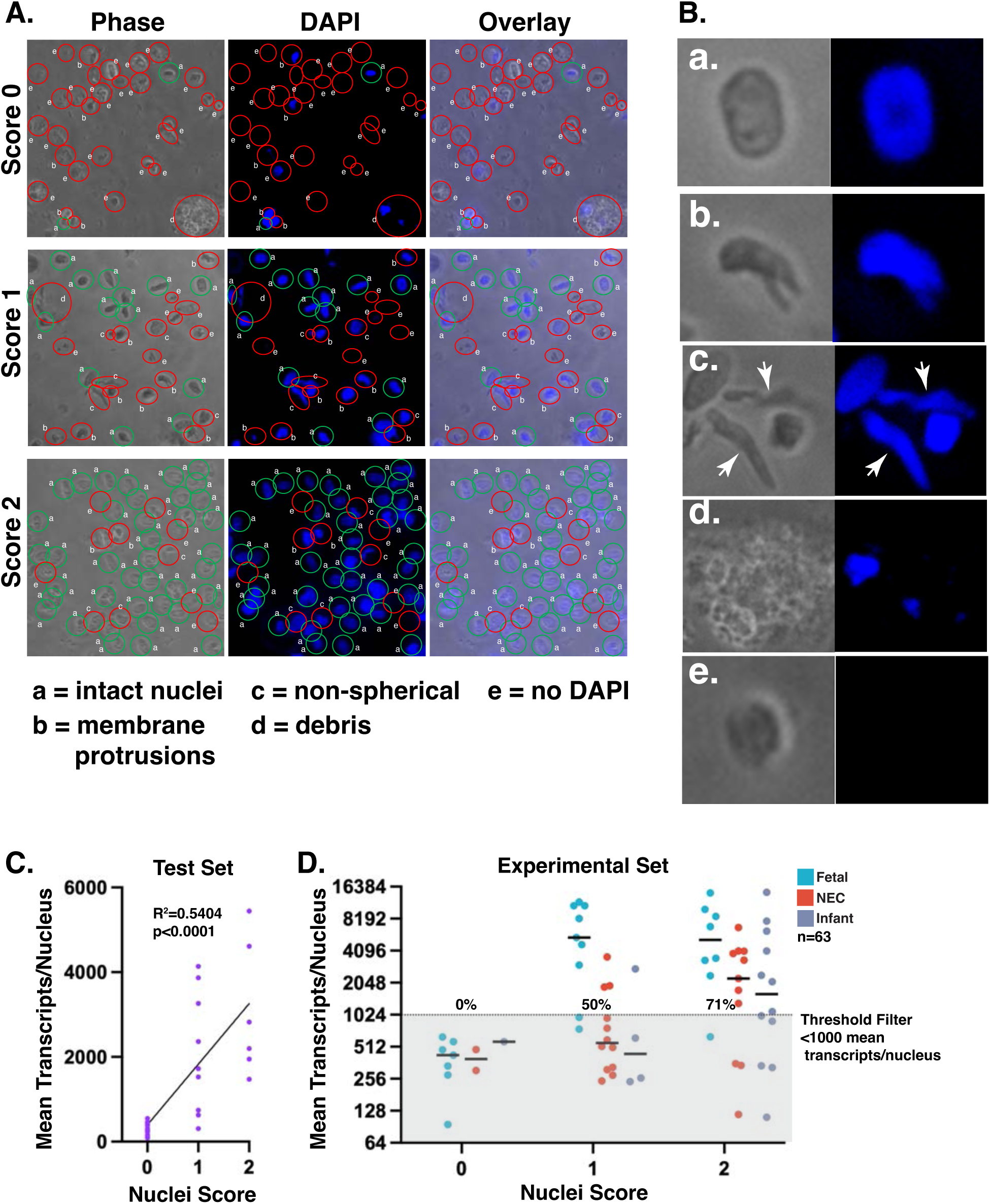
Nuclei morphology scoring predicts transcript recovery and guides sample selection for library preparation and sequencing. **A.** Representative phase-contrast, DAPI, and overlay images illustrating nuclei preparations assigned quality scores of 0, 1, and 2. Green outlines identify intact nuclei, whereas red outlines identify nuclei or image features with quality defects (defected denoted by letter code at bottom). **B.** Representative examples of morphological criteria used during scoring: a, intact nuclei; b, membrane protrusions; c, non-spherical nuclei; d, debris; and e, objects lacking DAPI signal. **C.** Relationship between nuclei score and mean transcripts recovered per nucleus in the initial test set; linear regression yielded R² = 0.5404 and P < 0.0001. **D.** Mean transcript recovery according to nuclei score in the experimental set (n = 63), with points colored by sample group. The dotted line indicates the 1,000-mean-transcripts-per-nucleus threshold used as a quality reference; 0%, 50%, and 71% of score 0, 1, and 2 preparations, respectively, exceeded this threshold.

### Library preparation and sequencing

Following nuclei isolation, fixation, and quality assessment, selected fixed nuclei samples underwent split-pool combinatorial barcoding and library preparation using the Evercode WT Mini v2 (Parse Biosciences; ECW02010 and ECW02110), and Evercode WT Mega v2 (Parse Biosciences; ECW02150) kits, according to the manufacturer’s instructions. Nuclear RNA was reverse-transcribed and labeled through three rounds of split-pool combinatorial barcoding. Following barcoding, nuclei were pooled and divided into sublibraries for lysis, cDNA amplification, and Illumina-compatible library construction.

Three separate library preparation and sequencing experiments were performed using the Mini v2 kits. Fixed nuclei from 8, 12, and 12 samples, respectively, were thawed and processed, with approximately 20,000, 10,000, and 10,000 barcoded nuclei targeted across 12 wells of the Round 1 barcoding plate, corresponding to target numbers of 2,500, 834, and 833 barcoded nuclei per sample, respectively. The resulting libraries were sequenced on an Illumina NextSeq 2000 at the UNC Integrated Genomics Cores (RRID SCR_022620) using a full P3 flow cell for the first experiment, and on an Illumina NovaSeq X at SeqMatic LLC (Fremont, CA) using two lanes of a 10B flow cell for both the second and third experiments. The first experiment used single indexing with a sequencing configuration of 74 cycles for Read 1, 6 cycles for Index 1, and 86 cycles for Read 2. The second and third experiments used dual indexing with 66 cycles for Read 1, 8 cycles for each index read, and 86 cycles for Read 2.

For the Mega v2 kit, fixed nuclei from 45 samples were thawed and processed, with approximately 1,000,000 barcoded nuclei targeted across the 96-well Round 1 barcoding plate and a target of 22,250 barcoded nuclei per sample. The pooled barcoded nuclei were divided into eight sublibraries for lysis, cDNA amplification, and Illumina-compatible library construction. Two of the eight sublibraries were selected for sequencing. These sublibraries were indexed using the Parse UDI WT plate, assessed using an Agilent Bioanalyzer, and submitted to SeqMatic LLC (Fremont, CA) for qPCR-based quality control, normalization, pooling, and sequencing on an Illumina NovaSeq X using all eight lanes of a 25B flow cell with a 300-cycle reagent kit. Sequencing was performed using 66 cycles for Read 1, 8 cycles for each index read, and 86 cycles for Read 2. Raw sequencing data were returned as FASTQ files.

### Data processing and filtering

FASTQ files were aligned to the human genome version GRCh38 using the Parse Biosciences pipeline (v1.4.1), and downstream analysis was performed with scanpy (v1.12).^1602^ Quality filtering thresholds were set on a per-sample basis with an absolute minimum threshold of at least 1,000 transcripts per nucleus. After filtering, read counts were normalized to reads per 4k and log-transformed. Gene expression associated with per-nucleus read depth was regressed out by linear regression, and 5475 highly variable genes were identified with the scanpy function pp.highly_variable_genes (min_mean=0.0125, max_mean=6, min_disp=0.2). Genes were scaled using Z-score normalization (max_value=10), and cell cycle phases were inferred using the scanpy function tl.score_genes_cell_cycle, following published methods.^1603^ Principal component analysis (PCA) of the highly variable gene set was then performed on the scaled data, and Harmony (harmonypy v0.2.0) was used to correct the PCA for batch effects.

### Clustering and doublet removal

An initial clustering run was performed, identifying several clusters characterized by high read counts, co-expression of multiple lineage markers, and a high Scrublet score, which were removed from the dataset as likely doublets. Data processing and clustering were then repeated, calculating the nearest neighbor graph (n_neighbors=20, n_pcs=40), performing Leiden clustering (resolution=0.6), and using PAGA^1604^ to initialize the UMAP dimensional reduction algorithm (min_dist=0.3, maxiter=None, spread=1, gamma=1). Sub-clustering was performed by re-running the Leiden algorithm on a subset of the data and was used to further resolve related cell types grouped by the initial Leiden clustering. This was done to resolve the mesothelial cells and the mixed neurons 2 (sensory-like), as they initially clustered together. B cells and activated B cells were initially separate clusters but had similar expression of canonical marker genes, as well as NEC-versus-infant differentially expressed genes, so the two clusters were merged into one “B cell” cluster.

### Cell identities and tissue compartment heatmaps

Differential gene expression between clusters was calculated using the scanpy function tl.rank_genes_groups, using t-tests followed by a Benjamini-Hochberg multiple-testing correction. DEGs distinguishing each cluster from the remaining clusters were used to assign cell identities by probing the literature and plotting canonical cell type markers (Figure 2B,F). To further characterize cell types within each tissue compartment, DEGs were calculated between each cell type and the remaining cell types within that compartment and plotted on heatmaps if their expression was predominantly specific to the corresponding cell type (Figure 6,14,22,27,31,35).

### Differential gene expression and Gene Ontology (GO) enrichment analysis

Differential gene expression between groups was calculated using the scanpy function tl.rank_genes_groups, using t-tests followed by a Benjamini-Hochberg multiple-testing correction. For GO enrichment analysis, lists of genes significantly (FDR-adjusted P value < 0.05) enriched or depleted in NEC samples relative to infant controls were analyzed using the Python package goatools (v1.5.2), comparing to the full GO basic database (release version 2025-10-10), again correcting for multiple testing with the Benjamini-Hochberg method. GO terms were shown if they were significantly enriched or depleted in NEC compared with infant controls, potentially relevant to NEC or the cell-type-specific disease response, and nonredundant. Closely related terms were generally represented by a single term (e.g., “inflammatory response” [GO:0006954] or “regulation of inflammatory response” [GO:0050727], rather than both).

### Common altered NEC-Associated DEG dot plots

NEC-versus-infant differentially expressed genes across cell types were compared to identify genes commonly altered in NEC. Upregulated (log2 fold change > 0 and FDR-adjusted P value < 0.05) and downregulated (log2 fold change < 0 and FDR-adjusted P value < 0.05) genes were considered separately. For each gene, the number of cell types in which it was significantly altered in the same direction (upregulated or downregulated) was counted. Genes significantly upregulated in at least 16 cell types were ranked by the number of cell types in which they were upregulated, and the top 30 were displayed in a dot plot (Fig. 4A). Genes significantly downregulated in at least 16 cell types were ranked and displayed in the same manner (Fig. 4B). Each dot represents a significant gene-cell-type pair, with dot size indicating the percentage of NEC nuclei within that cell type expressing the gene. If there is no dot present, the gene is not significantly altered in NEC in that cell type, or that the displayed gene was significantly altered in NEC but in less than 5% of NEC cells. For visualization, log2 fold-change values were converted to fold-change magnitudes (2^|log2FC|^) and grouped into discrete bins for color encoding, preventing extreme fold changes from obscuring differences among the remaining genes. Cell types with no significant DEGs in the corresponding direction were excluded from the x-axis ordering (UP: BEST4+ Cells, Dendritic Cells, Interstitial Cells of Cajal, Mesothelial Cells, Mixed Neurons 2 (Sensory-like), Tuft Cells; DOWN: B Cells, BEST4+ Cells, Dendritic Cells, Interstitial Cells of Cajal, Mesothelial Cells, Mixed Neurons 2 (Sensory-like), Tuft Cells). After plotting criteria were applied, any additional cell type with no plotted points among the selected genes was also removed.

### Cell death gene dot plots

The NEC-versus-infant differentially expressed genes calculated above were used and probed for cell death-associated gene expression changes in each cell type in NEC. Common cell death genes were categorized based on their known function in apoptotic, necroptotic, pyroptotic, and ferroptotic processes, as these are forms of cell death that can happen during intestinal disease. Genes involved in one of the cell death processes listed were plotted for a given cell type if they were significantly upregulated (log2 fold change > 0 and FDR-adjusted P value < 0.05) or downregulated (log2 fold change < 0 and FDR-adjusted P value < 0.05) in NEC. Dot color indicates NEC-vs-infant log2 fold change, with orange indicating increased expression in NEC and blue indicating decreased expression in NEC. Dot size indicates the percentage of NEC nuclei expressing each gene, and if there is no dot present, then the gene is not significantly altered in NEC in that cell type. To prevent high log2 fold change values from dominating the visual range, fold changes were displayed using a symmetric logarithmic color scale: values from −3 to +3 log2 fold change were scaled linearly, while values outside this range were log-compressed for visualization only (Fig. 5). Original log2 fold change values are retained in the exported tables.

### Gene categories for interpretability

The NEC-versus-infant control differentially expressed genes calculated above were used as well as NEC-versus-fetal differentially expressed genes to categorize genes for better interpreting biological significance in each cell type (Fig. 3A). Genes with expression in NEC significantly (FDR-adjusted P value < 0.05) higher than both fetal and infant controls or significantly lower than both fetal and infant controls in a given cell type were categorized as “NEC-specific change”, meaning the change in gene expression is more likely due to an active disease state. Genes with expression in NEC significantly different than infant controls, but not significantly different than fetal controls were categorized as “Fetal-like persistence/reversion”, meaning the difference in expression seen in NEC could be due to the premature state of the NEC infant where the expression hasn’t progressed to the proper mature state, or that in NEC this gene reverts to a fetal-like state. Genes with NEC expression significantly different than both infant and fetal, but where NEC expression was between the two controls (fetal<NEC<infant or fetal>NEC>infant) were classified as “Incomplete developmental change”, meaning the difference in expression seen in NEC could be due to a disease state or just due to normal development, where if the premature NEC infant had more time to mature, the NEC expression would match infant control. Genes with NEC expression significantly different than fetal control but not infant control were classified as “Normal developmental change retained”, meaning that even though the NEC infant is premature, this gene appears to reach a mature state of expression matching infant control. Genes with expression in NEC not significantly different than both fetal and infant control were classified as “No major difference”, meaning this gene does not appear to change in expression across developmental stages and it also does not change in NEC. These categories were used to interpret the gene expression changes seen across developmental controls and in disease to hypothesize which genes might be important in NEC pathogenesis.

### Top 30 NEC-specific gene dot plots

Using the gene categories above, the top “NEC-specific change” genes in each cell type were shown to give readers a list of potentially important genes regulating NEC pathogenesis in a given cell type (Fig. 3B). For each annotated cell type, FDR-adjusted P values (< 0.05) and effect sizes (log2 fold change) from the same differential expression pipeline used throughout the atlas were used, with upregulated genes defined as log2 fold change > 0 and downregulated genes defined as log2 fold change < 0. Cell types with no significant genes were excluded from reporting. To avoid sparse, hard-to-interpret signals, genes were further required to be detected in at least 10% of NEC cells (for upregulated genes in NEC) or 10% of infant control cells (for downregulated genes in NEC) within that cell type before they were eligible for ranking. Prior to ranking, features that are frequently significant in single-cell outputs but are not readily interpretable mechanistically were removed, including Ensembl-only identifiers (e.g., ENSG00000… without a mapped symbol), predicted/placeholder transcript-like entries (e.g., uncharacterized open reading frames), predominantly non-coding transcript classes typically used as annotation-depth or transcriptional-noise markers (e.g., LINC*, antisense *AS1/*AS2, and small RNA families such as MIR*, unless there was a strong a priori rationale), and pseudogenes (unless specifically relevant to the biological question). After applying these filters, two ranked outputs in each cell type were produced: (i) the top “NEC-specific change” genes ranked by significance, ordering genes by increasing FDR-adjusted P value (with deterministic tie-breaking by effect size magnitude), and (ii) the top “NEC-specific change” genes ranked by log2 fold change, ordering genes by increasing effect size (with direction retained). The top 30 upregulated genes by significance, the top 30 upregulated genes by log2 fold change, the top 30 downregulated genes by significance, and the top 30 downregulated genes by log2 fold change were plotted into separate dot plots. Dot size indicates the percentage of nuclei expressing each gene, and darker dots indicate higher relative mean expression. For each gene within each cell population, mean expression was normalized to the highest value among the three conditions (FC, NEC, and IC), with the highest assigned a value of 1 and the others scaled proportionally between 0 and 1. Gene names were colored orange if upregulated in NEC (log2 fold change > 0) or colored blue if downregulated in NEC (log2 fold change < 0).

### Curated gene story dot plots

Using the NEC-versus-infant differentially expressed genes identified above for each cell type, the literature was probed for roles of these genes in processes implicated in the initiation, progression, or resolution of NEC. Genes related to the same biological process or broader process category were grouped and displayed in dot plots, with dot size indicating the percentage of nuclei expressing each gene, and darker dots indicating higher relative mean expression. Mean-expression normalization and gene-name colors followed the definitions used for the top 30 NEC-specific gene dot plots described above.

### Cell-type-specific maturity projection & visualization

For each annotated cell type, fetal and infant control cells were used to define a normal fetal-to-infant developmental reference axis. For visualization, genes with zero mean expression in either the fetal or infant control group were excluded, as a spot-check analysis across several cell types indicated that most of these genes were expressed in fewer than 1% of NEC nuclei. These genes were retained in the complete results provided in the supplemental files.

Developmentally regulated genes were then identified by differential expression testing between fetal and infant controls within each cell type, excluding NEC samples. Results from this comparison determined the x and y position of each gene in the volcano plots, with log2 fold change between infant and fetal expression on the x-axis and -log10(FDR-adjusted P value) on the y-axis. Because genes with zero mean expression in fetal or infant controls were removed before differential expression testing, the multiple-testing correction was performed on a different set of genes. Consequently, the FDR-adjusted P values for NEC-versus-infant comparisons here differ slightly from those obtained in the original NEC-versus-infant analysis.

Genes were then evaluated for NEC-associated regulation by comparing NEC with infant controls, and genes with an FDR-adjusted P value < 0.05 in both the fetal-versus-infant and NEC-versus-infant comparisons were colored in the maturity projection plots. For each qualifying gene, NEC expression was normalized using the equation: (NEC mean − fetal mean) / (infant mean − fetal mean) to calculate a developmental projection score. With this scale, a score of 0 indicates that NEC mean expression of that gene is equal to the fetal control mean expression, and 1 indicates that NEC mean expression is equal to the infant control mean expression. Values between 0 and 1 indicate that NEC mean expression is between fetal and infant expression; values below 0 indicate fetal-like dysregulation (either NEC<fetal<infant or NEC>fetal>infant), and values above 1 indicate infant-like dysregulation (either fetal<infant<NEC or fetal>infant>NEC).

Genes were visualized using volcano plots, with the x-axis showing developmental projection score and the y-axis showing NEC-versus-infant significance as -log10(FDR-adjusted P value). Each gene was assigned to a developmental-position category based on where NEC expression fell relative to the fetal and infant means, using the gene categories above: fetal-like, between fetal and infant, beyond fetal, beyond infant, or not significant. To prevent highly significant genes outside the fetal-to-infant range from compressing the fetal-like genes near the bottom of the plot, beyond-fetal and beyond-infant genes more significant than the most significant fetal-like gene were not displayed. The transcription factors used for the developmental projection plots were obtained from a published transcription factor database (v1.01).^1605^

## Supporting information

Universal Cell Type DEGs

Number_Percent_Cells_Dataset

Maturity Volcano Plots Spreadsheets

GOA term enriched depleted

DEGs NEC vs Infant

Common DEGs for Dotplot

Within Lineage Cell Type DEGs

## Transcript Profiling

Data will be deposited in the NCBI Gene Expression Omnibus under accession number GSE351352. Python scripts supporting the major analysis steps will be made available upon request.

## Author Contributions

STM and MG conceptualized the study and provided financial support. MFM, CWS, RJB, GC, and LCF developed the nuclei-isolation protocol and prepared sequencing libraries. MFM, CWS, KB, HMD, and STM performed computational analyses, with KB providing computational, statistical, and bioinformatics support. MFM, CWS, KB, JB, RJB, and STM interpreted sequencing results and designed the analytical framework and data presentation. MFM, CWS, MG, and STM wrote the manuscript; MFM, CWS, KB, and STM wrote the Methods. STM, MFM, CWS, JB, and MG led analytical interpretation and narrative development for epithelial populations; LVL and MFM for enteric nervous system populations; VB, PB, and MFM for endothelial populations; DS and MFM for immune populations; and NM and CWS for stromal and smooth muscle populations. MG, LCF, CRR, DES, ABH, CRM, TAM, and BS provided patient tissue samples and clinical expertise. VP, LCF, LRB, CRR, DES, CMJ, GDM, NSA, and MG provided tissue quality assurance. JC provided the patient-family perspective and collaborated on CZI grant support. PB, NM, DS, VB, and LVL provided tissue-specific expertise and evaluated sequencing findings. All authors read, edited, and approved the manuscript.

## Acknowledgements

We thank the generous patient donors and their families, whose contributions made this study possible. We also thank Nikki McCoy and the UNC Center for Gastrointestinal Biology and Disease Advanced Analytics Core (RRID SCR_015613) for technical support; Henry Taylor, Caroline Hinesley, and members of the Magness laboratory for assistance; and Drs. Heather McCauley, Adam Gracz, Aadra Bhatt, Laura Ruprecht, and the UNC GI Stem Cell Group for useful discussions and feedback. High throughput sequencing was performed by the UNC Integrated Genomics Cores (RRID SCR_022620). We also acknowledge the participating tissue-procurement, neonatology, and pediatric surgical teams at UNC, Texas Children’s Hospital, Indiana University, Oregon Health & Science University, University of Pittsburgh, and Washington University that supported specimen acquisition.

## Generative AI Use

Generative artificial intelligence tools and large language models were used to assist with literature searches, spelling, grammar, formatting, language refinement to harmonize writing styles across sections contributed by multiple authors and streamlining code for selected data visualizations. The authors were solely responsible for the conception, design, execution, analysis, interpretation, and conclusions of the study; reviewed and validated all AI-assisted code and visualizations; verified all manuscript content and citations; and take full responsibility for the accuracy, integrity, and originality of the work.

## List of Abbreviations

Abbreviations used in the manuscript and figures. Gene and protein names, families, and complexes are excluded.

Abbreviation: Full term
A: Asian (demographic table)
AMPA: alpha-amino-3-hydroxy-5-methyl-4-isoxazolepropionic acid
ATP: adenosine triphosphate
B: Black (demographic table)
BH4: tetrahydrobiopterin
C1P: ceramide-1-phosphate
cAMP: cyclic adenosine monophosphate
cGMP: cyclic guanosine monophosphate
CM: circular muscle
CoA: coenzyme A
CpG: cytosine-phosphate-guanine
DAG: diacylglycerol
DAPI: 4′,6-diamidino-2-phenylindole
DEG: differentially expressed gene
DNA: deoxyribonucleic acid
EA: early absorptive
ECM: extracellular matrix
EEC: enteroendocrine cell
ENS: enteric nervous system
ER: endoplasmic reticulum
ERAD: endoplasmic reticulum-associated degradation
F: female (demographic table)
FC: fetal control (sample-group labels); fold change (gene-ranking headings)
FC.Dn: downregulated genes ranked by fold change
FC.Up: upregulated genes ranked by fold change
FDR: false discovery rate
GABA: gamma-aminobutyric acid
GI: gastrointestinal
GO: Gene Ontology
GSH: reduced glutathione
IBD: inflammatory bowel disease
IC: infant control
ICC: interstitial cell of Cajal
ID: identity or identification (tissue and cell-type labels)
IPAN: intrinsic primary afferent neuron
ISC: intestinal stem cell
ISR: integrated stress response
LEC: lymphatic endothelial cell
LM: longitudinal muscle
log2FC: base-2 logarithm of fold change
LP: lamina propria
LPM: lamina propria myocyte
LPS: lipopolysaccharide
M: mucosa (anatomical schematics); male (demographic table)
MA: mature absorptive
MM: muscularis mucosae
MOMP: mitochondrial outer membrane permeabilization
MP: myenteric plexus
mRNA: messenger ribonucleic acid
N: necrotizing enterocolitis sample group
N/S, n.s.: not significant
NAD: nicotinamide adenine dinucleotide
NEC: necrotizing enterocolitis
neg. reg.: negative regulation
NK: natural killer
NMDA: N-methyl-D-aspartate
NO: nitric oxide
nuc: nuclei
PC: Paneth cell
pos. reg.: positive regulation
Prolif.: proliferative
PUFA: polyunsaturated fatty acid
pval_adj / pval-adj: adjusted P value
RNA: ribonucleic acid
RNA-seq: ribonucleic acid sequencing
S: serosa
Sig.Dn: downregulated genes ranked by statistical significance
Sig.Up: upregulated genes ranked by statistical significance
SIP: smooth muscle cells, interstitial cells of Cajal, and platelet-derived growth factor receptor alpha-positive cells
SM: submucosa
SMC: smooth muscle cell
snRNA-seq: single-nucleus ribonucleic acid sequencing
SP: submucosal plexus
symlog: symmetric logarithmic scale
TA: transit-amplifying
TF: transcription factor
U: unknown or unavailable (demographic table)
UMAP: uniform manifold approximation and projection
UNC: University of North Carolina
W: White (demographic table)

